# WWOX P47T loss-of-function mutation induces epilepsy, progressive neuroinflammation, and cerebellar degeneration in mice phenocopying human SCAR12

**DOI:** 10.1101/2022.10.05.510979

**Authors:** Tabish Hussain, Kevin Sanchez, Jennifer Crayton, Dhurjhoti Saha, Collene Jeter, Yue Lu, Martin Abba, Ryan Seo, Jeffrey L Noebels, Laura Fonken, C Marcelo Aldaz

**Author notes:** **Correspondence:** C. Marcelo Aldaz, Department of Epigenetics and Molecular Carcinogenesis, The University of Texas MD Anderson Cancer Center, 1901 East Rd, Houston, TX 77054, USA.

## Abstract

*WWOX* gene loss-of-function (LoF) has been associated with neuropathologies resulting in developmental, epileptic, and ataxic phenotypes of varying severity based on the level of WWOX dysfunction. *WWOX* gene biallelic germline variant p.Pro47Thr (P47T) has been causally associated with a new form of autosomal recessive cerebellar ataxia with epilepsy and intellectual disability (SCAR12). This mutation affects the WW1 protein binding domain of WWOX, impairing its ability to interact with canonical proline-proline-X-tyrosine motifs in partner proteins. We generated a mutant knock-in mouse model of *Wwox* P47T that phenocopies SCAR12. *Wwox^P47T/P47T^* mice displayed epilepsy, profound social behavior and cognition deficits, and poor motor coordination, and unlike KO models that survive only for 1 month, live beyond 1 year of age. These deficits progressed with age, and mice became practically immobile, suggesting severe cerebellar dysfunction. *Wwox^P47T/P47T^* mice exhibited signs of progressive neuroinflammation with elevated astro-microgliosis that increased with age. The cerebellar cortex displayed significantly reduced molecular and granular layer thickness and a strikingly reduced number of Purkinje cells with degenerated dendrites. Transcriptome profiling from various brain regions from these Wwox LoF mice highlighted widespread changes in neuronal and glial pathways, enrichment of bioprocesses related to neuroinflammation and severe cerebellar dysfunction, activation of pathways compatible with compensatory neurogenesis along with major suppression of gene networks associated with excitability, neuronal cell differentiation and brain development. Our results show significant pathobiological effects and potential mechanisms through which WWOX LoF leads to epilepsy, cerebellar neurodegeneration, neuroinflammation, and ataxia. Additionally, the mouse model described here will be a useful tool for the study of WWOX in common neurodegenerative conditions in which it has been identified as a novel risk factor.

## INTRODUCTION

Over the past few years, abundant evidence has established the association of *WWOX* (WW domain-containing oxidoreductase) gene loss-of-function (LoF) with multiple central nervous system (CNS) pathologies [1]. In 2007, Gribaa et al. described a novel early-childhood onset recessive cerebellar ataxia (autosomal recessive spinocerebellar ataxia type 12 [SCAR12]; MIM:614322) associated with generalized tonic-clonic epilepsy developing at an early age (9-12 months) and severe intellectual disability in four siblings from a large consanguineous family [2]. Cerebellar ataxia and psychomotor retardation were noticed when affected children started to walk, which was delayed until the age of 2–3 years. These children also suffered from severe dysarthria, nystagmus, diminished reflexes, and displayed mild cerebellar atrophy as per brain magnetic resonance imaging (MRI). Using homozygosity mapping, the defective locus was mapped to a 19-Mb chromosomal region at ch16q21-q23 [2]. In 2014, by whole-exome sequencing, it was determined that siblings from the first family with SCAR12 harbored a homozygous missense mutation in the *WWOX* gene. The novel p.Pro47Thr germline mutation affected a highly conserved proline residue which is critical for the function of the first WW domain (WW1, protein-protein interaction domain) of the WWOX protein [3, 4]. We demonstrated that the p.Pro47Thr mutation renders WWOX unable to bind canonical proline-proline-X-tyrosine (PPxY) or similar motifs in interacting partner proteins, thus leading to a partial WWOX LoF, since expression of the protein remains intact and its enzymatic domain is unaffected [4].

After the initial SCAR12 report, multiple other familial cases bearing biallelic germline *WWOX* pathogenic variants leading to loss of function were identified, resulting in developmental, epileptic, and ataxic phenotypes of varying severity [5–9]. The most severe cases of WWOX complete LoF (i.e., null genotype) are now classified as Developmental and Epileptic Encephalopathy 28 (DEE28, MIM:616211), also known as WOREE syndrome (WWOX-related epileptic encephalopathy). These patients display refractory epilepsy, profoundly impaired psychomotor development, spastic quadriplegia, brain structural defects of varying degrees, microcephaly, ophthalmic abnormalities, reduced myelination, and commonly death within the first two years of life (Reviewed in [1, 9]).

Importantly, recent large-scale genome-wide association meta-analyses and biomarker studies have also identified *WWOX* as a risk gene for common neurodegenerative conditions such as Alzheimer’s disease, Parkinson’s disease, and multiple sclerosis [10–18]. The latter two pathologies include ataxia and impaired motor coordination, among significant clinical manifestations. Taken together, WWOX dysfunction is implicated in severe neurodevelopmental abnormalities, and it is a risk gene for a heterogeneous array of neurodegenerative conditions. However, it is largely unknown how total or partial *WWOX* LoF leads to these complex neurological conditions. Thus, understanding the role of WWOX in conditions associated with ataxia and neurodegeneration is important not only for the cases with germline mutation but potentially in more common neurodegenerative pathologies in which *WWOX* has been identified as a risk gene.

In this study, we aimed to determine the pathophysiological effects of WWOX LoF using a novel knock-in mutant mouse model carrying the same germline p.Pro47Thr mutation causally associated with SCAR12 in humans. We observed that *Wwox^P47T/P47T^* mice (i.e., *Wwox* P47T homozygous mutant mice) phenocopied the complex human SCAR12 neurological phenotypes. Like SCAR12 patients, these mice reached adulthood and displayed intense seizure activity, profound behavioral and cognitive deficits, and impaired motor function among other ataxia-related phenotypes. Characterization of cellular and molecular changes in brains from *Wwox^P47T/P47T^* mice revealed signs of significant neuroinflammation with progressive astro-microgliosis (i.e., increasing in severity with mouse age). Neuroinflammatory changes were further confirmed by transcriptome profiling analyses from various brain regions. Histomorphological analyses of the cerebellum (CB) indicated significant atrophy with reduced molecular and granular layer thickness and Purkinje cells (PC) degeneration in the cerebellar cortex of *Wwox^P47T/P47T^* mice. Our studies demonstrate for the first time that a single amino acid change impairing the protein-protein interactive properties of Wwox is sufficient to induce an array of CNS abnormalities that include epilepsy, widespread neuroinflammation and cerebellar neurodegeneration.

## RESULTS

### Generation of *Wwox^P47T/P47T^* mutant mice

We utilized CRISPR/Cas9 technology to generate mice carrying a constitutive knock-in mutation to replace the highly conserved proline residue at the 47 position of Wwox into threonine, thus, reproducing the exact same loss-of-function germline mutation found in SCAR12 patients. To this end, we used a single-strand oligonucleotide donor (SSOD) sequence of 145 nucleotides in length, agcagtgttaacttactctgttgtgggtctctattaca<u>gtcacactgaggagaagacccagtgggaacat**acc**aaaaccggcaagaggaaa cgggtcgcaggag</u>gtctgtatgccgtcccaagcagagaagcattaagtagctag, where the underlined sequence highlights exon 2 encoding nucleotides and shown in bold is codon ACC (Thr) (c.139C>A, c.141G>C) for replacement of codon CCG (Pro) (Supplementary Fig. S1). Wwox P47 founder lines were generated and genotyping performed as described in the Methods. After primary characterization, only one founder line was utilized for obtaining experimental mice used in these studies. Breeding of *Wwox^P47T/WT^* (i.e., heterozygous mice), resulted in viable pups having genotypes with the expected Mendelian ratios demonstrating that the knock-in mutation had no effect on embryonic development or viability (Supplementary Table 1).

As previously mentioned, the P47T mutation lies in the first WW domain of Wwox and renders this protein unable to bind the minimal core consensus PPxY motif in interacting partner proteins [4]. Thus, to evaluate the effects of the knocked-in P47T mutation on Wwox’s binding ability, we performed a comparative peptide pull-down assays using ligand biotinylated oligopeptides from the Dishevelled segment polarity protein 2 (Dvl2) and WW domain binding protein 1 (Wbp1). Both proteins are typical WW group I binders containing Pro-Pro-Pro-Tyr (PPPY) motifs [19, 20]. Protein pull-downs were performed on CB lysates from *Wwox^WT/WT^* (i.e., wildtype mice) and *Wwox^P47T/P47T^* mice. As shown in Figure 1a, Wwox protein from the *Wwox^WT/WT^* mice was successfully pulled down, whereas the *Wwox* P47T version from the *Wwox^P47T/P47T^* mice failed to interact with the PPPY containing Dvl2 and Wbp1 oligopeptides, confirming an efficient mutation and loss of the protein-protein interactive properties of the WW domain of Wwox. Therefore, the generated mouse model is a useful tool for analyzing the effects of Wwox WW domain LoF. To determine whether the P47T mutation affected the *Wwox* gene and protein expression, we measured mRNA and protein levels in CB samples. The average mRNA expression levels between *Wwox^WT/WT^* and *Wwox^P47T/P47T^* CB were comparable, with no significant difference in n=6 mice tested in each group (Supplementary Fig. S2a). Western blot analyses also showed similar levels of Wwox protein expression in homozygous mutant and wildtype mice (see Fig. 1a, Wwox 10% input panel). Thus, these data suggest that the P47T knock-in mutation does not significantly impact *Wwox* expression in mice reproducing observations in human samples from SCAR12 patients [4].

**Figure 1.**
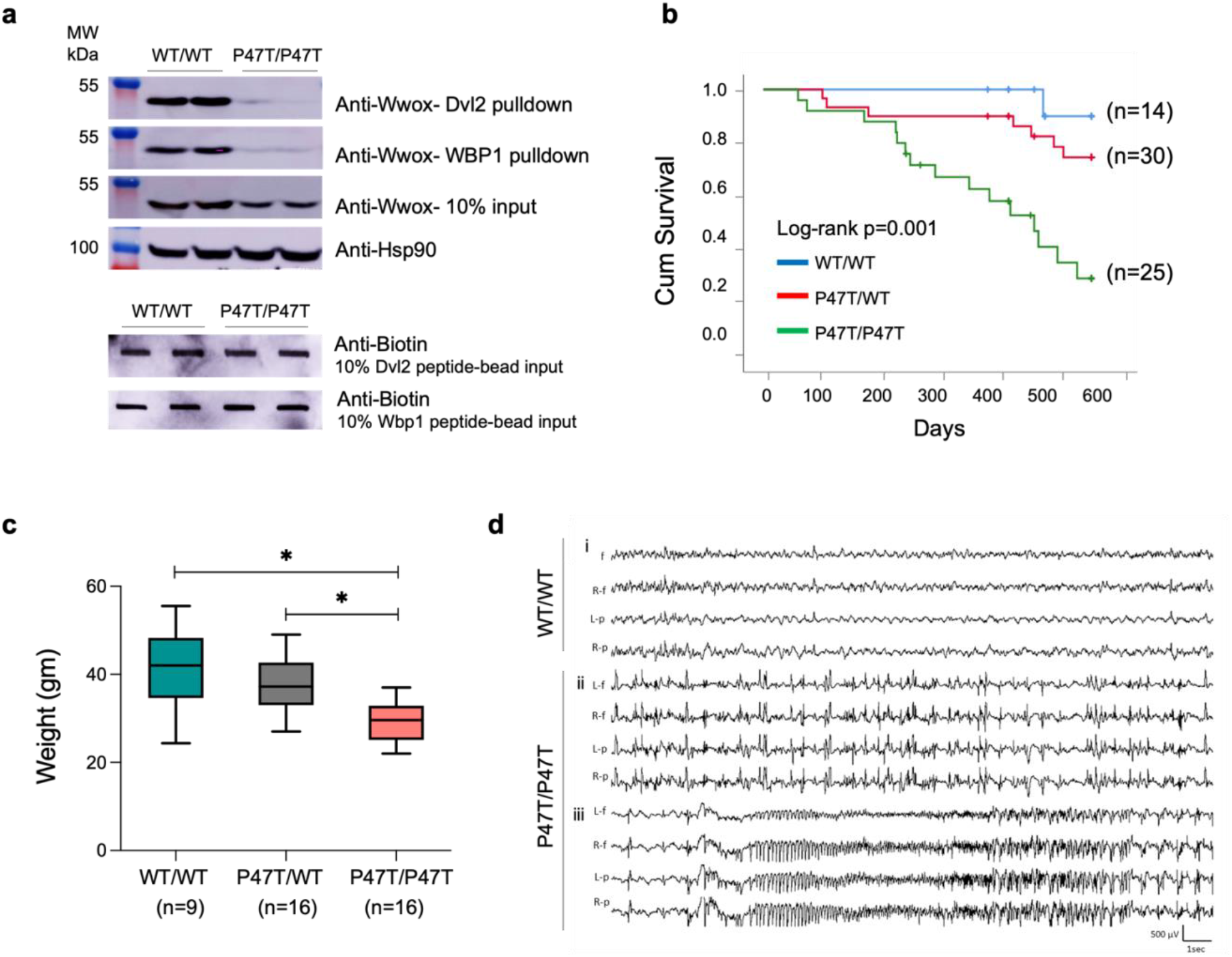
p.Pro47Thr mutation is sufficient to abrogate Wwox affinity for PPxY motifs. (a) Western blots showing the amount of Wwox pulled down from *Wwox^WT/WT^* (n=2) and *Wwox^P47T/P47T^* (n=2) CB tissue lysates using biotinylated Dvl2 (HPYSPQ**<u>PPPY</u>**HELSSY) (top panel) and Wbp1 (SGSGGTP**<u>PPPY</u>**TVG) (second panel) peptides. Third panel shows Wwox expression using 10% input in *Wwox^WT/WT^* and *Wwox^P47T/P47T^* CB samples. HSP90 was used as control to demonstrate equal amount of protein used for pull-down experiment. Two lowermost panels show slot-blots of 10% biotin labeled peptide-bead input to show incubation with equal amounts of peptides. **(b-d) *Wwox^P47T/P47T^* mice display reduced overall survival, hyperactive cortical spike discharges, and generalized spontaneous seizures.** (b) Kaplan-Meier comparative survival in days of a cohort of *Wwox^WT/WT^* wildtype (blue line, n=14), *Wwox^P47T/WT^* heterozygous (red line, n=30), and *Wwox^P47T/P47T^* homozygous (green line, n=25) mice. The *Wwox^P47T/P47T^* mice show significantly lower survival than the *Wwox^WT/WT^* and *Wwox^P47T/WT^* mice, p-value=0.001, Log-rank (Mantel-Cox) analysis (c) Box and Whisker plot showing total body weight in grams of *Wwox^WT/WT^* (n=9), *Wwox^P47T/WT^* (n=16), and *Wwox^P47T/P47T^* (n=16) mice. Box extends from 25^th^ to 75^th^ percentile, with horizontal line at median (50^th^ percentile), whiskers represent bottom and top 25% and extend down to the smallest and up to the largest value, *p-value < 0.005, One-Way ANOVA (Tukey’s post-hoc test). (d) Video-EEG recording profiles showing seizure activity in representative *Wwox^WT/WT^* and *Wwox^P47T/P47T^* mice from a group of n=3 mice/group. i. No abnormal epileptiform activity was detected in EEG from adult *Wwox^WT/WT^* littermates, ii. abnormal bilateral interictal spike discharges were frequent in *Wwox^P47T/P47T^* mutant mice, iii. generalized seizure discharge in the same mouse as ii. shows abrupt spike discharge followed by onset of fast rhythmic, and then arrhythmic patterns of cortical seizure. Traces: L-f: left frontal, R-f: right frontal, L-p: left parietal, R-p: right parietal.

### Effect of germline *Wwox* P47T mutation on survival and morbidity

We conducted a long-term survival study comparing mice from all three genotypes (i.e., *Wwox^WT/WT^, Wwox^P47T/WT^*, and *Wwox^P47T/P47T^*). In Figure 1b, the comparative overall long-term survival is plotted using the Kaplan-Meier method and analyzed by the Log-rank test. In line with observations in humans, where SCAR12 patients homozygous for the P47T mutation reached adulthood [2, 4], *Wwox^P47T/P47T^* mice showed a mean long-term survival of 393 days (±32ds.). However, their life span was shorter compared to 495 (±23ds) for *Wwox^P47T/WT^* and 542 (±8ds) for *Wwox^WT/WT^* mice. The difference in survival was statistically significant between the *Wwox^P47T/P47T^* and the *Wwox^WT/WT^* and *Wwox^P47T/WT^* groups (Log-rank p-value=0.001) (Fig. 1b). *Wwox^P47T/WT^* mice also displayed a reduced overall survival in comparison to the *Wwox^WT/WT^*, but the difference was not statistically significance (Fig. 1b). Throughout the observation period animals were monitored for signs of morbidity and euthanized when moribund. Mice were necropsied and histology samples were collected either at the end of the experimental period or after euthanasia due to poor health. Animal weight was monitored, and blood samples were collected for hematology and laboratory chemistry analyses. In addition to shorter survival, homozygous mutant mice showed a significant decrease in overall weight. Figure 1c displays the comparative mean weight of mice from the various genotypes with *Wwox^P47T/P47T^* mice at 29.22 (±4.28 gm), which was significantly lower than the mean weight 37.39 (±6.08 gm) of *Wwox^P47T/WT^* (p-value < 0.005) and 41.17 (±9.52 gm) of *Wwox^WT/WT^* mice (p-value < 0.005). At necropsy, no significant macroscopic organ abnormalities were observed. Relative organ weights (weight coefficients) were determined, and a slight increase in the kidney and liver weight coefficients was observed in the *Wwox^P47T/P47T^* group compared to the *Wwox^WT/WT^* (p-value < 0.05) (Supplementary Fig. S2). Blood chemistry and hematology parameters were also comparable with no significant differences between the three groups (Supplementary Tables 2 and 3).

Tissue histology showed that several *Wwox^P47T/WT^* (33%) and *Wwox^WT/WT^* (29%) mice developed neoplasia (mostly lung adenocarcinomas and lymphomas) and nephropathy as typically observed in aging mice (Supplementary Table 4). The lower incidence of neoplasia observed in the *Wwox^P47T/P47T^* mice (5%) is likely the consequence of the shorter lifespan for this group. Interestingly, and among the only other remarkable histopathological differences between groups, it was observed that various *Wwox^P47T/P47T^* and *Wwox^P47T/WT^* mice displayed signs of cholestasis with enlarged gallbladders, and a few animals had evidence of focal liver degeneration (Supplementary Table 4).

### *Wwox^P47T/P47T^* mice display chronic spontaneous generalized tonic-clonic seizures

We observed that *Wwox^P47T/P47T^* mice displayed signs of spontaneous seizures, occasionally beginning with wild running and jumping, which often progressed to contraction of limbs and repeated jerking movements followed by lethargy. To evaluate seizure and epileptic activity, we performed video EEG and monitored brain recordings. We examined *Wwox^WT/WT^* (n=3) and *Wwox^P47T/P47T^* (n=3) mice over prolonged periods (21-53 hours) and observed that adult homozygous mutant mice displayed significant seizure activity (Fig. 1c). Wildtype mice displayed infrequent spike activity (0-68/hour) but no evidence of seizures (Fig. 1ci, Supplementary Figs. S3a and b). We detected very frequent interictal cortical spike discharges (0-3098/hour) (Fig. 1cii) and repeated spontaneous generalized convulsive activity (11-22 seizures during 43-53 hour monitoring periods) in both male and female homozygous mutant mice. All three *Wwox^P47T/P47T^* mutant mice had multiple (>4) seizures during a 24 hour recording period (Supplementary Figs. S3a and b). The behavioral seizures were generalized tonic-clonic convulsive episodes, beginning with a sudden myoclonic jerk or jump, followed by whole-body rhythmic jerking movements. These seizures were accompanied by bilaterally symmetrical EEG discharges (Fig. 1ciii), which began with a single spike discharge, rapidly followed by complex hypersynchronous activity consisting of rapid spike patterns that evolved from regular to irregular activity before slowly terminating. Motor contractions began at a high frequency and progressively decreased in parallel with the spike discharge frequency. These seizures were relatively prolonged (up to 1 minute in duration) and were followed by postictal lethargy lasting up to 3 minutes, but were not lethal.

### *Wwox^P47T/P47T^* mice display severe deficits in social and motor behaviors

Since SCAR12 patients display severe intellectual disability and ataxia, we performed experiments to test social and motor behavior. Hind limb clasping is an indicator of disease occurrence and progression in several neurodegenerative mouse models, including cerebellar ataxias [21]. While *Wwox^WT/WT^* and *Wwox^P47T/WT^* mice exhibited normal reaction and their hindlimbs were consistently splayed outward and away from the abdomen, *Wwox^P47T/P47T^* showed abnormal hind limb clasping reflexes, where the hindlimbs were entirely retracted and touching the abdomen (Supplementary Fig. S4a).

In order to evaluate abnormalities in affective behaviors, we assessed performance in the open field test (OFT) and elevated plus maze (EPM). Mouse behavior was determined as the amount of time spent in the brightly lit, open space of the center zone in the OFT arena or of the open arms in the EPM. *Wwox^P47T/P47T^* mice exhibited a marked increase in anxiety-like behaviors in both behavioral tasks. The homozygous mutant mice spent significantly less time in the open areas compared to the wildtype mice, indicative of an anxiety-like phenotype, while the heterozygous mice displayed an intermediate behavioral phenotype (Figs. 2a and b, Supplementary Figs. S4b and c). Additionally, the *Wwox^P47T/P47T^* mice showed blunted locomotor activity in OFT and EPM and traveled less distance in OFT and on average took more than twice as long to first enter the open arms in EPM compared to *Wwox^WT/WT^* (Fig. 2a, Supplementary Figs. S4c). Mice are thigmotaxic in that they prefer to be in an enclosed space where their whiskers contact walls. However, spending more amount of time along the walled perimeter can also indicate elevated anxiety-like behaviors. We observed that *Wwox^P47T/P47T^* mice spent significantly more time in the closed arm of EPM and less time in open arms and the maze center zone than their wildtype counterparts (Fig. 2b).

**Figure 2.**
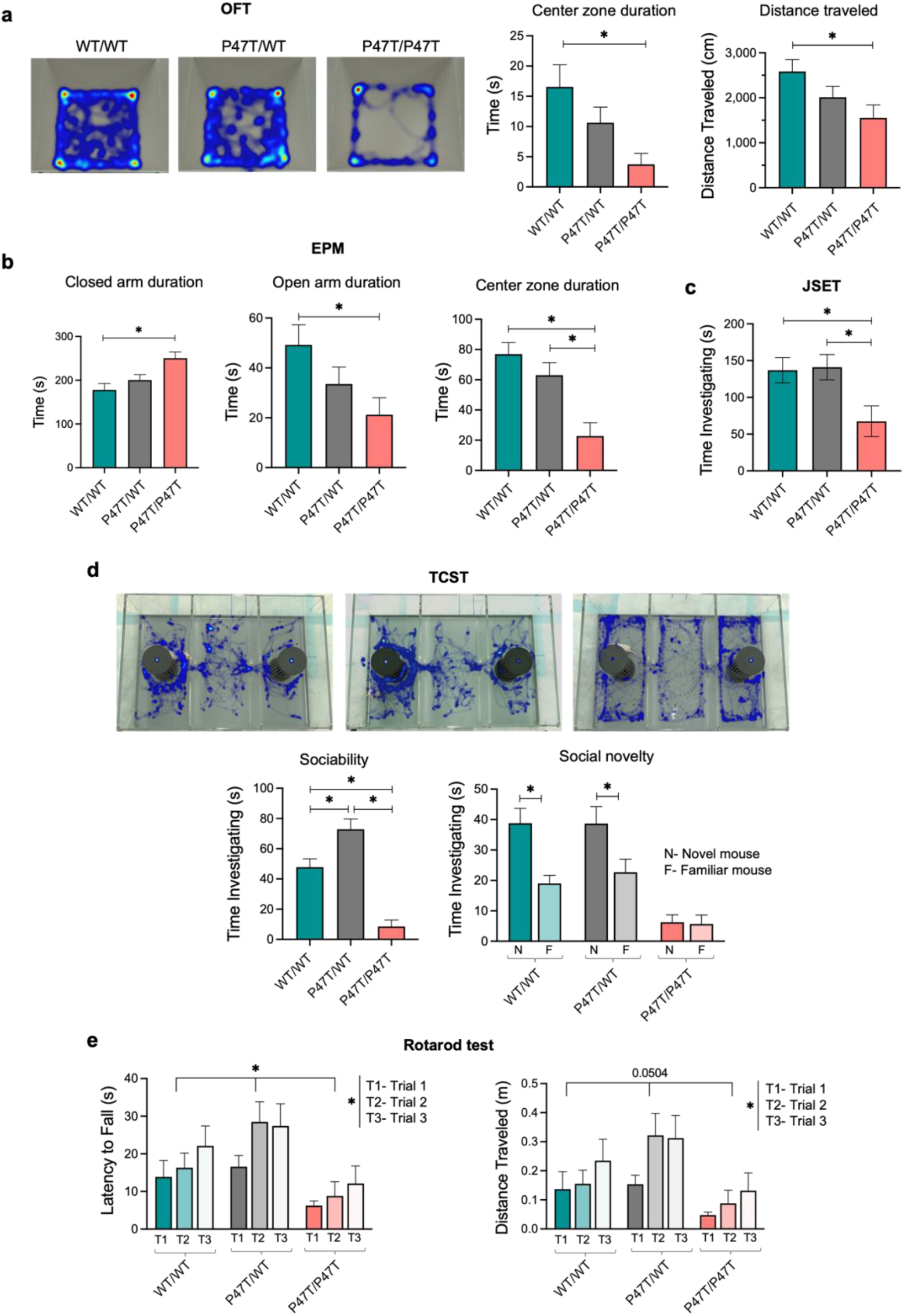
*Wwox^P47T/P47T^* mice display elevated anxiety-like behaviors, reduced sociability, and severely impaired motor function. (a) Representative images of Ethovision video tracking heatmaps depicting the time spent at different locations within the OFT arena over a 5 minute testing duration. Adjacent bar graphs show the quantitative measurement of time spent by test mice in the center zone of OFT arena and total distance traveled during the test duration. (b) Bar graphs showing quantitative measurement of time spent by test mice in closed arm, open arm, and center zone of EPM over a 5 minute testing duration. (c) Mice that underwent OFT were immediately put to JSET using sex-matched 4 weeks old juvenile mouse in the OFT arena. Bar graph showing the quantitative measurement of time spent by test mice investigating the juvenile mouse. (d) Mice underwent TCST that was divided into three 10 minutes sessions of habituation, sociability, and social novelty. Representative images of Ethovision video tracking heatmaps depicting the time spent by the test mouse in different chambers of TCST arena. Heatmaps show the social novelty phase of TCST, where both cages contain sex-matched 8 weeks old stimulus mouse. In the given heatmaps, the left chamber cage contain novel stimulus mouse and the right chamber cage contains familiar stimulus mouse. Bar graphs below the heatmaps show quantitative measurement of sociability measured by time investigating the first stimulus mouse (while the other chamber cage remained empty) and social novelty measured by time investigating the novel stimulus mouse in comparison to the familiar stimulus mouse. (e) Bar graphs showing quantitative measurement of latency to fall and distance traveled by mice in accelerating rotarod test. Mice were placed on the rotarod at 4 revolutions per minute (rpm) with an acceleration rate of 20 rpm/min. Rpm automatically increased as long as the mice remained running. Data was recorded for three trials per mouse with a 15 minute break in between trials. All behavioral tests were performed using *Wwox^WT/WT^* (n=19), *Wwox^P47T/WT^* (n=17), and *Wwox^P47T/P47T^* (n=16) mice and data are represented as mean ± SEM. Rotarod test was statically analyzed using Two-Way ANOVA, other behavioral test were statically analyzed using One-Way ANOVA (Tukey’s post-hoc test), *p-value < 0.05.

The three-chambered sociability test (TCST) assesses social cognition in the form of general sociability and interest in social novelty. This test is useful in quantifying deficits in social behavior in rodent models of CNS disorders. Mice typically prefer to spend more time with another mouse (i.e., sociability) and will investigate a novel mouse more so than a familiar one (i.e., social novelty). Juvenile social exploration (JSET) is another test that measures direct social interaction in mice. We observed that while *Wwox^WT/WT^* and *Wwox^P47T/WT^* mice spent similar amounts of time investigating a juvenile mouse during the JSET, there was a significantly lowered investigation time with the *Wwox^P47T/P47T^* mice (Fig. 2c). A similar pattern emerged during the TCST. During the sociability phase of this test, the wildtype and heterozygous mice had a clear preference for spending time with another mouse versus an empty cage. However, this preference was not seen with the homozygous mutant mice (Fig. 2d). Further, in the social novelty phase of the TCST *Wwox^WT/WT^* and *Wwox^P47T/WT^* mice investigated a novel mouse more in comparison to the familiar one; however, the *Wwox^P47T/P47T^* mice displayed no clear preference (Fig. 2d).

Assessment of sensorimotor coordination and motor learning was primarily done by rotarod test. Mice are naturally inclined to stay on the rotating cylinder to avoid falling. The length of time that a mouse spends on this accelerating rod reflects maturity of balance, coordination, and motor planning. All mice, regardless of genotype, took longer to fall and traversed a longer distance between the first and third trials of the rotarod test (Fig. 2e). However, *Wwox^P47T/P47T^* mutant mice took less time to fall off the rotarod compared to the *Wwox^WT/WT^* mice and, therefore, also had a lower distance traveled (Fig. 2e). These data indicate that *Wwox^P47T/P47T^* mice have impaired motor skills, which is also supported by evidence from the OFT and TCST, where reduced locomotor activity was observed in the same mice. Furthermore, and as mentioned in the survival studies, we observed that mice progressively lose motor activity with age to the point of almost total immobility and deteriorated health.

### *Wwox^P47T/P47T^* brains display progressive microgliosis and astrogliosis

Regardless of the cause, a common feature of neurodegenerative disorders is chronic immune activation [22]. It involves gliosis, a reactive inflammatory response causing hypertrophy and activation of different glial cells, including microglia and astrocytes [23]. In the normal physiological state, microglial cells are highly ramified with a small soma and fine, long processes, allowing constant surveillance of the brain microenvironment [24]. Upon activation by injury, infection, or neurodegenerative stimuli, ramified microglia initially transform into a pro-inflammatory state, with a swollen cell body and shorter thick processes [24]. We evaluated evidence of neuroinflammation by comparative confocal microscopy of microglial morphology (i.e., microgliosis) using immunohistochemical staining for ionized calcium binding adaptor molecule-1 (Iba1) in hippocampi subfields from *Wwox^WT/WT^* (n=3) and *Wwox^P47T/P47T^* (n=3) mice. In order to evaluate evidence of progressive inflammatory changes with aging, we analyzed the brains of mice at 80 and 250 days of age. In both age groups, brains from *Wwox^P47T/P47T^* mice displayed a significant increase in Iba1+ microglial cells in all hippocampal subfields (Fig. 3a and Supplementary Fig. S5a). The difference in the average microglial cell numbers for homozygous mutant mice in all three investigated hippocampal regions (i.e., CA1, CA3, and DG) was significantly higher compared to the wildtype mice at both age groups (p-value < 0.005) (Fig. 3b). A similar trend was observed for the area fraction (AF) occupied by Iba1+ cells, where hippocampi from *Wwox^P47T/P47T^* mice showed significantly higher AF in all inspected brain regions, at 80-days and 250-days (p-value < 0.005) (Fig. 3c).

**Figure 3.**
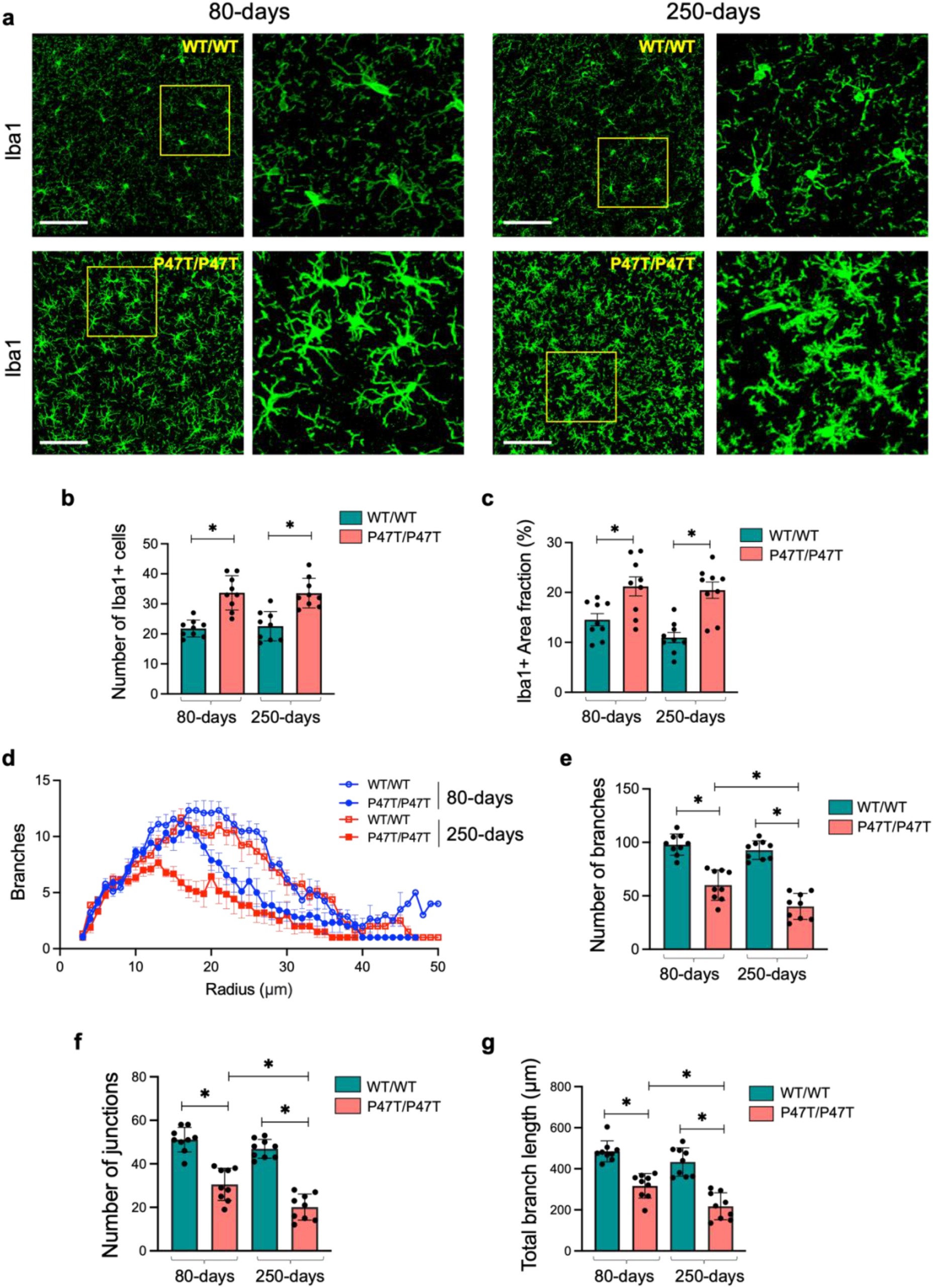
*Wwox^P47T/P47T^* hippocampi exhibit progressive microgliosis. (a) Representative high magnification confocal images illustrating comparative abundance and morphology of Iba1+ microglia in 425 μm^2^ coronal section of HPC CA3 region in *Wwox^WT/WT^* and *Wwox^P47T/P47T^* mice at 80 and 250 days of age as indicated. Yellow square shows the region selected and zoomed to show more intricate morphology as displayed adjacent to each image. Bar graphs show the average number of Iba1+ microglia (b) and Iba1+ area fraction % (c) in the CA1, CA3, and DG subfields taken together for *Wwox^WT/WT^* and *Wwox^P47T/P47T^* in 80 and 250-days groups. Each data point represents quantitation from a single subfield of n=3 mice/group. (d) Sholl analysis comparing number of branches vs. radius from microglia soma measured using representative cells from CA1, CA3, and DG subfields of each mouse. Line graph shows data from *Wwox^WT/WT^* 80-days (hollow blue circles), *Wwox^P47T/P47T^* 80-days (solid blue circles), *Wwox^WT/WT^* 250-days (hollow red squares), and *Wwox^P47T/P47T^* 250-days (solid red squares). Each data point represents average of 9 measurements from n=3 mice/group. Skeleton analysis comparing the number of branches (e), number of junctions (f), and total branch length (g) between *Wwox^WT/WT^* and *Wwox^P47T/P47T^* samples at 80 and 250 days of age as indicated. Each data point represents quantitation from the same microglial cells analyzed for Sholl analysis from CA1, CA3, and DG subfields of n=3 mice/group. Scale bar=100 μm, bar graph data is represented as mean ± SEM, *p-value < 0.005, unpaired Student’s *t*-test.

Microglia were also evaluated for evidence of a more inflammatory phenotype by quantitating microglial cell branching and complexity in representative cells from all hippocampal subfields of n=3 independent mice of each genotype subgroup. Using Sholl analysis, we created concentric circles originating from the microglial soma and subsequently analyzed microglial branching across all distances from the soma. In all hippocampal regions, *Wwox^P47T/P47T^* mice showed reduced branching in both age groups (Fig. 3d). Interestingly, homozygous mutant mice from the 250-days age group showed far more condensed branching compared to the younger (80-days) homozygous mice (Fig. 3d, solid red squares vs. solid blue circles). Additionally, the overall branching at the different concentric radius positions was comparable for the *Wwox^WT/WT^* mice in both age groups (Fig. 3d, hollow red squares vs. hollow blue circles). To complement these observations, after the Sholl analysis, we skeletonized the reconstructed microglia by using the Skeletonize3D plug-in on FIJI to compare the number of branches, junctions, and total branch length. In parallel with the results from the Sholl analysis, *Wwox^P47T/P47T^* mice from both age groups had a reduced number of branches and junctions and an overall lower total branch length (Figs. 3e-g). Similar to observations with Sholl analysis, the differences in the number of branches, junctions, and total branch length were more pronounced and significantly lower in the older homozygous mice (250-days) compared to homozygous mice at 80-days age (p-value < 0.005) (Figs. 3e-g).

We also investigated astrocyte activation (astrogliosis) using glial fibrillary acidic protein (Gfap) immunostaining and quantitated the density and AF occupied by the soma and processes of Gfap+ astrocytes. Similar to the observations of microglial cells, image analysis of Gfap immunostained hippocampal DG, CA1, and CA3 regions from the *Wwox^P47T/P47T^* revealed significantly elevated astrogliosis in both age groups (Figs. 4a and Supplementary Fig. S5b). The average number (Fig. 4b) and AF occupied by Gfap+ astrocytes (Fig. 4c) were significantly higher in the hippocampi of *Wwox^P47T/P47T^* mice (p-value < 0.01 for density and AF). Similar to observations with microglia activation, the level of astrogliosis was significantly higher in the 250-days *Wwox^P47T/P47T^* mice compared to the 80-days *Wwox^P47T/P47T^* group (p-value < 0.005, Figs. 4b and c).

**Figure 4.**
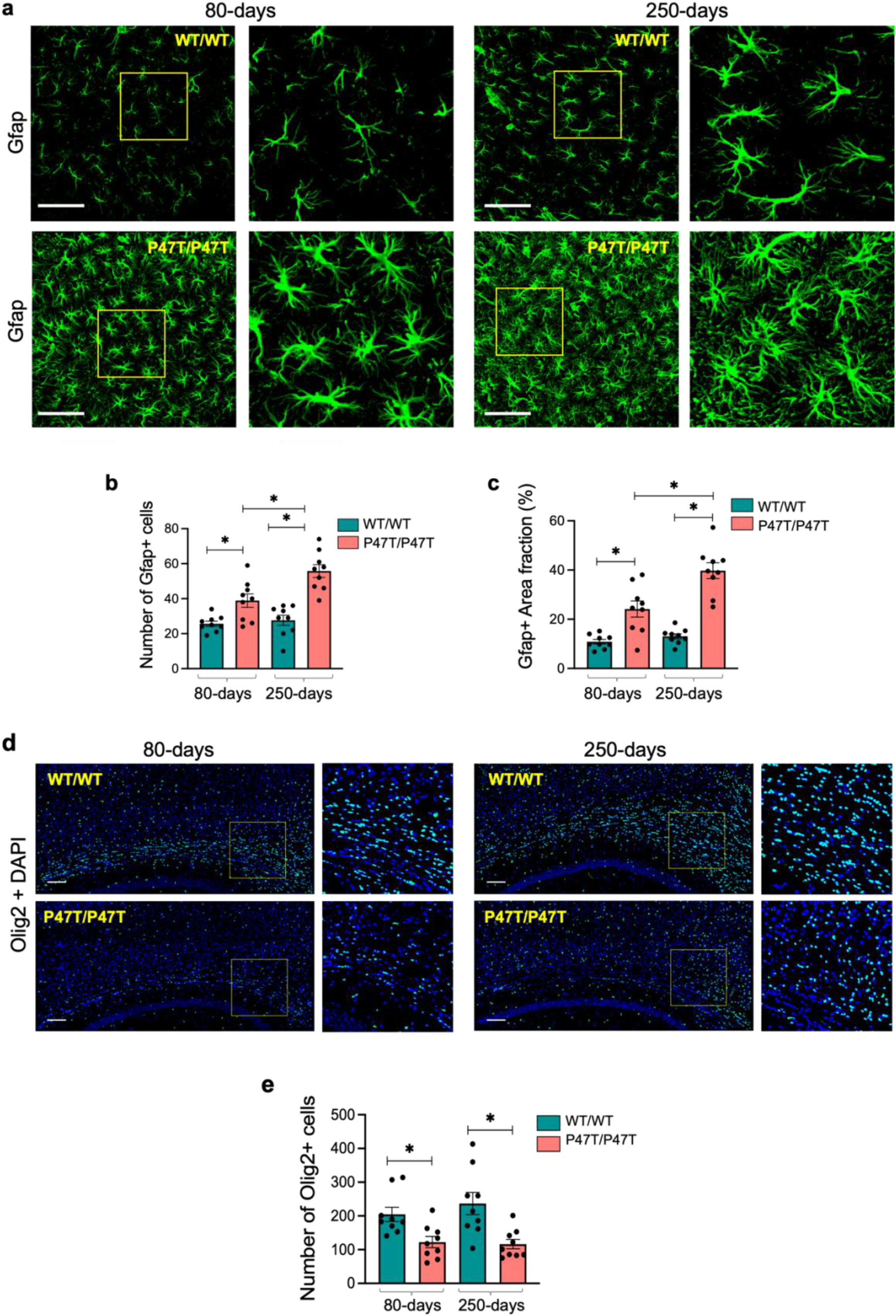
*Wwox^P47T/P47T^* hippocampi exhibit progressive astrogliosis and reduced oligodendrocyte numbers. (a) Representative high magnification confocal images illustrating comparative abundance and morphology of Gfap+ astrocytes in 425 μm^2^ coronal section of HPC CA3 region of *Wwox^WT/WT^* and *Wwox^P47T/P47T^* samples at 80 and 250 days of age as indicated. Yellow square shows the region selected and zoomed to show more intricate morphology as displayed adjacent to each image. Bar graphs compare the average number of Gfap1+ astrocytes (b) and Gfap+ area fraction % (c) in the CA1, CA3, and DG subfields taken together for *Wwox^WT/WT^* and *Wwox^P47T/P47T^* in 80 and 250-days groups. Each data point represents quantitation from a single subfield of n=3 mice/group. (d) Coronal sections of brain tissue immunostained for Olig2 pan-marker for labeling all oligodendrocyte lineages. Representative images showing number of Olig2 stained oligodendrocytes in the 1.5 x 1 mm^2^ corpus callosum section. Yellow square shows the region selected and zoomed as displayed adjacent to each image. (e) Bar graph showing quantitative measurement of Olig2 positive oligodendrocytes counted in three independent 500 μm^2^ regions spanning the entire imaged corpus callosum. Each data point shows measurement from a single independent region from n=3 mice/group. Scale bar=100 μm, data is represented as mean ± SEM, *p-value < 0.01, unpaired Student’s *t*-test.

In summary, these data indicate significant microgliosis and astrogliosis in brains from *Wwox^P47T/P47T^* mice that increase with aging and are indicative of a progressive neuroinflammatory process. It is worth noting that evaluations of both Iba1 and Gfap immunostainings were also performed in brain samples from *Wwox^P47T/WT^* mice; however no differences were observed when compared with the wildtype mice (data not shown). Thus, evidence of reactive astro-microgliosis was only observed in mice homozygous for the *Wwox* P47T mutation.

### *Wwox^P47T/P47T^* brains exhibit a reduced number of oligodendrocytes

Oligodendrocytes produce myelin sheath in CNS, which is critical for the insulation of axons and transmission of the proper action potential [25]. Since SCAR12 patients display a seizure phenotype and severely impaired cognitive and motor functions indicative of abnormal neurotransmission, we examined oligodendrocyte numbers and myelination in *Wwox^P47T/P47T^* brains. The number of oligodendrocytes was measured by immunostaining for Olig2, a pan-marker labeling all oligodendrocyte lineages [25]. The corpus callosum of *Wwox^P47T/P47T^* mice showed significantly reduced oligodendrocytes numbers compared to *Wwox^WT/WT^* (p-value < 0.01) (Figs. 4d and e). We also analyzed myelination by staining with myelin basic protein (Mbp) antibody in the parietal cortex above the corpus callosum. However, unlike observations for oligodendrocyte numbers, no significant differences in Mbp staining were detected when comparing *WwoxWT/WT* and *Wwox^P47T/P47T^* cortical regions (Supplementary Figs. 6a-d).

### *Wwox* loss of function leads to cerebellar atrophy and Purkinje cell degeneration

We examined brains from wildtype and homozygous mutant mice to determine whether the profound motor and social deficits observed in *Wwox^P47T/P47T^* mice arise from abnormal CB development. *Wwox^P47T/P47T^* mice displayed aberrant cerebellar morphology with fusion of interhemispheric fissure and cerebellar vermis lobules (Supplementary Fig. S7). We next analyzed cerebellar histomorphology by means of confocal microscopy. Image analyses of sagittal CB tissue sections from Wwox homozygous mutant mice revealed significant cerebellar hypoplasia and PC degeneration (Figs. 5a-c, Supplementary Fig. S8a). Cerebellar cortex from *Wwox^P47T/P47T^* mice displayed a striking decrease in the number of calbindin labeled PC bodies with deformed and degenerated dendrites in most folia of the cerebellar cortex (Figs. 5a-c). Figure 5b (left panel) shows the dense deposition of PC dendrites in the molecular layer of the *Wwox^WT/WT^* CB, while in *Wwox^P47T/P47T^* mice, severe dendritic atrophy and pruning of dendritic branches are evident (Fig. 5b, two right panels). Determination of PC and basket cell (Hcn1+) numbers indicate that while PC numbers were significantly reduced in the cerebellar cortex from *Wwox^P47T/P47T^* mice, the number of basket cells in the same regions remained unchanged (Figs. 5c and e). Next, we determined molecular layer (ML) thickness as a measure of dendritic span for the overall assessment of PC dendrites degeneration and also determined granular layer (GL) thickness by calretinin immunostaining. All measurements were performed on the preculminate and primary fissures between cerebellar lobules III-VI. The average molecular and granular layer thickness was significantly reduced in the cerebellar cortex from *Wwox^P47T/P47T^* mice in both age groups, indicative of significant CB atrophy in these animals (p-value < 0.001, Figs. 5f-h, Supplementary Fig. S8b). ML and GL thickness were comparable, and there were no age-related differences between the *Wwox^P47T/P47T^* mice at 80-days and 250-days (Figs. 5g and h).

**Figure 5.**
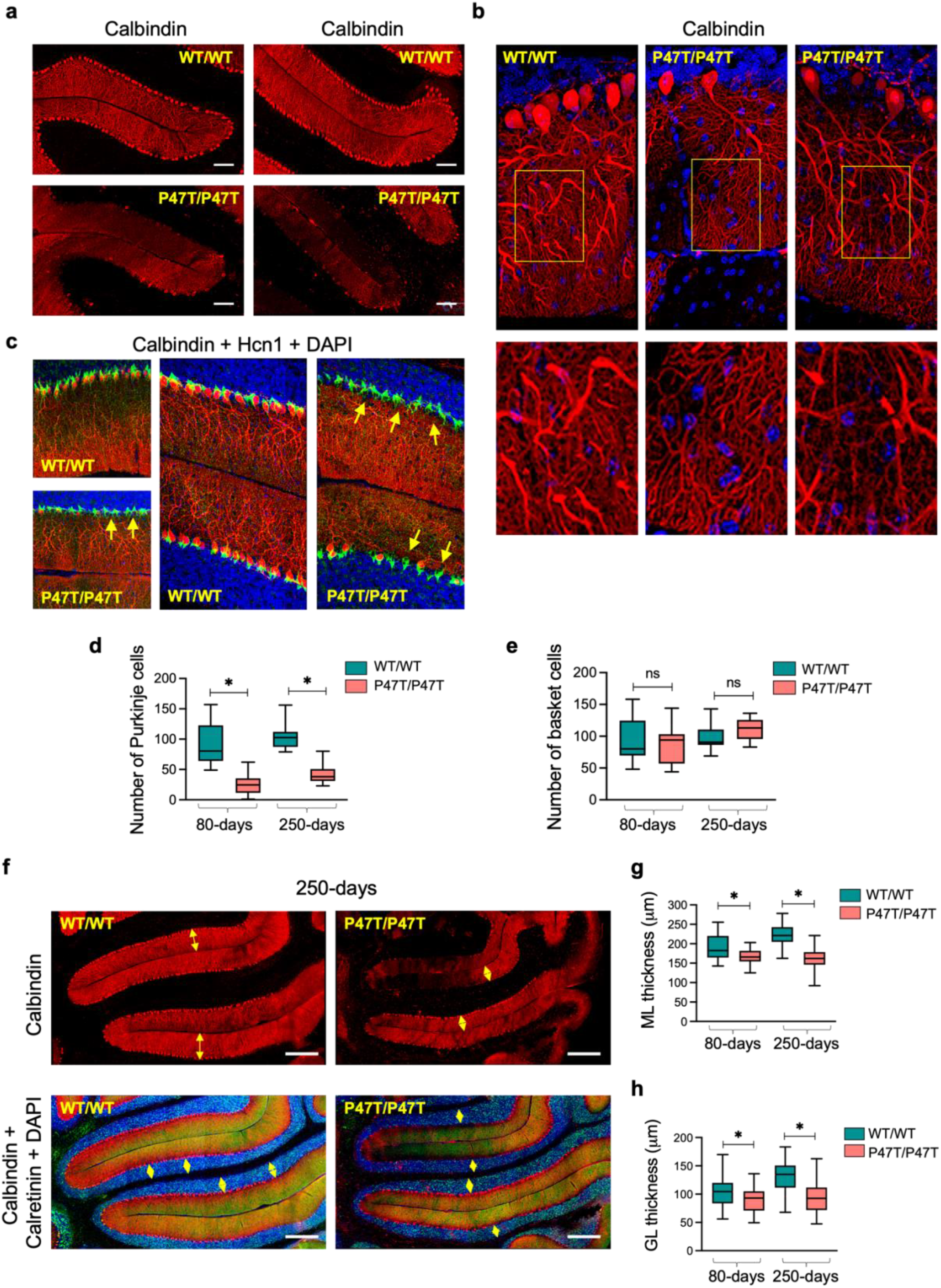
*Wwox* LoF leads to cerebellar atrophy and Purkinje cell degeneration. (a) Comparative confocal images of the fissure separating lobules II/III (left panel) and III/IV (right panel) in mid-sagittal sections of the vermis region of CB from *Wwox^WT/WT^* and *Wwox^P47T/P47T^* mice of 250-days age group. Samples were immunostained with anti-calbindin antibody identifying PC bodies and dendrites in molecular layer of CB cortex (red stain). Note the dramatic absence of PCs in folia of *Wwox^P47T/P47T^* CB. (b) Comparative high magnification images of PC bodies and dendrites (250-days age group). Loss of PCs and degenerating dendrites are evident in images from *Wwox^P47T/P47T^* CB compared to *Wwox^WT/WT^* controls. Yellow rectangle marks the region selected and zoomed to show more intricate morphology as displayed below each image. (c) Comparative immunostaining of PCs with anti-calbindin antibody (red cells) and basket cells with anti-Hcn1 antibody (green cells). Left panel top and bottom images show immunostaining of mice CB from 80-days age group, right two panels show images from 250-days age group. Note significant decreased thickness of the molecular layer in the *Wwox^P47T/P47T^* sample, with basket cells present and no PC bodies (yellow arrows). (d-e) Box and Whisker plots comparing the number of calbindin+ PCs (d) and number of Hcn1+ basket cells (e) in comparative 1000 μm^2^ sections from *Wwox^WT/WT^* and *Wwox^P47T/P47T^* pairs (n=3) at 80 and 250-days of age as indicated. (f) Representative calbindin immunofluorescence of PCs (top panel, red) and calretinin (green) and DAPI (blue) immunofluorescence of GL (bottom panel) in preculminate and primary fissures between cerebellar lobules III-VI of 250-days old *Wwox^WT/WT^* and *Wwox^P47T/P47T^* mice. Yellow arrows in the top panel mark the ML thickness and in the lower panel mark the GL thickness. (g-h) Box and Whisker plots showing the quantitative measurements of ML (g) and GL (h). Twenty four measurements, with 12 measurements each spanning the entire preculminate and primary fissures were taken from *Wwox^WT/WT^* (n=3) and *Wwox^P47T/P47T^* (*n=3*) at 80 and 250-days age groups. Fig. a Scale bar=100 μm, Fig. f *s*cale bar=200 μm, data is shown as box extending from 25^th^ to 75^th^ percentile, with horizontal line at median (50^th^ percentile), whiskers represent bottom and top 25% and extend down to the smallest and up to the largest value, *p-value < 0.05, unpaired Student’s *t*-test.

### Transcriptome profiling provides further evidence of neuroinflammation and glial cell dysfunction in forebrains of *Wwox* P47T mice

To gain insight into transcriptional changes associated with *Wwox* P47T mutation phenotypes, we first performed bulk RNA sequencing (RNA-seq) on forebrain tissue regions, prefrontal cortex (PFC), parietal cortex (CTX), and hippocampus (HPC) from *Wwox^WT/WT^* (n=5) and *Wwox^P47T/P47T^* (n=5) mice. Transcriptome data showed that *Wwox* gene expression was comparable between wildtype and homozygous samples across all three tissue regions (Supplementary Fig. S9), confirming that the P47T mutation does not affect Wwox mRNA expression in the brain.

Unsupervised hierarchical clustering of RNA-Seq profiles of the forebrain samples robustly segregated all different brain regions and separately clustered samples from P47T homozygous and wildtype mice (Fig. 6a). Differential gene expression (DGE) datasets (log2 FC > ±1, FDR < 0.05) identified a total of 425 DEGs in PFC (338 genes upregulated, 87 genes downregulated), 363 in CTX (312 genes upregulated, 51 genes downregulated), and 1508 DEGs in HPC (986 genes upregulated, 522 genes downregulated) (Supplementary File 1). Of these DEGs, an overlapping set of 208 transcripts was commonly dysregulated in all three forebrain tissue regions (Fig. 6b, Supplementary File 2). Out of the 208 common DEGs, 204 were dysregulated in the same direction in all three tissues, with 190 upregulated and 14 downregulated transcripts (Supplementary File 2). Among the common upregulated genes, we found genes related to neuroinflammation (*Tgfβ1, Oas1f, Ighg1, Il17rb, Ripk3*), neuronal differentiation (*Wnt9b, Ascl2, Lhx3*), synaptic transmission (*Drd4, Nptx2*) G protein signaling (*Npw, Rgs13*), lipid metabolism (*Acaa1b, Lipg, Plb1*), and calcium signaling (*Atp2a1, Stc2, Atp2c2’*), among others (Supplementary File 2).

**Figure 6.**
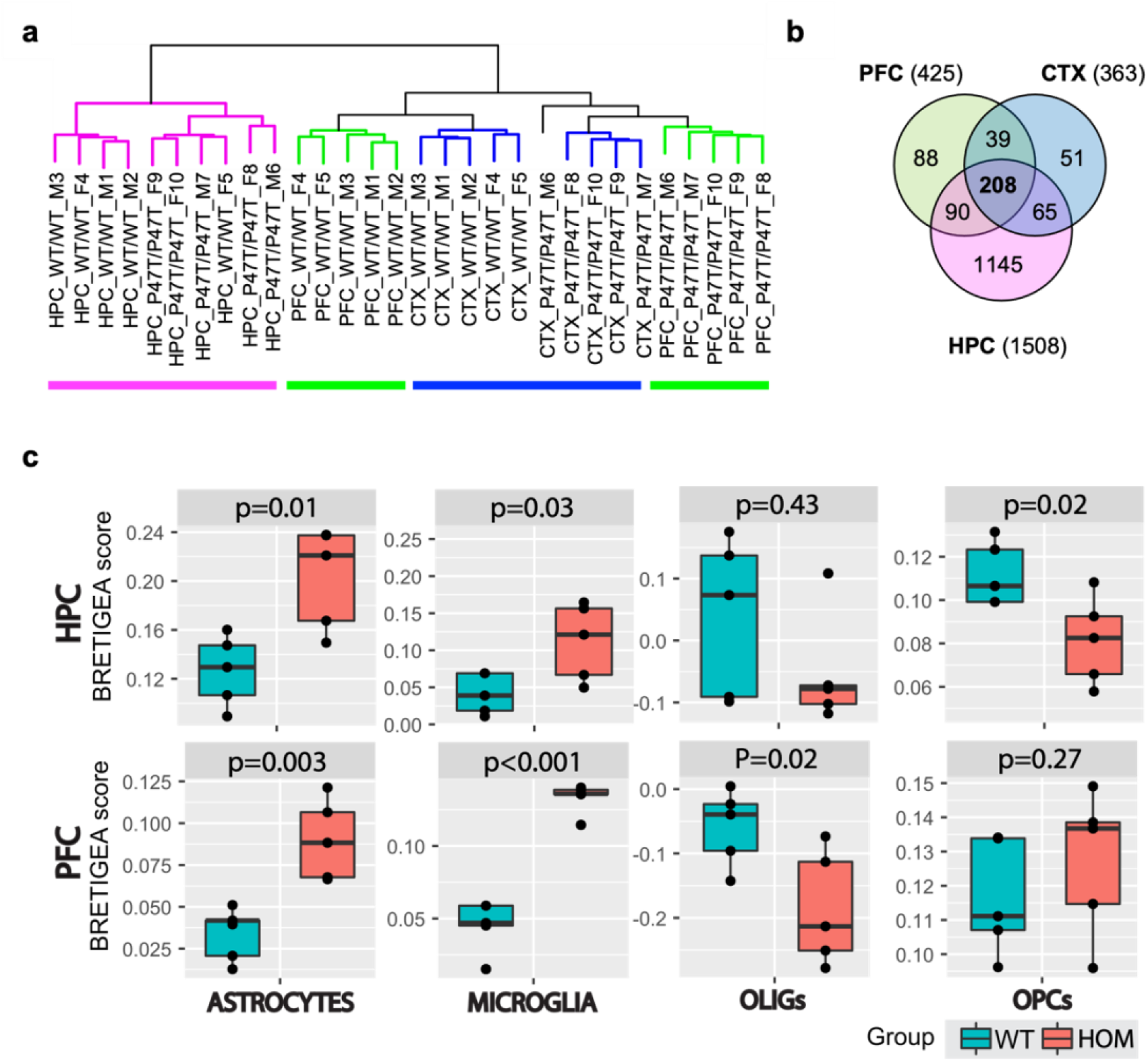
Unsupervised hierarchical clustering segregates clusters of forebrain regions. (a) Hierarchical clustering of RNA-Seq profiles showing segregated clusters of the three forebrain regions and within each region separation of *Wwox^WT/WT^* from *Wwox^P47T/P47T^* samples (n=5/tissue). Pink: HPC samples, green: PFC, blue: CTX. (b) Venn diagram of transcripts dysregulated in PFC (425), CTX (363), and HPC (1508) samples with log2 FC > ±1, and FDR < 0.05, 208 transcripts were commonly dysregulated in all three tissues. (c) Comparative BRETIGEA scores for HPC (top row) and PFC (bottom row) tissues from RNA-Seq profiles of *Wwox^WT/WT^* and *Wwox^P47T/P47T^* tissues (n=5). OLIGs= oligodendrocytes and OPCs= oligodendrocytes progenitor cells, respective p-values are indicated on top of the plot, Student’s *t*-test.

The R package BRETIGEA (**<u>Br</u>**ain c**<u>e</u>**ll **<u>t</u>**ype specific **<u>g</u>**ene **<u>e</u>**xpression **<u>a</u>**nalysis) was employed to estimate relative brain cell type representation from the RNA-seq profiles [26]. Figure 6c shows representative results of comparative cell type estimates in the *Wwox^WT/WT^ and Wwox^P47T/P47T^* HPC and PFC regions. Both brain regions demonstrate significantly higher scores for astrocyte and microglia gene expression signatures in the P47T homozygous mutant samples compared to the wildtype samples, which agrees with the confocal imaging observations of significant astro-microgliosis with Wwox LoF (Figs. 3 and 4a-c). It also shows reduced scores for mature oligodendrocytes (OLIGs) and oligodendrocytes progenitor cells (OPCs) in the HPC of *Wwox^P47T/P47T^* mice, which is also in line with the immunofluorescence imaging data, where a reduced number of Olig2+ oligodendrocytes was observed in the corpus callosum of homozygous mice (Figs. 4d and e)

Using gene set enrichment analysis (GSEA) of differential *Wwox^P47T/P47T^* and *Wwox^WT/WT^* gene expression profiles, we identified the most prominent activated or suppressed bioprocesses and pathways in each brain region. Representative results from the HPC and CTX are shown in Figure 7a and revealed major suppression of gene networks associated with brain development, glial cell differentiation, postsynaptic density, and activation of gene networks related to neuropeptide binding, inhibitory postsynaptic potential, G protein-coupled receptor activity, and activation of bioprocesses related to inflammation (Figs. 7a and b). Interestingly, GSEA of the CTX, despite representing the transcriptome profile of a different brain region, was particularly informative relating to the suppression of cerebellar biofunctions not only associated with CB development but also with PC differentiation, morphogenesis, and PC layer development (Fig. 7b). This is in strong agreement with our observations from CB imaging and phenotype of *Wwox^P47T/P47T^* mice. Similarly, PFC also showed enrichment of bioprocesses related to activation of inflammation (Supplementary Fig. S10), a theme that is also common in KEGG pathway enrichment analyses for all regions. (Supplementary Figs. S11-13). Suppression of GABAergic and glutamatergic synapses in CTX and HPC region was also detected in KEGG pathways suggesting abnormal neurotransmission likely related to the epileptic and seizure activity observed in *Wwox^P47T/P47T^* mice (Supplementary Figs. S11 and 12).

**Figure 7.**
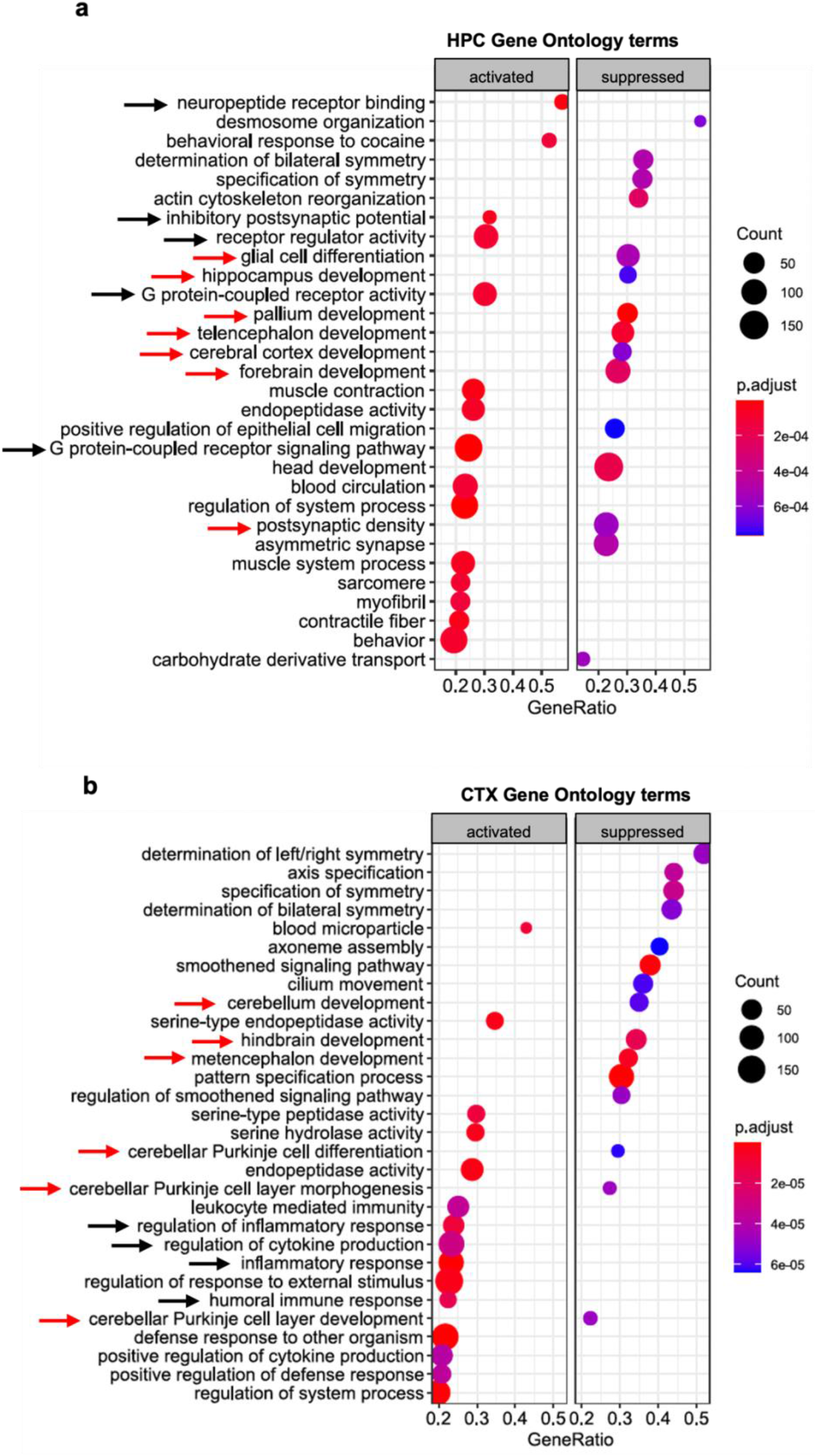
Transcriptome profiling provides further evidence of neuroinflammation and glial cell dysfunction in forebrains of *Wwox* P47T mice. (a-b) clusterProfiler plot showing Gene Ontology terms enriched in *Wwox^P47T/P47T^* HPC (a) and CTX (b) regions. Red arrows indicate gene networks associated with brain development and black arrows indicate bioprocesses related to G protein signaling in PFC and inflammation in CTX. Left panel shows activated and right panel shows suppressed GO terms. Color of the circle represents the adjusted p-value and size represents number of genes enriched.

IPA-based functional annotation of transcriptome alterations revealed top-activated and inhibited upstream regulator pathways in all three tissues (HPC, PFC, CTX). The cAMP response element binding protein 1 (CREB1) pathway was implicated as an activated upstream regulator in all three regions (Fig. 8a). Furthermore, ‘CREB Signaling in Neurons’ was also a top activated canonical pathway in IPA analysis (data not shown). Additional themes in the upstream regulator analysis include multiple inflammatory-related molecules and pathways such as TGFβ, TNF, NFκB, and other inflammatory cytokine pathways (Fig. 8b). Likewise, a top IPA canonical pathway activated in all three brain regions is that of ‘Phagosome formation’ (data not shown). GSEA Hallmark gene set analyses also identified enrichment of ‘Inflammatory Response’, ‘TNFA signaling via NFkB’, and ‘IL6-JAK-STAT3 signaling’ (Fig. 8b). All these observations strongly implicate neuroinflammatory changes in the pathology associated with Wwox LoF. Interestingly, the Hallmark gene-set analysis also revealed hypoxia, apoptosis, and glycolysis pathways enrichment in all three brain regions (Fig. 8b).

**Figure 8.**
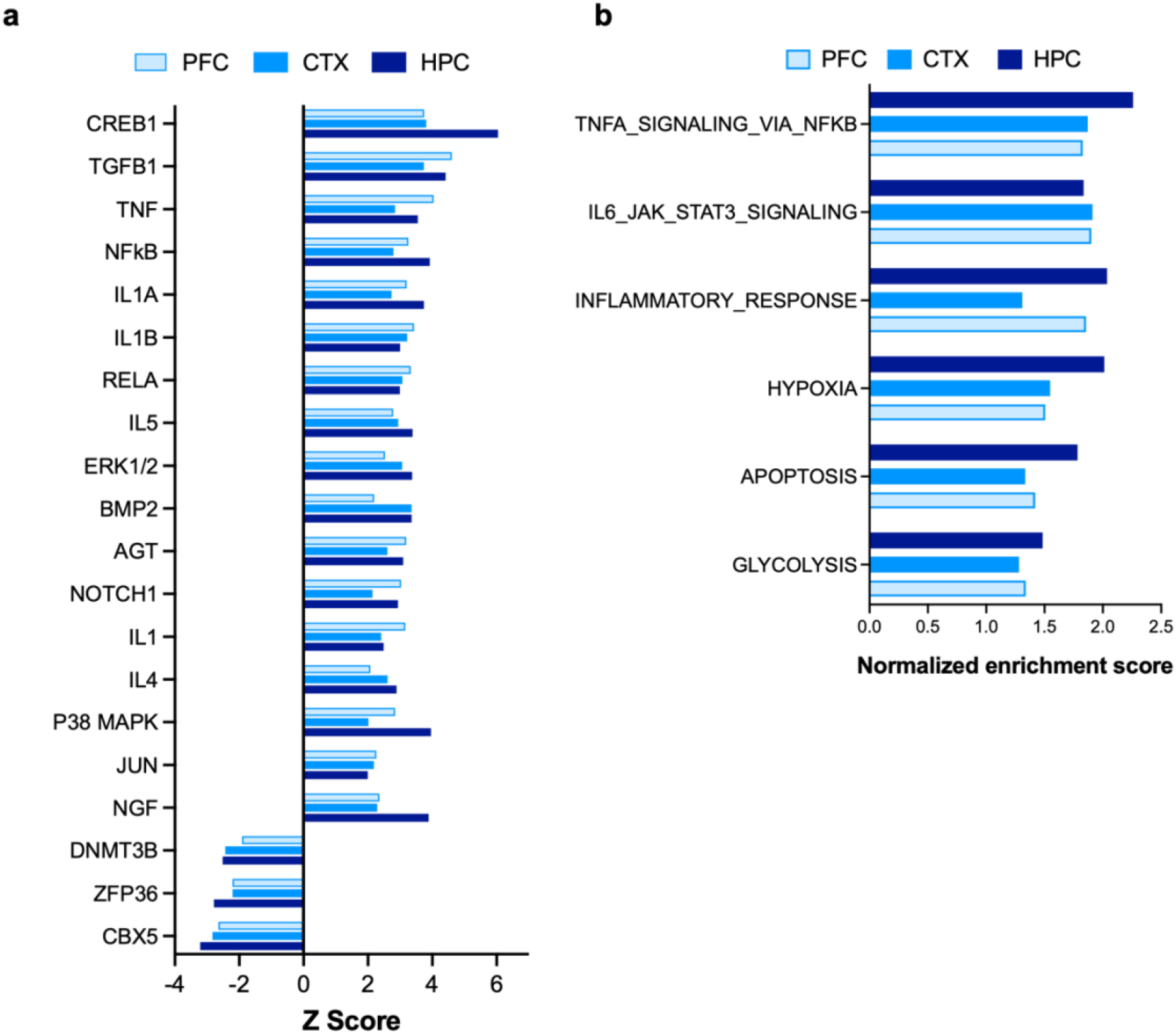
IPA upstream regulators and GSEA Hallmark gene set analysis. (a) Bar graph showing commonly upregulated and downregulated upstream regulators identified by comparative analysis using IPA in the three forebrain regions. RNA-Seq DEG profiles from PFC, CTX and HPC of *Wwox^P47T/P47T^* samples vs. *Wwox^WT/WT^* were compared to identify top common altered upstream regulators pathways. (b) Bar graph showing pathways and bioprocesses enriched by GSEA Hallmark gene set analysis in PFC, CTX, and HPC samples.

### Transcriptome profiling of *Wwox* P47T cerebella provides further evidence of dysfunction

Since *Wwox^P47T/P47T^* mice displayed motor abnormalities along with the evidence of significant cerebellar atrophy and PC degeneration, we sought to evaluate the transcriptional changes associated with Wwox LoF in CB. Like the forebrain tissues, *Wwox* gene expression was comparable between wildtype and P47T homozygous CB samples (Supplementary Fig. S9) and in agreement with earlier described results (Figs. 1a and S2a). Unsupervised hierarchical clustering of the CB RNA-Seq profiles segregated samples from P47T homozygous and wildtype mice (Fig. 9a). EdgeR analysis identified a total of 1059 DEGs (376 genes upregulated, 683 genes downregulated) comparing both groups at an FDR < 0.01 (Supplementary File 1). Annotation of CB transcriptional differences using IPA showed striking enrichment of diseases and biofunctions related to severe cerebellar dysfunction. Congenital neurological disorder and encephalopathy, motor dysfunction and movement disorder, and hypoplasia of the brain, were among the top enriched diseases and functions with a positive Z score in the range of 2.1 – 6.8 and highly significant p-values (-log10 p-values range of 3.6 – 10.9) (Fig. 9b). This implies that the CB is drastically affected by Wwox LoF in agreement with the ataxic phenotype observed and the described confocal studies. Interestingly, GSEA hallmark gene set analysis showed that the topmost enriched gene sets in CB belonged to lipid metabolism and reactive oxygen species pathway (Fig. 9c). In addition, it also showed enrichment of matching gene networks and pathways identified in forebrain tissues, including apoptosis, inflammatory response, IL6-JAK-STAT3 signaling, glycolysis, and hypoxia (Fig. 9c). Given these data and the significance of neuroinflammatory changes seen in forebrain samples, we investigated the occurrence of neuroinflammatory abnormalities in CB. We analyzed the expression of pro-inflammatory cytokines *IL1β* and *TNFα* by qRT-PCR in CB tissue and observed a significant ≥ 2-fold upregulation of mRNA of both cytokines (p-value < 0.05) in CB samples from *Wwox^P47T/P47T^* mice (n=6) compared to *WT* mice (n=6) (Figs 9d and e).

**Figure 9.**
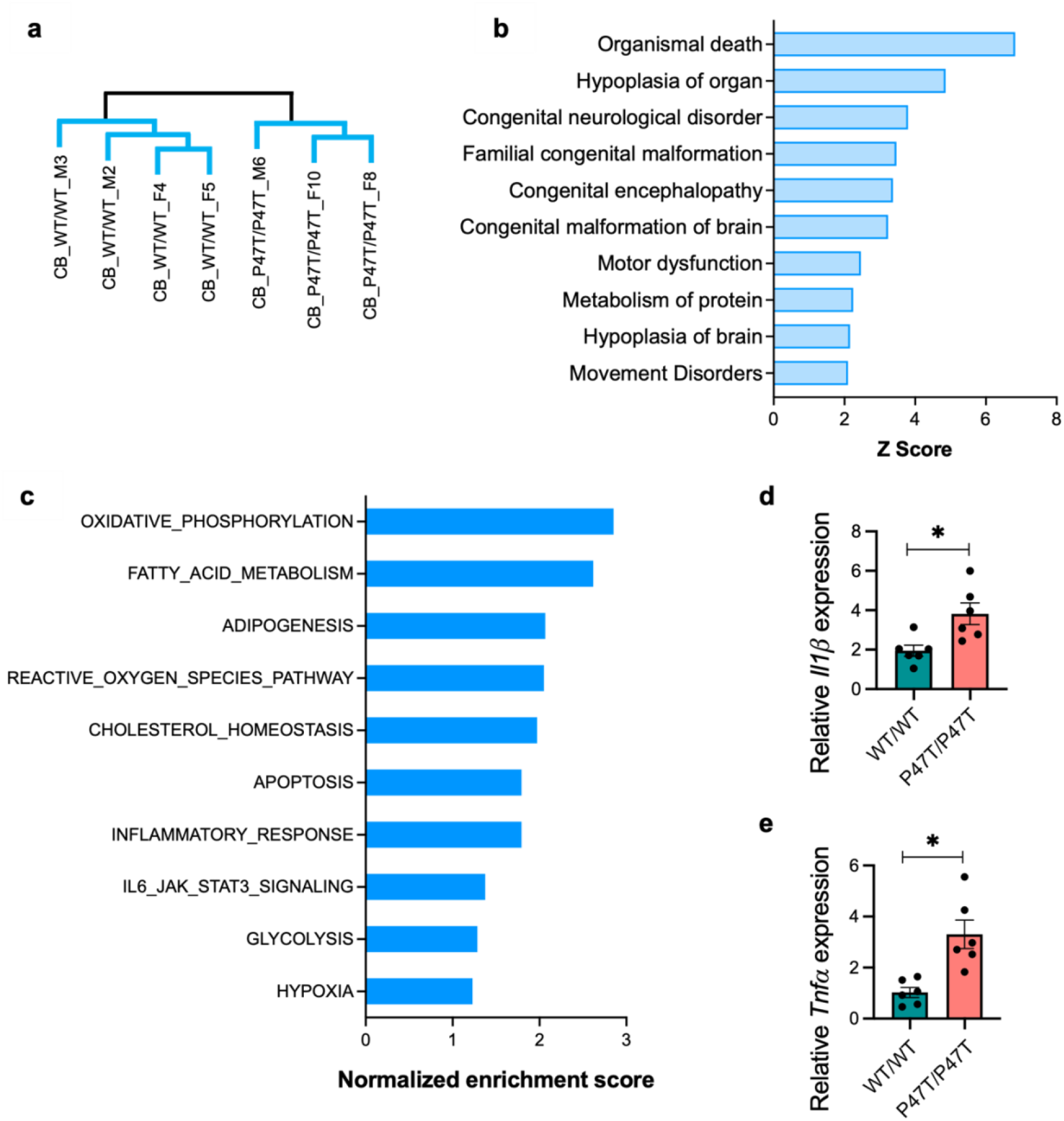
Transcriptome profiling provides further evidence of severe dysfunction in *Wwox P47T* cerebella. (a) Unsupervised hierarchical clustering of RNA-Seq profiles showing separated clusters of *Wwox^WT/WT^* (n=4) and *Wwox^P47T/P47T^* (n=3) CB samples. (b) Bar graph showing IPA based enrichment of diseases and functions primarily related to cerebellar dysfunction and movement disorder. (c) Bar graph showing pathways and bioprocesses enriched by GSEA Hallmark gene set analysis. (d-e) Bar graphs showing mean *IL1β* (d) and *TNFα* (e) mRNA expression measured by qRT-PCR in *Wwox^WT/WT^* and *Wwox^P47T/P47T^* CB (n=6 mice/group). Error bars represent ± SEM, *p-value < 0.05, unpaired Student’s *t*-test.

Overall, our RNA-Seq results highlight widespread changes in neuronal and glial pathways in Wwox LoF mice. Wwox LoF is associated with an increase in neuroinflammatory changes in addition to suppression of neurodevelopmental and differentiation bioprocesses, in particular affecting the hindbrain. These changes potentially play an important role in contributing to behavioral and motor pathology associated with Wwox LoF.

## DISCUSSION

We have generated a mouse model that phenocopies the clinicopathological features of the human SCAR12 syndrome by means of substituting a single amino acid in the Wwox protein (p.Pro47Thr). Our model confirms that this human condition, characterized by early-onset epilepsy, severe intellectual disability and cerebellar ataxia, occurs as a consequence of LoF of the Wwox protein. Indeed, *Wwox^P47T/P47T^* mice displayed severe chronic seizure activity, profound behavioral dysfunction, and significant motor impairment. This mouse model allowed us to gain the first insight into the complex brain abnormalities caused by the expression of a hypomorphic Wwox mutation which results in a defective protein incapable of binding partner proteins.

Among the most striking findings of our studies was the magnitude and extent of neuroinflammatory changes affecting the brain of *Wwox^P47T/P47T^* mice which included progressive astro-microgliosis increasing in severity with age. Transcriptome profiling studies highlighted a number of inflammatory pathways altered in Wwox LoF mice and pathognomonic of microglia pro-inflammatory gene signatures. In previous studies, we observed that Wwox deficiency had an association with neuroinflammation, since *Wwox-KO* hippocampi displayed increase in Iba1+ microglia, astrogliosis, and overexpression of inflammatory cytokines [27]. Interestingly, upregulation of *Wwox* expression is observed in microglial cells upon systemic treatment of mice with inflammatory agents such as lipopolysaccharide, suggesting that Wwox might play a role in CNS inflammatory response [1]. Indeed, evidence from studies in cancer and various inflammatory pathologies strongly suggest that Wwox regulates inflammation and direct associations between Wwox and the NF-kB and IL-6/JAK2/STAT3 inflammation signaling pathways have been demonstrated [1, 28–34]. Microglia are vital for healthy brain development and function, regulating brain development primarily through two routes: the release of diffusible factors (e.g., pro-inflammatory cytokines) and phagocytosis. The phagocytic capacity of microglia is particularly important during early brain development and subsequent maturation as they remove excessive synapses and sculpt the precise, organized, circuitry characteristic of the healthy adult brain [35]. Further, microglial dysfunction is implicated in several neurodevelopmental and neurodegenerative disorders; for example, microgliosis is known to contribute to deficits in social behavior in other contexts [36]. It is further hypothesized that in the diseased brain, interactions between damaged neurons and uncontrolled, chronic inflammation ultimately drives the progression of neurodegenerative diseases [22]. Based on our findings, we hypothesize that impaired Wwox function leads to microglial dysfunction, chronically enhancing neuroimmune activation and ultimately contributing to widespread brain pathology.

In behavioral tests, *Wwox^P47T/P47T^* mice displayed heightened anxiety-like behaviors and profound deficits in social interaction. There is growing evidence implicating immune system disruption in the pathogenesis of social stress and abnormal behavioral features [37, 38]. Challenges to the immune system result in a constellation of symptoms described as the sickness response, which are remarkably similar to hallmarks of neuropsychiatric disorders including social withdrawal, anxiety/agitation, cognitive impairments, anhedonia, and altered sleep [reviewed in [39]]. Importantly, while the sickness response typically rapidly resolves, sensitization or low grade activation of the neuroimmune system can cause exaggerated and prolonged expression of these behaviors [40]. Activation of the immune system during early developmental periods can also reshape neuronal circuitry resulting in persistent deficits in social behaviors [reviewed in [36]]. Since in *Wwox^P47T/P47T^* mice impaired Wwox function enhances neuroimmune activation increasing substantially in severity with mouse aging, we hypothesize that the observed social deficits and anxiety-like behavior in these mice are the result of the widespread neuropathology caused by chronic neuroinflammation.

In addition to neuroinflammatory signatures, transcriptome pathway analyses pointed to significant activation of CREB1 (cAMP response element binding protein 1) and NGF (nerve growth factor) downstream signaling among the most prominently affected pathways (see Fig. 8a). Activation of the CREB pathway can be induced by various growth factors and inflammatory signals leading to a variety of biological responses including neuronal excitation and neural cell proliferation and survival among others [41, 42]. For example, among the common genes upregulated in all forebrain regions and among downstream targets of CREB1, we detected significant upregulation of, NPTX2 (neuronal petraxin 2) a protein involved in excitatory synapse formation and BDNF (brain derived neurotrophic factor) both significantly upregulated in *Wwox^P47T/P47T^* brain tissues (Supplementary File 2). These changes suggest active neurogenesis as compensatory mechanisms to chronic inflammation and neurodegeneration.

Wwox is expressed in all neuronal and glial cell types, but its expression is comparatively higher in cerebellar cortex than in all other CNS structures [1]. Our studies demonstrate that the *Wwox* P47T mutation led to cerebellar cortex atrophy with significant loss of Purkinje cells and dendritic degeneration ultimately leading to the motor and in part behavioral deficits observed affecting *Wwox^P47T/P47T^* mice. These observations were further supported by transcriptome profiles from mutant mouse cerebella indicating gene dysregulations associated with congenital neurological disorders, cerebellar dysfunction, and movement disorders. The degeneration observed affecting specifically Purkinje cells suggests the critical need of full Wwox function for the normal development or differentiation of this critical cerebellar neuronal cell type.

Brains of *Wwox^P47T/P47T^* also showed evidence of a decrease in oligodendrocytes numbers supported by histological evidence from corpus callosum and BRETIGEA estimates of RNA-Seq data. These observations agree with previous studies in *Wwox* whole body and neuron-specific *KO* mice [43, 44] and with the lde/lde rat model of spontaneous Wwox mutation [45]. All of these models of complete Wwox deficiency showed a significant decrease in the number of mature oligodendrocytes and an association with hypomyelination. In our studies, however, despite observing a decrease in oligodendrocyte numbers, we did not detect significant differences in myelination by Mbp staining when comparing Wwox wildtype with homozygous mutant mice.

Autosomal recessive cerebellar ataxias (ARCAs) such as SCAR12, are rare heterogeneous neurodegenerative hereditary conditions that usually implicate genes that play roles in a cluster of molecular themes such as mitochondrial metabolism, oxidative stress, DNA repair, genome stability and lipid metabolism (Reviewed in [46]). Indeed, we and others have demonstrated that WWOX is implicated in various such molecular themes, including: DNA repair/genome stability [47–49], mitochondrial respiration/oxidative stress [50–52], and lipid metabolism [53–55]. CNS pathologies with demonstrable germline *WWOX* LoF are rare, however, it is estimated that approximately 50% of patients affected by ARCAs remain without a molecular diagnosis despite next-generation sequencing advances [46]. This may reflect, in many cases, causality by mechanisms other than germline mutations such as epigenetic causes, gene dosage defects, dysregulation at the protein level, and posttranslational modifications, among other possibilities [46]. WWOX’s protein interactome is vast and complex, as we and others have demonstrated [20], and this protein is indeed a prime target for dysregulation by mechanisms other than specific germline mutation.

In previous studies modeling Wwox deficiency, we used full *Wwox-KO* mice, which to a certain extent recapitulate the most severe human WOREE syndrome. Like WOREE children, *Wwox-KO* mice displayed spontaneous seizures, motor deficits, ultimately reaching almost total immobility, and a short lifespan (~3 weeks) [4, 53, 56]. Studies by others using whole body and neuronal conditional *Wwox-KO* models substantiated our original observations [43, 44]. Even though these studies have been instrumental in building on our understanding of Wwox LoF in CNS disorders, one of their major limitations of *Wwox-KO* mice is the short life span of only 3 weeks, making them unsuitable models for long-term CNS and behavioral studies. Contrary to *Wwox-KO* models, our *Wwox^P47T/P47T^* mice reached adulthood and have a long life span (mean overall survival of almost 400 days). In addition, the introduction of the P47T mutation had no detrimental effect on normal mouse growth or reproduction in *Wwox^P47T/WT^* mice. Thus, the mouse model here described will not only be useful for the study of the rare ARCA such as SCAR12, but is likely more generalizable to understand the role of WWOX in the pathogenesis of a variety of far more common neurodegenerative conditions where this gene has been identified as a risk factor (e.g., Alzheimer’s disease, Parkinson’s disease) [11, 13–18]. Finally, this novel mouse model of neurodegeneration can provide a useful *in vivo* tool for testing promising therapeutic strategies for neurodevelopmental and neurodegenerative conditions in which WWOX loss of function plays a pathogenic role.

## MATERIAL AND METHODS

### Development of *Wwox* P47T mouse model

*Wwox^P47T/P47T^* mutant mice were developed using a CRISPR/Cas9 Knock-in system. *Wwox* sgRNA and Cas9 were *in vitro* transcribed (IVT) into mRNAs and a ready to inject Cas9 and sgRNA premix was synthesized by Horizon Discovery (St. Louis, MO). FVB/6N female mice were used as embryo donors and ICR females were used as the surrogates. Ovulated 6-8 week old female FVB/6N mice were mated to FVB/6N males, and the next morning the fertilized eggs from the oviducts were collected. *Wwox* P47T SSOD (3g/μl), *Wwox* sgRNA/<u>PAM</u> (cagtgggaacatccgaaaac<u>CGG</u>) (20 ng/μl), *Cas9* mRNA (30ng/ul) were mixed in 100 μl of microinjection buffer. The oligo, sgRNA, and Cas9 mixture was micro-injected into the pronucleus of the zygotes at the 1-cell stage, in M2 medium (Sigma). Injected zygotes were cultured in EmbryoMaxx AA medium (Millipore) at 37 °C and 5% CO2 for 1 day. The surviving embryos at the 2-cell stage were transferred into the infundibulum of ICR surrogate female mice. To identify target mice from the resulting litter, DNA from mouse tails was PCR amplified and genotype was confirmed by Sanger sequencing. Multiple pups were screened by sequencing until the founder pups with the required Wwox mutation were obtained. Founder mice were further crossed multiple times to clean up any spurious CRISPR artifacts in other genomic regions thus generating final colonies. *Wwox^P47T/WT^* mice were used to maintain the line and bred to homozygosity along with heterozygous and wildtype littermate controls for further investigations. Subsequently, experimental animals were genotyped using custom Kompetitive Allele Specific PCR (KASP, LGC Biosearch Technologies, UK) genotyping assay [57]. All animals were bred and kept in a clean, modified-barrier animal facility, fed regular commercial mouse diet (Harlan Lab., Indianapolis, IN) under controlled light (12L:12D) and temperature (20–24 °C). Comparative survival was assessed in a cohort of *Wwox^WT/WT^* (n=14, 7 males and 7 females), *Wwox^P47T/WT^* (n=30, 21 males and 9 females), and *WwoxP47T/P47T*

(n=25, 15 males and 10 females) mice. All animal research was conducted in facilities accredited by the Association for Assessment and Accreditation of Laboratory Animal Care International (AAALAC) at the University of Texas, MD Anderson Cancer Center, following international guidelines and all research was approved by the corresponding Institutional Animal Care and Use Committee (IACUC).

### Oligopeptide pull-down assay

*In vitro* peptide pull-down assay was performed using whole CB lysates from *Wwox^WT/WT^* and *Wwox^P47T/P47T^* (n=2/group) of either sex. Biotinylated Dvl2 (HPYSPQ**<u>PPPY</u>**HELSSY) and Wbp1 (SGSGGTP**<u>PPPY</u>**TVG) peptides (15 μg) were incubated with 30 μl magnetic MyOne™ Streptavidin C1 Dynabeads™ in 300 μl 1x PBS with 0.1% BSA for 2 hours at 4 °C. After incubation, beads were washed in 1x PBS with 0.1% BSA three times and 10% peptide-bead complex was aliquoted to be used as input. Remaining peptide-bead complex was incubated with 1000 μg CB lysate for 6 hours at 4 °C. After incubation peptide-bead-protein complex was washed using lysis buffer (25 mM Tris-HCl pH 7.4, 150 mM NaCl, 1 mM EDTA, 1% NP-40 and 5% glycerol with protease inhibitor cocktail) three times. Finally, purified peptide-bead-protein complex was resuspended in 40 μl lysis buffer, mixed with 5x protein loading dye, and boiled for 5-7 minutes to break biotin streptavidin bond. CB lysate (100 μg, 10 %) was used as input, protein separation was done by 10% SDS-PAGE and analyzed by western blotting using 1:1000 dilution of rabbit anti-Wwox and anti-Hsp90 antibodies followed by 1:10000 anti-rabbit secondary antibody. Slot-blot assay was done to visualize 10% biotin labeled peptide-bead (~2kDa) input to show incubation with equal amounts of peptides. Slot-blots were incubated with 1:1000 rabbit anti-Biotin antibody and 1:10000 anti-rabbit secondary antibody.

### Mouse necropsy, histology, and blood hematology and chemistry

Mice were euthanized by CO2 asphyxiation at moribund stage. Blood was collected immediately following euthanasia by intracardiac puncture on the left ventricle. Fresh whole blood from each individual animal was used for hematological analysis on ABAXIS VETSCAN^®^ HM5 Hematology Analyzer. For blood chemistry, serum was collected by allowing the blood to clot at room temperature for 30 minute followed by centrifugation for 10 minutes at 14000 rpm. Serum from each individual mouse was analyzed using automated ABAXIS VETSCAN^®^ VS2 chemistry analyzer. Mouse organs were harvested and immediately weighed using an analytical balance. Mouse weight and blood analyses were done in *Wwox^WT/WT^* (n=9, 4 males and 5 females), *Wwox^P47T/WT^* (n=16, 13 males and 3 females), and *Wwox^P47T/P47T^* (n=16, 8 males and 8 females) mice. Organs were then fixed in formalin for 24-48 hours, followed by dehydration and embedding in paraffin. Tissue sections from *Wwox^WT/WT^* (n=7, 3 males and 4 females), *Wwox^P47T/WT^* (n=19, 14 males and 5 females), and *Wwox^P47T/P47T^* (n=21, 11 males and 10 females) mice were stained in H&E for histological analysis.

### Video Electroencephalography (EEG)

Mice were anesthetized with isoflurane (2.0–4% in oxygen, Patterson Veterinary Vaporizer), and silver wire electrodes (0.005” diameter) soldered to a connector were surgically implanted bilaterally into the subdural space over frontal and parietal cortex. Mice were allowed to recover for 14 days before recording. Simultaneous video-EEG and behavioral monitoring (Labchart 8.0, ADI Systems) was performed during 24-hour sessions in adult (aged >6 weeks) *Wwof^WT/WT^* (n=3) and *Wwox^P47T/P47T^* (n=3) mice of either sex. EEG was recorded while mice moved freely in the test cages. All EEG signals were amplified by a g.BSAMP biosignal amplifier (Austria), digitized by PowerLab with a 0.1 or 0.3 0.5 Hz high-pass and 60 Hz low-pass filter (ADInstruments, Dunedin, New Zealand) and acquired via Labchart 8.0 (ADInstruments). EEGs were reviewed by two trained observers.

### Behavioral testing

To analyze social, motor, and anxiety-like behavior, *Wwox^WT/WT^* (n=19, 7 males and 12 females), *Wwox^P47T/WT^* (n=17, 10 males and 7 females), and *Wwox^P47T/P47T^* (n=16, 8 males and 8 females) mice underwent a series of tests. The behavioral tests were performed from least to most stressful for the mouse in the order: OFT, JSET, TCST, EPM, and rotarod test. During these tests, the arenas were cleaned with 70% ethanol and allowed to air dry between each experimental mouse. These behavioral tasks were performed early during the light phase, and each test was spaced out by at least 24 hours, unless otherwise specified.

### Open Field Test

Mice were placed into an unfamiliar, inescapable square arena (50 cm x 50 cm) for 5 minutes. The central area is considered more anxiogenic relative to the edges. Anxiety-like behavior was measured as the amount of time spent in the center of the open field. Exploratory behavior was scored using automated tracking with EthoVision software (Noldus, Leesburg, VA).

### Juvenile Social Exploration Test

Immediately following the OFT, a juvenile mouse (sex-matched and 4 weeks old) was added to the arena for 5 minutes to observe for changes in social behavior. The amount of time the experimental mouse spent investigating the juvenile mouse (e.g., sniffing, pinning, allogrooming, and chasing) was manually scored by a genotype-blind investigator using EthoVision.

### Elevated Plus Maze

The apparatus used in this test was an opaque plexiglass maze lifted to a height of 40 cm with arms that were 5 cm wide and 30.5 cm long. The two closed arms were surrounded by walls that were 18.5 cm tall. Mice were placed onto the central platform facing a random open arm and tracker for 5 minutes. The open arms are considered more anxiogenic than the closed arms. The number of entries and total time spent in the open and closed arms were manually scored by a genotype-blind investigator using EthoVision software.

### Three-Chambered Sociability Test

A three-chambered social apparatus (120 cm x 40 cm x 40 cm) with two 7 cm x 15 cm stimulus cages were used as the testing arena. The experimental mouse was placed in the middle chamber of the apparatus, with doors to the two side chambers closed. A 10 minute habituation period was commenced when the doors to the side chambers were removed, allowing the mouse access to the entire apparatus. During the sociability test, one of the stimulus cages, randomly selected, held a novel mouse (sex-matched and 8 weeks old) while the other stimulus cage remained empty. The mouse was initially placed in the center chamber, and its interaction with the novel mouse vs. empty cage was recorded for 10 minutes. During the social novelty phase, the original stimulus mouse and a novel mouse (also sex-matched and 8 weeks old) were each placed into the stimulus cages. The experimental animal was then reintroduced to the center chamber and allowed to interact with the familiar and novel stimulus animals for ten minutes. Across the three phases of this behavioral task, the time spent in each chamber and time spent interacting with the novel or familiar mouse were automatically scored using EthoVision.

### Rotarod

The rotarod test was conducted to assess motor performance in mice. Each mouse performed this task three times with a 15-minute break in between tests. Mice were placed on the rotarod at 4 revolutions per minute (rpm) with an acceleration rate of 20 rpm/min. Rpm would automatically increase as long as the mice remained running. Two flanges on the mouse’s side prevent the mouse from leaving the rod. For each trial, the time until mice fell along with the distance traveled was recorded.

### Brain Tissue Preparation

Imaging experiments were performed using brain samples from *Wwox^WT/WT^* and *Wwox^P47T/P47T^* mice (n=3/group of either sex) at 80 and 250 days of age. For brain tissue collection, animals were deeply anesthetized and transcardially perfused with 4% paraformaldehyde (PFA) in PBS (pH 7.4). Perfused brains were collected and cerebrum was separated from CB. Both tissues were post fixed in 4% PFA for 24 hr at 4°C. Samples were then placed in PBS containing 15% sucrose for 4 hours at 4°C followed by immersion in 30% sucrose in PBS at 4°C until the tissues sank to the bottom of the tube. Prior to vibratome sectioning, CB tissues were stored in a solution containing 30% glycerol, 30% ethylene glycol and 10% 2x phosphate buffer at −20°C. Cerebrum was embedded and stored in Tissue-Tek OCT compound (Thermofisher) at −80°C. For immunofluorescent (IF) staining of markers in HPC, free floating (in 0.01M PBS w/ 0.01% sodium azide) 25 μm coronal cerebrum cryosections were collected with a Microm Leica CM3050S cryostat. For markers in CB, 65 μm free floating (in 0.01M PBS w/ 0.01% sodium azide) sagittal sections were cut using a Leica VT1200S vibratome.

### Immunofluorescence imaging

For immunostaining, floating tissue sections were permeabilized in 0.5% Triton X-100 in 1x PBS (PBT) for 30 min to 1h, followed by blocking for 1 h in a buffer containing 5% to 7% normal goat serum (NGS) or normal donkey serum (NDS). HPC sections were incubated at 4°C overnight and CB sections were incubated at 4°C for 48 h with marker specific primary antibodies diluted in blocking solution. After incubation with primary antibody, sections were washed four times while shaking in PBS with one wash containing 0.05% Tween-20 (PBST) followed by incubation with corresponding Alexa Fluor-488, or Alexa Fluor-568 conjugated secondary antibodies diluted in blocking buffer for 12 or 24 hr at 4°C. For the Iba1 antibody 0.1% Triton X-100 was used during permeabilization and Tween-20 was omitted from washes. For nuclear counterstain, tissues were treated with 10 ng of 4’,6-diamidino-2-phenylindole (DAPI) per ml for 2 or 4 hours at RT. After three additional washes tissues were mounted in glass dishes using ProLong Gold Antifade Mountant (Thermofisher), cover slipped and cured overnight. Labeled sections were visualized and imaged using Zeiss LSM 880 laser confocal microscope with 20x, 40x air objectives, or a 63x oil objective. The acquired images were processed, and pseudocolor images were generated using Zeiss Zen blue. Antibodies used were the following. Primary antibodies and final dilutions used were as follows: chicken anti-Gfap, 1:1000 (CPCA-Gfap); rabbit anti-Iba1, 1:1000 (RPCA-Iba1); mouse anti-calbindin, 1:1000 (MCA-467); rabbit anti-calretenin (RPCA-Calret) from Encor Biotechnology; rabbit anti-Olig2, 1:100 (ab109186) from Abcam; anti-Hcn1, 1:250 (PA5-78675) from Sigma.

### Image analyses

For analyses of HPC, Iba1, and Gfap immunostained confocal images (425 μm^2^) were acquired from the DG, CA1, and CA3 subfields at 40x magnification. All images for a specific marker in one age group were processed and acquired simultaneously by using identical confocal settings. Cell counting was done using dual label with DAPI. Activated astrocytes and microglia counts were determined by assessing the number of Iba1+ or Gfap+ cells per 425 μm^2^ image, respectively. AF analysis was done using various features of FIJI. First, the FFT Bandpass Filter was applied to remove background noise, the image was then converted into an 8-bit grayscale image and a despeckle step was done to eliminate the salt and pepper noise. The image was then auto-thresholded and was visually inspected to ensure that an accurate AF was obtained. Percentage AF of the resulting image was then calculated using analyze feature in FIJI. Cell count and AF analyses were done in all three regions per mouse, and with n=3 mice per experimental group. For microglia Sholl and skeleton analysis, images processed for AF analysis were taken and single microglial cell were isolated. The analysis was performed using the Sholl Analysis plug-in on FIJI. The start radius was defined at 2 μm from the center of the microglial cell soma, and the step size was set to 1 μm. Finally, the characteristics of traced microglia were measured using the analyze function in FIJI. Only representative microglia that had clear soma and processes were selected and analyzed. At least, one microglial cell per DG, CA1, and CA3 regions for each mouse was analyzed with n=3 mice per experimental group. Skeleton analysis was done using traced microglial cell created for Sholl analysis. First, the microglial cell images were skeletonized using the Skeletonize plug-in function in FIJI followed by measuring the features using the Analyze Skeleton plug-in function in FIJI. A cutoff value was set at 1 μm such that branches shorter than this were excluded from the analysis to avoid capturing small specks of noise that would artificially inflate branching. This plug-in generated values reflecting the total length and branching of microglial cells processes which were averaged to generate a single value per cells for statistical comparisons.

For analysis of CB degeneration markers, whole CB images and images from preculminate and primary fissures between cerebellar lobules III-VI were captured at 10x and 20x magnifications. The entire CB was scanned for regions of atrophy and PC degeneration and the exact comparative regions between *Wwox^WT/WT^* and *Wwox^P47T/P47T^* CB were imaged at 20x and 40x magnification. Few comparative regions with severe atrophy, PC loss and dendritic degeneration were also images at 63x magnification to capture finer morphological differences. All images for a specific marker in one age group were processed and acquired simultaneously by using identical confocal settings. Calbindin+ PC and Hcn1+ basket cell counting was done using dual labelled images in comparative 1000 μm^2^ sections from *Wwox^WT/WT^* and *Wwox^P47T/P47T^* pairs (n=3). The quantitation of different sections from animals with same genotype and age were averaged and plotted using GraphPad Prism 9. Measurements of ML thickness were done using calbindin staining by measuring the perpendicular distance from the base of the PC body to the end of the dendrite at the pial surface and GL thickness was measured based on calretinin and DAPI dual staining. Twenty four measurements, with 12 measurements each in preculminate and primary fissures per animal were taken. For oligodendrocyte quantitation oligodendrocyte transcription factor 2 (Olig2) stained images (1 x 1.5 mm^2^) were captured in the corpus callosum and cells were counted in three independent 500 μm^2^ section spanning the entire imaged corpus callosum region. All quantitative data was averaged from three mice per group and plotted using GraphPad Prism 9.

### RNA-Seq and data analysis

RNA was isolated from HPC, PFC, (parietal) CTX, and CB from *Wwox^WT/WT^* and *Wwox^P47T/P47T^* mice, 150-280 days of age, n=5 mice/group (2 males and 3 females). Mice were euthanized, brains were extracted, and different tissue regions were collected on a cold plate and flash frozen in liquid N2. Total RNA was extracted using TRIzol reagent (Invitrogen) following standard manufacturer’s protocol. RNA concentration and integrity were measured on an Agilent 2100 Bioanalyzer (Agilent Technologies). RNA samples with RNA integrity number (RIN) over 8.0 were considered for RNA sequencing. The samples were processed for directional mRNA-seq library construction using the ScriptSeq v2 RNA-Seq Library Preparation Kit (Epicentre) according to the manufacturer’s protocol. We performed 76 nt paired-end sequencing using an Illumina HiSeq3000 platform and obtained ~40 million tags per sample. The short-sequenced reads were mapped to the mouse reference genome (mm10) by the splice junction aligner TopHat V2.0.10. We employed several R/Bioconductor packages to calculate gene expression abundance at the whole-genome level using the aligned records (BAM files) and to identify differentially expressed genes between *Wwox^WT/WT^* and *Wwox^P47T/P47T^* mice. Briefly, to identify differentially expressed genes between wildtype and *Wwox-P47T* homozygous mutant samples, we computed fold change using the EdgeR Bioconductor package based on the normalized log2 based count per million values [58]. Data integration and visualization of differentially expressed transcripts between *Wwox^WT/WT^* and *Wwox^P47T/P47T^* mice brain regions (log2 fold change [log2 FC] > ±1, *p*-value <.05, and false discovery rate [FDR] <0.05) was done with the Multiexperiment Viewer Software. Functional enrichment analysis of dysregulated transcripts was performed using the Ingenuity Pathway Analysis (IPA, Qiagen), and gene set enrichment analysis (GSEA) [59] with R/Bioconductor packages clusterProfiler and enrichplot [60]. The R package BRETIGEA (**<u>Br</u>**ain c**<u>e</u>**ll **<u>t</u>**ype specific **<u>g</u>**ene **<u>e</u>**xpression **<u>a</u>**nalysis) was employed to estimate relative brain cell type proportions from the RNA-seq profiles [26]. Functional enrichment analysis of dysregulated transcripts was performed using the Ingenuity Pathway Analysis (IPA) database (http://www.ingenuity.com/index.html).

### Quantitative RT-PCR (qRT-PCR)

Total RNA was extracted and quantified from *Wwox^WT/WT^* and *Wwox^P47T/P47T^* n=6 mice/group (3 males and 3 females) and cDNA was synthesized using High-Capacity cDNA Reverse Transcription Kit (Applied Biosystems) following manufacturer’s instructions. The relative expression level for specific genes was determined in triplicate by qRT-PCR using the SYBR Green-based method. After normalization to 18s RNA expression, the average fold change between *Wwox^WT/WT^* and *Wwox^P47T/P47T^* samples was calculated using the 2^-(ΔΔCt) method described elsewhere [61]. Following primers were used for qRT-PCR-*IL1β* F-5’-GAAATGCCACCTTTTGACAGTG-3’, R-5’-TGGATGCTCTCATCAGGACAG-3’; *TNFα* F-5’-CCTGTAGCCCACGTCGTAG-3’, R-5’-GGGAGTAGACAAGGTACAACCC-3’; and 18s 5’-GCAATTATTCCCCATGAACG-3’, R-5’-GGCCTCACTAAACCATCCAA-3’.

### Statistical analyses

Mouse survival analysis was performed using SPSS statistical software by the Kaplan-Meier method and Log-rank test was applied to compare survival curves. Statistical significance analysis for comparison of means from two groups was done by Student’s two-tailed unpaired *t*-test when data followed normal distribution, and by Mann-Whitney U test for data with non-normal distribution. For comparison of means from more than two groups, One-Way or Two-Way ANOVA with Tukey’s post-hoc test was performed. Generation of quantitation graphs and statistical analyses was done using GraphPad Prism 9. All data are presented as mean ± SEM and differences between the group means were considered statistically significant at p < 0.05.

## Acknowledgements

Authors would like to acknowledge funding by National Ataxia Foundation, FF2022-00060843-HM, The UT Austin/MD Anderson Cancer Center Collaboration 00060803-Y1, The CPRIT Core Facility Support Award (RP170628) to C.M.A., R01AG078758 and R01AG062716 to L.F., and NINDS NS29709 and Blue Bird Circle Foundation to J.L.N. Authors would like to thank the University of Texas MD Anderson Cancer Center (UTMDACC) Research Animal Support Facility and Laboratory Animal Genetic Services for animal support and Advanced Technology Genomics Core (ATGC) for sequencing facility (P30 NIH CA016672).

## Author contributions

T.H.- Designed and performed experiments, data analysis, manuscript writing; K.S.- designed and analyzed behavioral tests; J.C.- confocal imaging; D.S.- oligopeptide pull-down assay; C.J.- confocal imaging; Y.L.- bioinformatics analysis; M.A.- bioinformatics analysis; R.S.- video EEG; J.L.N.- video EEG; L.F.- experimental designing, intellectual input for improving the manuscript; C.M.A.- conceptualized the study, experimental designing, manuscript writing, correspondence

## Materials & Correspondence

T.H.; C.M.A

