## Supplementary material for "WWOX P47T loss-of-function mutation induces epilepsy, progressive neuroinflammation, and cerebellar degeneration in mice phenocopying human SCAR12": Wwox-P47T_Supplementary File 1.pdf

### PFC

| ID | Symbol | Expr Log Ratio | Expr p-value | Expr FDR |
|---|---|---|---|---|
| ENSMUSG00000059631.6 | 1500035N22Rik | -1.132844783 | 0.00120726 | 0.021744876 |
| ENSMUSG00000085549.7 | 1700047F07Rik | -1.859341806 | 9.81E-06 | 4.98E-04 |
| ENSMUSG00000097178.7 | 2310002F09Rik | 2.095548668 | 2.73E-09 | 4.79E-07 |
| ENSMUSG00000085338.1 | 2410004I01Rik | 1.560757152 | 2.02E-05 | 8.90E-04 |
| ENSMUSG00000115625.1 | 2900040C04Rik | -3.622405805 | 0.00125646 | 0.022359118 |
| ENSMUSG00000085129.1 | 5031425F14Rik | 3.586794549 | 4.54E-09 | 7.01E-07 |
| ENSMUSG00000086822.2 | 5330413P13Rik | 1.017646711 | 1.20E-05 | 5.87E-04 |
| ENSMUSG00000110148.1 | 5830408C22Rik | 1.638016781 | 1.25E-04 | 0.003843749 |
| ENSMUSG00000034185.9 | 6430628N08Rik | 3.187290888 | 1.43E-04 | 0.004264874 |
| ENSMUSG00000098097.7 | 6530403H02Rik | 1.25935197 | 6.18E-04 | 0.013135621 |
| ENSMUSG00000118154.1 | 9330117O12Rik | -1.411988591 | 4.94E-06 | 2.89E-04 |
| ENSMUSG00000108242.1 | 9330118I20Rik | 1.042547389 | 1.53E-06 | 1.11E-04 |
| ENSMUSG00000117916.1 | 9630028I04Rik | 2.402506269 | 5.37E-15 | 4.43E-12 |
| ENSMUSG00000105891.4 | A230001M10Rik | -1.256567201 | 1.08E-04 | 0.003412543 |
| ENSMUSG00000089633.2 | A230009B12Rik | -1.281357011 | 6.63E-04 | 0.013846701 |
| ENSMUSG00000092071.1 | A230065N10Rik | 1.019809401 | 3.07E-04 | 0.007626352 |
| ENSMUSG00000097666.2 | A330094K24Rik | 1.110858286 | 0.00148587 | 0.02527058 |
| ENSMUSG00000047878.13 | A4GALT | 1.155667345 | 2.72E-05 | 0.001124423 |
| ENSMUSG00000010651.4 | Acaa1b | 2.318567621 | 8.62E-24 | 1.78E-20 |
| ENSMUSG00000030607.7 | Acan | 1.996615144 | 0.0028514 | 0.040418961 |
| ENSMUSG00000001348.15 | Acp5 | 1.288912122 | 2.23E-04 | 0.005971099 |
| ENSMUSG00000006457.4 | Actn3 | 1.011017842 | 3.48E-04 | 0.00834415 |
| ENSMUSG00000011256.16 | ADAM19 | 1.207607119 | 6.32E-07 | 5.22E-05 |
| ENSMUSG00000053399.9 | ADAMTS18 | 1.253589837 | 0.00131083 | 0.02309644 |
| ENSMUSG00000032363.15 | ADAMTS7 | 1.182462505 | 7.27E-04 | 0.014724099 |
| ENSMUSG00000031994.13 | ADAMTS8 | 1.01496584 | 2.66E-04 | 0.006778487 |
| ENSMUSG00000024256.7 | ADCYAP1 | 1.850854982 | 4.49E-13 | 2.25E-10 |
| ENSMUSG00000023918.12 | ADGRF4 | 1.872689017 | 1.23E-06 | 9.08E-05 |
| ENSMUSG00000040706.4 | AGMAT | 1.783005713 | 3.71E-07 | 3.33E-05 |
| ENSMUSG00000084826.7 | AI847159 | 1.39875185 | 2.09E-04 | 0.005734581 |
| ENSMUSG00000021214.14 | AKR1C3 | 1.561178059 | 1.90E-06 | 1.32E-04 |
| ENSMUSG00000019102.10 | ALDH3A1 | 3.338932009 | 1.02E-29 | 5.60E-26 |
| ENSMUSG00000054204.6 | ALKAL2 | 2.088198076 | 4.07E-04 | 0.009540966 |
| ENSMUSG00000032807.5 | ALOX12B | -1.808889982 | 2.38E-07 | 2.39E-05 |
| ENSMUSG00000025701.12 | ALOX5 | 2.691938935 | 5.20E-14 | 3.43E-11 |
| ENSMUSG00000031465.6 | ANGPT2 | 1.707566741 | 5.87E-04 | 0.012660851 |
| ENSMUSG00000074771.10 | ANKEF1 | 1.054716182 | 4.72E-04 | 0.010777726 |
| ENSMUSG00000078137.1 | ANKRD63 | -1.405625263 | 0.00184839 | 0.02943516 |
| ENSMUSG00000068246.6 | Apol9a/Apol9b | 1.852140836 | 5.80E-04 | 0.012612575 |
| ENSMUSG00000051517.14 | ARHGEF39 | 1.572610443 | 8.44E-05 | 0.00278268 |
| ENSMUSG00000021200.14 | ASB2 | 1.504270818 | 1.84E-11 | 5.74E-09 |
| ENSMUSG00000020052.9 | ASCL1 | -1.029585785 | 2.75E-05 | 0.001134276 |
| ENSMUSG00000009248.6 | ASCL2 | 4.121589286 | 9.33E-06 | 4.85E-04 |

| | | | | |
|---|---|---|---|---|
| ENSMUSG00000037686.6 | ASPG | 1.60635275 | 1.17E-06 | 8.74E-05 |
| ENSMUSG00000030730.12 | ATP2A1 | 3.722690083 | 6.93E-17 | 7.93E-14 |
| ENSMUSG00000034112.9 | ATP2C2 | 2.119581536 | 3.47E-06 | 2.17E-04 |
| ENSMUSG00000020566.19 | ATP6V1C2 | 1.158831083 | 0.00144206 | 0.024754865 |
| ENSMUSG00000079445.3 | B3GNT7 | 2.471744346 | 6.30E-11 | 1.68E-08 |
| ENSMUSG00000059479.7 | B3GNT8 | 1.793083615 | 1.26E-14 | 9.49E-12 |
| ENSMUSG00000046415.11 | B430212C06Rik | 1.294492758 | 6.30E-04 | 0.01331416 |
| ENSMUSG00000041372.10 | B4GALNT3 | -1.114197518 | 0.00266625 | 0.038725055 |
| ENSMUSG00000018126.15 | BAIAP2L2 | -1.204894331 | 1.11E-09 | 2.18E-07 |
| ENSMUSG00000047507.12 | BAIAP3 | -1.221426763 | 0.00126359 | 0.02243751 |
| ENSMUSG00000035021.13 | BAZ1A | 1.88208024 | 2.19E-04 | 0.005893161 |
| ENSMUSG00000085041.1 | BB031773 | 1.369172911 | 5.18E-04 | 0.011563102 |
| ENSMUSG00000040363.15 | BCOR | 1.067664614 | 0.00178226 | 0.028714304 |
| ENSMUSG00000048482.14 | BDNF | 2.860645998 | 4.46E-08 | 5.50E-06 |
| ENSMUSG00000061132.13 | BLNK | 1.221046164 | 3.59E-05 | 0.001379024 |
| ENSMUSG00000027485.15 | BPIFB1 | 2.670640628 | 1.84E-09 | 3.45E-07 |
| ENSMUSG00000053216.15 | BTN2A2 | -1.49198979 | 1.42E-06 | 1.04E-04 |
| ENSMUSG00000071317.4 | BVES | 1.422082035 | 2.23E-04 | 0.005968433 |
| ENSMUSG00000047642.15 | C12orf56 | 1.653962974 | 5.52E-06 | 3.17E-04 |
| ENSMUSG00000036907.9 | C1QL2 | 3.289592906 | 2.00E-33 | 3.30E-29 |
| ENSMUSG00000050157.6 | C22orf15 | 1.288873654 | 7.52E-05 | 0.002525091 |
| ENSMUSG00000090066.2 | C4orf54 | 1.12848211 | 5.47E-04 | 0.012015772 |
| ENSMUSG00000033053.4 | C9orf135 | 1.300800028 | 2.11E-04 | 0.005767186 |
| ENSMUSG00000010066.15 | CACNA2D2 | 1.04667228 | 3.64E-09 | 5.99E-07 |
| ENSMUSG00000023964.15 | CALCR | 2.418726592 | 0.00139488 | 0.024221944 |
| ENSMUSG00000032246.14 | CALML4 | -2.308880746 | 0.00236959 | 0.03557403 |
| ENSMUSG00000020424.3 | CASTOR1 | 1.032557986 | 4.88E-07 | 4.24E-05 |
| ENSMUSG00000028348.7 | CAVIN4 | 1.491592322 | 0.00191857 | 0.030302518 |
| ENSMUSG00000046318.16 | CCBE1 | -1.217201397 | 4.11E-07 | 3.65E-05 |
| ENSMUSG00000044033.16 | CCDC141 | 1.041918082 | 1.57E-07 | 1.69E-05 |
| ENSMUSG00000047810.9 | CCDC88B | 1.864551101 | 3.27E-20 | 5.40E-17 |
| ENSMUSG00000072082.7 | CCNF | 1.564518714 | 2.06E-05 | 9.03E-04 |
| ENSMUSG00000008845.9 | CD163 | 1.275843405 | 0.00175879 | 0.02852402 |
| ENSMUSG00000028865.14 | CD164L2 | 1.321028369 | 7.69E-09 | 1.11E-06 |
| ENSMUSG00000028076.12 | CD1D | 1.901282095 | 4.82E-06 | 2.84E-04 |
| ENSMUSG00000022667.18 | CD200R1 | 1.537044655 | 6.53E-05 | 0.002274502 |
| ENSMUSG00000026012.2 | CD28 | 2.971156461 | 4.53E-12 | 1.74E-09 |
| ENSMUSG00000016494.9 | CD34 | -1.177500148 | 2.51E-13 | 1.34E-10 |
| ENSMUSG00000012819.15 | CDH23 | 1.024718744 | 7.19E-04 | 0.014692731 |
| ENSMUSG00000034918.8 | CDHR2 | 1.323044289 | 1.02E-05 | 5.15E-04 |
| ENSMUSG00000023067.14 | CDKN1A | 2.213645865 | 5.41E-05 | 0.001955561 |
| ENSMUSG00000037664.13 | Cdkn1c | -1.163041464 | 5.24E-05 | 0.001904307 |
| ENSMUSG00000037628.10 | CDKN3 | 1.906047297 | 0.00119224 | 0.021534039 |
| ENSMUSG00000076433.4 | CEP295NL | 1.906547114 | 2.86E-04 | 0.007218996 |
| ENSMUSG00000028294.15 | CFAP206 | -1.053005376 | 0.00156779 | 0.026258079 |

| | | | | |
|---|---|---|---|---|
| ENSMUSG00000021194.6 | CHGA | 1.265988268 | 3.71E-21 | 6.81E-18 |
| ENSMUSG00000022041.10 | CHRNA2 | 1.476696025 | 3.00E-05 | 0.001207913 |
| ENSMUSG00000042269.15 | CIBAR2 | 1.601789719 | 1.41E-06 | 1.04E-04 |
| ENSMUSG00000070803.6 | CITED4 | -1.357895607 | 1.45E-11 | 4.71E-09 |
| ENSMUSG00000047230.6 | CLDN2 | -3.016325007 | 5.88E-04 | 0.012666523 |
| ENSMUSG00000033633.15 | CLEC18B | 1.671302088 | 6.22E-07 | 5.21E-05 |
| ENSMUSG00000022949.9 | CLIC6 | -2.099298509 | 9.50E-04 | 0.018121772 |
| ENSMUSG00000039462.4 | COL10A1 | 2.229703204 | 0.00232242 | 0.034929357 |
| ENSMUSG00000001435.15 | COL18A1 | 1.207121778 | 4.03E-06 | 2.44E-04 |
| ENSMUSG00000004098.7 | COL5A3 | 1.916989903 | 6.62E-06 | 3.69E-04 |
| ENSMUSG00000057606.14 | COLQ | 1.298198759 | 9.79E-05 | 0.003165243 |
| ENSMUSG00000039070.5 | CPA4 | 1.600701148 | 2.48E-04 | 0.006501591 |
| ENSMUSG00000021680.8 | CRHBP | 1.476019204 | 1.31E-08 | 1.80E-06 |
| ENSMUSG00000018634.10 | CRHR1 | -1.122293251 | 3.53E-07 | 3.22E-05 |
| ENSMUSG00000007888.15 | CRLF1 | 2.035794907 | 6.86E-12 | 2.44E-09 |
| ENSMUSG00000006546.3 | CRYBA2 | 2.42399611 | 5.49E-06 | 3.16E-04 |
| ENSMUSG00000012123.17 | CRYBG2 | 1.263944114 | 3.03E-05 | 0.001212807 |
| ENSMUSG00000026456.18 | CYB5R1 | 1.087788502 | 1.41E-10 | 3.41E-08 |
| ENSMUSG00000024087.4 | CYP1B1 | 1.977791132 | 2.62E-06 | 1.73E-04 |
| ENSMUSG00000063415.12 | CYP26B1 | 1.257543411 | 4.63E-04 | 0.010660332 |
| ENSMUSG00000062563.15 | CYS1 | -1.116459517 | 2.96E-05 | 0.001200982 |
| ENSMUSG00000109198.1 | D7Bwg0826e | 2.033621015 | 1.03E-06 | 7.93E-05 |
| ENSMUSG00000035910.15 | DCDC2 | 1.373412123 | 3.82E-12 | 1.54E-09 |
| ENSMUSG00000027068.6 | DHRS9 | 2.267899817 | 8.78E-04 | 0.017083743 |
| ENSMUSG00000075707.5 | DIO3 | -2.035897685 | 2.74E-05 | 0.001130999 |
| ENSMUSG00000020871.8 | DLX4 | -1.382328922 | 0.00131293 | 0.02309644 |
| ENSMUSG00000086552.3 | Dlx4os | -3.846263655 | 2.07E-10 | 4.82E-08 |
| ENSMUSG00000060962.12 | DMKN | 2.063674442 | 1.86E-08 | 2.44E-06 |
| ENSMUSG00000000730.13 | DNMT3L | 2.389036386 | 6.76E-09 | 1.01E-06 |
| ENSMUSG00000042737.9 | DPM3 | 1.259402396 | 5.65E-06 | 3.22E-04 |
| ENSMUSG00000031786.7 | DRC7 | -2.168537148 | 5.36E-04 | 0.011887789 |
| ENSMUSG00000025496.5 | DRD4 | 3.493277588 | 3.57E-07 | 3.22E-05 |
| ENSMUSG00000054889.10 | DSP | 2.139003753 | 4.46E-11 | 1.27E-08 |
| ENSMUSG00000026247.13 | ECEL1 | 1.835456539 | 8.11E-04 | 0.016013867 |
| ENSMUSG00000026051.8 | ECRG4 | -3.101838576 | 5.45E-04 | 0.012002062 |
| ENSMUSG00000022441.17 | EFCAB6 | 3.018163564 | 6.10E-10 | 1.27E-07 |
| ENSMUSG00000043439.5 | EPOP | -1.144057126 | 2.88E-06 | 1.87E-04 |
| ENSMUSG00000019768.16 | ESR1 | 1.178770253 | 0.00194578 | 0.030602554 |
| ENSMUSG00000021255.17 | ESRRB | 1.351234045 | 2.73E-06 | 1.78E-04 |
| ENSMUSG00000003382.18 | ETV3 | 1.223037386 | 1.62E-04 | 0.004672931 |
| ENSMUSG00000118505.1 | ETV3L | 5.964695746 | 8.24E-06 | 4.39E-04 |
| ENSMUSG00000021675.4 | F2RL2 | 1.481708068 | 6.56E-05 | 0.002280322 |
| ENSMUSG00000089665.2 | Fcor | 1.055951554 | 2.53E-05 | 0.001058393 |
| ENSMUSG00000070504.9 | FCRL6 | 3.278976967 | 0.00202419 | 0.031535284 |
| ENSMUSG00000021732.14 | FGF10 | -1.997320952 | 6.93E-12 | 2.44E-09 |

| | | | | |
|---|---|---|---|---|
| ENSMUSG00000028874.14 | FGR | 4.917389123 | 8.78E-26 | 2.07E-22 |
| ENSMUSG00000051435.11 | FHAD1 | 1.162909011 | 4.41E-08 | 5.47E-06 |
| ENSMUSG00000045326.13 | FNDC7 | 2.168123636 | 3.26E-06 | 2.07E-04 |
| ENSMUSG00000048721.2 | FNDC9 | 3.430700482 | 9.54E-06 | 4.91E-04 |
| ENSMUSG00000001827.12 | FOLR1 | -3.570751576 | 5.39E-04 | 0.011934692 |
| ENSMUSG00000027004.3 | FRZB | -1.424628632 | 1.38E-09 | 2.64E-07 |
| ENSMUSG00000031340.8 | Gabre | 1.684617435 | 0.00160621 | 0.026721938 |
| ENSMUSG00000056880.12 | GADL1 | 1.111035145 | 0.00150203 | 0.025496594 |
| ENSMUSG00000024907.7 | GAL | 3.805194328 | 9.47E-07 | 7.37E-05 |
| ENSMUSG00000067724.5 | GBX1 | 2.916739117 | 3.39E-08 | 4.28E-06 |
| ENSMUSG00000051136.1 | GHSR | 3.762876712 | 7.30E-06 | 3.98E-04 |
| ENSMUSG00000048582.7 | GJA3 | 2.842681416 | 1.99E-05 | 8.80E-04 |
| ENSMUSG00000047197.1 | GJD3 | 1.14445411 | 3.06E-05 | 0.001222219 |
| ENSMUSG00000000263.15 | GLRA1 | 3.833130013 | 5.10E-12 | 1.91E-09 |
| ENSMUSG00000075437.4 | Gm11681 | -1.339759411 | 0.0015243 | 0.025633761 |
| ENSMUSG00000086196.7 | Gm13571 | 2.599289283 | 0.00278963 | 0.039989482 |
| ENSMUSG00000085225.1 | Gm14637 | 3.49352277 | 3.04E-05 | 0.00121653 |
| ENSMUSG00000091177.2 | Gm15494 | -1.107831167 | 0.00117661 | 0.021328712 |
| ENSMUSG00000087652.1 | Gm15918 | 1.168508613 | 0.00328035 | 0.044844159 |
| ENSMUSG00000085055.2 | Gm15958 | -1.484117794 | 0.00112941 | 0.020792796 |
| ENSMUSG00000087411.1 | Gm16048 | 2.458123185 | 8.81E-11 | 2.24E-08 |
| ENSMUSG00000090050.1 | Gm16339 | 1.006352591 | 3.48E-04 | 0.00834415 |
| ENSMUSG00000083218.2 | Gm16425 | 1.712659361 | 3.32E-07 | 3.08E-05 |
| ENSMUSG00000102548.1 | Gm16701 | 1.256470487 | 3.43E-05 | 0.001331608 |
| ENSMUSG00000086289.3 | Gm16933 | 1.265193319 | 0.00237959 | 0.03569174 |
| ENSMUSG00000091542.1 | Gm17167 (includes | -1.27112324 | 1.35E-04 | 0.004084026 |
| ENSMUSG00000079669.2 | Gm17396 | 1.192931175 | 5.55E-09 | 8.40E-07 |
| ENSMUSG00000097183.2 | Gm17501 | 1.466923743 | 0.00216869 | 0.033130247 |
| ENSMUSG00000117767.1 | Gm18786 | 1.258983562 | 6.86E-05 | 0.002356389 |
| ENSMUSG00000098088.1 | Gm26916 | 1.354001483 | 2.88E-04 | 0.00725723 |
| ENSMUSG00000099644.1 | Gm28626 | 3.175999928 | 2.34E-09 | 4.19E-07 |
| ENSMUSG00000100980.1 | Gm29100 | 1.78764662 | 0.00352995 | 0.047393167 |
| ENSMUSG00000100546.1 | Gm29483 | 1.000409931 | 2.50E-04 | 0.006524722 |
| ENSMUSG00000100596.1 | Gm29502 | 1.304132119 | 3.62E-04 | 0.008621233 |
| ENSMUSG00000112035.1 | Gm30539 | 1.465089576 | 1.04E-04 | 0.003303799 |
| ENSMUSG00000109844.2 | Gm31816 | 1.119516989 | 0.00179492 | 0.028860866 |
| ENSMUSG00000114271.1 | Gm32224 | 1.167939438 | 0.00344171 | 0.046587253 |
| ENSMUSG00000117254.1 | Gm34567 | 1.472646672 | 2.43E-05 | 0.001025662 |
| ENSMUSG00000107272.1 | Gm34583 | 1.435625533 | 2.77E-05 | 0.001137886 |
| ENSMUSG00000116114.1 | Gm35853 | 2.276885625 | 4.02E-06 | 2.44E-04 |
| ENSMUSG00000110179.1 | Gm39168 | 2.498639001 | 5.64E-09 | 8.46E-07 |
| ENSMUSG00000109868.1 | Gm39244 | 1.150554826 | 2.18E-05 | 9.47E-04 |
| ENSMUSG00000113845.1 | Gm39473 | -1.796730805 | 0.00259234 | 0.037985765 |
| ENSMUSG00000106651.1 | Gm42608 | 1.001581607 | 3.31E-04 | 0.008042684 |
| ENSMUSG00000105152.1 | Gm42864 | -2.331980289 | 1.35E-12 | 6.02E-10 |

| | | | | |
|---|---|---|---|---|
| ENSMUSG00000104807.1 | Gm42865 | -2.334396204 | 2.37E-07 | 2.39E-05 |
| ENSMUSG00000104850.1 | Gm42901 | 1.524843776 | 5.58E-06 | 3.19E-04 |
| ENSMUSG00000086804.8 | Gm43154 | 2.20006339 | 2.02E-12 | 8.79E-10 |
| ENSMUSG00000097532.2 | Gm4349 | -1.170133643 | 7.14E-04 | 0.014623515 |
| ENSMUSG00000109130.1 | Gm45187 | -1.72120635 | 1.73E-06 | 1.21E-04 |
| ENSMUSG00000109604.1 | Gm45346 | 2.109428303 | 7.22E-06 | 3.95E-04 |
| ENSMUSG00000109957.1 | Gm45353 | 1.274530814 | 5.43E-05 | 0.001956126 |
| ENSMUSG00000114544.1 | Gm49146 | -1.548584972 | 2.35E-05 | 0.001002441 |
| ENSMUSG00000081436.1 | Gm5912 | 1.478583815 | 4.80E-04 | 0.010873131 |
| ENSMUSG00000073532.3 | Gm7276 | 1.030488922 | 0.00152613 | 0.025638334 |
| ENSMUSG00000041468.7 | GPR12 | -1.116763359 | 4.98E-06 | 2.90E-04 |
| ENSMUSG00000066197.5 | GPR139 | 2.07302365 | 3.41E-09 | 5.75E-07 |
| ENSMUSG00000045509.7 | GPR150 | -1.702652653 | 3.16E-08 | 4.01E-06 |
| ENSMUSG00000049649.8 | GPR3 | 1.393489142 | 0.00340401 | 0.04615252 |
| ENSMUSG00000063234.5 | GPR84 | 2.434596603 | 5.83E-05 | 0.002076032 |
| ENSMUSG00000051043.16 | GPRC5C | 2.211337594 | 3.57E-09 | 5.95E-07 |
| ENSMUSG00000024517.17 | GRP | -2.571496855 | 9.55E-10 | 1.90E-07 |
| ENSMUSG00000042638.14 | GUCY2C | 1.521289328 | 1.14E-04 | 0.003557624 |
| ENSMUSG00000020890.11 | GUCY2D | 1.196199158 | 2.27E-04 | 0.006052249 |
| ENSMUSG00000030074.9 | GXYLT2 | 1.321162156 | 3.32E-05 | 0.001300841 |
| ENSMUSG00000020295.1 | Hbq1a | -1.709177403 | 9.87E-08 | 1.12E-05 |
| ENSMUSG00000032338.9 | HCN4 | 1.66631035 | 2.16E-09 | 3.91E-07 |
| ENSMUSG00000047171.8 | HELT | 2.091779007 | 1.69E-04 | 0.004835067 |
| ENSMUSG00000021260.8 | HHIPL1 | 1.253358252 | 2.03E-11 | 6.22E-09 |
| ENSMUSG00000055632.18 | HMCN2 | 3.275760793 | 1.92E-09 | 3.56E-07 |
| ENSMUSG00000031722.10 | HP | 2.126539728 | 7.40E-04 | 0.014914754 |
| ENSMUSG00000043155.4 | HPDL | 1.293080642 | 3.17E-04 | 0.007811295 |
| ENSMUSG00000051456.4 | HSPB3 | 1.814438951 | 6.13E-07 | 5.19E-05 |
| ENSMUSG00000006221.7 | HSPB7 | 1.877483125 | 5.98E-06 | 3.38E-04 |
| ENSMUSG00000037406.7 | HTRA4 | 2.954171815 | 4.10E-10 | 9.03E-08 |
| ENSMUSG00000037405.8 | ICAM1 | 1.756927638 | 1.58E-05 | 7.29E-04 |
| ENSMUSG00000000732.9 | ICOSLG/LOC102723 | 1.019479888 | 3.57E-04 | 0.008521612 |
| ENSMUSG00000032394.6 | IGDCC3 | 1.005651816 | 9.08E-05 | 0.002962143 |
| ENSMUSG00000076614.7 | IGHG1 | 6.239034261 | 1.81E-13 | 9.98E-11 |
| ENSMUSG00000037995.15 | IGSF9 | 1.367232755 | 9.37E-06 | 4.85E-04 |
| ENSMUSG00000015966.17 | IL17RB | 3.573991989 | 5.32E-10 | 1.16E-07 |
| ENSMUSG00000041324.14 | INHBA | 2.54705801 | 1.38E-04 | 0.004165648 |
| ENSMUSG00000025498.15 | IRF7 | 1.140186371 | 8.00E-04 | 0.015819538 |
| ENSMUSG00000032243.8 | ITGA11 | 1.744851914 | 2.47E-06 | 1.64E-04 |
| ENSMUSG00000117975.1 | ITPRIP | 1.644586091 | 1.75E-04 | 0.004950581 |
| ENSMUSG00000048534.7 | JAML | 1.852337761 | 3.79E-18 | 5.22E-15 |
| ENSMUSG00000020216.13 | JSRP1 | 1.58433294 | 1.44E-04 | 0.004286594 |
| ENSMUSG00000035407.8 | KANK4 | 1.327414132 | 4.15E-12 | 1.63E-09 |
| ENSMUSG00000039672.12 | KCNE2 | -4.592015907 | 1.13E-04 | 0.003518666 |
| ENSMUSG00000074575.4 | KCNG1 | -1.071117177 | 9.56E-05 | 0.003106901 |

| | | | | |
|---|---|---|---|---|
| ENSMUSG00000045246.11 | KCNG4 | 1.192410918 | 2.00E-07 | 2.07E-05 |
| ENSMUSG00000025221.16 | KCNIP2 | -1.071174864 | 6.26E-07 | 5.22E-05 |
| ENSMUSG00000009545.14 | KCNQ1 | 1.907395145 | 4.68E-06 | 2.78E-04 |
| ENSMUSG00000042514.11 | KLHL14 | -1.198186224 | 1.01E-04 | 0.003238445 |
| ENSMUSG00000074001.3 | KLHL40 | 1.736790324 | 1.81E-04 | 0.005121257 |
| ENSMUSG00000108444.1 | Klk2-ps | 3.468026272 | 1.09E-10 | 2.68E-08 |
| ENSMUSG00000064023.4 | KLK8 | 1.454887804 | 1.16E-07 | 1.29E-05 |
| ENSMUSG00000046834.7 | KRT1 | 1.10092049 | 1.11E-05 | 5.54E-04 |
| ENSMUSG00000023043.7 | KRT18 | -2.526764897 | 0.00339024 | 0.046036614 |
| ENSMUSG00000064201.8 | Krt2 | 2.261806064 | 7.48E-09 | 1.09E-06 |
| ENSMUSG00000061527.7 | KRT5 | 2.269663725 | 0.00176275 | 0.028531777 |
| ENSMUSG00000023039.17 | KRT7 | 1.829492746 | 2.07E-04 | 0.005687754 |
| ENSMUSG00000022986.5 | KRT75 | 5.376420038 | 1.08E-07 | 1.21E-05 |
| ENSMUSG00000041700.9 | LHFPL1 | 2.836579964 | 3.01E-06 | 1.93E-04 |
| ENSMUSG00000026934.15 | LHX3 | 4.069194793 | 3.38E-12 | 1.39E-09 |
| ENSMUSG00000053846.5 | LIPG | 1.586192275 | 4.23E-04 | 0.009856373 |
| ENSMUSG00000020782.18 | LLGL2 | 1.494917349 | 6.54E-08 | 7.66E-06 |
| ENSMUSG00000054966.13 | LMNTD1 | 1.22255195 | 4.28E-04 | 0.009938077 |
| ENSMUSG00000026686.14 | LMX1A | -2.957654913 | 8.51E-04 | 0.016650669 |
| ENSMUSG00000022759.15 | Lrrc74b | 2.085954345 | 5.06E-11 | 1.39E-08 |
| ENSMUSG00000044847.13 | LSM11 | 1.306249248 | 2.50E-07 | 2.44E-05 |
| ENSMUSG00000018819.10 | LSP1 | 1.011717832 | 6.31E-06 | 3.56E-04 |
| ENSMUSG00000013766.11 | Ly6g6e | 1.916945325 | 1.47E-04 | 0.00436084 |
| ENSMUSG00000032530.14 | LYZL4 | 2.433873908 | 9.82E-05 | 0.00316582 |
| ENSMUSG00000044313.13 | MAB21L3 | 3.317767949 | 3.88E-06 | 2.38E-04 |
| ENSMUSG00000047591.5 | MAFA | 1.219846489 | 2.55E-05 | 0.001062281 |
| ENSMUSG00000061143.15 | MAML3 | 1.176833895 | 1.49E-04 | 0.004391207 |
| ENSMUSG00000022269.13 | MARCHF11 | 1.273986424 | 1.20E-08 | 1.68E-06 |
| ENSMUSG00000013974.3 | MCEMP1 | 2.526982008 | 6.82E-10 | 1.41E-07 |
| ENSMUSG00000026355.11 | MCM6 | 1.027075199 | 3.43E-04 | 0.008260644 |
| ENSMUSG00000032776.9 | MCTP2 | 1.249098273 | 2.63E-05 | 0.001089811 |
| ENSMUSG00000090667.2 | Mdfic2 | -3.043277425 | 3.79E-10 | 8.45E-08 |
| ENSMUSG00000029659.16 | MEDAG | 1.417771335 | 1.05E-06 | 8.00E-05 |
| ENSMUSG00000036466.17 | MEGF11 | 1.176231652 | 9.13E-06 | 4.78E-04 |
| ENSMUSG00000057751.14 | MEGF6 | 1.183446574 | 2.65E-08 | 3.45E-06 |
| ENSMUSG00000068117.10 | MEI1 | 1.045443084 | 6.26E-04 | 0.013270697 |
| ENSMUSG00000001493.9 | MEOX1 | 1.219445445 | 0.00162308 | 0.026884126 |
| ENSMUSG00000034739.17 | MFRP | -3.146750449 | 0.00174026 | 0.028303398 |
| ENSMUSG00000074813.14 | Morrbid | 2.093916523 | 1.59E-08 | 2.14E-06 |
| ENSMUSG00000047502.14 | MROH7 | 1.033976372 | 1.59E-10 | 3.75E-08 |
| ENSMUSG00000020814.13 | Mxra7 | -1.083592374 | 5.85E-13 | 2.84E-10 |
| ENSMUSG00000020061.18 | MYBPC1 | 2.669064113 | 3.23E-32 | 2.66E-28 |
| ENSMUSG00000002100.15 | MYBPC3 | 3.036786799 | 1.30E-28 | 3.57E-25 |
| ENSMUSG00000020908.14 | MYH3 | -1.296299299 | 1.32E-05 | 6.34E-04 |
| ENSMUSG00000061086.12 | MYL4 | 1.551216228 | 7.20E-17 | 7.93E-14 |

| | | | | |
|---|---|---|---|---|
| ENSMUSG00000022836.11 | MYLK | 1.208722845 | 9.84E-11 | 2.46E-08 |
| ENSMUSG00000042678.17 | MYO15A | 2.729796774 | 2.90E-14 | 1.99E-11 |
| ENSMUSG00000026697.10 | MYOC | -1.894127602 | 1.87E-04 | 0.005277665 |
| ENSMUSG00000031837.14 | NECAB2 | 1.02275611 | 1.98E-07 | 2.05E-05 |
| ENSMUSG00000004891.16 | Nes | 1.171214182 | 1.28E-05 | 6.16E-04 |
| ENSMUSG00000047180.8 | NEURL3 | 1.11449931 | 0.00130733 | 0.023065414 |
| ENSMUSG00000048540.8 | NHLH2 | 1.438648012 | 3.68E-06 | 2.28E-04 |
| ENSMUSG00000048528.7 | NKX1-2 | 6.886894923 | 7.34E-15 | 5.77E-12 |
| ENSMUSG00000038745.15 | NLRP6 | -1.785171245 | 4.17E-06 | 2.51E-04 |
| ENSMUSG00000025723.12 | NMB | -1.057721087 | 9.08E-04 | 0.017519192 |
| ENSMUSG00000020447.5 | NPC1L1 | -1.293965361 | 0.00176747 | 0.028531777 |
| ENSMUSG00000041616.9 | NPPA | 1.790369355 | 2.24E-04 | 0.005981937 |
| ENSMUSG00000059991.7 | NPTX2 | 3.400036307 | 1.05E-06 | 8.00E-05 |
| ENSMUSG00000071230.7 | NPW | 5.123964212 | 3.29E-11 | 9.69E-09 |
| ENSMUSG00000029819.6 | NPY | 1.136723051 | 4.33E-05 | 0.001622775 |
| ENSMUSG00000049134.15 | NRAP | 1.338151973 | 2.41E-05 | 0.00101848 |
| ENSMUSG00000040632.16 | NRL | 1.04383563 | 0.00117592 | 0.021328712 |
| ENSMUSG00000039481.5 | NRTN | 1.304303704 | 2.25E-11 | 6.75E-09 |
| ENSMUSG00000049107.13 | NTF3 | -2.146968688 | 1.21E-07 | 1.32E-05 |
| ENSMUSG00000074121.3 | NTF4 | 1.744758514 | 2.60E-05 | 0.001080857 |
| ENSMUSG00000061356.13 | NUGGC | 3.632550231 | 1.21E-13 | 6.89E-11 |
| ENSMUSG00000046719.7 | NXPH3 | -1.254057512 | 6.23E-08 | 7.35E-06 |
| ENSMUSG00000053765.4 | Oas1f | 7.272187295 | 9.70E-09 | 1.38E-06 |
| ENSMUSG00000061462.18 | OBSCN | 1.47291137 | 7.99E-05 | 0.002653188 |
| ENSMUSG00000022026.7 | OLFM4 | 1.419603331 | 7.21E-05 | 0.002443633 |
| ENSMUSG00000050511.1 | OPRD1 | 1.051020302 | 2.15E-05 | 9.38E-04 |
| ENSMUSG00000062372.13 | OTOF | -1.400403823 | 2.88E-04 | 0.007251213 |
| ENSMUSG00000050201.9 | OTOP2 | 1.614499549 | 5.29E-05 | 0.001918831 |
| ENSMUSG00000018862.11 | OTOP3 | 2.679440348 | 5.33E-05 | 0.001929307 |
| ENSMUSG00000044055.5 | OTOS | 1.97396556 | 0.00224802 | 0.034080108 |
| ENSMUSG00000021848.16 | OTX2 | -2.647304459 | 1.97E-04 | 0.005489392 |
| ENSMUSG00000020787.14 | P2RX1 | 1.5123234 | 7.80E-04 | 0.015467632 |
| ENSMUSG00000028370.7 | PAPPA | 3.200951901 | 9.30E-06 | 4.84E-04 |
| ENSMUSG00000022439.9 | PARVG | 1.940339349 | 1.48E-29 | 6.12E-26 |
| ENSMUSG00000035873.8 | PAWR | 1.218099847 | 6.71E-04 | 0.013934698 |
| ENSMUSG00000036422.10 | PCDH8 | 1.26824881 | 0.00168716 | 0.027723109 |
| ENSMUSG00000103442.5 | PCDHA1 | 1.875397692 | 0.00346585 | 0.046768996 |
| ENSMUSG00000103800.1 | PCDHA9 | 2.074702922 | 0.0025319 | 0.037232178 |
| ENSMUSG00000044254.6 | PCSK9 | 2.140754679 | 3.85E-05 | 0.001464328 |
| ENSMUSG00000055044.12 | PDLIM1 | 1.241935802 | 7.13E-05 | 0.002424363 |
| ENSMUSG00000031636.7 | Pdlim3 | 1.874588563 | 1.95E-05 | 8.63E-04 |
| ENSMUSG00000031212.3 | Pgr15l | 3.032408678 | 1.02E-05 | 5.15E-04 |
| ENSMUSG00000046207.14 | PIK3R6 | 1.456416199 | 1.45E-11 | 4.71E-09 |
| ENSMUSG00000029423.10 | PIWIL1 | 1.529466396 | 1.19E-06 | 8.82E-05 |
| ENSMUSG00000021822.3 | PLAU | 1.073904883 | 6.85E-08 | 7.86E-06 |

| | | | | |
|---|---|---|---|---|
| ENSMUSG00000029134.14 | PLB1 | 1.577313277 | 3.51E-05 | 0.001360155 |
| ENSMUSG00000031557.16 | PLEKHA2 | 1.151888991 | 2.01E-06 | 1.37E-04 |
| ENSMUSG00000032068.14 | PLET1 | 3.148886967 | 7.97E-06 | 4.27E-04 |
| ENSMUSG00000035486.14 | PLK5 | -2.20357001 | 1.48E-09 | 2.81E-07 |
| ENSMUSG00000031727.8 | PMFBP1 | 1.718069735 | 1.73E-08 | 2.30E-06 |
| ENSMUSG00000002228.7 | PPM1J | 3.31561664 | 1.13E-11 | 3.81E-09 |
| ENSMUSG00000040734.14 | PPP1R13L | 1.042973262 | 5.08E-04 | 0.011408328 |
| ENSMUSG00000052221.8 | PPP1R36 | 1.613767389 | 5.43E-08 | 6.50E-06 |
| ENSMUSG00000037086.3 | PRR32 | -3.63546952 | 0.00205379 | 0.031897115 |
| ENSMUSG00000045027.7 | PRSS22 | 1.108186699 | 6.13E-05 | 0.002153284 |
| ENSMUSG00000039405.7 | PRSS23 | 1.752177884 | 1.24E-15 | 1.20E-12 |
| ENSMUSG00000030623.4 | Prss23os | 1.751717302 | 2.26E-07 | 2.32E-05 |
| ENSMUSG00000024124.10 | Prss30 | 1.904210431 | 2.28E-05 | 9.84E-04 |
| ENSMUSG00000057729.12 | PRTN3 | 1.898738344 | 6.50E-14 | 4.13E-11 |
| ENSMUSG00000068744.12 | PSRC1 | -2.049279357 | 1.64E-11 | 5.20E-09 |
| ENSMUSG00000032322.14 | PSTPIP1 | 1.7997397 | 1.31E-16 | 1.35E-13 |
| ENSMUSG00000028378.6 | PTGR1 | 2.158267791 | 6.68E-13 | 3.15E-10 |
| ENSMUSG00000047250.13 | PTGS1 | 1.631544996 | 1.16E-08 | 1.64E-06 |
| ENSMUSG00000032487.8 | PTGS2 | 2.388237636 | 4.65E-04 | 0.010681859 |
| ENSMUSG00000025946.13 | PTH2R | -1.268330211 | 3.65E-04 | 0.008674183 |
| ENSMUSG00000097993.7 | Ptprv | -1.524561774 | 0.00285731 | 0.040433758 |
| ENSMUSG00000030559.8 | RAB38 | -1.852949762 | 0.00115925 | 0.021106784 |
| ENSMUSG00000004952.13 | RASA4 | 1.422783978 | 9.61E-14 | 5.88E-11 |
| ENSMUSG00000089809.9 | RASGEF1B | -1.103700842 | 4.05E-09 | 6.38E-07 |
| ENSMUSG00000025795.8 | RASSF3 | -1.211753579 | 1.06E-13 | 6.25E-11 |
| ENSMUSG00000041534.9 | RBP3 | 4.299563449 | 5.22E-09 | 7.98E-07 |
| ENSMUSG00000074269.10 | REC114 | 1.750724811 | 1.32E-06 | 9.70E-05 |
| ENSMUSG00000020275.9 | REL | 1.08546547 | 4.66E-04 | 0.010681859 |
| ENSMUSG00000022176.11 | REM2 | 1.313656697 | 0.00123295 | 0.022086063 |
| ENSMUSG00000030110.13 | RET | 1.790922997 | 2.89E-14 | 1.99E-11 |
| ENSMUSG00000012705.16 | RETN | 2.345743063 | 1.29E-05 | 6.19E-04 |
| ENSMUSG00000051079.8 | RGS13 | 3.089732434 | 2.69E-08 | 3.47E-06 |
| ENSMUSG00000026360.9 | RGS2 | 1.416466934 | 6.81E-05 | 0.002341481 |
| ENSMUSG00000044456.16 | RIN3 | 1.341959784 | 1.98E-09 | 3.63E-07 |
| ENSMUSG00000022221.14 | RIPK3 | 3.081075104 | 3.59E-15 | 3.12E-12 |
| ENSMUSG00000036492.12 | RNF39 | 1.163478883 | 5.69E-04 | 0.012428091 |
| ENSMUSG00000037593.12 | RSKR | -1.133496775 | 0.00113687 | 0.020860333 |
| ENSMUSG00000085925.3 | RTL1 | 3.345046919 | 4.50E-05 | 0.001670055 |
| ENSMUSG00000034009.14 | RXFP1 | -1.221769497 | 1.92E-05 | 8.55E-04 |
| ENSMUSG00000012017.4 | SCARF2 | 1.527397529 | 6.45E-12 | 2.37E-09 |
| ENSMUSG00000038936.13 | SCCPDH | 1.685113623 | 5.54E-29 | 1.83E-25 |
| ENSMUSG00000050711.7 | SCG2 | 1.43463126 | 1.33E-08 | 1.81E-06 |
| ENSMUSG00000046480.6 | SCN4B | -1.121191918 | 0.00163774 | 0.027072715 |
| ENSMUSG00000034810.7 | SCN7A | 2.40568777 | 5.95E-10 | 1.27E-07 |
| ENSMUSG00000038580.13 | Sct | 3.361004131 | 9.35E-10 | 1.88E-07 |

| | | | | |
|---|---|---|---|---|
| ENSMUSG00000040364.8 | Sec1 | -1.295511545 | 2.61E-04 | 0.006687219 |
| ENSMUSG00000028064.17 | SEMA4A | 1.080810419 | 4.04E-09 | 6.38E-07 |
| ENSMUSG00000063232.16 | SERPINA11 | 3.172157593 | 2.36E-05 | 0.001002441 |
| ENSMUSG00000079014.4 | Serpina3g (includes | 2.105770818 | 1.35E-05 | 6.43E-04 |
| ENSMUSG00000041449.16 | Serpina3h | 1.306317805 | 0.00339267 | 0.046036614 |
| ENSMUSG00000037411.10 | SERPINE1 | 2.426170807 | 0.00250961 | 0.037069543 |
| ENSMUSG00000056306.5 | SERTM1 | 1.092383806 | 3.11E-04 | 0.007720184 |
| ENSMUSG00000027996.13 | SFRP2 | 1.623657225 | 1.25E-05 | 6.06E-04 |
| ENSMUSG00000018822.7 | SFRP5 | 1.361111754 | 0.00144938 | 0.02482887 |
| ENSMUSG00000005057.13 | SH2B2 | 1.026576965 | 2.16E-04 | 0.005849452 |
| ENSMUSG00000034460.9 | SIX4 | -1.087300253 | 4.71E-05 | 0.001735206 |
| ENSMUSG00000022372.14 | SLA | -1.107993079 | 8.99E-10 | 1.83E-07 |
| ENSMUSG00000063652.10 | Slc22a21 | 1.298891471 | 2.14E-04 | 0.005821117 |
| ENSMUSG00000023828.3 | SLC22A3 | -1.215154391 | 9.68E-05 | 0.003138965 |
| ENSMUSG00000034224.14 | SLC38A8 | 3.186966594 | 2.63E-15 | 2.41E-12 |
| ENSMUSG00000039728.15 | SLC6A5 | 5.03170077 | 7.43E-07 | 5.93E-05 |
| ENSMUSG00000013611.14 | SNX31 | 1.249617858 | 0.00147699 | 0.025171265 |
| ENSMUSG00000063434.6 | SORCS3 | 1.007570226 | 4.93E-04 | 0.01113095 |
| ENSMUSG00000036169.6 | SOSTDC1 | -2.359708858 | 0.00305107 | 0.042506528 |
| ENSMUSG00000043461.13 | SPTSSB | -1.326374243 | 2.60E-07 | 2.53E-05 |
| ENSMUSG00000050824.12 | SSTR5 | 1.385055856 | 3.53E-05 | 0.001362573 |
| ENSMUSG00000014813.9 | STC1 | -1.499344247 | 4.66E-08 | 5.71E-06 |
| ENSMUSG00000020303.2 | STC2 | 2.836150053 | 2.46E-07 | 2.43E-05 |
| ENSMUSG00000029123.8 | STK32B | 1.634981046 | 1.85E-08 | 2.44E-06 |
| ENSMUSG00000027744.4 | STOML3 | -2.090125141 | 0.0012331 | 0.022086063 |
| ENSMUSG00000028860.13 | SYTL1 | 1.357584053 | 4.72E-06 | 2.80E-04 |
| ENSMUSG00000054453.11 | SYTL5 | 1.408850826 | 4.82E-07 | 4.21E-05 |
| ENSMUSG00000055865.8 | TAFA3 | 2.798298471 | 3.76E-10 | 8.45E-08 |
| ENSMUSG00000039813.14 | TBC1D2 | 1.030118361 | 3.73E-04 | 0.008826808 |
| ENSMUSG00000054003.13 | TDRD9 | 3.55682759 | 1.27E-12 | 5.84E-10 |
| ENSMUSG00000002603.15 | TGFB1 | 1.478930715 | 5.98E-10 | 1.27E-07 |
| ENSMUSG00000030782.17 | TGFB1I1 | 1.338929896 | 3.56E-07 | 3.22E-05 |
| ENSMUSG00000020317.14 | THEG | 1.287670854 | 0.0014271 | 0.024600312 |
| ENSMUSG00000037731.5 | THEMIS2 | 1.526395261 | 5.41E-08 | 6.50E-06 |
| ENSMUSG00000035686.8 | THRSP | 1.041792868 | 3.50E-07 | 3.22E-05 |
| ENSMUSG00000053626.5 | TLL1 | 1.740522122 | 4.79E-04 | 0.010867845 |
| ENSMUSG00000025013.15 | TLL2 | 1.596166734 | 0.0017057 | 0.027935789 |
| ENSMUSG00000027995.10 | TLR2 | 1.359368982 | 0.00272628 | 0.039251809 |
| ENSMUSG00000038540.14 | TMC3 | -1.40872136 | 0.0018427 | 0.029401288 |
| ENSMUSG00000056498.13 | TMEM154 | 4.758971163 | 8.34E-09 | 1.20E-06 |
| ENSMUSG00000059900.14 | TMEM40 | 2.193830363 | 5.56E-08 | 6.60E-06 |
| ENSMUSG00000061702.10 | TMEM91 | 1.6042517 | 1.43E-10 | 3.41E-08 |
| ENSMUSG00000024034.13 | TMPRSS3 | 2.488058026 | 3.92E-09 | 6.28E-07 |
| ENSMUSG00000016942.6 | TMPRSS6 | 2.618972761 | 3.66E-06 | 2.27E-04 |
| ENSMUSG00000028364.15 | TNC | 1.980635283 | 8.52E-06 | 4.51E-04 |

| | | | | |
|---|---|---|---|---|
| ENSMUSG00000024401.14 | TNF | 4.240961139 | 0.00179741 | 0.028860866 |
| ENSMUSG00000021281.15 | TNFAIP2 | 1.62190299 | 7.03E-04 | 0.014442916 |
| ENSMUSG00000091898.8 | TNNC1 | 1.792887783 | 1.06E-17 | 1.34E-14 |
| ENSMUSG00000035458.15 | TNNI3 | 1.760713159 | 1.36E-04 | 0.004101274 |
| ENSMUSG00000070867.4 | TRABD2B | -1.013984569 | 1.23E-07 | 1.34E-05 |
| ENSMUSG00000079259.2 | TRIM71 | 1.777173838 | 2.10E-04 | 0.005744871 |
| ENSMUSG00000056596.8 | TRNP1 | 1.315470014 | 1.01E-19 | 1.51E-16 |
| ENSMUSG00000061808.4 | TTR | -5.81611728 | 5.97E-06 | 3.38E-04 |
| ENSMUSG00000026668.10 | UCMA | 3.241583359 | 8.34E-12 | 2.87E-09 |
| ENSMUSG00000018845.14 | UNC45B | 1.067968362 | 1.07E-05 | 5.35E-04 |
| ENSMUSG00000022435.6 | UPK3A | 1.222574832 | 3.53E-05 | 0.001362573 |
| ENSMUSG00000045288.10 | USH1G | 2.179927011 | 2.68E-06 | 1.76E-04 |
| ENSMUSG00000022479.15 | VDR | 1.312191165 | 0.00331138 | 0.045193434 |
| ENSMUSG00000037428.14 | VGF | 1.389238835 | 6.86E-07 | 5.56E-05 |
| ENSMUSG00000000983.13 | Wfdc18 | 2.174068099 | 4.17E-09 | 6.50E-07 |
| ENSMUSG00000029671.15 | WNT16 | 1.560797465 | 1.03E-04 | 0.003294848 |
| ENSMUSG00000018486.2 | WNT9B | 4.142619228 | 3.29E-07 | 3.07E-05 |
| ENSMUSG00000033454.6 | ZBTB1 | 1.009912652 | 2.67E-04 | 0.006791936 |
| ENSMUSG00000039981.6 | ZC3H12D | 1.567604685 | 4.97E-05 | 0.001818379 |
| ENSMUSG00000096696.1 | Zfp960/Zfp97 | 1.68512116 | 0.00221589 | 0.033664421 |
| ENSMUSG00000011267.8 | ZNF296 | -1.171497468 | 5.63E-07 | 4.82E-05 |
| ENSMUSG00000043903.4 | ZNF469 | 1.325485874 | 3.36E-06 | 2.12E-04 |

### CTX

| ID | Symbol | Expr Log Ratio | Expr p-value | Expr FDR |
|---|---|---|---|---|
| ENSMUSG00000097178.7 | 2310002F09Rik | 1.051299945 | 0.001964933 | 0.031503795 |
| ENSMUSG00000085338.1 | 2410004I01Rik | 1.068217473 | 0.002181443 | 0.03388932 |
| ENSMUSG00000026736.2 | 4930426L09Rik | 1.022363151 | 3.27E-04 | 0.008538489 |
| ENSMUSG00000085129.1 | 5031425F14Rik | 3.063730259 | 5.19E-07 | 5.14E-05 |
| ENSMUSG00000054944.7 | 5330416C01Rik | 1.16667948 | 1.50E-05 | 7.90E-04 |
| ENSMUSG00000110140.1 | 5430421F17Rik | 1.833216158 | 3.95E-05 | 0.001694375 |
| ENSMUSG00000110148.1 | 5830408C22Rik | 1.594679936 | 1.94E-04 | 0.005728878 |
| ENSMUSG00000117916.1 | 9630028I04Rik | 1.841527493 | 1.42E-09 | 3.73E-07 |
| ENSMUSG00000117172.1 | A230051N06Rik | 1.006985398 | 8.95E-08 | 1.19E-05 |
| ENSMUSG00000097622.2 | A330033J07Rik | -1.195487634 | 0.002925952 | 0.041369159 |
| ENSMUSG00000113342.1 | AA414992 | 1.207673234 | 5.95E-04 | 0.013114868 |
| ENSMUSG00000118572.1 | AC122821.1 | -1.44427281 | 5.33E-04 | 0.01216218 |
| ENSMUSG00000114458.1 | AC129085.1 | 1.030650875 | 3.94E-05 | 0.001694375 |
| ENSMUSG00000010651.4 | Acaa1b | 1.939163986 | 1.51E-17 | 2.54E-14 |
| ENSMUSG00000030607.7 | Acan | 2.275361659 | 7.48E-04 | 0.01569994 |
| ENSMUSG00000011256.16 | ADAM19 | 1.11779385 | 3.93E-06 | 2.72E-04 |
| ENSMUSG00000053399.9 | ADAMTS18 | 1.400830392 | 3.79E-04 | 0.009559015 |
| ENSMUSG00000024256.7 | ADCYAP1 | 1.881921284 | 2.01E-13 | 1.66E-10 |
| ENSMUSG00000023918.12 | ADGRF4 | 2.318426342 | 3.63E-09 | 8.11E-07 |
| ENSMUSG00000030790.15 | ADM | 1.611980813 | 7.55E-04 | 0.015754844 |
| ENSMUSG00000050541.14 | ADRA1B | 1.10662834 | 3.26E-08 | 5.18E-06 |
| ENSMUSG00000058620.5 | ADRA2B | -1.320570158 | 5.97E-04 | 0.013135026 |
| ENSMUSG00000040706.4 | AGMAT | 2.25318956 | 2.28E-10 | 7.86E-08 |
| ENSMUSG00000031980.10 | AGT | -1.050812581 | 1.68E-05 | 8.60E-04 |
| ENSMUSG00000021575.16 | AHRR | 1.137234572 | 3.42E-05 | 0.001529178 |
| ENSMUSG00000084826.7 | AI847159 | 1.443001079 | 2.22E-04 | 0.006361402 |
| ENSMUSG00000029762.6 | AKR1B10 | 2.384757931 | 4.64E-05 | 0.001895388 |
| ENSMUSG00000021214.14 | AKR1C3 | 1.48187856 | 4.88E-06 | 3.20E-04 |
| ENSMUSG00000019102.10 | ALDH3A1 | 2.582813266 | 5.35E-19 | 1.10E-15 |
| ENSMUSG00000054204.6 | ALKAL2 | 1.910108132 | 0.00104649 | 0.020355398 |
| ENSMUSG00000032807.5 | ALOX12B | -1.53450024 | 1.03E-05 | 5.82E-04 |
| ENSMUSG00000025701.12 | ALOX5 | 2.009199063 | 7.13E-09 | 1.37E-06 |
| ENSMUSG00000068246.6 | Apol9a/Apol9b | 2.427720992 | 1.75E-05 | 8.93E-04 |
| ENSMUSG00000009248.6 | ASCL2 | 4.233901885 | 5.50E-06 | 3.48E-04 |
| ENSMUSG00000037686.6 | ASPG | 1.417719084 | 1.57E-05 | 8.25E-04 |
| ENSMUSG00000030730.12 | ATP2A1 | 3.567041 | 3.19E-15 | 4.05E-12 |
| ENSMUSG00000034112.9 | ATP2C2 | 2.034166884 | 7.74E-06 | 4.65E-04 |
| ENSMUSG00000109890.2 | AU023762 | 1.621717346 | 0.002591974 | 0.038115633 |
| ENSMUSG00000084843.1 | B230312C02Rik | 1.009551326 | 1.15E-04 | 0.003802102 |
| ENSMUSG00000079445.3 | B3GNT7 | 2.110332715 | 1.38E-08 | 2.44E-06 |
| ENSMUSG00000059479.7 | B3GNT8 | 1.774254976 | 6.00E-14 | 5.51E-11 |
| ENSMUSG00000046415.11 | B430212C06Rik | 1.177713656 | 0.001462131 | 0.025858361 |
| ENSMUSG00000047507.12 | BAIAP3 | -1.591272762 | 2.74E-05 | 0.001277479 |

| | | | | |
|---|---|---|---|---|
| ENSMUSG00000035021.13 | BAZ1A | 1.4556787 | 0.003683952 | 0.047907954 |
| ENSMUSG00000040363.15 | BCOR | 1.00294488 | 0.003296798 | 0.044696202 |
| ENSMUSG00000048482.14 | BDNF | 3.158882628 | 2.43E-09 | 5.82E-07 |
| ENSMUSG00000061132.13 | BLNK | 1.875463638 | 4.59E-10 | 1.40E-07 |
| ENSMUSG00000032726.12 | BMP8A | 1.434189761 | 0.001584168 | 0.027137922 |
| ENSMUSG00000027485.15 | BPIFB1 | 2.630688641 | 1.65E-09 | 4.22E-07 |
| ENSMUSG00000036907.9 | C1QL2 | 1.611151287 | 1.79E-10 | 6.73E-08 |
| ENSMUSG00000055172.10 | C1R | 1.11606978 | 1.55E-04 | 0.004793862 |
| ENSMUSG00000091956.2 | C2CD4B | 2.472537454 | 4.55E-05 | 0.001879213 |
| ENSMUSG00000078815.8 | CACNG6 | 1.111484689 | 3.77E-04 | 0.009545676 |
| ENSMUSG00000028348.7 | CAVIN4 | 1.671889003 | 5.11E-04 | 0.011809354 |
| ENSMUSG00000047810.9 | CCDC88B | 1.628359791 | 5.59E-16 | 8.40E-13 |
| ENSMUSG00000072082.7 | CCNF | 1.702572309 | 3.86E-06 | 2.70E-04 |
| ENSMUSG00000042417.5 | CCNO | -1.242554279 | 8.63E-05 | 0.003052541 |
| ENSMUSG00000028865.14 | CD164L2 | 1.356609559 | 2.24E-09 | 5.52E-07 |
| ENSMUSG00000028076.12 | CD1D | 1.933102141 | 3.01E-06 | 2.21E-04 |
| ENSMUSG00000022667.18 | CD200R1 | 1.333950466 | 4.36E-04 | 0.010568178 |
| ENSMUSG00000026012.2 | CD28 | 3.081938987 | 1.15E-12 | 8.22E-10 |
| ENSMUSG00000025163.6 | CD7 | 2.16227633 | 2.75E-08 | 4.53E-06 |
| ENSMUSG00000023067.14 | CDKN1A | 2.217659517 | 5.26E-05 | 0.0020882 |
| ENSMUSG00000037628.10 | CDKN3 | 2.084653643 | 4.09E-04 | 0.010070824 |
| ENSMUSG00000076433.4 | CEP295NL | 2.072214478 | 7.48E-05 | 0.002734241 |
| ENSMUSG00000021194.6 | CHGA | 1.381894605 | 9.09E-25 | 3.00E-21 |
| ENSMUSG00000042269.15 | CIBAR2 | 1.228434859 | 2.00E-04 | 0.00586842 |
| ENSMUSG00000037594.11 | CLBA1 | 1.031427365 | 1.90E-04 | 0.005616922 |
| ENSMUSG00000033633.15 | CLEC18B | 1.805488626 | 8.13E-08 | 1.10E-05 |
| ENSMUSG00000042489.15 | CLSPN | -1.293595224 | 6.01E-04 | 0.013172806 |
| ENSMUSG00000039462.4 | COL10A1 | 2.830726744 | 9.72E-05 | 0.003373658 |
| ENSMUSG00000045672.15 | COL27A1 | 1.51944678 | 6.66E-05 | 0.002485497 |
| ENSMUSG00000004098.7 | COL5A3 | 2.098400569 | 8.30E-07 | 7.66E-05 |
| ENSMUSG00000039070.5 | CPA4 | 1.603537328 | 1.18E-04 | 0.003906462 |
| ENSMUSG00000027230.9 | CREB3L1 | 1.138021741 | 6.59E-05 | 0.00247819 |
| ENSMUSG00000021680.8 | CRHBP | 1.477371311 | 1.25E-08 | 2.25E-06 |
| ENSMUSG00000007888.15 | CRLF1 | 1.174687264 | 3.76E-05 | 0.001642389 |
| ENSMUSG00000006546.3 | CRYBA2 | 3.226876545 | 4.78E-09 | 9.87E-07 |
| ENSMUSG00000012123.17 | CRYBG2 | 1.648476225 | 4.36E-08 | 6.61E-06 |
| ENSMUSG00000117220.1 | CT025671.1 | 1.171339856 | 4.76E-06 | 3.17E-04 |
| ENSMUSG00000044258.10 | Ctla2a/Ctla2b | 1.597695896 | 6.36E-05 | 0.002410266 |
| ENSMUSG00000026456.18 | CYB5R1 | 1.285295858 | 3.81E-14 | 3.93E-11 |
| ENSMUSG00000024087.4 | CYP1B1 | 1.492674574 | 3.09E-04 | 0.008133422 |
| ENSMUSG00000109198.1 | D7Bwg0826e | 2.167179468 | 1.86E-07 | 2.12E-05 |
| ENSMUSG00000035910.15 | DCDC2 | 1.066471011 | 4.99E-08 | 7.49E-06 |
| ENSMUSG00000007379.15 | DENND2C | 1.915436784 | 5.31E-05 | 0.002097436 |
| ENSMUSG00000027068.6 | DHRS9 | 2.940179253 | 2.15E-05 | 0.00106357 |
| ENSMUSG00000075707.5 | DIO3 | -2.113449325 | 6.88E-06 | 4.19E-04 |

| | | | | |
|---|---|---|---|---|
| ENSMUSG00000020871.8 | DLX4 | -1.261660617 | 0.001329675 | 0.024168686 |
| ENSMUSG00000086552.3 | Dlx4os | -1.394382024 | 0.001511438 | 0.026388975 |
| ENSMUSG00000060962.12 | DMKN | 1.18654339 | 8.19E-04 | 0.016892565 |
| ENSMUSG00000000730.13 | DNMT3L | 2.614799917 | 3.45E-11 | 1.58E-08 |
| ENSMUSG00000042737.9 | DPM3 | 1.447700599 | 1.93E-07 | 2.17E-05 |
| ENSMUSG00000025496.5 | DRD4 | 3.836923268 | 3.48E-08 | 5.42E-06 |
| ENSMUSG00000054889.10 | DSP | 1.334837065 | 2.10E-05 | 0.001044747 |
| ENSMUSG00000027368.6 | DUSP2 | 1.3226873 | 0.001241225 | 0.022953627 |
| ENSMUSG00000031530.6 | DUSP4 | 1.343673601 | 0.002270224 | 0.034585314 |
| ENSMUSG00000034765.6 | DUSP5 | 1.246001269 | 0.003128664 | 0.04318749 |
| ENSMUSG00000022441.17 | EFCAB6 | 3.415496634 | 5.92E-12 | 3.15E-09 |
| ENSMUSG00000045394.9 | EPCAM | 1.232278018 | 0.002305543 | 0.034928609 |
| ENSMUSG00000043439.5 | EPOP | -1.077727065 | 1.02E-05 | 5.82E-04 |
| ENSMUSG00000021255.17 | ESRRB | 1.163178316 | 5.59E-05 | 0.002187255 |
| ENSMUSG00000003382.18 | ETV3 | 1.264264596 | 9.82E-05 | 0.003392407 |
| ENSMUSG00000118505.1 | ETV3L | 5.888703162 | 1.21E-05 | 6.53E-04 |
| ENSMUSG00000021675.4 | F2RL2 | 1.94153999 | 4.09E-07 | 4.20E-05 |
| ENSMUSG00000047115.15 | FAM221A | 1.216216785 | 6.88E-04 | 0.014618857 |
| ENSMUSG00000042377.8 | FAM83G | 1.738957132 | 0.001519859 | 0.02647568 |
| ENSMUSG00000089665.2 | Fcor | 1.010491722 | 3.93E-05 | 0.00169261 |
| ENSMUSG00000070504.9 | FCRL6 | 4.568840506 | 1.12E-07 | 1.43E-05 |
| ENSMUSG00000021732.14 | FGF10 | -1.307465728 | 1.58E-06 | 1.27E-04 |
| ENSMUSG00000028874.14 | FGR | 4.057005922 | 1.48E-19 | 3.49E-16 |
| ENSMUSG00000051435.11 | FHAD1 | 1.16830429 | 5.21E-08 | 7.62E-06 |
| ENSMUSG00000045326.13 | FNDC7 | 2.057478476 | 6.44E-06 | 3.98E-04 |
| ENSMUSG00000048721.2 | FNDC9 | 3.055741024 | 5.99E-05 | 0.002298766 |
| ENSMUSG00000024912.6 | FOSL1 | 3.744342099 | 7.63E-04 | 0.015879783 |
| ENSMUSG00000027004.3 | FRZB | -1.385250106 | 4.98E-09 | 1.02E-06 |
| ENSMUSG00000056880.12 | GADL1 | 1.42198129 | 5.11E-05 | 0.002046816 |
| ENSMUSG00000024907.7 | GAL | 3.168694851 | 2.39E-05 | 0.00113599 |
| ENSMUSG00000021903.11 | GALNT15 | 1.147567047 | 0.001847965 | 0.030125654 |
| ENSMUSG00000051136.1 | GHSR | 2.118976123 | 0.003425132 | 0.045773447 |
| ENSMUSG00000048582.7 | GJA3 | 3.406760762 | 2.25E-07 | 2.47E-05 |
| ENSMUSG00000047197.1 | GJD3 | 1.068322097 | 7.90E-05 | 0.002854001 |
| ENSMUSG00000056888.11 | GLIPR1 | 1.307919883 | 0.001234229 | 0.022849832 |
| ENSMUSG00000000263.15 | GLRA1 | 2.636926947 | 5.00E-07 | 4.98E-05 |
| ENSMUSG00000074776.4 | Gm10754 | -1.467923703 | 6.92E-04 | 0.014668754 |
| ENSMUSG00000084375.1 | Gm11637 | 1.074090495 | 1.25E-04 | 0.004085373 |
| ENSMUSG00000075437.4 | Gm11681 | -1.309422131 | 5.89E-04 | 0.013060663 |
| ENSMUSG00000087496.1 | Gm13523 | 1.303991677 | 5.07E-06 | 3.30E-04 |
| ENSMUSG00000085397.1 | Gm13524 | 1.321272049 | 2.85E-04 | 0.007642837 |
| ENSMUSG00000086196.7 | Gm13571 | 3.179885019 | 2.68E-04 | 0.007359381 |
| ENSMUSG00000091556.8 | Gm14569 | -1.607390328 | 0.002072831 | 0.032756685 |
| ENSMUSG00000085225.1 | Gm14637 | 3.613128036 | 8.90E-06 | 5.21E-04 |
| ENSMUSG00000087556.1 | Gm15764 | 2.036537965 | 5.03E-05 | 0.002024817 |

| | | | | |
|---|---|---|---|---|
| ENSMUSG00000087411.1 | Gm16048 | 2.409548302 | 2.65E-09 | 6.17E-07 |
| ENSMUSG00000083218.2 | Gm16425 | 1.11837356 | 0.001251742 | 0.023096392 |
| ENSMUSG00000102548.1 | Gm16701 | 1.08026326 | 2.78E-04 | 0.007547858 |
| ENSMUSG00000086289.3 | Gm16933 | 1.39002646 | 0.00129586 | 0.02373759 |
| ENSMUSG00000117767.1 | Gm18786 | 1.533626069 | 5.48E-06 | 3.48E-04 |
| ENSMUSG00000092974.2 | Gm25072 | 1.211390643 | 0.002521828 | 0.037259594 |
| ENSMUSG00000097684.1 | Gm26645 | -1.309509298 | 3.28E-05 | 0.001474651 |
| ENSMUSG00000098088.1 | Gm26916 | 1.083616039 | 0.001554862 | 0.026865014 |
| ENSMUSG00000099644.1 | Gm28626 | 3.113926041 | 3.91E-09 | 8.50E-07 |
| ENSMUSG00000100980.1 | Gm29100 | 2.020029561 | 0.001098463 | 0.021093043 |
| ENSMUSG00000100546.1 | Gm29483 | 2.194826981 | 7.08E-13 | 5.32E-10 |
| ENSMUSG00000100417.2 | Gm29595 | 1.066849165 | 2.57E-04 | 0.007166603 |
| ENSMUSG00000113689.1 | Gm29676 | 2.592782421 | 2.20E-04 | 0.006339111 |
| ENSMUSG00000112035.1 | Gm30539 | 1.055929457 | 0.003539428 | 0.04664814 |
| ENSMUSG00000114271.1 | Gm32224 | 1.146183609 | 0.003750322 | 0.048460728 |
| ENSMUSG00000117254.1 | Gm34567 | 1.490245495 | 2.75E-05 | 0.001277479 |
| ENSMUSG00000107272.1 | Gm34583 | 1.321015898 | 1.32E-04 | 0.004258184 |
| ENSMUSG00000103785.1 | Gm35025 | 1.413618429 | 6.70E-05 | 0.002492684 |
| ENSMUSG00000116114.1 | Gm35853 | 3.12009593 | 1.04E-09 | 2.76E-07 |
| ENSMUSG00000102240.1 | Gm37717 | 1.985675365 | 8.62E-04 | 0.017609877 |
| ENSMUSG00000103622.1 | Gm38391 | 1.29823838 | 7.15E-08 | 9.92E-06 |
| ENSMUSG00000110179.1 | Gm39168 | 1.934392691 | 5.26E-06 | 3.39E-04 |
| ENSMUSG00000110618.1 | Gm39822 | 1.187121183 | 1.11E-06 | 9.65E-05 |
| ENSMUSG00000116946.1 | Gm41442 | 2.054048426 | 2.34E-04 | 0.00656226 |
| ENSMUSG00000105152.1 | Gm42864 | -2.040596471 | 2.35E-10 | 7.87E-08 |
| ENSMUSG00000104807.1 | Gm42865 | -1.799195617 | 3.82E-05 | 0.001658696 |
| ENSMUSG00000104850.1 | Gm42901 | 1.256898963 | 2.24E-04 | 0.006395527 |
| ENSMUSG00000086804.8 | Gm43154 | 1.657558579 | 5.69E-08 | 8.16E-06 |
| ENSMUSG00000109130.1 | Gm45187 | -1.443055622 | 7.23E-05 | 0.002671022 |
| ENSMUSG00000109604.1 | Gm45346 | 1.876963802 | 4.73E-05 | 0.001923745 |
| ENSMUSG00000109957.1 | Gm45353 | 1.028307198 | 0.00152593 | 0.026500649 |
| ENSMUSG00000110086.1 | Gm45623 | -1.060659903 | 9.81E-06 | 5.62E-04 |
| ENSMUSG00000081436.1 | Gm5912 | 1.730097732 | 2.30E-05 | 0.001108465 |
| ENSMUSG00000097806.3 | Gm6556 | -1.398336959 | 3.59E-07 | 3.77E-05 |
| ENSMUSG00000073532.3 | Gm7276 | 1.239520983 | 1.48E-04 | 0.004682664 |
| ENSMUSG00000029816.10 | GPNMB | 1.344420055 | 4.43E-04 | 0.010672347 |
| ENSMUSG00000066197.5 | GPR139 | 1.182281521 | 3.94E-04 | 0.009808303 |
| ENSMUSG00000043441.5 | GPR149 | -1.15136238 | 0.001894689 | 0.030765882 |
| ENSMUSG00000045509.7 | GPR150 | -1.219397074 | 4.50E-05 | 0.00186345 |
| ENSMUSG00000042816.4 | GPR151 | 1.284958645 | 2.68E-04 | 0.007359381 |
| ENSMUSG00000049649.8 | GPR3 | 1.666935195 | 5.04E-04 | 0.011701837 |
| ENSMUSG00000051043.16 | GPRC5C | 1.755571837 | 1.80E-06 | 1.40E-04 |
| ENSMUSG00000050069.3 | GREM2 | 1.003935154 | 3.23E-08 | 5.18E-06 |
| ENSMUSG00000042638.14 | GUCY2C | 1.250968382 | 0.001724846 | 0.028822086 |
| ENSMUSG00000030074.9 | GXYLT2 | 1.351883626 | 1.66E-05 | 8.55E-04 |

| | | | | |
|---|---|---|---|---|
| ENSMUSG00000032338.9 | HCN4 | 1.095889125 | 6.49E-05 | 0.00245085 |
| ENSMUSG00000028778.15 | HCRTR1 | 1.094650752 | 0.001383838 | 0.024759157 |
| ENSMUSG00000047171.8 | HELT | 2.155880667 | 1.51E-04 | 0.004728891 |
| ENSMUSG00000021260.8 | HHIPL1 | 1.186272891 | 2.08E-10 | 7.62E-08 |
| ENSMUSG00000038403.10 | HJV | -1.231032986 | 0.00278336 | 0.040003834 |
| ENSMUSG00000055632.18 | HMCN2 | 3.117075909 | 7.12E-09 | 1.37E-06 |
| ENSMUSG00000031722.10 | HP | 2.488465313 | 3.73E-06 | 2.63E-04 |
| ENSMUSG00000043155.4 | HPDL | 1.225803092 | 6.04E-04 | 0.013204149 |
| ENSMUSG00000051456.4 | HSPB3 | 2.041101053 | 5.91E-08 | 8.34E-06 |
| ENSMUSG00000037406.7 | HTRA4 | 3.177896961 | 2.88E-11 | 1.36E-08 |
| ENSMUSG00000076614.7 | IGHG1 | 3.752133592 | 2.65E-07 | 2.88E-05 |
| ENSMUSG00000015966.17 | IL17RB | 3.390117588 | 2.71E-09 | 6.22E-07 |
| ENSMUSG00000026073.13 | IL1R2 | 1.633189331 | 3.76E-04 | 0.009535671 |
| ENSMUSG00000041324.14 | INHBA | 3.113125601 | 5.35E-06 | 3.42E-04 |
| ENSMUSG00000032243.8 | ITGA11 | 1.340920986 | 2.50E-04 | 0.006974573 |
| ENSMUSG00000015533.9 | ITGA2 | 1.369303554 | 0.001760428 | 0.029247195 |
| ENSMUSG00000020216.13 | JSRP1 | 2.017436756 | 1.75E-06 | 1.38E-04 |
| ENSMUSG00000038292.14 | KASH5 | 1.083747413 | 1.29E-05 | 6.91E-04 |
| ENSMUSG00000059852.7 | KCNG2 | -1.123854134 | 9.99E-08 | 1.31E-05 |
| ENSMUSG00000045246.11 | KCNG4 | 1.089503731 | 2.11E-06 | 1.61E-04 |
| ENSMUSG00000009545.14 | KCNQ1 | 1.711792102 | 2.96E-05 | 0.001359805 |
| ENSMUSG00000027115.14 | KIF18A | 1.240672293 | 0.002044618 | 0.032430353 |
| ENSMUSG00000074001.3 | KLHL40 | 1.87902514 | 5.54E-05 | 0.002173722 |
| ENSMUSG00000108444.1 | Klk2-ps | 3.644145681 | 1.70E-10 | 6.54E-08 |
| ENSMUSG00000030713.6 | KLK7 | 2.348973336 | 3.75E-04 | 0.009535671 |
| ENSMUSG00000064023.4 | KLK8 | 1.118752937 | 3.55E-05 | 0.001577405 |
| ENSMUSG00000023039.17 | KRT7 | 2.911465759 | 2.18E-08 | 3.71E-06 |
| ENSMUSG00000022986.5 | KRT75 | 6.093574675 | 9.91E-09 | 1.82E-06 |
| ENSMUSG00000066652.5 | LEFTY2 | -1.324313776 | 0.002108982 | 0.033137704 |
| ENSMUSG00000042793.13 | LGR6 | 1.354935383 | 2.92E-05 | 0.001349993 |
| ENSMUSG00000041700.9 | LHFPL1 | 3.258615582 | 3.44E-08 | 5.40E-06 |
| ENSMUSG00000026934.15 | LHX3 | 3.14715173 | 1.66E-09 | 4.22E-07 |
| ENSMUSG00000019230.14 | LHX9 | -1.278486413 | 1.07E-04 | 0.003606531 |
| ENSMUSG00000053846.5 | LIPG | 1.484552309 | 9.38E-04 | 0.018831554 |
| ENSMUSG00000020782.18 | LLGL2 | 1.318981758 | 1.22E-06 | 1.04E-04 |
| ENSMUSG00000090291.3 | LRRC10B | -1.117913889 | 0.003474452 | 0.046044852 |
| ENSMUSG00000022759.15 | Lrrc74b | 1.298969814 | 2.18E-05 | 0.001073883 |
| ENSMUSG00000044847.13 | LSM11 | 1.165916203 | 3.90E-06 | 2.71E-04 |
| ENSMUSG00000018819.10 | LSP1 | 1.175493041 | 1.94E-07 | 2.17E-05 |
| ENSMUSG00000034634.7 | LY6D | 1.719815339 | 2.71E-04 | 0.007409418 |
| ENSMUSG00000013766.11 | Ly6g6e | 2.83599219 | 5.49E-08 | 7.96E-06 |
| ENSMUSG00000026344.9 | LYPD1 | -1.022813654 | 4.52E-06 | 3.05E-04 |
| ENSMUSG00000032530.14 | LYZL4 | 2.617470054 | 3.02E-05 | 0.001379519 |
| ENSMUSG00000044313.13 | MAB21L3 | 1.988248622 | 0.002790237 | 0.040032997 |
| ENSMUSG00000047591.5 | MAFA | 1.042614317 | 2.87E-04 | 0.007646705 |

| | | | | |
|---|---|---|---|---|
| ENSMUSG00000022269.13 | MARCHF11 | 1.083311377 | 1.08E-06 | 9.55E-05 |
| ENSMUSG00000013974.3 | MCEMP1 | 2.160156255 | 2.55E-08 | 4.25E-06 |
| ENSMUSG00000029659.16 | MEDAG | 1.556984325 | 1.03E-07 | 1.33E-05 |
| ENSMUSG00000068117.10 | MEI1 | 1.022503704 | 8.31E-04 | 0.017081984 |
| ENSMUSG00000039208.15 | METRNL | 1.1246156 | 1.13E-04 | 0.003764839 |
| ENSMUSG00000101860.4 | Mindy4b-ps | -1.025076125 | 5.73E-05 | 0.002233453 |
| ENSMUSG00000075020.6 | Mir670hg | 1.864363639 | 2.84E-04 | 0.007642837 |
| ENSMUSG00000048416.15 | MLF1 | 1.680120786 | 0.003489879 | 0.046179377 |
| ENSMUSG00000074813.14 | Morrbid | 1.556882109 | 9.70E-06 | 5.58E-04 |
| ENSMUSG00000050808.13 | MUC15 | 1.308551573 | 1.02E-04 | 0.003480701 |
| ENSMUSG00000042485.7 | MUSTN1 | 1.125487613 | 0.001334737 | 0.024168686 |
| ENSMUSG00000020061.18 | MYBPC1 | 3.112732332 | 5.15E-42 | 8.51E-38 |
| ENSMUSG00000002100.15 | MYBPC3 | 3.051707292 | 7.17E-28 | 3.95E-24 |
| ENSMUSG00000020908.14 | MYH3 | -1.125299779 | 1.59E-04 | 0.004893883 |
| ENSMUSG00000061086.12 | MYL4 | 1.395281159 | 6.48E-14 | 5.64E-11 |
| ENSMUSG00000031698.14 | MYLK3 | 1.271721124 | 0.001900436 | 0.030828876 |
| ENSMUSG00000042678.17 | MYO15A | 2.163624979 | 7.10E-10 | 2.02E-07 |
| ENSMUSG00000026697.10 | MYOC | -2.113172609 | 2.72E-05 | 0.001270867 |
| ENSMUSG00000021032.13 | NGB | -1.41899456 | 5.91E-05 | 0.002274467 |
| ENSMUSG00000048540.8 | NHLH2 | 1.091163669 | 1.73E-04 | 0.005242729 |
| ENSMUSG00000048528.7 | NKX1-2 | 7.153511036 | 1.79E-15 | 2.47E-12 |
| ENSMUSG00000038745.15 | NLRP6 | -1.302196938 | 3.95E-04 | 0.009813918 |
| ENSMUSG00000041616.9 | NPPA | 2.81839262 | 1.46E-07 | 1.71E-05 |
| ENSMUSG00000059991.7 | NPTX2 | 3.209926529 | 3.35E-06 | 2.44E-04 |
| ENSMUSG00000071230.7 | NPW | 4.965198403 | 8.57E-11 | 3.72E-08 |
| ENSMUSG00000029819.6 | NPY | 1.579629327 | 1.90E-08 | 3.30E-06 |
| ENSMUSG00000049134.15 | NRAP | 1.847692981 | 2.75E-09 | 6.22E-07 |
| ENSMUSG00000039114.16 | NRN1 | 1.038317973 | 2.23E-04 | 0.006374396 |
| ENSMUSG00000049107.13 | NTF3 | -1.186086542 | 0.001496435 | 0.026254676 |
| ENSMUSG00000074121.3 | NTF4 | 2.101233294 | 1.58E-06 | 1.27E-04 |
| ENSMUSG00000061356.13 | NUGGC | 3.348555545 | 4.28E-12 | 2.62E-09 |
| ENSMUSG00000053765.4 | Oas1f | 6.016315485 | 7.78E-07 | 7.30E-05 |
| ENSMUSG00000034990.15 | OTOA | 1.358936367 | 8.92E-04 | 0.018059259 |
| ENSMUSG00000062372.13 | OTOF | -1.566281026 | 5.25E-05 | 0.0020882 |
| ENSMUSG00000091455.4 | OTOGL | 1.298550735 | 0.001136241 | 0.021542923 |
| ENSMUSG00000050201.9 | OTOP2 | 2.222092035 | 7.29E-08 | 1.00E-05 |
| ENSMUSG00000018862.11 | OTOP3 | 2.747356186 | 3.23E-05 | 0.001463005 |
| ENSMUSG00000028370.7 | PAPPA | 3.503023362 | 1.39E-06 | 1.16E-04 |
| ENSMUSG00000022439.9 | PARVG | 1.566466769 | 3.32E-20 | 9.13E-17 |
| ENSMUSG00000112129.1 | PBLD | 1.016878428 | 6.06E-05 | 0.002315642 |
| ENSMUSG00000036422.10 | PCDH8 | 1.528932924 | 1.69E-04 | 0.005146078 |
| ENSMUSG00000055044.12 | PDLIM1 | 1.325365085 | 2.19E-05 | 0.001078476 |
| ENSMUSG00000046207.14 | PIK3R6 | 1.368868761 | 2.38E-10 | 7.87E-08 |
| ENSMUSG00000029423.10 | PIWIL1 | 2.359760527 | 3.85E-10 | 1.20E-07 |
| ENSMUSG00000021822.3 | PLAU | 1.153903417 | 6.24E-09 | 1.23E-06 |

| | | | | |
|---|---|---|---|---|
| ENSMUSG00000029134.14 | PLB1 | 1.777925304 | 3.52E-06 | 2.54E-04 |
| ENSMUSG00000031557.16 | PLEKHA2 | 1.214649911 | 5.66E-07 | 5.49E-05 |
| ENSMUSG00000032068.14 | PLET1 | 3.25068908 | 2.84E-06 | 2.11E-04 |
| ENSMUSG00000035486.14 | PLK5 | -2.524515255 | 5.90E-12 | 3.15E-09 |
| ENSMUSG00000002228.7 | PPM1J | 3.455498164 | 2.58E-12 | 1.65E-09 |
| ENSMUSG00000040734.14 | PPP1R13L | 1.038562934 | 5.38E-04 | 0.012263929 |
| ENSMUSG00000075410.13 | Prcd | -1.180206214 | 4.78E-07 | 4.79E-05 |
| ENSMUSG00000045027.7 | PRSS22 | 1.290502465 | 6.72E-06 | 4.11E-04 |
| ENSMUSG00000039405.7 | PRSS23 | 1.879078308 | 1.54E-17 | 2.54E-14 |
| ENSMUSG00000030623.4 | Prss23os | 1.697983629 | 7.07E-06 | 4.29E-04 |
| ENSMUSG00000057729.12 | PRTN3 | 1.132151896 | 4.89E-06 | 3.20E-04 |
| ENSMUSG00000068744.12 | PSRC1 | -1.06169951 | 3.03E-04 | 0.008013562 |
| ENSMUSG00000032322.14 | PSTPIP1 | 1.374851208 | 1.69E-10 | 6.54E-08 |
| ENSMUSG00000028378.6 | PTGR1 | 2.222534333 | 5.67E-14 | 5.51E-11 |
| ENSMUSG00000047250.13 | PTGS1 | 1.508913116 | 1.21E-07 | 1.49E-05 |
| ENSMUSG00000032487.8 | PTGS2 | 2.998539294 | 1.80E-05 | 9.10E-04 |
| ENSMUSG00000037992.16 | RARA | 1.011452362 | 0.001288668 | 0.023671927 |
| ENSMUSG00000004952.13 | RASA4 | 1.311534622 | 5.73E-12 | 3.15E-09 |
| ENSMUSG00000041534.9 | RBP3 | 2.573392316 | 3.35E-05 | 0.001501699 |
| ENSMUSG00000074269.10 | REC114 | 2.106008712 | 4.29E-09 | 9.07E-07 |
| ENSMUSG00000040121.5 | REP15 | 1.029204051 | 7.42E-05 | 0.00271792 |
| ENSMUSG00000030110.13 | RET | 1.294955897 | 2.00E-08 | 3.44E-06 |
| ENSMUSG00000012705.16 | RETN | 2.34382089 | 5.71E-06 | 3.60E-04 |
| ENSMUSG00000051079.8 | RGS13 | 2.769294264 | 4.49E-07 | 4.55E-05 |
| ENSMUSG00000026360.9 | RGS2 | 1.435197743 | 5.46E-05 | 0.002144908 |
| ENSMUSG00000044456.16 | RIN3 | 1.04682545 | 2.00E-06 | 1.54E-04 |
| ENSMUSG00000022221.14 | RIPK3 | 2.755152214 | 3.01E-13 | 2.36E-10 |
| ENSMUSG00000036492.12 | RNF39 | 1.329947709 | 8.86E-05 | 0.003119888 |
| ENSMUSG00000112505.1 | RP23-118L13.6 | 1.105281909 | 1.83E-04 | 0.005441389 |
| ENSMUSG00000112255.1 | RP24-323H7.5 | 1.0978256 | 1.05E-04 | 0.003546202 |
| ENSMUSG00000111844.1 | RP24-94A19.4 | 1.62506226 | 0.003038635 | 0.042348772 |
| ENSMUSG00000028174.12 | RPE65 | -1.525409009 | 0.003205721 | 0.043860218 |
| ENSMUSG00000038936.13 | SCCPDH | 1.585151017 | 6.13E-26 | 2.53E-22 |
| ENSMUSG00000050195.8 | Scd4 | 1.812335407 | 0.003430176 | 0.045773447 |
| ENSMUSG00000050711.7 | SCG2 | 1.360098927 | 6.74E-08 | 9.43E-06 |
| ENSMUSG00000034810.7 | SCN7A | 1.785837762 | 2.81E-06 | 2.10E-04 |
| ENSMUSG00000038580.13 | Sct | 3.57126012 | 5.26E-12 | 3.10E-09 |
| ENSMUSG00000028064.17 | SEMA4A | 1.185496438 | 1.21E-10 | 4.87E-08 |
| ENSMUSG00000023232.17 | SERINC2 | 1.061898586 | 7.99E-04 | 0.016557857 |
| ENSMUSG00000063232.16 | SERPINA11 | 3.730675007 | 1.19E-06 | 1.02E-04 |
| ENSMUSG00000079012.11 | SERPINA3 | 1.279852903 | 4.84E-04 | 0.011412588 |
| ENSMUSG00000079014.4 | Serpina3g (include | 2.109601248 | 9.29E-06 | 5.40E-04 |
| ENSMUSG00000041449.16 | Serpina3h | 1.912621126 | 2.46E-05 | 0.001159316 |
| ENSMUSG00000038224.12 | SERPINF2 | 2.098307785 | 0.002209694 | 0.034133997 |
| ENSMUSG00000008384.8 | SERTAD1 | 1.206806461 | 0.001809906 | 0.029710531 |

| | | | | |
|---|---|---|---|---|
| ENSMUSG00000027996.13 | SFRP2 | 1.603942867 | 1.60E-05 | 8.29E-04 |
| ENSMUSG00000050010.8 | SHISA3 | -1.595528809 | 0.002692523 | 0.039106706 |
| ENSMUSG00000034224.14 | SLC38A8 | 1.839688107 | 4.60E-07 | 4.63E-05 |
| ENSMUSG00000039728.15 | SLC6A5 | 3.943443062 | 4.32E-05 | 0.001803205 |
| ENSMUSG00000013611.14 | SNX31 | 1.99221403 | 8.31E-07 | 7.66E-05 |
| ENSMUSG00000072663.12 | SPEF2 | -1.05418471 | 1.86E-04 | 0.005538209 |
| ENSMUSG00000020303.2 | STC2 | 2.987590023 | 5.73E-08 | 8.16E-06 |
| ENSMUSG00000028860.13 | SYTL1 | 1.247713083 | 2.28E-05 | 0.001100163 |
| ENSMUSG00000055865.8 | TAFA3 | 3.133938491 | 2.21E-10 | 7.75E-08 |
| ENSMUSG00000026547.15 | TAGLN2 | 1.044675304 | 0.001062724 | 0.020574238 |
| ENSMUSG00000039813.14 | TBC1D2 | 1.02832461 | 3.84E-04 | 0.009626956 |
| ENSMUSG00000007877.2 | TCAP | -1.181554485 | 6.63E-06 | 4.08E-04 |
| ENSMUSG00000068079.5 | TCF15 | 1.000434747 | 1.13E-04 | 0.003764839 |
| ENSMUSG00000054003.13 | TDRD9 | 3.335211844 | 1.96E-11 | 9.52E-09 |
| ENSMUSG00000002603.15 | TGFB1 | 1.263190855 | 1.01E-07 | 1.32E-05 |
| ENSMUSG00000030782.17 | TGFB1I1 | 1.155140984 | 1.03E-05 | 5.83E-04 |
| ENSMUSG00000022218.16 | TGM1 | 3.439344924 | 0.00203479 | 0.03231012 |
| ENSMUSG00000020317.14 | THEG | 1.789927442 | 2.31E-05 | 0.001109276 |
| ENSMUSG00000037731.5 | THEMIS2 | 1.148216436 | 4.05E-05 | 0.0017199 |
| ENSMUSG00000035686.8 | THRSP | 1.385313115 | 1.67E-11 | 8.53E-09 |
| ENSMUSG00000053626.5 | TLL1 | 2.24998707 | 8.26E-06 | 4.87E-04 |
| ENSMUSG00000056498.13 | TMEM154 | 3.786947898 | 1.37E-06 | 1.15E-04 |
| ENSMUSG00000045036.15 | TMEM232 | 1.210360811 | 3.64E-06 | 2.59E-04 |
| ENSMUSG00000059900.14 | TMEM40 | 3.236667681 | 1.89E-14 | 2.23E-11 |
| ENSMUSG00000061702.10 | TMEM91 | 1.211349541 | 9.65E-07 | 8.66E-05 |
| ENSMUSG00000024034.13 | TMPRSS3 | 1.579596156 | 2.97E-05 | 0.001361682 |
| ENSMUSG00000028965.13 | TNFRSF9 | 1.216913716 | 5.28E-04 | 0.012101608 |
| ENSMUSG00000091898.8 | TNNC1 | 2.507990732 | 1.40E-30 | 1.15E-26 |
| ENSMUSG00000035458.15 | TNNI3 | 1.565978962 | 3.91E-04 | 0.009756471 |
| ENSMUSG00000056596.8 | TRNP1 | 1.095281198 | 2.86E-14 | 3.15E-11 |
| ENSMUSG00000030351.5 | TSPAN11 | -1.085181246 | 5.83E-06 | 3.66E-04 |
| ENSMUSG00000001473.7 | TUBB6 | 2.528784619 | 1.54E-04 | 0.004786477 |
| ENSMUSG00000026668.10 | UCMA | 3.34718876 | 2.24E-12 | 1.54E-09 |
| ENSMUSG00000062151.13 | UNC13C | -1.054109827 | 2.12E-10 | 7.62E-08 |
| ENSMUSG00000029546.12 | UNCX | -1.232643178 | 0.003171464 | 0.043521138 |
| ENSMUSG00000049436.5 | UPK1B | -1.343572554 | 0.001462499 | 0.025858361 |
| ENSMUSG00000022435.6 | UPK3A | 1.275120406 | 1.29E-05 | 6.91E-04 |
| ENSMUSG00000045288.10 | USH1G | 2.499532155 | 1.30E-07 | 1.56E-05 |
| ENSMUSG00000022479.15 | VDR | 1.36394984 | 0.00204629 | 0.032430353 |
| ENSMUSG00000037428.14 | VGF | 1.334318879 | 1.79E-06 | 1.40E-04 |
| ENSMUSG00000032528.5 | VIPR1 | 1.005491779 | 1.26E-05 | 6.77E-04 |
| ENSMUSG00000028753.12 | VWA5B1 | -1.152524891 | 5.61E-05 | 0.002191152 |
| ENSMUSG00000000983.13 | Wfdc18 | 2.513129151 | 1.70E-11 | 8.53E-09 |
| ENSMUSG00000026167.14 | WNT10A | 1.248394888 | 1.87E-05 | 9.41E-04 |
| ENSMUSG00000029671.15 | WNT16 | 1.348441131 | 6.58E-04 | 0.014118382 |

| | | | | |
|---|---|---|---|---|
| ENSMUSG00000027840.5 | WNT2B | -2.054802032 | 1.16E-07 | 1.45E-05 |
| ENSMUSG00000018486.2 | WNT9B | 4.301332122 | 1.24E-07 | 1.49E-05 |
| ENSMUSG00000079243.4 | XIRP1 | 2.884586558 | 1.53E-04 | 0.004777193 |
| ENSMUSG00000039981.6 | ZC3H12D | 2.099201287 | 8.97E-09 | 1.68E-06 |
| ENSMUSG00000014198.15 | ZNF385C | 1.036470521 | 5.03E-04 | 0.011701837 |

### HPC

| ID | Symbol | Expr Log Ratio | Expr p-value | Expr FDR |
|---|---|---|---|---|
| ENSMUSG00000085151.7 | 1110018N20Rik | 1.268073327 | 8.19E-11 | 1.97E-09 |
| ENSMUSG00000085944.1 | 1700003D09Rik | 2.104658848 | 1.98E-12 | 6.05E-11 |
| ENSMUSG00000089730.2 | 1700007P06Rik | 1.395657501 | 2.38E-05 | 2.05E-04 |
| ENSMUSG00000051054.4 | 1700014D04Rik | 2.395834954 | 1.56E-07 | 2.20E-06 |
| ENSMUSG00000085549.7 | 1700047F07Rik | 4.641543582 | 3.70E-11 | 9.48E-10 |
| ENSMUSG00000100599.2 | 1700120C14Rik | 1.07489109 | 1.20E-06 | 1.40E-05 |
| ENSMUSG00000051606.5 | 2010001K21Rik | -1.568857188 | 4.51E-04 | 0.002703207 |
| ENSMUSG00000097178.7 | 2310002F09Rik | -2.117053786 | 1.64E-10 | 3.75E-09 |
| ENSMUSG00000085338.1 | 2410004I01Rik | 1.272673399 | 2.87E-04 | 0.001841028 |
| ENSMUSG00000051297.8 | 2410124H12Rik | -2.308053013 | 0.002653291 | 0.01212261 |
| ENSMUSG00000102386.1 | 2900022M07Rik | 1.613884791 | 0.00390909 | 0.016763103 |
| ENSMUSG00000051339.10 | 2900026A02Rik | -1.1891594 | 1.59E-20 | 1.39E-18 |
| ENSMUSG00000109162.1 | 2900027M19Rik | 1.265922725 | 0.003091956 | 0.013792697 |
| ENSMUSG00000115625.1 | 2900040C04Rik | -2.842163039 | 0.003331883 | 0.014672723 |
| ENSMUSG00000085642.8 | 3110053B16Rik | 1.865603016 | 0.014476884 | 0.049096891 |
| ENSMUSG00000107585.1 | 3300002P13Rik | -1.045770234 | 1.32E-04 | 9.34E-04 |
| ENSMUSG00000086943.2 | 4732414G09Rik | 1.248557654 | 8.59E-07 | 1.04E-05 |
| ENSMUSG00000108456.1 | 4732496C06Rik | 1.111599677 | 4.87E-05 | 3.88E-04 |
| ENSMUSG00000115936.1 | 4833415N18Rik | 3.91840827 | 5.47E-10 | 1.15E-08 |
| ENSMUSG00000104586.1 | 4921539H07Rik | -2.042113925 | 1.98E-12 | 6.05E-11 |
| ENSMUSG00000114953.1 | 4930519K11Rik | 1.2472458 | 2.79E-05 | 2.37E-04 |
| ENSMUSG00000097587.2 | 4930578M01Rik | -1.954581644 | 8.77E-17 | 4.98E-15 |
| ENSMUSG00000097216.3 | 4932441J04Rik | 1.043392446 | 4.29E-07 | 5.56E-06 |
| ENSMUSG00000110427.1 | 4933406B17Rik | -3.555429437 | 5.56E-16 | 2.81E-14 |
| ENSMUSG00000113960.1 | 4933412O06Rik | -1.62344731 | 9.39E-10 | 1.90E-08 |
| ENSMUSG00000105203.1 | 4933428P19Rik | -1.009876907 | 1.88E-05 | 1.67E-04 |
| ENSMUSG00000085129.1 | 5031425F14Rik | 2.290790858 | 8.85E-05 | 6.57E-04 |
| ENSMUSG00000086822.2 | 5330413P13Rik | 1.175077642 | 3.96E-07 | 5.17E-06 |
| ENSMUSG00000054944.7 | 5330416C01Rik | 1.827504943 | 5.13E-11 | 1.28E-09 |
| ENSMUSG00000110140.1 | 5430421F17Rik | 2.660894312 | 1.99E-08 | 3.28E-07 |
| ENSMUSG00000103477.1 | 5930409G06Rik | 1.726891754 | 4.06E-11 | 1.03E-09 |
| ENSMUSG00000114590.1 | 5930438M14Rik | 1.217245577 | 0.006340386 | 0.025001224 |
| ENSMUSG00000110067.2 | 6330411D24Rik | 1.809026528 | 3.08E-06 | 3.32E-05 |
| ENSMUSG00000109898.1 | 6330420H09Rik | -5.787506947 | 4.45E-21 | 4.20E-19 |
| ENSMUSG00000108228.2 | 6430584L05Rik | 1.527384852 | 9.95E-15 | 4.27E-13 |
| ENSMUSG00000034185.9 | 6430628N08Rik | 2.711128198 | 7.56E-04 | 0.004186359 |
| ENSMUSG00000073067.10 | 9130019P16Rik | -1.089992221 | 4.04E-05 | 3.29E-04 |
| ENSMUSG00000087022.4 | 9130024F11Rik | 2.37843809 | 2.14E-21 | 2.16E-19 |
| ENSMUSG00000108242.1 | 9330118I20Rik | 1.195188241 | 2.31E-07 | 3.16E-06 |
| ENSMUSG00000103502.1 | 9330121J05Rik | -1.413795338 | 1.91E-12 | 5.86E-11 |
| ENSMUSG00000056031.13 | 9330154J02Rik | 1.385691844 | 2.62E-04 | 0.001700105 |
| ENSMUSG00000109536.1 | 9330162G02Rik | 1.346255292 | 1.31E-06 | 1.52E-05 |
| ENSMUSG00000054457.5 | 9430021M05Rik | 1.095141296 | 1.53E-06 | 1.76E-05 |

| | | | | |
|---|---|---|---|---|
| ENSMUSG00000097462.7 | 9530026P05Rik | -1.240194815 | 1.91E-08 | 3.18E-07 |
| ENSMUSG00000117916.1 | 9630028I04Rik | 2.536374089 | 2.92E-15 | 1.35E-13 |
| ENSMUSG00000097493.3 | 9930014A18Rik | -1.384050621 | 2.26E-11 | 5.97E-10 |
| ENSMUSG00000115276.1 | 9930017N22Rik | 1.069944963 | 0.008431029 | 0.03162881 |
| ENSMUSG00000109394.2 | A230057D06Rik | 1.169109922 | 8.39E-07 | 1.02E-05 |
| ENSMUSG00000109122.1 | A230103L15Rik | 1.880222698 | 4.10E-04 | 0.002491422 |
| ENSMUSG00000030111.9 | A2M | -1.44499845 | 1.44E-05 | 1.32E-04 |
| ENSMUSG00000052014.6 | A330070K13Rik | -1.220377424 | 5.29E-04 | 0.003096343 |
| ENSMUSG00000109321.1 | A330076H08Rik | 1.15205739 | 1.63E-09 | 3.16E-08 |
| ENSMUSG00000097666.2 | A330094K24Rik | 2.646842613 | 1.18E-12 | 3.75E-11 |
| ENSMUSG00000087305.8 | A430035B10Rik | 1.07233147 | 1.32E-08 | 2.26E-07 |
| ENSMUSG00000047878.13 | A4GALT | 1.134406425 | 3.28E-05 | 2.74E-04 |
| ENSMUSG00000045238.9 | A730035I17Rik | -1.092757686 | 1.36E-06 | 1.58E-05 |
| ENSMUSG00000103529.1 | A730089K16Rik | -1.073252329 | 1.88E-04 | 0.001270358 |
| ENSMUSG00000097182.2 | A830009L08Rik | 3.24021612 | 7.92E-24 | 1.08E-21 |
| ENSMUSG00000084890.1 | A830036E02Rik | 2.52848725 | 5.38E-10 | 1.13E-08 |
| ENSMUSG00000091890.2 | A830073O21Rik | 1.078709066 | 1.52E-06 | 1.76E-05 |
| ENSMUSG00000114961.1 | A930002C04Rik | 3.176606358 | 1.08E-18 | 7.44E-17 |
| ENSMUSG00000056735.5 | A930024E05Rik | -1.223227488 | 1.85E-12 | 5.68E-11 |
| ENSMUSG00000113342.1 | AA414992 | 2.900943855 | 8.63E-14 | 3.16E-12 |
| ENSMUSG00000085042.8 | Abhd11os | 1.443572131 | 1.51E-05 | 1.38E-04 |
| ENSMUSG00000038459.10 | ABHD17C | -1.099579266 | 3.34E-27 | 6.35E-25 |
| ENSMUSG00000035258.15 | Abi3bp | 2.85510089 | 7.54E-12 | 2.14E-10 |
| ENSMUSG00000116226.1 | AC105358.1 | 1.163357921 | 3.36E-04 | 0.002101393 |
| ENSMUSG00000117901.1 | AC109619.1 | 1.522030047 | 2.14E-10 | 4.82E-09 |
| ENSMUSG00000115406.1 | AC115735.4 | 2.089865175 | 7.23E-17 | 4.13E-15 |
| ENSMUSG00000113055.1 | AC115910.1 | 3.002187386 | 5.28E-07 | 6.72E-06 |
| ENSMUSG00000114193.1 | AC122828.4 | 1.163424561 | 0.008522075 | 0.031948592 |
| ENSMUSG00000116610.1 | AC129186.1 | -1.054239819 | 0.003959842 | 0.01695872 |
| ENSMUSG00000115810.1 | AC132863.2 | 1.259732719 | 3.40E-11 | 8.75E-10 |
| ENSMUSG00000118550.1 | AC135859.1 | 1.427226156 | 0.002079816 | 0.009869563 |
| ENSMUSG00000118094.1 | AC142100.3 | 1.840022265 | 7.17E-06 | 7.05E-05 |
| ENSMUSG00000115373.1 | AC154575.1 | 1.274054188 | 4.34E-04 | 0.002614923 |
| ENSMUSG00000114624.1 | AC156025.2 | 1.836898984 | 0.001008012 | 0.005330229 |
| ENSMUSG00000114709.1 | AC157650.3 | 1.046729544 | 0.006536396 | 0.025676031 |
| ENSMUSG00000114970.1 | AC164978.1 | 1.586354255 | 2.39E-04 | 0.001568985 |
| ENSMUSG00000116081.1 | AC165278.1 | 1.369446394 | 1.02E-05 | 9.68E-05 |
| ENSMUSG00000116243.1 | AC165278.2 | 2.194238944 | 8.27E-13 | 2.67E-11 |
| ENSMUSG00000116766.1 | AC171205.1 | 2.134663424 | 2.41E-04 | 0.001579428 |
| ENSMUSG00000113309.1 | AC173210.2 | -1.367538524 | 2.25E-06 | 2.49E-05 |
| ENSMUSG00000010651.4 | Acaa1b | 2.291471726 | 5.17E-23 | 6.42E-21 |
| ENSMUSG00000038007.14 | ACER2 | 1.282257156 | 3.94E-05 | 3.22E-04 |
| ENSMUSG00000044534.8 | ACKR2 | 1.176896585 | 2.83E-06 | 3.07E-05 |
| ENSMUSG00000042540.12 | Acot5 | -1.701266809 | 5.46E-09 | 9.90E-08 |
| ENSMUSG00000011256.16 | ADAM19 | 1.636025706 | 2.73E-11 | 7.10E-10 |

| | | | | |
|---|---|---|---|---|
| ENSMUSG00000027318.17 | ADAM33 | 3.081454244 | 2.42E-11 | 6.35E-10 |
| ENSMUSG00000059901.12 | ADAMTS14 | -1.355900744 | 1.19E-04 | 8.51E-04 |
| ENSMUSG00000049538.14 | ADAMTS16 | 1.244200637 | 1.26E-04 | 8.93E-04 |
| ENSMUSG00000053399.9 | ADAMTS18 | 1.771655262 | 9.16E-06 | 8.80E-05 |
| ENSMUSG00000022894.6 | ADAMTS5 | 1.227584365 | 1.37E-05 | 1.27E-04 |
| ENSMUSG00000043822.18 | ADAMTSL5 | 1.068248995 | 3.49E-08 | 5.48E-07 |
| ENSMUSG00000024256.7 | ADCYAP1 | 3.146128587 | 1.47E-31 | 3.99E-29 |
| ENSMUSG00000044017.16 | ADGRD1 | 2.779416994 | 3.42E-09 | 6.38E-08 |
| ENSMUSG00000023918.12 | ADGRF4 | 1.322674197 | 4.53E-04 | 0.00271432 |
| ENSMUSG00000030790.15 | ADM | 1.971936597 | 4.61E-05 | 3.69E-04 |
| ENSMUSG00000020178.6 | ADORA2A | 3.890067907 | 1.51E-25 | 2.62E-23 |
| ENSMUSG00000031448.12 | ADPRHL1 | 1.049897947 | 0.004722508 | 0.019560445 |
| ENSMUSG00000050541.14 | ADRA1B | 2.821200819 | 2.76E-40 | 1.43E-37 |
| ENSMUSG00000027335.9 | ADRA1D | -1.02102619 | 1.27E-07 | 1.82E-06 |
| ENSMUSG00000031489.15 | ADRB3 | 1.878831347 | 1.04E-09 | 2.10E-08 |
| ENSMUSG00000040706.4 | AGMAT | 3.586702638 | 4.07E-21 | 3.89E-19 |
| ENSMUSG00000031980.10 | AGT | -1.325184476 | 6.67E-08 | 1.00E-06 |
| ENSMUSG00000072812.4 | AHNAK2 | 1.775815339 | 4.31E-08 | 6.64E-07 |
| ENSMUSG00000084826.7 | AI847159 | 2.25656418 | 9.12E-08 | 1.34E-06 |
| ENSMUSG00000091415.5 | AK9 | -1.422621541 | 6.56E-04 | 0.003716444 |
| ENSMUSG00000029762.6 | AKR1B10 | 2.946226283 | 9.19E-07 | 1.11E-05 |
| ENSMUSG00000021214.14 | AKR1C3 | 1.899283464 | 8.32E-09 | 1.46E-07 |
| ENSMUSG00000118394.1 | AL607088.1 | 4.275760307 | 8.20E-18 | 5.21E-16 |
| ENSMUSG00000019102.10 | ALDH3A1 | 4.143365255 | 1.77E-39 | 8.13E-37 |
| ENSMUSG00000020636.14 | ALLC | 1.821662451 | 8.63E-07 | 1.04E-05 |
| ENSMUSG00000032807.5 | ALOX12B | -2.238429844 | 2.13E-10 | 4.82E-09 |
| ENSMUSG00000025701.12 | ALOX5 | 2.001810406 | 8.05E-09 | 1.42E-07 |
| ENSMUSG00000048218.6 | AMIGO2 | -1.967738199 | 1.99E-23 | 2.55E-21 |
| ENSMUSG00000031465.6 | ANGPT2 | 1.530800159 | 0.001756359 | 0.008558429 |
| ENSMUSG00000047606.4 | ANKRD34C | 1.723216468 | 2.09E-11 | 5.56E-10 |
| ENSMUSG00000021866.15 | ANXA11 | -1.007765966 | 5.33E-11 | 1.33E-09 |
| ENSMUSG00000038242.12 | Aox4 | 1.080392701 | 0.001404039 | 0.007106845 |
| ENSMUSG00000003309.14 | AP1M2 | 2.631700742 | 2.09E-11 | 5.56E-10 |
| ENSMUSG00000068246.6 | Apol9a/Apol9b | 4.077097264 | 1.38E-10 | 3.22E-09 |
| ENSMUSG00000042797.9 | AQP11 | -1.119728571 | 7.07E-16 | 3.56E-14 |
| ENSMUSG00000019987.9 | ARG1 | 2.593369069 | 1.81E-04 | 0.00122891 |
| ENSMUSG00000037148.8 | ARHGAP10 | 1.749253968 | 1.06E-19 | 8.41E-18 |
| ENSMUSG00000041225.16 | ARHGAP12 | -1.400467346 | 8.30E-25 | 1.28E-22 |
| ENSMUSG00000030047.14 | ARHGAP25 | 1.575839149 | 1.51E-14 | 6.30E-13 |
| ENSMUSG00000041444.14 | ARHGAP32 | 1.273392256 | 1.39E-17 | 8.50E-16 |
| ENSMUSG00000033697.16 | ARHGAP39 | -1.099799134 | 7.66E-20 | 6.20E-18 |
| ENSMUSG00000031355.16 | ARHGAP6 | 1.184935829 | 2.89E-04 | 0.001851396 |
| ENSMUSG00000052921.13 | ARHGEF15 | 1.274343769 | 4.08E-20 | 3.41E-18 |
| ENSMUSG00000036885.14 | ARHGEF26 | -1.087540353 | 1.11E-23 | 1.48E-21 |
| ENSMUSG00000021662.11 | ARHGEF28 | -1.124844149 | 8.07E-22 | 8.65E-20 |

| | | | | |
|---|---|---|---|---|
| ENSMUSG00000051517.14 | ARHGEF39 | 2.237994312 | 2.72E-08 | 4.39E-07 |
| ENSMUSG00000031133.12 | ARHGEF6 | -1.234868 | 2.66E-18 | 1.78E-16 |
| ENSMUSG00000034936.2 | ARL4D | 1.332479754 | 0.001942846 | 0.009329504 |
| ENSMUSG00000007656.13 | ARPP19 | 2.171140177 | 2.69E-36 | 1.06E-33 |
| ENSMUSG00000032503.18 | ARPP21 | 1.539188482 | 1.37E-22 | 1.58E-20 |
| ENSMUSG00000031382.14 | ASB11 | 1.538746887 | 3.80E-04 | 0.002333881 |
| ENSMUSG00000067081.13 | ASB18 | -1.165472515 | 5.03E-06 | 5.14E-05 |
| ENSMUSG00000021200.14 | ASB2 | 1.064346746 | 2.22E-06 | 2.45E-05 |
| ENSMUSG00000009248.6 | ASCL2 | 3.874289889 | 2.03E-05 | 1.79E-04 |
| ENSMUSG00000097918.1 | ASCL5 | 2.355575 | 6.35E-09 | 1.14E-07 |
| ENSMUSG00000023017.10 | ASIC1 | 1.252516196 | 7.38E-21 | 6.77E-19 |
| ENSMUSG00000033007.15 | ASIC4 | 1.055373023 | 7.85E-07 | 9.57E-06 |
| ENSMUSG00000037686.6 | ASPG | 1.059964752 | 0.001126029 | 0.005859725 |
| ENSMUSG00000037400.17 | ATP11B | 1.086320929 | 5.24E-16 | 2.68E-14 |
| ENSMUSG00000030730.12 | ATP2A1 | 2.474598256 | 5.09E-09 | 9.26E-08 |
| ENSMUSG00000026463.17 | ATP2B4 | 1.54098631 | 1.78E-16 | 9.71E-15 |
| ENSMUSG00000034112.9 | ATP2C2 | 1.95006265 | 1.74E-05 | 1.56E-04 |
| ENSMUSG00000078958.9 | Atp6ap1l | 3.192829214 | 3.28E-30 | 7.63E-28 |
| ENSMUSG00000020566.19 | ATP6V1C2 | 1.534531436 | 2.17E-05 | 1.89E-04 |
| ENSMUSG00000109890.2 | AU023762 | 2.825833848 | 5.46E-07 | 6.92E-06 |
| ENSMUSG00000026432.7 | AVPR1B | -3.168459917 | 2.80E-12 | 8.43E-11 |
| ENSMUSG00000097414.2 | B130046B21Rik | -1.34084373 | 2.71E-15 | 1.26E-13 |
| ENSMUSG00000097547.8 | B230110C06Rik | 1.01099965 | 9.09E-04 | 0.004881768 |
| ENSMUSG00000092492.1 | B230208B08Rik | 1.763820462 | 1.56E-08 | 2.63E-07 |
| ENSMUSG00000089706.7 | B230216N24Rik | 2.404446232 | 2.30E-13 | 7.97E-12 |
| ENSMUSG00000113701.1 | B230303A05Rik | -1.578053958 | 1.88E-13 | 6.63E-12 |
| ENSMUSG00000084843.1 | B230312C02Rik | 1.042550788 | 7.12E-05 | 5.39E-04 |
| ENSMUSG00000074892.9 | B3GALT5 | -1.30412061 | 5.88E-17 | 3.40E-15 |
| ENSMUSG00000079445.3 | B3GNT7 | 2.433167698 | 1.18E-10 | 2.78E-09 |
| ENSMUSG00000059479.7 | B3GNT8 | 1.171001382 | 3.07E-07 | 4.09E-06 |
| ENSMUSG00000046415.11 | B430212C06Rik | -2.222709328 | 7.82E-12 | 2.22E-10 |
| ENSMUSG00000098202.2 | B830012L14Rik | 1.285804154 | 0.002717615 | 0.012380331 |
| ENSMUSG00000109473.1 | B930025P03Rik | -1.361372692 | 9.87E-04 | 0.005235122 |
| ENSMUSG00000034384.11 | BARHL2 | 1.240606326 | 8.95E-04 | 0.004821469 |
| ENSMUSG00000032033.11 | BARX2 | 2.691331242 | 3.54E-16 | 1.86E-14 |
| ENSMUSG00000035021.13 | BAZ1A | 2.561676747 | 9.33E-07 | 1.12E-05 |
| ENSMUSG00000085041.1 | BB031773 | 2.973127206 | 4.09E-11 | 1.04E-09 |
| ENSMUSG00000041660.8 | BBOX1 | -1.598945237 | 2.21E-10 | 4.96E-09 |
| ENSMUSG00000115783.1 | Bc1 | 1.118091968 | 1.25E-05 | 1.17E-04 |
| ENSMUSG00000048251.15 | BCL11B | -1.025667333 | 2.04E-06 | 2.28E-05 |
| ENSMUSG00000000317.11 | BCL6B | 1.32761841 | 0.011759644 | 0.041628888 |
| ENSMUSG00000040363.15 | BCOR | 1.369481464 | 6.91E-05 | 5.27E-04 |
| ENSMUSG00000021070.6 | BDKRB2 | 1.450374698 | 0.002905283 | 0.013083677 |
| ENSMUSG00000048482.14 | BDNF | 2.267717126 | 8.97E-06 | 8.63E-05 |
| ENSMUSG00000020169.4 | BEST3 | -2.09330009 | 2.45E-08 | 3.99E-07 |

| | | | | |
|---|---|---|---|---|
| ENSMUSG00000074489.9 | BGLAP | 3.163865637 | 6.91E-10 | 1.43E-08 |
| ENSMUSG00000044243.3 | BHLHA9 | -1.030141822 | 3.14E-05 | 2.63E-04 |
| ENSMUSG00000025128.7 | BHLHE22 | -2.017120742 | 5.43E-31 | 1.34E-28 |
| ENSMUSG00000061132.13 | BLNK | 1.834732228 | 1.02E-09 | 2.06E-08 |
| ENSMUSG00000029335.5 | BMP3 | 3.105824381 | 1.83E-18 | 1.23E-16 |
| ENSMUSG00000032726.12 | BMP8A | 2.978660747 | 2.32E-07 | 3.17E-06 |
| ENSMUSG00000028115.16 | BNIPL | 1.548421468 | 0.00448192 | 0.018771097 |
| ENSMUSG00000027485.15 | BPIFB1 | 2.340373362 | 3.52E-08 | 5.49E-07 |
| ENSMUSG00000035131.14 | BRINP3 | 1.556794058 | 4.26E-31 | 1.08E-28 |
| ENSMUSG00000040298.6 | BTBD16 | -1.872203829 | 5.24E-08 | 7.98E-07 |
| ENSMUSG00000062098.11 | BTBD3 | -1.029400789 | 4.67E-22 | 5.07E-20 |
| ENSMUSG00000117926.1 | C030004G16Rik | 2.064872033 | 3.28E-04 | 0.002056547 |
| ENSMUSG00000086600.8 | C030005K06Rik | 1.065943998 | 0.001507286 | 0.007526861 |
| ENSMUSG00000047642.15 | C12orf56 | 2.518856196 | 4.94E-11 | 1.24E-09 |
| ENSMUSG00000021098.14 | C14orf39 | 1.344335952 | 4.49E-05 | 3.59E-04 |
| ENSMUSG00000042460.5 | C1GALT1 | 1.315568307 | 1.75E-09 | 3.39E-08 |
| ENSMUSG00000039349.5 | C1orf115 | 1.513061858 | 2.73E-17 | 1.62E-15 |
| ENSMUSG00000055172.10 | C1R | 1.293478111 | 1.25E-05 | 1.16E-04 |
| ENSMUSG00000038521.17 | C1S | 1.021143977 | 0.001911076 | 0.009195662 |
| ENSMUSG00000024164.15 | C3 | -1.264455121 | 0.003846116 | 0.016554412 |
| ENSMUSG00000026405.14 | C4bp | -3.211365201 | 6.80E-12 | 1.95E-10 |
| ENSMUSG00000074737.2 | C530025M09Rik | 1.219973671 | 3.97E-06 | 4.16E-05 |
| ENSMUSG00000112663.1 | C630031E19Rik | -4.234595492 | 6.54E-10 | 1.36E-08 |
| ENSMUSG00000079105.4 | C7 | 2.300956026 | 1.32E-08 | 2.25E-07 |
| ENSMUSG00000114469.2 | C730002L08Rik | 3.127464407 | 3.81E-21 | 3.66E-19 |
| ENSMUSG00000040978.4 | C7orf57 | -1.472729415 | 2.03E-05 | 1.79E-04 |
| ENSMUSG00000056158.14 | CA10 | 1.669386832 | 9.28E-26 | 1.65E-23 |
| ENSMUSG00000032373.15 | CA12 | -1.256420476 | 6.14E-04 | 0.003513917 |
| ENSMUSG00000041261.9 | CA8 | 1.406096829 | 9.28E-09 | 1.62E-07 |
| ENSMUSG00000028463.14 | CA9 | 1.201459584 | 0.001557909 | 0.007739863 |
| ENSMUSG00000009075.2 | CABP7 | -3.238408213 | 2.10E-42 | 1.12E-39 |
| ENSMUSG00000028532.14 | CACHD1 | -1.655617266 | 5.33E-28 | 1.07E-25 |
| ENSMUSG00000020866.17 | CACNA1G | 1.224017925 | 8.52E-10 | 1.74E-08 |
| ENSMUSG00000024112.16 | CACNA1H | -1.019807706 | 8.90E-07 | 1.07E-05 |
| ENSMUSG00000010066.15 | CACNA2D2 | 1.424041336 | 1.75E-15 | 8.54E-14 |
| ENSMUSG00000066189.9 | CACNG3 | 1.104653119 | 1.12E-07 | 1.63E-06 |
| ENSMUSG00000053395.15 | CACNG8 | -1.109363904 | 2.80E-16 | 1.49E-14 |
| ENSMUSG00000017978.18 | CADPS2 | -1.255805846 | 5.16E-21 | 4.78E-19 |
| ENSMUSG00000003657.9 | CALB2 | -1.880811735 | 5.67E-07 | 7.15E-06 |
| ENSMUSG00000023964.15 | CALCR | 2.704708364 | 2.12E-04 | 0.001415077 |
| ENSMUSG00000049872.8 | CALHM5 | -1.168794773 | 2.73E-07 | 3.66E-06 |
| ENSMUSG00000032246.14 | CALML4 | -1.94970134 | 0.006004563 | 0.023911107 |
| ENSMUSG00000030272.15 | CAMK1 | 1.122611314 | 3.82E-18 | 2.51E-16 |
| ENSMUSG00000024617.16 | CAMK2A | -1.338831743 | 5.62E-14 | 2.12E-12 |
| ENSMUSG00000053819.16 | CAMK2D | 1.696840158 | 2.23E-28 | 4.60E-26 |

| | | | | |
|---|---|---|---|---|
| ENSMUSG00000046447.3 | CAMK2N1 | 1.072280006 | 1.03E-11 | 2.89E-10 |
| ENSMUSG00000032936.13 | CAMKV | -1.150016057 | 3.76E-09 | 6.97E-08 |
| ENSMUSG00000040712.16 | CAMTA2 | -1.251312434 | 1.98E-16 | 1.07E-14 |
| ENSMUSG00000097638.1 | Carlr | 1.331340473 | 2.48E-07 | 3.36E-06 |
| ENSMUSG00000025888.6 | CASP1 | 1.021087257 | 1.10E-04 | 7.92E-04 |
| ENSMUSG00000051980.13 | CASR | 1.801064978 | 0.011253728 | 0.040173816 |
| ENSMUSG00000028977.17 | CASZ1 | 1.262296992 | 7.15E-11 | 1.74E-09 |
| ENSMUSG00000050623.5 | CATSPERZ | 1.649908388 | 8.08E-06 | 7.87E-05 |
| ENSMUSG00000028348.7 | CAVIN4 | 1.780913887 | 2.23E-04 | 0.001477995 |
| ENSMUSG00000006362.16 | CBFA2T3 | -1.003621974 | 4.13E-07 | 5.36E-06 |
| ENSMUSG00000031654.16 | CBLN1 | 1.632710389 | 1.25E-10 | 2.91E-09 |
| ENSMUSG00000024647.14 | CBLN2 | 1.218332636 | 2.71E-08 | 4.39E-07 |
| ENSMUSG00000067578.3 | CBLN4 | 1.401231548 | 5.54E-06 | 5.61E-05 |
| ENSMUSG00000046318.16 | CCBE1 | -2.93008069 | 7.74E-32 | 2.13E-29 |
| ENSMUSG00000022768.17 | CCDC116 | 1.0343958 | 2.14E-07 | 2.94E-06 |
| ENSMUSG00000050625.4 | CCDC121 | -1.01690242 | 0.001191615 | 0.006157173 |
| ENSMUSG00000019767.12 | CCDC170 | -1.116180787 | 0.005352673 | 0.021739805 |
| ENSMUSG00000029875.5 | CCDC184 | 1.211135347 | 7.43E-07 | 9.11E-06 |
| ENSMUSG00000048038.8 | CCDC187 | 1.236683913 | 9.47E-04 | 0.005063605 |
| ENSMUSG00000108900.1 | Ccdc194 | -1.824081163 | 2.85E-04 | 0.001831272 |
| ENSMUSG00000090946.3 | CCDC71L | -1.165296003 | 6.23E-15 | 2.75E-13 |
| ENSMUSG00000022665.15 | CCDC80 | 1.18805178 | 4.74E-04 | 0.002816424 |
| ENSMUSG00000032878.16 | CCDC85A | -1.011381713 | 8.74E-11 | 2.09E-09 |
| ENSMUSG00000096257.2 | CCER2 | 1.655388234 | 2.63E-09 | 4.96E-08 |
| ENSMUSG00000030898.15 | CCKBR | 2.56373619 | 2.06E-62 | 2.43E-59 |
| ENSMUSG00000031780.2 | CCL17 | 1.018242197 | 1.68E-04 | 0.00115068 |
| ENSMUSG00000019997.11 | CCN2 | 1.86202068 | 3.88E-06 | 4.08E-05 |
| ENSMUSG00000005124.10 | CCN4 | 1.777572802 | 1.19E-05 | 1.11E-04 |
| ENSMUSG00000027793.6 | CCNA1 | -1.370356202 | 7.02E-07 | 8.67E-06 |
| ENSMUSG00000028212.16 | CCNE2 | 1.075603358 | 4.28E-07 | 5.55E-06 |
| ENSMUSG00000072082.7 | CCNF | 1.589464644 | 1.52E-05 | 1.38E-04 |
| ENSMUSG00000044707.7 | Ccnjl | -1.802613926 | 4.37E-15 | 1.99E-13 |
| ENSMUSG00000042417.5 | CCNO | -1.061016587 | 8.69E-04 | 0.004703127 |
| ENSMUSG00000046186.8 | CD109 | -1.945638412 | 8.92E-09 | 1.56E-07 |
| ENSMUSG00000028865.14 | CD164L2 | 1.6405094 | 1.12E-12 | 3.55E-11 |
| ENSMUSG00000028076.12 | CD1D | 1.936765099 | 2.86E-06 | 3.10E-05 |
| ENSMUSG00000022667.18 | CD200R1 | 3.256143605 | 5.26E-14 | 1.99E-12 |
| ENSMUSG00000026012.2 | CD28 | 4.381853164 | 4.64E-20 | 3.85E-18 |
| ENSMUSG00000029084.5 | CD38 | -1.448760547 | 1.80E-16 | 9.80E-15 |
| ENSMUSG00000023274.14 | CD4 | 4.889674179 | 1.24E-10 | 2.90E-09 |
| ENSMUSG00000005087.17 | CD44 | 1.206769532 | 0.006724762 | 0.026259807 |
| ENSMUSG00000024670.17 | CD6 | 1.987862787 | 5.29E-11 | 1.32E-09 |
| ENSMUSG00000025163.6 | CD7 | 4.377819585 | 5.57E-21 | 5.14E-19 |
| ENSMUSG00000040452.9 | CDH12 | 1.182717713 | 1.13E-10 | 2.66E-09 |
| ENSMUSG00000040420.15 | CDH18 | 1.425889374 | 2.33E-26 | 4.23E-24 |

| | | | | |
|---|---|---|---|---|
| ENSMUSG00000053166.14 | CDH22 | 1.069587068 | 1.19E-09 | 2.36E-08 |
| ENSMUSG00000012819.15 | CDH23 | 1.020288654 | 7.11E-04 | 0.003971043 |
| ENSMUSG00000059674.7 | CDH24 | -1.010491823 | 1.19E-04 | 8.55E-04 |
| ENSMUSG00000039155.15 | CDH26 | -1.910359704 | 0.001585641 | 0.007855547 |
| ENSMUSG00000061048.8 | CDH3 | -1.532353572 | 0.007814568 | 0.029721276 |
| ENSMUSG00000000305.12 | CDH4 | 1.127278761 | 1.30E-08 | 2.23E-07 |
| ENSMUSG00000039385.5 | CDH6 | 1.323095001 | 1.73E-11 | 4.70E-10 |
| ENSMUSG00000026312.17 | CDH7 | 1.768989821 | 7.20E-19 | 5.08E-17 |
| ENSMUSG00000026023.16 | CDK15 | 2.219481993 | 1.09E-11 | 3.04E-10 |
| ENSMUSG00000023067.14 | CDKN1A | 3.365531417 | 4.80E-09 | 8.77E-08 |
| ENSMUSG00000037628.10 | CDKN3 | 3.510371927 | 3.37E-08 | 5.31E-07 |
| ENSMUSG00000039518.7 | Cdsn | 2.227114252 | 4.80E-12 | 1.40E-10 |
| ENSMUSG00000006585.3 | CDT1 | 1.122186854 | 2.11E-04 | 0.001408905 |
| ENSMUSG00000071226.11 | CECR2 | -1.592685005 | 3.13E-14 | 1.24E-12 |
| ENSMUSG00000029177.9 | CENPA | 1.522521021 | 1.68E-04 | 0.00115068 |
| ENSMUSG00000045328.11 | CENPE | -1.026819157 | 7.04E-04 | 0.003937478 |
| ENSMUSG00000076433.4 | CEP295NL | 4.553379069 | 3.72E-13 | 1.26E-11 |
| ENSMUSG00000075256.9 | CERKL | -1.632782724 | 3.79E-12 | 1.12E-10 |
| ENSMUSG00000026649.14 | CFAP126 | -1.128675226 | 0.003775414 | 0.016282891 |
| ENSMUSG00000071550.14 | CFAP44 | -1.033212726 | 1.10E-04 | 7.95E-04 |
| ENSMUSG00000020904.11 | CFAP52 | -1.530207744 | 0.001759777 | 0.008572556 |
| ENSMUSG00000079502.8 | CFAP77 | -1.161321476 | 0.009138162 | 0.033828201 |
| ENSMUSG00000027313.3 | CHAC1 | 1.173047942 | 0.005110312 | 0.020878698 |
| ENSMUSG00000021919.8 | CHAT | 3.805366657 | 2.05E-06 | 2.29E-05 |
| ENSMUSG00000021194.6 | CHGA | 1.512673634 | 3.74E-29 | 7.81E-27 |
| ENSMUSG00000078185.3 | CHML | 1.006288295 | 7.85E-09 | 1.39E-07 |
| ENSMUSG00000022860.15 | CHODL | 1.490430536 | 2.54E-04 | 0.001657784 |
| ENSMUSG00000031283.16 | CHRDL1 | 2.04401533 | 1.26E-33 | 3.85E-31 |
| ENSMUSG00000045613.9 | CHRM2 | 1.342706517 | 2.88E-10 | 6.32E-09 |
| ENSMUSG00000074939.3 | CHRM5 | -1.535368415 | 9.47E-05 | 6.95E-04 |
| ENSMUSG00000027107.3 | CHRNA1 | -1.095482286 | 5.97E-04 | 0.003437685 |
| ENSMUSG00000022041.10 | CHRNA2 | 2.103875317 | 1.57E-08 | 2.64E-07 |
| ENSMUSG00000030525.8 | CHRNA7 | -1.276423052 | 3.62E-21 | 3.52E-19 |
| ENSMUSG00000031492.13 | CHRNB3 | 2.50221587 | 1.70E-06 | 1.93E-05 |
| ENSMUSG00000060402.8 | CHST8 | -1.479793158 | 8.17E-20 | 6.58E-18 |
| ENSMUSG00000047161.12 | CHST9 | -1.223840038 | 0.011967515 | 0.042228963 |
| ENSMUSG00000058152.8 | CHSY3 | 1.341884022 | 1.48E-14 | 6.19E-13 |
| ENSMUSG00000042269.15 | CIBAR2 | 1.709270331 | 5.58E-07 | 7.05E-06 |
| ENSMUSG00000032578.7 | CISH | 1.02557147 | 1.88E-06 | 2.12E-05 |
| ENSMUSG00000029516.19 | CIT | 1.405684048 | 4.11E-15 | 1.88E-13 |
| ENSMUSG00000070803.6 | CITED4 | 1.649428777 | 6.06E-14 | 2.26E-12 |
| ENSMUSG00000054400.16 | CKLF | 1.011389032 | 4.86E-07 | 6.24E-06 |
| ENSMUSG00000037594.11 | CLBA1 | 1.200639168 | 1.50E-05 | 1.37E-04 |
| ENSMUSG00000047230.6 | CLDN2 | -2.79333856 | 7.69E-04 | 0.004245731 |
| ENSMUSG00000038064.7 | CLDN22 | -2.286765422 | 4.97E-08 | 7.61E-07 |

| | | | | |
|---|---|---|---|---|
| ENSMUSG00000055976.6 | CLDN23 | 1.076376354 | 5.69E-04 | 0.003296234 |
| ENSMUSG00000033633.15 | CLEC18B | 2.554504182 | 1.77E-13 | 6.28E-12 |
| ENSMUSG00000033082.10 | CLEC1A | -2.015835571 | 9.84E-10 | 1.99E-08 |
| ENSMUSG00000022949.9 | CLIC6 | -1.699160883 | 0.006141727 | 0.024357466 |
| ENSMUSG00000021097.15 | CLMN | -1.650729151 | 3.94E-25 | 6.44E-23 |
| ENSMUSG00000043850.15 | CLRN1 | -2.069570209 | 1.28E-07 | 1.83E-06 |
| ENSMUSG00000024873.7 | CNIH2 | -1.575449155 | 2.15E-43 | 1.27E-40 |
| ENSMUSG00000022025.13 | CNMD | -1.723770992 | 1.65E-04 | 0.001135823 |
| ENSMUSG00000039488.15 | CNTN5 | 1.105485684 | 5.82E-06 | 5.86E-05 |
| ENSMUSG00000030092.14 | CNTN6 | 1.961787592 | 6.30E-27 | 1.17E-24 |
| ENSMUSG00000038048.8 | Cntnap5c | -1.287843683 | 4.84E-11 | 1.22E-09 |
| ENSMUSG00000020173.17 | COBL | 2.786193707 | 2.00E-42 | 1.10E-39 |
| ENSMUSG00000020953.17 | COCH | 3.028822483 | 1.61E-24 | 2.37E-22 |
| ENSMUSG00000039462.4 | COL10A1 | 5.583992368 | 6.21E-11 | 1.52E-09 |
| ENSMUSG00000025064.15 | Col17a1 | 1.853053348 | 1.96E-07 | 2.71E-06 |
| ENSMUSG00000028197.4 | COL24A1 | 3.352880936 | 4.76E-11 | 1.20E-09 |
| ENSMUSG00000058897.18 | COL25A1 | -1.129722219 | 3.41E-09 | 6.36E-08 |
| ENSMUSG00000045672.15 | COL27A1 | 2.055863068 | 1.13E-07 | 1.64E-06 |
| ENSMUSG00000068794.7 | COL28A1 | 1.36282185 | 6.01E-07 | 7.54E-06 |
| ENSMUSG00000031503.13 | COL4A2 | 1.083590609 | 4.91E-05 | 3.90E-04 |
| ENSMUSG00000026837.16 | COL5A1 | 2.090326875 | 1.33E-09 | 2.63E-08 |
| ENSMUSG00000004098.7 | COL5A3 | 2.27333964 | 1.16E-07 | 1.67E-06 |
| ENSMUSG00000001119.7 | COL6A1 | 1.312665811 | 3.94E-04 | 0.002407073 |
| ENSMUSG00000032572.9 | Col6a4 | 1.822323403 | 8.64E-08 | 1.28E-06 |
| ENSMUSG00000056174.8 | COL8A2 | -1.856742316 | 0.001201152 | 0.006198696 |
| ENSMUSG00000028626.5 | COL9A2 | 1.07478586 | 2.50E-06 | 2.75E-05 |
| ENSMUSG00000057606.14 | COLQ | 1.350977187 | 1.66E-04 | 0.001135823 |
| ENSMUSG00000025981.13 | COQ10B | 1.205349501 | 5.99E-08 | 9.07E-07 |
| ENSMUSG00000020836.15 | CORO6 | 1.875349924 | 2.25E-08 | 3.68E-07 |
| ENSMUSG00000025488.9 | Cox8b | -1.67233466 | 0.003521015 | 0.015346014 |
| ENSMUSG00000039070.5 | CPA4 | 3.438009851 | 8.75E-13 | 2.82E-11 |
| ENSMUSG00000042501.12 | CPA6 | 1.28317765 | 0.014261613 | 0.048626605 |
| ENSMUSG00000039714.9 | CPLX3 | 2.040284887 | 1.64E-16 | 8.95E-15 |
| ENSMUSG00000032564.16 | CPNE4 | -1.762665834 | 1.99E-19 | 1.50E-17 |
| ENSMUSG00000024008.16 | CPNE5 | 2.32591444 | 1.21E-22 | 1.42E-20 |
| ENSMUSG00000022212.15 | CPNE6 | -1.79799023 | 4.66E-22 | 5.07E-20 |
| ENSMUSG00000034796.14 | CPNE7 | -1.68419693 | 1.24E-08 | 2.13E-07 |
| ENSMUSG00000026090.16 | CRACDL | -1.013801141 | 4.77E-09 | 8.72E-08 |
| ENSMUSG00000089978.1 | Crb1-ps | 2.705790816 | 1.60E-05 | 1.45E-04 |
| ENSMUSG00000035041.8 | CREB3L3 | -1.649955635 | 6.14E-07 | 7.69E-06 |
| ENSMUSG00000049796.4 | CRH | 2.512392955 | 2.06E-14 | 8.43E-13 |
| ENSMUSG00000021680.8 | CRHBP | 1.692647288 | 9.11E-11 | 2.18E-09 |
| ENSMUSG00000006546.3 | CRYBA2 | 4.481980876 | 3.49E-14 | 1.36E-12 |
| ENSMUSG00000012123.17 | CRYBG2 | 2.657887613 | 4.14E-17 | 2.42E-15 |
| ENSMUSG00000030905.5 | CRYM | -2.216208189 | 1.88E-05 | 1.67E-04 |

| | | | | |
|---|---|---|---|---|
| ENSMUSG00000024846.6 | CST6 | 1.060867306 | 3.63E-08 | 5.64E-07 |
| ENSMUSG00000079597.2 | CSTA | 2.186660804 | 6.18E-05 | 4.80E-04 |
| ENSMUSG00000117220.1 | CT025671.1 | 3.705315457 | 3.73E-29 | 7.81E-27 |
| ENSMUSG00000030970.16 | CTBP2 | -1.176947735 | 4.27E-23 | 5.39E-21 |
| ENSMUSG00000044258.10 | Ctla2a/Ctla2b | 1.496471627 | 1.70E-04 | 0.001161696 |
| ENSMUSG00000028111.4 | CTSK | 1.175448292 | 1.57E-06 | 1.81E-05 |
| ENSMUSG00000042589.18 | CUX2 | 2.665049525 | 2.69E-45 | 1.78E-42 |
| ENSMUSG00000019590.16 | CYB561 | 1.319127907 | 3.86E-25 | 6.38E-23 |
| ENSMUSG00000027015.4 | CYBRD1 | -1.195217533 | 6.29E-06 | 6.29E-05 |
| ENSMUSG00000039519.6 | CYP7B1 | -1.322210078 | 4.94E-23 | 6.18E-21 |
| ENSMUSG00000062563.15 | CYS1 | -1.366670053 | 2.41E-07 | 3.28E-06 |
| ENSMUSG00000092130.2 | D030025P21Rik | 3.808644159 | 0.00953851 | 0.035082173 |
| ENSMUSG00000086296.2 | D030055H07Rik | 2.238511651 | 2.73E-13 | 9.33E-12 |
| ENSMUSG00000102260.1 | D330025C20Rik | -3.242551407 | 6.44E-17 | 3.69E-15 |
| ENSMUSG00000109198.1 | D7Bwg0826e | 2.653767585 | 6.92E-10 | 1.43E-08 |
| ENSMUSG00000048826.7 | DACT2 | 1.220778067 | 3.42E-14 | 1.34E-12 |
| ENSMUSG00000042096.15 | DAO | -1.532620379 | 5.85E-07 | 7.37E-06 |
| ENSMUSG00000059213.7 | DDN | -1.687389747 | 8.63E-04 | 0.00466902 |
| ENSMUSG00000007379.15 | DENND2C | 1.930838944 | 4.79E-05 | 3.82E-04 |
| ENSMUSG00000027173.7 | DEPDC7 | -1.205708721 | 1.03E-06 | 1.22E-05 |
| ENSMUSG00000022419.16 | DEPTOR | 1.506494979 | 5.04E-15 | 2.26E-13 |
| ENSMUSG00000022861.17 | DGKG | -1.538636644 | 1.84E-33 | 5.54E-31 |
| ENSMUSG00000040479.12 | DGKZ | -1.110475606 | 8.86E-13 | 2.85E-11 |
| ENSMUSG00000027068.6 | DHRS9 | 3.729467791 | 2.98E-07 | 3.98E-06 |
| ENSMUSG00000040620.15 | DHX33 | -1.225238456 | 3.76E-21 | 3.63E-19 |
| ENSMUSG00000085604.1 | Dhx58os | 1.161148864 | 0.004474151 | 0.018750101 |
| ENSMUSG00000075707.5 | DIO3 | -2.830361076 | 2.87E-09 | 5.40E-08 |
| ENSMUSG00000028031.6 | DKK2 | 1.627456695 | 4.89E-04 | 0.0028887 |
| ENSMUSG00000030792.8 | DKKL1 | 3.485061692 | 4.56E-68 | 1.08E-64 |
| ENSMUSG00000047428.14 | DLK2 | 1.03639431 | 4.01E-05 | 3.26E-04 |
| ENSMUSG00000014773.13 | DLL1 | -1.033854702 | 9.13E-06 | 8.78E-05 |
| ENSMUSG00000029754.13 | DLX6 | 1.016776913 | 7.37E-08 | 1.10E-06 |
| ENSMUSG00000060962.12 | DMKN | 3.39475482 | 3.11E-18 | 2.07E-16 |
| ENSMUSG00000042372.5 | DMRT3 | -2.343464811 | 1.93E-11 | 5.18E-10 |
| ENSMUSG00000056752.16 | DNAH9 | -2.021760327 | 1.15E-21 | 1.22E-19 |
| ENSMUSG00000000730.13 | DNMT3L | 2.161142362 | 2.01E-08 | 3.31E-07 |
| ENSMUSG00000052301.14 | DOC2A | 1.739800361 | 3.07E-23 | 3.91E-21 |
| ENSMUSG00000020848.7 | DOC2B | -1.433147007 | 1.42E-11 | 3.88E-10 |
| ENSMUSG00000038608.15 | DOCK10 | -1.429521318 | 1.10E-19 | 8.73E-18 |
| ENSMUSG00000035954.10 | DOCK4 | -1.509722625 | 6.67E-31 | 1.62E-28 |
| ENSMUSG00000025558.19 | DOCK9 | -1.185277883 | 7.12E-16 | 3.57E-14 |
| ENSMUSG00000027560.4 | DOK5 | 1.479437815 | 1.70E-11 | 4.61E-10 |
| ENSMUSG00000021221.15 | DPF3 | -1.706601065 | 2.40E-25 | 4.04E-23 |
| ENSMUSG00000042737.9 | DPM3 | 1.617057793 | 7.13E-09 | 1.26E-07 |
| ENSMUSG00000036815.17 | DPP10 | 1.861532968 | 9.63E-39 | 4.30E-36 |

| | | | | |
|---|---|---|---|---|
| ENSMUSG00000035000.8 | DPP4 | 1.914913513 | 1.02E-11 | 2.86E-10 |
| ENSMUSG00000031786.7 | DRC7 | -2.061813055 | 5.00E-04 | 0.002949557 |
| ENSMUSG00000021478.6 | DRD1 | 1.777853893 | 4.07E-05 | 3.30E-04 |
| ENSMUSG00000032259.8 | DRD2 | 2.801444983 | 6.24E-07 | 7.79E-06 |
| ENSMUSG00000025496.5 | DRD4 | 5.064627852 | 9.81E-12 | 2.76E-10 |
| ENSMUSG00000039358.3 | DRD5 | -3.017607601 | 7.86E-34 | 2.54E-31 |
| ENSMUSG00000059898.9 | DSC3 | 1.570189268 | 0.010136844 | 0.036953606 |
| ENSMUSG00000032087.11 | DSCAML1 | -1.051140552 | 2.88E-09 | 5.41E-08 |
| ENSMUSG00000044393.15 | DSG2 | -1.277110722 | 1.99E-04 | 0.001338879 |
| ENSMUSG00000054889.10 | DSP | -2.546672995 | 2.58E-15 | 1.21E-13 |
| ENSMUSG00000037474.13 | DTL | 2.921373925 | 7.05E-25 | 1.10E-22 |
| ENSMUSG00000018648.15 | DUSP14 | 1.302837033 | 2.38E-10 | 5.31E-09 |
| ENSMUSG00000042662.16 | DUSP15 | 1.058757394 | 2.55E-13 | 8.78E-12 |
| ENSMUSG00000027368.6 | DUSP2 | 1.499213812 | 2.72E-04 | 0.0017538 |
| ENSMUSG00000039661.14 | DUSP26 | 1.32176831 | 1.72E-23 | 2.22E-21 |
| ENSMUSG00000031383.8 | DUSP9 | 1.95729686 | 1.07E-06 | 1.27E-05 |
| ENSMUSG00000034467.7 | DYNLRB2 | -1.155608689 | 0.004998832 | 0.020509492 |
| ENSMUSG00000047671.8 | DYNLT4 | -1.421260591 | 0.001571441 | 0.00779771 |
| ENSMUSG00000033788.15 | DYSF | 1.173952369 | 2.53E-08 | 4.12E-07 |
| ENSMUSG00000086843.3 | E030013I19Rik | -1.620902653 | 5.90E-07 | 7.42E-06 |
| ENSMUSG00000087216.1 | E130111B04Rik | -5.276262735 | 1.30E-06 | 1.52E-05 |
| ENSMUSG00000087231.7 | E230016M11Rik | 1.172223127 | 6.36E-06 | 6.35E-05 |
| ENSMUSG00000044719.5 | E230025N22Rik | 1.690631874 | 4.44E-05 | 3.55E-04 |
| ENSMUSG00000027490.17 | E2F1 | 1.159294673 | 2.03E-13 | 7.10E-12 |
| ENSMUSG00000053117.11 | E330013P04Rik | -2.678398342 | 1.84E-15 | 8.92E-14 |
| ENSMUSG00000026247.13 | ECEL1 | 2.279804702 | 4.12E-05 | 3.33E-04 |
| ENSMUSG00000028601.18 | ECHDC2 | -1.244967998 | 3.75E-14 | 1.45E-12 |
| ENSMUSG00000043631.8 | ECM2 | -1.07896311 | 2.56E-07 | 3.46E-06 |
| ENSMUSG00000026051.8 | ECRG4 | -2.848141682 | 0.001198307 | 0.006185948 |
| ENSMUSG00000022441.17 | EFCAB6 | 2.163600659 | 4.45E-06 | 4.60E-05 |
| ENSMUSG00000040659.3 | EFHD2 | 1.53736282 | 1.71E-22 | 1.96E-20 |
| ENSMUSG00000048915.12 | EFNA5 | 2.45694659 | 1.03E-29 | 2.27E-27 |
| ENSMUSG00000063600.14 | Egfem1 | -1.125538883 | 7.83E-06 | 7.64E-05 |
| ENSMUSG00000000402.2 | EGFL6 | -1.258720034 | 3.33E-04 | 0.002086838 |
| ENSMUSG00000038418.7 | EGR1 | 1.452595488 | 0.007262174 | 0.02799429 |
| ENSMUSG00000024772.9 | EHD1 | -1.281780897 | 2.16E-34 | 7.59E-32 |
| ENSMUSG00000043969.4 | EMX2 | -1.578827942 | 4.97E-16 | 2.55E-14 |
| ENSMUSG00000087095.2 | Emx2os | -1.876903961 | 2.47E-21 | 2.45E-19 |
| ENSMUSG00000022425.16 | ENPP2 | -1.278652627 | 0.005159892 | 0.021050014 |
| ENSMUSG00000032446.14 | EOMES | -1.850480475 | 0.014405998 | 0.048930615 |
| ENSMUSG00000024376.7 | EPB41L4A | -1.090664673 | 7.48E-09 | 1.32E-07 |
| ENSMUSG00000026235.14 | EPHA4 | -1.356528487 | 2.06E-20 | 1.79E-18 |
| ENSMUSG00000055540.15 | EPHA6 | -1.421649574 | 1.09E-10 | 2.60E-09 |
| ENSMUSG00000028289.12 | EPHA7 | -1.129031802 | 1.25E-15 | 6.15E-14 |
| ENSMUSG00000028661.8 | EPHA8 | 1.825628307 | 8.93E-10 | 1.82E-08 |

| | | | | |
|---|---|---|---|---|
| ENSMUSG00000028664.14 | EPHB2 | -1.05058845 | 2.06E-11 | 5.51E-10 |
| ENSMUSG00000010080.15 | EPN3 | -1.366490956 | 1.84E-05 | 1.64E-04 |
| ENSMUSG00000044726.8 | ERICH5 | -3.230118783 | 5.36E-10 | 1.13E-08 |
| ENSMUSG00000021255.17 | ESRRB | 1.250537965 | 1.55E-05 | 1.41E-04 |
| ENSMUSG00000070644.13 | ETNK2 | 1.263980599 | 6.58E-04 | 0.003725804 |
| ENSMUSG00000003382.18 | ETV3 | 1.235141359 | 1.40E-04 | 9.84E-04 |
| ENSMUSG00000118505.1 | ETV3L | 4.196421284 | 4.38E-04 | 0.002637281 |
| ENSMUSG00000050248.11 | EVC2 | -1.008739037 | 1.29E-07 | 1.84E-06 |
| ENSMUSG00000034282.3 | EVPL | 1.191040627 | 2.31E-06 | 2.55E-05 |
| ENSMUSG00000034584.3 | EXPH5 | 1.326671195 | 1.35E-09 | 2.65E-08 |
| ENSMUSG00000025932.14 | EYA1 | -1.094384826 | 4.54E-11 | 1.15E-09 |
| ENSMUSG00000021675.4 | F2RL2 | 4.131416579 | 1.18E-20 | 1.07E-18 |
| ENSMUSG00000026579.8 | F5 | -1.761166342 | 0.005080062 | 0.020801425 |
| ENSMUSG00000052125.16 | F730043M19Rik | -1.145733823 | 6.37E-17 | 3.66E-15 |
| ENSMUSG00000026655.15 | FAM107B | -1.020876964 | 2.92E-14 | 1.17E-12 |
| ENSMUSG00000028995.14 | FAM126A | -1.011412051 | 2.26E-11 | 5.97E-10 |
| ENSMUSG00000026153.15 | FAM135A | 1.009714655 | 1.40E-08 | 2.39E-07 |
| ENSMUSG00000071719.2 | FAM155B | -1.453212289 | 5.59E-17 | 3.24E-15 |
| ENSMUSG00000015484.3 | FAM163A | -1.010645131 | 6.19E-05 | 4.80E-04 |
| ENSMUSG00000027495.4 | FAM210B | -1.249677174 | 6.60E-40 | 3.31E-37 |
| ENSMUSG00000045655.9 | FAM216B | -1.145545279 | 0.004151425 | 0.017614756 |
| ENSMUSG00000047115.15 | FAM221A | 1.407465314 | 9.31E-05 | 6.85E-04 |
| ENSMUSG00000041930.7 | FAM222A | 1.258221019 | 6.28E-14 | 2.34E-12 |
| ENSMUSG00000046546.3 | FAM43A | 1.033519921 | 1.18E-05 | 1.11E-04 |
| ENSMUSG00000042377.8 | FAM83G | 2.088810445 | 1.77E-04 | 0.00120655 |
| ENSMUSG00000046761.13 | FAM83H | 1.522470417 | 4.23E-11 | 1.07E-09 |
| ENSMUSG00000034023.16 | FANCD2 | 1.553608939 | 1.95E-07 | 2.70E-06 |
| ENSMUSG00000046743.6 | FAT4 | -1.55128626 | 7.69E-08 | 1.14E-06 |
| ENSMUSG00000024598.9 | FBN2 | -1.027453078 | 3.47E-04 | 0.002156396 |
| ENSMUSG00000047730.17 | FCGBP | -2.263149093 | 3.73E-11 | 9.54E-10 |
| ENSMUSG00000070504.9 | FCRL6 | 3.83421034 | 2.91E-06 | 3.15E-05 |
| ENSMUSG00000021732.14 | FGF10 | -3.678099862 | 9.49E-38 | 4.02E-35 |
| ENSMUSG00000022523.10 | FGF12 | 1.042146607 | 1.69E-23 | 2.20E-21 |
| ENSMUSG00000031137.17 | FGF13 | -1.659597977 | 2.39E-46 | 1.72E-43 |
| ENSMUSG00000031230.3 | FGF16 | -2.010101788 | 6.54E-05 | 5.03E-04 |
| ENSMUSG00000031074.14 | FGF3 | 3.070711843 | 3.87E-04 | 0.002368587 |
| ENSMUSG00000048373.8 | FGFBP1 | 1.321433706 | 0.002413474 | 0.011228455 |
| ENSMUSG00000028874.14 | FGR | 4.710669728 | 8.10E-24 | 1.10E-21 |
| ENSMUSG00000051435.11 | FHAD1 | 1.108949353 | 2.22E-07 | 3.05E-06 |
| ENSMUSG00000041842.15 | FHDC1 | 1.101546974 | 6.14E-06 | 6.15E-05 |
| ENSMUSG00000051000.17 | FHIP1A | -1.271758186 | 4.02E-08 | 6.21E-07 |
| ENSMUSG00000034295.10 | FHOD3 | 2.522284715 | 9.15E-63 | 1.16E-59 |
| ENSMUSG00000026841.7 | FIBCD1 | -3.552110523 | 8.63E-43 | 4.92E-40 |
| ENSMUSG00000047787.8 | FLRT1 | 1.312926568 | 7.72E-19 | 5.40E-17 |
| ENSMUSG00000040170.13 | FMO2 | 1.125347195 | 0.001398238 | 0.007085151 |

| | | | | |
|---|---|---|---|---|
| ENSMUSG00000039735.16 | Fnbp1l | 1.932142169 | 4.91E-73 | 2.03E-69 |
| ENSMUSG00000045326.13 | FNDC7 | 3.573728586 | 1.94E-12 | 5.93E-11 |
| ENSMUSG00000048721.2 | FNDC9 | 4.086732936 | 3.08E-07 | 4.10E-06 |
| ENSMUSG00000001827.12 | FOLR1 | -2.601321343 | 0.007557834 | 0.028951534 |
| ENSMUSG00000003545.3 | FOSB | 2.325514551 | 0.0043293 | 0.018275577 |
| ENSMUSG00000029563.16 | Foxp2 | 2.704438012 | 7.96E-16 | 3.99E-14 |
| ENSMUSG00000048285.9 | FRMD6 | 1.094625043 | 8.06E-05 | 6.02E-04 |
| ENSMUSG00000036131.12 | FRMD7 | -1.466644808 | 0.006359212 | 0.025067381 |
| ENSMUSG00000027004.3 | FRZB | -2.012766599 | 5.21E-17 | 3.03E-15 |
| ENSMUSG00000057092.12 | FXYD3 | -1.004126222 | 0.002599295 | 0.011913617 |
| ENSMUSG00000078612.9 | FYB2 | -1.286215711 | 5.83E-06 | 5.87E-05 |
| ENSMUSG00000086308.1 | G630016G05Rik | 1.723726604 | 9.29E-07 | 1.11E-05 |
| ENSMUSG00000031343.13 | GABRA3 | 1.178511003 | 1.48E-19 | 1.13E-17 |
| ENSMUSG00000055078.7 | GABRA5 | -2.310097675 | 2.97E-27 | 5.70E-25 |
| ENSMUSG00000055026.13 | GABRG3 | 1.045682456 | 7.76E-07 | 9.48E-06 |
| ENSMUSG00000026787.3 | GAD2 | 1.103172782 | 3.02E-14 | 1.20E-12 |
| ENSMUSG00000015312.9 | GADD45B | 1.679656489 | 0.001403694 | 0.007106845 |
| ENSMUSG00000056880.12 | GADL1 | 1.046703983 | 0.00218857 | 0.010317453 |
| ENSMUSG00000024907.7 | GAL | 2.980097691 | 6.20E-05 | 4.80E-04 |
| ENSMUSG00000047658.6 | GAL3ST3 | -1.138069398 | 4.79E-16 | 2.47E-14 |
| ENSMUSG00000021903.11 | GALNT15 | 1.344625869 | 2.80E-04 | 0.001802231 |
| ENSMUSG00000021130.8 | GALNT16 | -1.727062525 | 4.21E-38 | 1.83E-35 |
| ENSMUSG00000034040.16 | GALNT17 | -1.611202342 | 7.11E-40 | 3.45E-37 |
| ENSMUSG00000026994.9 | GALNT3 | -1.785414716 | 6.04E-14 | 2.26E-12 |
| ENSMUSG00000038860.15 | GARNL3 | 1.664662098 | 4.16E-52 | 4.04E-49 |
| ENSMUSG00000067724.5 | GBX1 | 2.787586605 | 1.01E-07 | 1.48E-06 |
| ENSMUSG00000037580.10 | GCH1 | 1.098062206 | 1.56E-04 | 0.001076434 |
| ENSMUSG00000041798.15 | GCK | -1.540666565 | 2.58E-06 | 2.81E-05 |
| ENSMUSG00000091387.2 | GCNT4 | 1.727173329 | 2.74E-11 | 7.10E-10 |
| ENSMUSG00000021943.7 | GDF10 | -1.402963974 | 1.50E-09 | 2.94E-08 |
| ENSMUSG00000022144.4 | GDNF | 1.739068356 | 5.41E-04 | 0.003157773 |
| ENSMUSG00000035314.10 | GDPD5 | 1.790410026 | 2.22E-16 | 1.19E-14 |
| ENSMUSG00000025089.15 | GFRA1 | -1.145958343 | 2.72E-14 | 1.09E-12 |
| ENSMUSG00000022103.10 | GFRA2 | 1.844617295 | 2.96E-20 | 2.53E-18 |
| ENSMUSG00000039131.15 | GIPC2 | 1.505126501 | 5.66E-09 | 1.02E-07 |
| ENSMUSG00000030406.7 | GIPR | 1.632957145 | 1.60E-04 | 0.001105045 |
| ENSMUSG00000047197.1 | GJD3 | 1.260686238 | 4.55E-06 | 4.69E-05 |
| ENSMUSG00000036395.15 | GLB1L2 | -1.626451097 | 6.25E-05 | 4.84E-04 |
| ENSMUSG00000032252.14 | GLCE | 1.311991334 | 3.29E-09 | 6.16E-08 |
| ENSMUSG00000024827.10 | GLDC | -1.302974643 | 8.78E-12 | 2.48E-10 |
| ENSMUSG00000024027.8 | GLP1R | -1.50971501 | 1.74E-04 | 0.001185014 |
| ENSMUSG00000000263.15 | GLRA1 | 3.744440905 | 1.40E-11 | 3.85E-10 |
| ENSMUSG00000038257.10 | GLRA3 | 1.79833146 | 1.64E-11 | 4.47E-10 |
| ENSMUSG00000022491.5 | Glycam1 | -1.825458439 | 0.012485006 | 0.043715741 |
| ENSMUSG00000062319.2 | Gm10115 | 2.345647244 | 2.65E-04 | 0.001712992 |

| | | | | |
|---|---|---|---|---|
| ENSMUSG00000062561.3 | Gm10118 | 1.159975029 | 1.69E-08 | 2.83E-07 |
| ENSMUSG00000066697.3 | Gm10172 | 2.11725742 | 4.43E-04 | 0.002662194 |
| ENSMUSG00000099907.1 | Gm10421 | 2.446960926 | 1.95E-08 | 3.22E-07 |
| ENSMUSG00000073787.4 | Gm10575 | 1.23492219 | 1.21E-07 | 1.74E-06 |
| ENSMUSG00000074235.4 | Gm10649 | -1.290422581 | 3.37E-12 | 1.01E-10 |
| ENSMUSG00000074776.4 | Gm10754 | -1.548057182 | 3.26E-04 | 0.002048242 |
| ENSMUSG00000096977.2 | Gm10827 | 1.783573563 | 6.75E-07 | 8.37E-06 |
| ENSMUSG00000087042.1 | Gm11611 | 1.210630502 | 3.15E-04 | 0.001985068 |
| ENSMUSG00000075437.4 | Gm11681 | -3.632784201 | 4.22E-21 | 4.00E-19 |
| ENSMUSG00000087675.1 | Gm11762 | 1.263938963 | 5.30E-07 | 6.74E-06 |
| ENSMUSG00000086774.1 | Gm11915 | 1.162621965 | 3.40E-05 | 2.83E-04 |
| ENSMUSG00000084898.1 | Gm12371 | 2.72287143 | 1.32E-23 | 1.74E-21 |
| ENSMUSG00000086320.1 | Gm12840 | 1.278872236 | 0.003751095 | 0.016203396 |
| ENSMUSG00000086606.1 | Gm13205 | 1.55113529 | 2.15E-15 | 1.02E-13 |
| ENSMUSG00000073716.6 | Gm13241 | 2.376185589 | 0.00371385 | 0.016059315 |
| ENSMUSG00000087496.1 | Gm13523 | 2.421362173 | 1.09E-14 | 4.64E-13 |
| ENSMUSG00000085397.1 | Gm13524 | 2.107861893 | 1.03E-07 | 1.50E-06 |
| ENSMUSG00000086196.7 | Gm13571 | 3.378274567 | 1.73E-04 | 0.001181472 |
| ENSMUSG00000087467.1 | Gm13601 | 4.35459576 | 7.44E-19 | 5.23E-17 |
| ENSMUSG00000087301.7 | Gm13629 | 1.058150449 | 5.19E-07 | 6.62E-06 |
| ENSMUSG00000086496.1 | Gm14204 | 1.026591431 | 9.23E-10 | 1.87E-08 |
| ENSMUSG00000087524.1 | Gm14285 | -1.039999712 | 3.51E-05 | 2.91E-04 |
| ENSMUSG00000086098.1 | Gm14291 | 2.290346554 | 4.85E-18 | 3.14E-16 |
| ENSMUSG00000091556.8 | Gm14569 | -2.89527589 | 5.26E-09 | 9.56E-08 |
| ENSMUSG00000085225.1 | Gm14637 | 5.7122997 | 4.45E-10 | 9.51E-09 |
| ENSMUSG00000087083.2 | Gm15083 | 1.3284139 | 0.01258498 | 0.044003463 |
| ENSMUSG00000084932.1 | Gm15156 | 1.330093209 | 6.06E-04 | 0.00348107 |
| ENSMUSG00000086034.1 | Gm15201 | 1.375680607 | 2.34E-07 | 3.19E-06 |
| ENSMUSG00000087400.8 | Gm15270 | 1.778348943 | 3.87E-05 | 3.17E-04 |
| ENSMUSG00000086291.1 | Gm15513 | 1.476929565 | 2.09E-06 | 2.33E-05 |
| ENSMUSG00000085527.1 | Gm15535 | 1.356526202 | 1.67E-07 | 2.34E-06 |
| ENSMUSG00000086873.1 | Gm15672 | 1.050925578 | 7.59E-06 | 7.43E-05 |
| ENSMUSG00000085565.1 | Gm15721 | 2.41532599 | 5.33E-11 | 1.33E-09 |
| ENSMUSG00000087556.1 | Gm15764 | 2.078131813 | 3.88E-05 | 3.17E-04 |
| ENSMUSG00000087652.1 | Gm15918 | 1.517010905 | 3.05E-04 | 0.001938191 |
| ENSMUSG00000085055.2 | Gm15958 | -2.810339397 | 3.35E-10 | 7.27E-09 |
| ENSMUSG00000107132.1 | Gm15997 | 1.481665 | 0.008059318 | 0.030504603 |
| ENSMUSG00000087411.1 | Gm16048 | 1.960915387 | 5.63E-07 | 7.11E-06 |
| ENSMUSG00000087435.1 | Gm16323 | 1.087864963 | 5.09E-05 | 4.04E-04 |
| ENSMUSG00000090050.1 | Gm16339 | 1.211725554 | 2.12E-05 | 1.85E-04 |
| ENSMUSG00000066477.13 | Gm16551 | -4.087296582 | 5.25E-10 | 1.11E-08 |
| ENSMUSG00000087088.2 | Gm16638 | -1.049685116 | 9.13E-10 | 1.85E-08 |
| ENSMUSG00000102548.1 | Gm16701 | 1.774437568 | 9.91E-09 | 1.73E-07 |
| ENSMUSG00000086534.2 | Gm16758 | -2.058320004 | 1.57E-07 | 2.20E-06 |
| ENSMUSG00000091542.1 | Gm17167 (include: | 1.03421421 | 0.005320081 | 0.021641607 |

| | | | | |
|---|---|---|---|---|
| ENSMUSG00000097305.2 | Gm17276 | 1.25267399 | 1.73E-09 | 3.36E-08 |
| ENSMUSG00000091191.2 | Gm17334 | -1.160419424 | 2.28E-05 | 1.97E-04 |
| ENSMUSG00000091604.2 | Gm17349 | 1.061062417 | 3.78E-04 | 0.002318486 |
| ENSMUSG00000079669.2 | Gm17396 | 1.268124923 | 2.64E-10 | 5.87E-09 |
| ENSMUSG00000097183.2 | Gm17501 | 2.217015735 | 8.35E-06 | 8.10E-05 |
| ENSMUSG00000097482.2 | Gm17634 | 1.351226182 | 9.85E-04 | 0.005226841 |
| ENSMUSG00000097486.2 | Gm17733 | 1.331921208 | 7.82E-05 | 5.86E-04 |
| ENSMUSG00000111092.1 | Gm17875 | 1.595376673 | 0.010757573 | 0.038735168 |
| ENSMUSG00000117246.1 | Gm17891 | 3.815233489 | 1.49E-15 | 7.28E-14 |
| ENSMUSG00000115632.1 | Gm17922 | 1.749230295 | 4.80E-04 | 0.002844149 |
| ENSMUSG00000117767.1 | Gm18786 | 1.519579477 | 7.00E-06 | 6.91E-05 |
| ENSMUSG00000109372.1 | Gm19410 | 1.219881688 | 7.23E-07 | 8.89E-06 |
| ENSMUSG00000116903.1 | Gm19522 | -1.614321067 | 4.86E-09 | 8.87E-08 |
| ENSMUSG00000107317.1 | Gm19719 | 1.206659698 | 3.10E-04 | 0.001957424 |
| ENSMUSG00000108237.1 | Gm19757 | -1.046721716 | 0.003975317 | 0.017011759 |
| ENSMUSG00000113495.1 | Gm19792 | 1.779210383 | 0.001469118 | 0.007369691 |
| ENSMUSG00000105302.1 | Gm19817 | 2.313406052 | 3.41E-04 | 0.002124871 |
| ENSMUSG00000102331.1 | Gm19938 | 1.310572778 | 0.009258543 | 0.034227799 |
| ENSMUSG00000113429.1 | Gm20063 | 1.905535735 | 1.32E-08 | 2.26E-07 |
| ENSMUSG00000093677.1 | Gm20712 | 1.028363964 | 5.99E-11 | 1.47E-09 |
| ENSMUSG00000103428.1 | Gm20754 | -1.457011049 | 1.64E-11 | 4.46E-10 |
| ENSMUSG00000078747.9 | Gm20878 (include: | 1.104909888 | 5.54E-06 | 5.61E-05 |
| ENSMUSG00000118426.1 | Gm2102 | 1.887586493 | 1.09E-05 | 1.03E-04 |
| ENSMUSG00000097789.2 | Gm2115 | -3.332531551 | 2.82E-57 | 3.10E-54 |
| ENSMUSG00000092974.2 | Gm25072 | 1.445073634 | 3.54E-04 | 0.002196013 |
| ENSMUSG00000064585.1 | Gm25129 | 1.230723158 | 0.001289684 | 0.00658968 |
| ENSMUSG00000097296.1 | Gm26532 | 1.023816866 | 3.31E-04 | 0.002074324 |
| ENSMUSG00000097730.3 | Gm26588 | -1.0249224 | 4.40E-06 | 4.56E-05 |
| ENSMUSG00000096967.3 | Gm26621 | 1.070809288 | 0.002351436 | 0.010981793 |
| ENSMUSG00000097683.2 | Gm26644 | -4.255185089 | 3.55E-32 | 1.01E-29 |
| ENSMUSG00000097596.1 | Gm26673 | 2.021052815 | 2.02E-10 | 4.57E-09 |
| ENSMUSG00000097911.2 | Gm26691 | 1.068281662 | 1.42E-04 | 9.92E-04 |
| ENSMUSG00000097801.1 | Gm26777 | -1.104855786 | 1.68E-06 | 1.92E-05 |
| ENSMUSG00000097892.2 | Gm26801 | -1.263414076 | 1.73E-05 | 1.55E-04 |
| ENSMUSG00000097311.7 | Gm26871 | 1.049210589 | 0.004039833 | 0.017229802 |
| ENSMUSG00000097797.6 | Gm26901 | 1.012371972 | 0.002815771 | 0.012746614 |
| ENSMUSG00000097558.1 | Gm26902 | 1.014509815 | 4.44E-04 | 0.002668923 |
| ENSMUSG00000098088.1 | Gm26916 | 1.036972738 | 0.002398889 | 0.011171812 |
| ENSMUSG00000097248.9 | Gm2694 | 1.160367122 | 1.09E-09 | 2.19E-08 |
| ENSMUSG00000107008.3 | Gm2762 | -2.198213688 | 2.70E-15 | 1.26E-13 |
| ENSMUSG00000098374.1 | Gm28043 | 1.110044712 | 0.003165071 | 0.01405809 |
| ENSMUSG00000100213.1 | Gm28151 | 1.728836971 | 3.66E-20 | 3.07E-18 |
| ENSMUSG00000099644.1 | Gm28626 | 3.115161986 | 3.91E-09 | 7.22E-08 |
| ENSMUSG00000099760.1 | Gm28800 | 2.20408996 | 6.77E-20 | 5.53E-18 |
| ENSMUSG00000100980.1 | Gm29100 | 2.306168561 | 2.29E-04 | 0.001513686 |

| | | | | |
|---|---|---|---|---|
| ENSMUSG00000100393.1 | Gm29491 | -1.745034277 | 2.17E-09 | 4.14E-08 |
| ENSMUSG00000100596.1 | Gm29502 | 1.5523143 | 4.12E-05 | 3.33E-04 |
| ENSMUSG00000100417.2 | Gm29595 | 1.728132412 | 6.76E-09 | 1.21E-07 |
| ENSMUSG00000112466.1 | Gm29674 | 1.571459794 | 3.32E-08 | 5.23E-07 |
| ENSMUSG00000113689.1 | Gm29676 | 1.888835206 | 0.005338556 | 0.021693138 |
| ENSMUSG00000118219.1 | Gm29695 | 2.504277545 | 1.27E-17 | 7.80E-16 |
| ENSMUSG00000114708.1 | Gm30177 | -1.997025825 | 9.34E-05 | 6.87E-04 |
| ENSMUSG00000111066.1 | Gm30313 | 2.279803899 | 6.77E-04 | 0.003814782 |
| ENSMUSG00000118369.1 | Gm30541 | 2.071572532 | 1.20E-17 | 7.35E-16 |
| ENSMUSG00000106139.1 | Gm30648 | 2.205215871 | 1.51E-07 | 2.13E-06 |
| ENSMUSG00000110447.1 | Gm31518 | 1.99216371 | 3.11E-07 | 4.12E-06 |
| ENSMUSG00000117351.1 | Gm31615 | -2.016686953 | 0.001238712 | 0.006362701 |
| ENSMUSG00000111669.1 | Gm31698 | -2.645417843 | 1.24E-08 | 2.13E-07 |
| ENSMUSG00000109844.2 | Gm31816 | -1.160012558 | 4.36E-05 | 3.50E-04 |
| ENSMUSG00000103114.1 | Gm32200 | 2.327610516 | 1.23E-10 | 2.88E-09 |
| ENSMUSG00000114271.1 | Gm32224 | 2.756306743 | 1.81E-10 | 4.13E-09 |
| ENSMUSG00000103659.1 | Gm32569 | -1.379269983 | 0.010958223 | 0.039322922 |
| ENSMUSG00000092090.2 | Gm3294 | 1.535860807 | 1.96E-07 | 2.71E-06 |
| ENSMUSG00000109853.1 | Gm33045 | 2.11791248 | 5.91E-09 | 1.07E-07 |
| ENSMUSG00000117604.1 | Gm33228 | 1.233203409 | 1.95E-05 | 1.73E-04 |
| ENSMUSG00000110390.1 | Gm33320 | 1.609390444 | 1.18E-04 | 8.45E-04 |
| ENSMUSG00000113252.1 | Gm33489 | 2.201339971 | 8.59E-04 | 0.004650834 |
| ENSMUSG00000104237.1 | Gm33533 | 1.290846268 | 9.31E-05 | 6.85E-04 |
| ENSMUSG00000112805.1 | Gm34574 | 1.003006249 | 0.002762527 | 0.012550309 |
| ENSMUSG00000114743.1 | Gm34672 | 2.433718814 | 9.77E-18 | 6.11E-16 |
| ENSMUSG00000115912.1 | Gm35569 | 1.325697917 | 2.62E-04 | 0.00170056 |
| ENSMUSG00000115311.1 | Gm35823 | 2.035301119 | 3.61E-20 | 3.04E-18 |
| ENSMUSG00000116114.1 | Gm35853 | 5.784270713 | 1.18E-22 | 1.40E-20 |
| ENSMUSG00000110996.1 | Gm36251 | 3.066972221 | 3.05E-13 | 1.04E-11 |
| ENSMUSG00000106692.1 | Gm36447 | -2.201667012 | 1.12E-12 | 3.55E-11 |
| ENSMUSG00000112630.1 | Gm36908 | -1.729041723 | 2.15E-08 | 3.53E-07 |
| ENSMUSG00000104415.1 | Gm37069 | -1.962529975 | 1.36E-06 | 1.58E-05 |
| ENSMUSG00000103046.1 | Gm37309 | 1.19645184 | 0.00715459 | 0.027676481 |
| ENSMUSG00000103947.1 | Gm37345 | 1.035388129 | 0.001831369 | 0.008868983 |
| ENSMUSG00000091227.1 | Gm3755 | 2.572464317 | 8.39E-06 | 8.13E-05 |
| ENSMUSG00000102747.1 | Gm37602 | 1.656123767 | 0.003988326 | 0.017049223 |
| ENSMUSG00000103831.1 | Gm37608 | 1.51278877 | 0.002618147 | 0.011983392 |
| ENSMUSG00000103540.1 | Gm37668 | 3.812072432 | 0.010471684 | 0.037880923 |
| ENSMUSG00000102796.1 | Gm37711 | 1.946355415 | 3.60E-16 | 1.88E-14 |
| ENSMUSG00000102240.1 | Gm37717 | 5.898910478 | 1.61E-14 | 6.68E-13 |
| ENSMUSG00000102813.1 | Gm37795 | 2.256641749 | 1.78E-05 | 1.59E-04 |
| ENSMUSG00000102816.1 | Gm37814 | 1.845193385 | 0.002523171 | 0.011639007 |
| ENSMUSG00000103761.1 | Gm37859 | 1.286707287 | 0.009096714 | 0.033704989 |
| ENSMUSG00000102234.1 | Gm37885 | -1.82427809 | 2.15E-11 | 5.72E-10 |
| ENSMUSG00000103914.1 | Gm38073 | 4.375196281 | 1.24E-07 | 1.79E-06 |

| | | | | |
|---|---|---|---|---|
| ENSMUSG00000103082.1 | Gm38124 | 1.485349917 | 3.28E-05 | 2.74E-04 |
| ENSMUSG00000102465.1 | Gm38198 | 1.512727984 | 0.007578037 | 0.029015467 |
| ENSMUSG00000104277.1 | Gm38299 | 2.815777923 | 6.73E-12 | 1.93E-10 |
| ENSMUSG00000105218.4 | Gm38413 | 1.110545682 | 0.002903503 | 0.013079229 |
| ENSMUSG00000111360.1 | Gm38642 | 3.392898446 | 1.46E-11 | 3.98E-10 |
| ENSMUSG00000109450.1 | Gm39043 | 6.180038638 | 1.87E-11 | 5.03E-10 |
| ENSMUSG00000110179.1 | Gm39168 | 3.590248323 | 2.39E-13 | 8.24E-12 |
| ENSMUSG00000109868.1 | Gm39244 | 2.438002604 | 2.91E-17 | 1.72E-15 |
| ENSMUSG00000111389.1 | Gm39465 | 4.575486483 | 2.27E-04 | 0.001498137 |
| ENSMUSG00000110618.1 | Gm39822 | 1.526921258 | 5.85E-10 | 1.22E-08 |
| ENSMUSG00000113029.1 | Gm40578 | -1.0605842 | 4.41E-05 | 3.54E-04 |
| ENSMUSG00000113216.1 | Gm40841 | 1.004287601 | 4.18E-05 | 3.37E-04 |
| ENSMUSG00000097146.1 | Gm4211 | 1.088933661 | 1.73E-05 | 1.55E-04 |
| ENSMUSG00000105736.1 | Gm42456 | 2.152026279 | 1.15E-09 | 2.31E-08 |
| ENSMUSG00000106651.1 | Gm42608 | 1.20422113 | 1.97E-05 | 1.74E-04 |
| ENSMUSG00000104871.1 | Gm42639 | 3.524734821 | 2.84E-07 | 3.80E-06 |
| ENSMUSG00000105922.1 | Gm42769 | 1.699469236 | 0.002073598 | 0.009845714 |
| ENSMUSG00000105983.1 | Gm42770 | 1.73190497 | 0.00510302 | 0.020854063 |
| ENSMUSG00000105700.1 | Gm42772 | 1.605160118 | 0.001978421 | 0.009470042 |
| ENSMUSG00000105895.1 | Gm42829 | -1.017611021 | 3.13E-10 | 6.84E-09 |
| ENSMUSG00000107021.1 | Gm42853 | 1.376231397 | 0.003026597 | 0.013554735 |
| ENSMUSG00000105152.1 | Gm42864 | -2.525104482 | 5.29E-15 | 2.37E-13 |
| ENSMUSG00000104807.1 | Gm42865 | -2.026520938 | 2.61E-06 | 2.84E-05 |
| ENSMUSG00000104850.1 | Gm42901 | 1.383148021 | 5.37E-05 | 4.24E-04 |
| ENSMUSG00000105568.1 | Gm42971 | 2.559182824 | 2.07E-05 | 1.82E-04 |
| ENSMUSG00000107374.1 | Gm43172 | 2.418972907 | 3.05E-06 | 3.29E-05 |
| ENSMUSG00000106683.1 | Gm43263 | 1.381165565 | 9.44E-04 | 0.005050646 |
| ENSMUSG00000106408.1 | Gm43321 | 1.285099341 | 0.002462123 | 0.011405188 |
| ENSMUSG00000104951.1 | Gm43413 | 1.125727314 | 0.012069576 | 0.042543647 |
| ENSMUSG00000097532.2 | Gm4349 | 1.034822322 | 0.002840356 | 0.01284734 |
| ENSMUSG00000105613.1 | Gm43684 | 1.356184963 | 8.78E-05 | 6.52E-04 |
| ENSMUSG00000105366.1 | Gm43719 | 3.335759026 | 2.80E-22 | 3.13E-20 |
| ENSMUSG00000107690.1 | Gm44044 | 1.119176226 | 1.01E-06 | 1.20E-05 |
| ENSMUSG00000107826.1 | Gm44242 | 1.844557403 | 0.001134923 | 0.005897456 |
| ENSMUSG00000108033.1 | Gm44284 | 1.220019559 | 4.71E-04 | 0.002798465 |
| ENSMUSG00000108943.1 | Gm44559 | 1.858700105 | 0.001512211 | 0.007544437 |
| ENSMUSG00000108880.1 | Gm44560 | 1.630182426 | 4.12E-05 | 3.33E-04 |
| ENSMUSG00000109362.1 | Gm44562 | 2.336231418 | 5.97E-05 | 4.65E-04 |
| ENSMUSG00000109370.1 | Gm44675 | 1.791750936 | 0.002111861 | 0.010010126 |
| ENSMUSG00000108920.1 | Gm44678 | 2.250944696 | 2.63E-04 | 0.001705622 |
| ENSMUSG00000109366.1 | Gm44698 | 1.410338097 | 4.01E-07 | 5.23E-06 |
| ENSMUSG00000108802.1 | Gm44769 | 1.695847556 | 1.69E-04 | 0.001156783 |
| ENSMUSG00000109095.1 | Gm44799 | 1.817292409 | 0.003159153 | 0.014035581 |
| ENSMUSG00000109575.1 | Gm44831 | 1.364918183 | 7.46E-04 | 0.004132994 |
| ENSMUSG00000108871.1 | Gm45102 | -1.016519598 | 8.16E-04 | 0.004461817 |

| | | | | |
|---|---|---|---|---|
| ENSMUSG00000109130.1 | Gm45187 | -3.170410916 | 8.33E-19 | 5.81E-17 |
| ENSMUSG00000109031.1 | Gm45201 | 1.324304249 | 0.011571266 | 0.041067675 |
| ENSMUSG00000109604.1 | Gm45346 | 2.492937013 | 2.38E-07 | 3.23E-06 |
| ENSMUSG00000109674.1 | Gm45470 | 1.440589275 | 1.47E-06 | 1.70E-05 |
| ENSMUSG00000110296.1 | Gm45533 | 2.079273697 | 3.63E-08 | 5.64E-07 |
| ENSMUSG00000110086.1 | Gm45623 | 1.075077122 | 1.60E-05 | 1.45E-04 |
| ENSMUSG00000109992.1 | Gm45650 | -1.19561491 | 1.01E-05 | 9.63E-05 |
| ENSMUSG00000117713.1 | Gm46637 | 1.265115739 | 0.001076619 | 0.005645356 |
| ENSMUSG00000112812.1 | Gm47415 | 8.043477525 | 1.80E-21 | 1.83E-19 |
| ENSMUSG00000054412.5 | Gm4793 | 1.204677009 | 3.14E-05 | 2.63E-04 |
| ENSMUSG00000112666.1 | Gm48485 | 1.003923683 | 0.01232387 | 0.043292148 |
| ENSMUSG00000112003.1 | Gm4864 | -1.003841568 | 4.39E-06 | 4.55E-05 |
| ENSMUSG00000116597.1 | Gm536 | 1.348682175 | 0.010931014 | 0.039233812 |
| ENSMUSG00000081436.1 | Gm5912 | 4.105379216 | 2.40E-16 | 1.28E-14 |
| ENSMUSG00000074171.2 | Gm6658 | 1.359687946 | 1.19E-08 | 2.05E-07 |
| ENSMUSG00000073532.3 | Gm7276 | 1.734835108 | 1.91E-07 | 2.66E-06 |
| ENSMUSG00000106917.1 | Gm7832 | -3.200157992 | 2.01E-14 | 8.25E-13 |
| ENSMUSG00000114041.2 | Gm9269 | 1.697807294 | 6.53E-07 | 8.11E-06 |
| ENSMUSG00000029360.3 | Gm9754 | 1.335519267 | 0.001255088 | 0.006438564 |
| ENSMUSG00000108391.1 | Gm9768 | 1.265048297 | 1.20E-07 | 1.73E-06 |
| ENSMUSG00000053214.7 | Gm9899 | 1.281481967 | 8.33E-11 | 2.00E-09 |
| ENSMUSG00000054945.4 | Gm9958 | 1.118922518 | 5.92E-05 | 4.61E-04 |
| ENSMUSG00000113175.1 | Gm9973 | -1.253815224 | 4.02E-06 | 4.21E-05 |
| ENSMUSG00000068428.7 | GMNC | -2.982921883 | 2.88E-07 | 3.85E-06 |
| ENSMUSG00000024697.4 | GNA14 | -1.241531889 | 1.21E-04 | 8.63E-04 |
| ENSMUSG00000021556.12 | GOLM1 | -1.183202642 | 1.84E-29 | 4.00E-27 |
| ENSMUSG00000029314.14 | GPAT3 | 1.411899826 | 7.42E-12 | 2.11E-10 |
| ENSMUSG00000031119.4 | GPC4 | -1.54750962 | 1.46E-16 | 8.16E-15 |
| ENSMUSG00000041468.7 | GPR12 | -1.00510091 | 3.13E-05 | 2.62E-04 |
| ENSMUSG00000066197.5 | GPR139 | 3.200970315 | 1.48E-18 | 9.99E-17 |
| ENSMUSG00000045509.7 | GPR150 | -1.070662386 | 3.66E-04 | 0.002256886 |
| ENSMUSG00000042804.13 | GPR153 | 1.336914748 | 4.39E-11 | 1.11E-09 |
| ENSMUSG00000040836.15 | GPR161 | -2.47705632 | 5.81E-34 | 2.00E-31 |
| ENSMUSG00000053164.6 | GPR21 | 2.624470701 | 1.06E-17 | 6.56E-16 |
| ENSMUSG00000049649.8 | GPR3 | 2.234086837 | 4.96E-06 | 5.08E-05 |
| ENSMUSG00000118401.1 | GPR52 | 1.658142634 | 2.47E-06 | 2.72E-05 |
| ENSMUSG00000046922.7 | GPR6 | 2.423952108 | 5.29E-05 | 4.19E-04 |
| ENSMUSG00000040372.2 | GPR63 | -1.82044316 | 1.96E-14 | 8.08E-13 |
| ENSMUSG00000068696.6 | GPR88 | 4.311463003 | 8.75E-15 | 3.79E-13 |
| ENSMUSG00000051043.16 | GPRC5C | 2.060687439 | 2.89E-08 | 4.63E-07 |
| ENSMUSG00000074259.10 | GRAMD2A | 1.374961048 | 7.39E-11 | 1.79E-09 |
| ENSMUSG00000019312.10 | GRB7 | 1.560217832 | 1.78E-12 | 5.50E-11 |
| ENSMUSG00000036523.16 | GREB1 | 1.28748733 | 9.56E-04 | 0.005097268 |
| ENSMUSG00000074934.3 | GREM1 | -1.57635159 | 5.42E-07 | 6.87E-06 |
| ENSMUSG00000050069.3 | GREM2 | 1.391334291 | 3.20E-14 | 1.27E-12 |

| | | | | |
|---|---|---|---|---|
| ENSMUSG00000020656.16 | GRHL1 | -1.451385114 | 7.13E-19 | 5.05E-17 |
| ENSMUSG00000037188.7 | GRHL3 | 1.299885182 | 1.90E-05 | 1.68E-04 |
| ENSMUSG00000020524.16 | GRIA1 | -1.470326451 | 1.09E-33 | 3.40E-31 |
| ENSMUSG00000001985.9 | GRIK3 | 1.399349467 | 2.59E-14 | 1.05E-12 |
| ENSMUSG00000032017.14 | GRIK4 | -1.464527845 | 8.51E-23 | 1.02E-20 |
| ENSMUSG00000030098.15 | GRIP2 | 1.410554578 | 1.42E-19 | 1.10E-17 |
| ENSMUSG00000031450.8 | GRK1 | 1.063221753 | 5.81E-05 | 4.54E-04 |
| ENSMUSG00000063239.17 | GRM4 | 1.491922577 | 2.59E-14 | 1.05E-12 |
| ENSMUSG00000024211.15 | GRM8 | 1.515134672 | 1.23E-20 | 1.10E-18 |
| ENSMUSG00000024517.17 | GRP | -1.993812038 | 6.16E-07 | 7.70E-06 |
| ENSMUSG00000046182.8 | GSG1L | 1.680969984 | 1.27E-15 | 6.26E-14 |
| ENSMUSG00000004035.12 | GSTM2 | -1.0706966 | 1.27E-12 | 4.01E-11 |
| ENSMUSG00000042638.14 | GUCY2C | 1.313802732 | 0.001063945 | 0.005593755 |
| ENSMUSG00000030074.9 | GXYLT2 | 1.42824261 | 5.86E-06 | 5.89E-05 |
| ENSMUSG00000069270.6 | H2AC15 | -1.043531232 | 0.001076836 | 0.005645356 |
| ENSMUSG00000075297.10 | H60b/H60c | 3.07237577 | 1.31E-12 | 4.13E-11 |
| ENSMUSG00000020295.1 | Hbq1a | -1.134784002 | 3.31E-05 | 2.76E-04 |
| ENSMUSG00000032338.9 | HCN4 | 2.239123389 | 4.55E-15 | 2.05E-13 |
| ENSMUSG00000045471.4 | HCRT | 4.151727641 | 0.00120744 | 0.006227255 |
| ENSMUSG00000028778.15 | HCRTR1 | 1.173048458 | 6.33E-04 | 0.003606478 |
| ENSMUSG00000047171.8 | HELT | 5.209308199 | 9.20E-14 | 3.35E-12 |
| ENSMUSG00000029798.11 | HERC6 | 1.954905802 | 8.36E-23 | 1.01E-20 |
| ENSMUSG00000028946.7 | HES3 | 1.024093445 | 0.008876037 | 0.03306539 |
| ENSMUSG00000028864.7 | HGF | 1.374830544 | 6.05E-07 | 7.58E-06 |
| ENSMUSG00000021260.8 | HHIPL1 | 1.557048527 | 1.53E-16 | 8.47E-15 |
| ENSMUSG00000020080.8 | HKDC1 | 1.41390157 | 1.46E-13 | 5.20E-12 |
| ENSMUSG00000067235.14 | HLA-A | 1.038187664 | 0.014735126 | 0.04984348 |
| ENSMUSG00000055632.18 | HMCN2 | 3.458179879 | 2.72E-10 | 6.00E-09 |
| ENSMUSG00000015217.11 | Hmgb3 | 1.343993157 | 9.02E-25 | 1.37E-22 |
| ENSMUSG00000007617.18 | HOMER1 | 1.528634514 | 4.52E-04 | 0.002707623 |
| ENSMUSG00000025813.14 | HOMER2 | -1.673210494 | 9.56E-31 | 2.29E-28 |
| ENSMUSG00000003573.15 | HOMER3 | -2.663025859 | 8.64E-66 | 1.58E-62 |
| ENSMUSG00000052566.8 | HOOK2 | 1.274947019 | 2.20E-11 | 5.83E-10 |
| ENSMUSG00000059325.14 | HOPX | -1.463009049 | 3.20E-13 | 1.09E-11 |
| ENSMUSG00000087541.1 | Hopxos | -1.595010154 | 4.17E-08 | 6.43E-07 |
| ENSMUSG00000031722.10 | HP | 1.850154519 | 3.68E-04 | 0.002267376 |
| ENSMUSG00000028785.13 | HPCA | -1.367674891 | 5.59E-25 | 8.95E-23 |
| ENSMUSG00000071379.2 | HPCAL1 | 1.07567365 | 1.60E-09 | 3.11E-08 |
| ENSMUSG00000001249.14 | HPN | 1.096577573 | 4.87E-04 | 0.002879542 |
| ENSMUSG00000035273.14 | HPSE | 1.332782737 | 1.96E-06 | 2.20E-05 |
| ENSMUSG00000053004.9 | HRH1 | 1.131499073 | 2.69E-14 | 1.08E-12 |
| ENSMUSG00000034987.4 | HRH2 | 1.052583536 | 1.14E-07 | 1.65E-06 |
| ENSMUSG00000046607.6 | Hrk | -1.165940523 | 9.88E-09 | 1.72E-07 |
| ENSMUSG00000034773.16 | HROB | 1.16117372 | 2.21E-06 | 2.45E-05 |
| ENSMUSG00000046321.8 | HS3ST2 | 2.854865778 | 6.60E-23 | 8.07E-21 |

| | | | | |
|---|---|---|---|---|
| ENSMUSG00000078591.1 | HS3ST4 | -1.132144546 | 1.60E-05 | 1.45E-04 |
| ENSMUSG00000044499.11 | HS3ST5 | 1.8831979 | 5.73E-14 | 2.16E-12 |
| ENSMUSG00000016194.14 | HSD11B1 | -1.383156097 | 6.49E-11 | 1.59E-09 |
| ENSMUSG00000059970.7 | HSPA2 | -1.519430741 | 7.09E-32 | 1.98E-29 |
| ENSMUSG00000051456.4 | HSPB3 | 2.989361994 | 9.00E-14 | 3.28E-12 |
| ENSMUSG00000006221.7 | HSPB7 | 1.756267377 | 2.42E-05 | 2.08E-04 |
| ENSMUSG00000021721.5 | HTR1A | -1.413744453 | 8.97E-08 | 1.32E-06 |
| ENSMUSG00000070687.11 | HTR1D | 2.771880462 | 1.29E-10 | 3.01E-09 |
| ENSMUSG00000034997.5 | HTR2A | 1.319602227 | 2.32E-11 | 6.10E-10 |
| ENSMUSG00000026322.9 | HTR4 | -2.207659565 | 4.69E-14 | 1.79E-12 |
| ENSMUSG00000050534.8 | Htr5b | -3.163779517 | 2.88E-15 | 1.34E-13 |
| ENSMUSG00000037406.7 | HTRA4 | 1.951780492 | 1.63E-05 | 1.46E-04 |
| ENSMUSG00000031551.12 | IDO1 | 2.766753592 | 6.68E-08 | 1.00E-06 |
| ENSMUSG00000003541.6 | IER3 | 1.190907129 | 9.95E-05 | 7.28E-04 |
| ENSMUSG00000056708.5 | IER5 | 1.216433307 | 7.20E-08 | 1.08E-06 |
| ENSMUSG00000023046.6 | IGFBP6 | 1.829414435 | 1.85E-13 | 6.54E-12 |
| ENSMUSG00000051985.12 | IGFN1 | 3.913345177 | 4.89E-10 | 1.04E-08 |
| ENSMUSG00000076614.7 | IGHG1 | 4.601647396 | 2.42E-09 | 4.58E-08 |
| ENSMUSG00000031111.16 | IGSF1 | -1.261400996 | 7.56E-06 | 7.40E-05 |
| ENSMUSG00000037995.15 | IGSF9 | 2.072790409 | 5.04E-11 | 1.26E-09 |
| ENSMUSG00000034275.18 | IGSF9B | 2.242559883 | 7.20E-11 | 1.75E-09 |
| ENSMUSG00000027776.12 | IL12A | 2.459068609 | 3.59E-14 | 1.40E-12 |
| ENSMUSG00000001741.12 | IL16 | -3.686632753 | 2.58E-65 | 4.26E-62 |
| ENSMUSG00000015966.17 | IL17RB | 3.818289509 | 5.46E-11 | 1.35E-09 |
| ENSMUSG00000043088.16 | IL17RE | -1.261659271 | 0.005365378 | 0.021780692 |
| ENSMUSG00000026073.13 | IL1R2 | 1.761538299 | 1.21E-04 | 8.63E-04 |
| ENSMUSG00000059203.10 | IL1RAPL2 | 1.415104945 | 2.15E-09 | 4.10E-08 |
| ENSMUSG00000044244.18 | IL20RB | -3.748759987 | 3.09E-21 | 3.04E-19 |
| ENSMUSG00000074695.3 | IL22 | 5.183761655 | 2.06E-04 | 0.001382063 |
| ENSMUSG00000041324.14 | INHBA | 3.015603146 | 9.57E-06 | 9.15E-05 |
| ENSMUSG00000042106.5 | INKA1 | -1.407934541 | 5.55E-12 | 1.60E-10 |
| ENSMUSG00000068154.5 | INSM1 | -1.083267285 | 3.57E-04 | 0.002209116 |
| ENSMUSG00000064065.15 | IPCEF1 | 1.049891349 | 1.16E-13 | 4.18E-12 |
| ENSMUSG00000021676.10 | IQGAP2 | -2.830948741 | 1.29E-48 | 1.02E-45 |
| ENSMUSG00000046192.4 | IQUB | -1.172217426 | 2.98E-04 | 0.001901672 |
| ENSMUSG00000020227.10 | IRAK3 | 1.055978974 | 5.61E-06 | 5.67E-05 |
| ENSMUSG00000027009.18 | ITGA4 | -1.442371672 | 1.45E-10 | 3.36E-09 |
| ENSMUSG00000000555.8 | ITGA5 | 1.464012869 | 3.48E-04 | 0.00216184 |
| ENSMUSG00000026768.10 | ITGA8 | -1.642678461 | 1.03E-17 | 6.43E-16 |
| ENSMUSG00000032925.16 | ITGBL1 | -3.085633208 | 8.34E-23 | 1.01E-20 |
| ENSMUSG00000019762.4 | IYD | -4.332122422 | 8.28E-37 | 3.33E-34 |
| ENSMUSG00000048534.7 | JAML | 2.080677832 | 3.32E-16 | 1.75E-14 |
| ENSMUSG00000042686.5 | JPH1 | -1.34018803 | 2.64E-24 | 3.80E-22 |
| ENSMUSG00000025318.13 | JPH3 | -1.021992832 | 1.28E-12 | 4.05E-11 |
| ENSMUSG00000022208.12 | JPH4 | -1.333918792 | 2.33E-15 | 1.10E-13 |

| | | | | |
|---|---|---|---|---|
| ENSMUSG00000020216.13 | JSRP1 | 2.971465437 | 1.35E-11 | 3.72E-10 |
| ENSMUSG00000038292.14 | KASH5 | 1.465256543 | 6.27E-09 | 1.13E-07 |
| ENSMUSG00000025213.11 | KAZALD1 | 1.879542276 | 3.45E-18 | 2.28E-16 |
| ENSMUSG00000055675.6 | KBTBD11 | -1.069538987 | 1.24E-10 | 2.90E-09 |
| ENSMUSG00000045534.4 | KCNA5 | 2.039828997 | 4.29E-13 | 1.43E-11 |
| ENSMUSG00000018470.8 | KCNAB3 | 1.342397394 | 7.83E-13 | 2.53E-11 |
| ENSMUSG00000039672.12 | KCNE2 | -2.95368283 | 0.006000624 | 0.023901182 |
| ENSMUSG00000035165.14 | KCNE3 | 2.326084819 | 7.66E-04 | 0.00423519 |
| ENSMUSG00000051726.6 | KCNF1 | 1.28041977 | 2.58E-07 | 3.48E-06 |
| ENSMUSG00000074575.4 | KCNG1 | -1.307105301 | 2.05E-06 | 2.29E-05 |
| ENSMUSG00000059852.7 | KCNG2 | -3.350009902 | 4.62E-51 | 4.02E-48 |
| ENSMUSG00000058248.12 | KCNH1 | 1.023391024 | 2.30E-13 | 7.98E-12 |
| ENSMUSG00000035355.15 | KCNH4 | 2.192838669 | 2.58E-11 | 6.75E-10 |
| ENSMUSG00000001901.15 | KCNH6 | 1.012146656 | 1.32E-05 | 1.23E-04 |
| ENSMUSG00000059742.10 | KCNH7 | 1.180623177 | 1.90E-07 | 2.65E-06 |
| ENSMUSG00000053519.17 | KCNIP1 | 1.113232731 | 1.21E-19 | 9.54E-18 |
| ENSMUSG00000025221.16 | KCNIP2 | -1.066987321 | 6.76E-07 | 8.38E-06 |
| ENSMUSG00000079056.12 | KCNIP3 | 1.337618978 | 2.92E-20 | 2.51E-18 |
| ENSMUSG00000042529.14 | KCNJ12 | 1.062274314 | 1.16E-17 | 7.13E-16 |
| ENSMUSG00000079436.3 | KCNJ13 | -1.821042181 | 0.008090706 | 0.030595357 |
| ENSMUSG00000044216.7 | KCNJ4 | 1.069737441 | 3.36E-04 | 0.002101393 |
| ENSMUSG00000049265.7 | KCNK3 | 1.154336962 | 2.45E-13 | 8.46E-12 |
| ENSMUSG00000009545.14 | KCNQ1 | 1.774250457 | 1.56E-05 | 1.42E-04 |
| ENSMUSG00000101609.2 | KCNQ1OT1 | 1.455967207 | 4.85E-05 | 3.86E-04 |
| ENSMUSG00000058740.14 | KCNT1 | 2.049140058 | 1.37E-20 | 1.22E-18 |
| ENSMUSG00000022342.6 | KCNV1 | 1.167811983 | 3.82E-05 | 3.14E-04 |
| ENSMUSG00000098557.1 | KCTD12 | -1.251508109 | 4.16E-14 | 1.60E-12 |
| ENSMUSG00000046523.5 | KCTD4 | -1.050340831 | 7.14E-08 | 1.07E-06 |
| ENSMUSG00000028758.12 | KIF17 | 1.055059022 | 3.41E-15 | 1.57E-13 |
| ENSMUSG00000005672.12 | KIT | -1.519982146 | 3.52E-22 | 3.87E-20 |
| ENSMUSG00000058488.7 | KL | -2.548699992 | 2.39E-04 | 0.001571476 |
| ENSMUSG00000005148.8 | KLF5 | 2.15337044 | 9.55E-13 | 3.05E-11 |
| ENSMUSG00000047485.6 | KLHL34 | -1.145940071 | 3.63E-06 | 3.86E-05 |
| ENSMUSG00000025597.13 | KLHL4 | 1.147419421 | 2.69E-06 | 2.93E-05 |
| ENSMUSG00000074001.3 | KLHL40 | 2.100613116 | 7.69E-06 | 7.53E-05 |
| ENSMUSG00000030693.10 | KLK10 | -3.319823622 | 3.22E-11 | 8.29E-10 |
| ENSMUSG00000108444.1 | Klk2-ps | 2.140689577 | 3.04E-05 | 2.56E-04 |
| ENSMUSG00000064023.4 | KLK8 | -1.788262403 | 2.88E-11 | 7.45E-10 |
| ENSMUSG00000029414.11 | KNTC1 | -1.704610472 | 4.36E-09 | 8.00E-08 |
| ENSMUSG00000020912.4 | KRT12 | 3.829881177 | 8.91E-30 | 1.99E-27 |
| ENSMUSG00000023043.7 | KRT18 | -2.016926413 | 0.013973736 | 0.047846211 |
| ENSMUSG00000064201.8 | Krt2 | -2.695873021 | 8.61E-13 | 2.78E-11 |
| ENSMUSG00000061527.7 | KRT5 | -2.70625924 | 3.76E-06 | 3.97E-05 |
| ENSMUSG00000023039.17 | KRT7 | 3.779065538 | 3.89E-11 | 9.95E-10 |
| ENSMUSG00000063661.6 | KRT73 | -1.329687644 | 1.71E-06 | 1.94E-05 |

| | | | | |
|---|---|---|---|---|
| ENSMUSG00000022986.5 | KRT75 | 4.403762995 | 3.19E-06 | 3.43E-05 |
| ENSMUSG00000049382.10 | KRT8 | -2.473808287 | 0.004629439 | 0.019253806 |
| ENSMUSG00000037185.9 | KRT80 | 3.096358823 | 2.64E-17 | 1.57E-15 |
| ENSMUSG00000018334.18 | KSR1 | -1.440105961 | 1.21E-22 | 1.42E-20 |
| ENSMUSG00000024421.16 | LAMA3 | 1.016368344 | 1.59E-05 | 1.44E-04 |
| ENSMUSG00000019846.11 | LAMA4 | 1.099055692 | 1.81E-08 | 3.02E-07 |
| ENSMUSG00000002900.17 | LAMB1 | -1.213411075 | 2.59E-12 | 7.79E-11 |
| ENSMUSG00000026479.13 | LAMC2 | 3.0002852 | 3.13E-28 | 6.38E-26 |
| ENSMUSG00000027270.14 | LAMP5 | 1.19887089 | 0.008017717 | 0.030388932 |
| ENSMUSG00000025762.16 | Larp1b | 1.013224644 | 2.61E-19 | 1.91E-17 |
| ENSMUSG00000021959.16 | LATS2 | -1.186525566 | 6.50E-13 | 2.14E-11 |
| ENSMUSG00000016024.9 | LBP | -1.905926272 | 0.002791858 | 0.012655709 |
| ENSMUSG00000026354.8 | LCT | -3.466210884 | 9.24E-15 | 3.98E-13 |
| ENSMUSG00000038793.8 | LEFTY1 | -3.829962398 | 2.75E-22 | 3.09E-20 |
| ENSMUSG00000066652.5 | LEFTY2 | -2.198236798 | 1.86E-07 | 2.60E-06 |
| ENSMUSG00000042793.13 | LGR6 | -1.437766816 | 6.33E-06 | 6.32E-05 |
| ENSMUSG00000045312.12 | LHFPL2 | -1.461347843 | 1.66E-25 | 2.85E-23 |
| ENSMUSG00000018698.15 | LHX1 | -1.614509941 | 1.02E-05 | 9.68E-05 |
| ENSMUSG00000026934.15 | LHX3 | 4.270918434 | 3.74E-14 | 1.45E-12 |
| ENSMUSG00000096225.7 | LHX8 | 3.382811677 | 2.63E-04 | 0.001705622 |
| ENSMUSG00000019230.14 | LHX9 | -4.552925323 | 4.08E-37 | 1.68E-34 |
| ENSMUSG00000040699.13 | LIMD2 | -1.103548461 | 1.93E-13 | 6.80E-12 |
| ENSMUSG00000063804.8 | LIN28B | 1.932159589 | 1.96E-16 | 1.06E-14 |
| ENSMUSG00000053846.5 | LIPG | 2.59025357 | 2.89E-08 | 4.64E-07 |
| ENSMUSG00000044626.13 | LIPH | 1.03979976 | 0.001788591 | 0.008692403 |
| ENSMUSG00000020782.18 | LLGL2 | 1.277065048 | 2.32E-06 | 2.56E-05 |
| ENSMUSG00000036111.8 | LMO1 | -2.766479916 | 7.06E-34 | 2.33E-31 |
| ENSMUSG00000026686.14 | LMX1A | -2.464301934 | 2.92E-04 | 0.001868374 |
| ENSMUSG00000044471.12 | Lncpint | 1.01881559 | 5.49E-04 | 0.003202898 |
| ENSMUSG00000111938.1 | LOC105244649 | 1.482481796 | 1.46E-07 | 2.07E-06 |
| ENSMUSG00000097640.7 | LOC105244755 | -1.060249971 | 3.49E-07 | 4.60E-06 |
| ENSMUSG00000049929.7 | LPAR4 | -1.450030599 | 3.42E-08 | 5.37E-07 |
| ENSMUSG00000015568.16 | LPL | -1.505717722 | 3.26E-10 | 7.08E-09 |
| ENSMUSG00000072568.4 | LRATD2 | -1.53945115 | 7.18E-14 | 2.66E-12 |
| ENSMUSG00000073779.7 | Lrp8os2 | 3.260836832 | 2.17E-12 | 6.59E-11 |
| ENSMUSG00000090291.3 | LRRC10B | -1.116961827 | 0.003502031 | 0.015275368 |
| ENSMUSG00000030125.11 | LRRC23 | -1.070798647 | 6.81E-05 | 5.21E-04 |
| ENSMUSG00000028584.3 | LRRC38 | 1.13453953 | 4.37E-08 | 6.73E-07 |
| ENSMUSG00000045201.6 | LRRC3B | 1.01217194 | 1.47E-16 | 8.18E-15 |
| ENSMUSG00000022759.15 | Lrrc74b | 1.364108673 | 8.85E-06 | 8.53E-05 |
| ENSMUSG00000036273.15 | LRRK2 | 1.110556604 | 6.14E-10 | 1.28E-08 |
| ENSMUSG00000026443.3 | LRRN2 | -1.504697908 | 8.65E-18 | 5.47E-16 |
| ENSMUSG00000036295.5 | LRRN3 | -1.029103654 | 2.94E-07 | 3.92E-06 |
| ENSMUSG00000043110.2 | LRRN4 | -2.370874856 | 1.69E-12 | 5.25E-11 |
| ENSMUSG00000060780.2 | LRRTM1 | -1.192734064 | 8.51E-11 | 2.04E-09 |

| | | | | |
|---|---|---|---|---|
| ENSMUSG00000042846.8 | LRRTM3 | 1.317693629 | 6.07E-20 | 4.98E-18 |
| ENSMUSG00000071342.5 | LSMEM1 | 1.159004355 | 3.30E-04 | 0.002069002 |
| ENSMUSG00000018819.10 | LSP1 | 1.925780506 | 1.06E-16 | 6.00E-15 |
| ENSMUSG00000020377.15 | LTC4S | -1.069204242 | 4.46E-04 | 0.002675305 |
| ENSMUSG00000048706.3 | LURAP1L | -1.209470291 | 2.43E-17 | 1.46E-15 |
| ENSMUSG00000034634.7 | LY6D | 6.559980534 | 1.33E-19 | 1.04E-17 |
| ENSMUSG00000013766.11 | Ly6g6e | 2.356218669 | 4.08E-06 | 4.27E-05 |
| ENSMUSG00000032530.14 | LYZL4 | 3.673460302 | 2.84E-08 | 4.56E-07 |
| ENSMUSG00000044313.13 | MAB21L3 | 2.950808575 | 2.27E-05 | 1.96E-04 |
| ENSMUSG00000074622.4 | MAFB | 1.235257851 | 1.54E-12 | 4.80E-11 |
| ENSMUSG00000031224.6 | MAGEE2 | -1.625898507 | 3.95E-10 | 8.50E-09 |
| ENSMUSG00000033207.7 | MAMDC2 | -1.164267652 | 4.65E-12 | 1.36E-10 |
| ENSMUSG00000031925.17 | Maml2 | -1.094027458 | 9.39E-15 | 4.04E-13 |
| ENSMUSG00000031303.8 | MAP3K15 | -1.146521943 | 0.001868835 | 0.009031883 |
| ENSMUSG00000053137.7 | MAPK11 | 1.435372655 | 1.80E-12 | 5.56E-11 |
| ENSMUSG00000042688.16 | MAPK6 | 1.495137772 | 1.91E-07 | 2.65E-06 |
| ENSMUSG00000022269.13 | MARCHF11 | 1.636312049 | 5.63E-13 | 1.86E-11 |
| ENSMUSG00000047945.6 | MARCKSL1 | -1.049925124 | 2.37E-09 | 4.50E-08 |
| ENSMUSG00000022324.15 | MATN2 | -1.981872894 | 5.65E-23 | 6.96E-21 |
| ENSMUSG00000047259.3 | MC4R | -1.380991102 | 4.52E-04 | 0.002707623 |
| ENSMUSG00000013974.3 | MCEMP1 | 3.166454572 | 7.57E-14 | 2.79E-12 |
| ENSMUSG00000026669.14 | MCM10 | 1.002564957 | 4.89E-04 | 0.002888867 |
| ENSMUSG00000090667.2 | Mdfic2 | 4.071304906 | 6.21E-14 | 2.32E-12 |
| ENSMUSG00000043557.16 | MDGA1 | -2.09376209 | 3.42E-13 | 1.16E-11 |
| ENSMUSG00000029659.16 | MEDAG | 2.746467378 | 4.31E-19 | 3.10E-17 |
| ENSMUSG00000005583.16 | MEF2C | 2.249864638 | 1.59E-82 | 1.31E-78 |
| ENSMUSG00000068117.10 | MEI1 | 2.918861146 | 1.26E-18 | 8.58E-17 |
| ENSMUSG00000020160.18 | Meis1 | 1.080704561 | 4.05E-05 | 3.29E-04 |
| ENSMUSG00000027210.20 | MEIS2 | 1.13414438 | 1.07E-05 | 1.01E-04 |
| ENSMUSG00000022780.5 | MELTF | 1.386075619 | 6.83E-06 | 6.76E-05 |
| ENSMUSG00000039208.15 | METRNL | 1.002422312 | 5.58E-04 | 0.003240244 |
| ENSMUSG00000045555.3 | METTL24 | -1.457900371 | 3.83E-06 | 4.04E-05 |
| ENSMUSG00000042436.12 | MFAP4 | -1.08447623 | 5.05E-04 | 0.002971899 |
| ENSMUSG00000034739.17 | MFRP | -2.647975937 | 0.00581669 | 0.023269548 |
| ENSMUSG00000030877.11 | Mfsd13b | -1.740460095 | 7.07E-05 | 5.37E-04 |
| ENSMUSG00000100252.6 | Mir124-2hg | 1.316052836 | 2.51E-15 | 1.18E-13 |
| ENSMUSG00000075020.6 | Mir670hg | 2.286241551 | 1.17E-05 | 1.10E-04 |
| ENSMUSG00000097391.8 | Mirg | 1.026262256 | 8.51E-05 | 6.34E-04 |
| ENSMUSG00000097636.8 | Mirt1 | -1.061483854 | 1.24E-05 | 1.16E-04 |
| ENSMUSG00000074217.8 | Misp3 | 1.028928783 | 6.13E-04 | 0.003512086 |
| ENSMUSG00000058183.14 | MMEL1 | 1.213268801 | 5.27E-07 | 6.71E-06 |
| ENSMUSG00000074813.14 | Morrbid | 1.759601419 | 7.22E-07 | 8.88E-06 |
| ENSMUSG00000047502.14 | MROH7 | 1.279922645 | 4.45E-15 | 2.01E-13 |
| ENSMUSG00000051246.3 | MSANTD1 | -1.031962306 | 3.42E-04 | 0.002132203 |
| ENSMUSG00000063011.7 | MSLN | 1.395381019 | 8.17E-04 | 0.004464292 |

| | | | | |
|---|---|---|---|---|
| ENSMUSG00000048450.10 | MSX1 | -1.244335543 | 9.32E-05 | 6.85E-04 |
| ENSMUSG00000091510.4 | Mtag2 | 1.057111388 | 1.00E-05 | 9.55E-05 |
| ENSMUSG00000052105.17 | MTCL1 | 1.189611343 | 3.19E-09 | 5.98E-08 |
| ENSMUSG00000038065.13 | Mturn | -1.089821401 | 1.40E-19 | 1.09E-17 |
| ENSMUSG00000050808.13 | MUC15 | 2.523508133 | 1.80E-11 | 4.88E-10 |
| ENSMUSG00000042485.7 | MUSTN1 | 1.445742353 | 4.44E-05 | 3.55E-04 |
| ENSMUSG00000020061.18 | MYBPC1 | 4.836039859 | 1.95E-81 | 1.07E-77 |
| ENSMUSG00000002100.15 | MYBPC3 | 3.510521663 | 6.00E-34 | 2.02E-31 |
| ENSMUSG00000028654.13 | MYCL | -2.023214296 | 1.82E-31 | 4.82E-29 |
| ENSMUSG00000056328.14 | MYH1 | 2.338363365 | 2.61E-08 | 4.23E-07 |
| ENSMUSG00000033196.17 | MYH2 | 2.490036613 | 4.65E-13 | 1.55E-11 |
| ENSMUSG00000057003.12 | MYH4 | 1.291711548 | 0.001304144 | 0.006655324 |
| ENSMUSG00000085348.1 | Myhas | 4.483306541 | 4.62E-14 | 1.77E-12 |
| ENSMUSG00000061816.15 | MYL1 | 1.32647984 | 1.16E-08 | 2.01E-07 |
| ENSMUSG00000061086.12 | MYL4 | 4.251784195 | 2.03E-88 | 3.35E-84 |
| ENSMUSG00000031698.14 | MYLK3 | 1.840972327 | 1.07E-05 | 1.01E-04 |
| ENSMUSG00000042678.17 | MYO15A | 2.458520812 | 8.15E-12 | 2.31E-10 |
| ENSMUSG00000042064.13 | MYO3B | 1.236812707 | 7.63E-05 | 5.74E-04 |
| ENSMUSG00000025885.18 | MYO5B | -1.890005793 | 1.71E-12 | 5.28E-11 |
| ENSMUSG00000033590.8 | MYO5C | 2.139143016 | 7.45E-09 | 1.32E-07 |
| ENSMUSG00000026697.10 | MYOC | -2.642946225 | 2.51E-07 | 3.40E-06 |
| ENSMUSG00000049173.8 | MYOZ3 | 2.034920072 | 3.43E-05 | 2.85E-04 |
| ENSMUSG00000020067.8 | MYPN | 2.198076329 | 2.86E-10 | 6.28E-09 |
| ENSMUSG00000044968.16 | NAPEPLD | -1.123772967 | 2.93E-19 | 2.13E-17 |
| ENSMUSG00000043924.16 | NCMAP | 1.33389406 | 1.99E-07 | 2.75E-06 |
| ENSMUSG00000040536.15 | NECAB1 | 1.281386223 | 2.76E-19 | 2.02E-17 |
| ENSMUSG00000022656.15 | NECTIN3 | -1.033774746 | 7.31E-15 | 3.20E-13 |
| ENSMUSG00000022055.7 | NEFL | -1.060642993 | 1.34E-13 | 4.80E-12 |
| ENSMUSG00000037738.11 | NEK5 | -2.047390357 | 7.12E-05 | 5.39E-04 |
| ENSMUSG00000079434.8 | NEU2 | 1.465262593 | 2.64E-11 | 6.90E-10 |
| ENSMUSG00000034701.9 | NEUROD1 | -1.425696005 | 1.70E-24 | 2.49E-22 |
| ENSMUSG00000038255.6 | NEUROD2 | -1.064701984 | 1.72E-14 | 7.14E-13 |
| ENSMUSG00000037984.9 | NEUROD6 | -1.961373882 | 7.09E-25 | 1.10E-22 |
| ENSMUSG00000039103.12 | NEXN | 2.113595854 | 2.25E-12 | 6.84E-11 |
| ENSMUSG00000026259.14 | NGEF | 1.103972242 | 8.63E-12 | 2.44E-10 |
| ENSMUSG00000000120.6 | NGFR | 1.03285575 | 0.002170069 | 0.010253651 |
| ENSMUSG00000051251.3 | NHLH1 | -1.323866344 | 0.003296449 | 0.014555498 |
| ENSMUSG00000039835.16 | NHSL1 | -1.127315822 | 3.66E-16 | 1.91E-14 |
| ENSMUSG00000055761.14 | NKAIN3 | -1.328702025 | 1.32E-06 | 1.54E-05 |
| ENSMUSG00000021567.15 | NKD2 | -2.134632605 | 3.29E-18 | 2.18E-16 |
| ENSMUSG00000060621.6 | NKPD1 | 1.212034671 | 6.51E-05 | 5.01E-04 |
| ENSMUSG00000048528.7 | NKX1-2 | 8.07516075 | 2.47E-17 | 1.48E-15 |
| ENSMUSG00000049871.14 | NLRC3 | 1.020100156 | 2.31E-06 | 2.55E-05 |
| ENSMUSG00000049709.4 | NLRP10 | 2.112771431 | 2.45E-05 | 2.10E-04 |
| ENSMUSG00000038745.15 | NLRP6 | -1.183684728 | 0.001350309 | 0.006865459 |

| | | | | |
|---|---|---|---|---|
| ENSMUSG00000025723.12 | NMB | -1.012405662 | 0.00125543 | 0.006438564 |
| ENSMUSG00000055994.15 | NOD2 | 1.787439198 | 1.41E-06 | 1.64E-05 |
| ENSMUSG00000033774.4 | NPBWR1 | -1.187482416 | 0.001461984 | 0.007340592 |
| ENSMUSG00000020090.7 | NPFFR1 | -1.352132227 | 2.83E-08 | 4.56E-07 |
| ENSMUSG00000041616.9 | NPPA | 2.690554641 | 4.99E-07 | 6.40E-06 |
| ENSMUSG00000026241.5 | NPPC | 1.334133635 | 1.85E-06 | 2.09E-05 |
| ENSMUSG00000027931.12 | NPR1 | -1.064627127 | 9.68E-06 | 9.24E-05 |
| ENSMUSG00000059991.7 | NPTX2 | 4.045945429 | 1.70E-08 | 2.84E-07 |
| ENSMUSG00000071230.7 | NPW | 4.92961672 | 1.10E-10 | 2.60E-09 |
| ENSMUSG00000029819.6 | NPY | 2.314540131 | 9.18E-16 | 4.57E-14 |
| ENSMUSG00000028004.12 | NPY2R | -3.272391445 | 2.60E-19 | 1.91E-17 |
| ENSMUSG00000031618.13 | NR3C2 | -2.289858038 | 3.22E-63 | 4.43E-60 |
| ENSMUSG00000028341.9 | NR4A3 | -1.66218231 | 7.31E-04 | 0.004065678 |
| ENSMUSG00000049134.15 | NRAP | 3.254553251 | 2.73E-22 | 3.08E-20 |
| ENSMUSG00000040632.16 | NRL | -1.627181673 | 3.29E-08 | 5.20E-07 |
| ENSMUSG00000049107.13 | NTF3 | -2.788640003 | 4.26E-13 | 1.42E-11 |
| ENSMUSG00000028072.6 | NTRK1 | 2.419151245 | 2.48E-05 | 2.12E-04 |
| ENSMUSG00000027568.9 | NTSR1 | 2.268809968 | 6.92E-13 | 2.27E-11 |
| ENSMUSG00000061356.13 | NUGGC | 4.359216389 | 3.60E-16 | 1.88E-14 |
| ENSMUSG00000030717.9 | NUPR1 | 1.29308249 | 0.001924656 | 0.009255612 |
| ENSMUSG00000090061.9 | NWD2 | 1.141298602 | 2.60E-13 | 8.93E-12 |
| ENSMUSG00000031410.14 | Nxf7 | 6.089350521 | 1.08E-23 | 1.45E-21 |
| ENSMUSG00000069132.3 | NXPH2 | 1.778668912 | 3.92E-04 | 0.002396709 |
| ENSMUSG00000040258.6 | NXPH4 | 1.798356607 | 5.83E-10 | 1.22E-08 |
| ENSMUSG00000053765.4 | Oas1f | 7.130145188 | 7.83E-08 | 1.16E-06 |
| ENSMUSG00000061462.18 | OBSCN | 1.722539229 | 4.18E-06 | 4.36E-05 |
| ENSMUSG00000032172.8 | OLFM2 | 1.775942897 | 1.28E-27 | 2.52E-25 |
| ENSMUSG00000022026.7 | OLFM4 | -1.197792408 | 1.24E-04 | 8.81E-04 |
| ENSMUSG00000038463.8 | OLFML2B | -3.451155768 | 1.73E-68 | 4.77E-65 |
| ENSMUSG00000059864.2 | Olfr10 (includes ot | 3.832088562 | 9.53E-07 | 1.14E-05 |
| ENSMUSG00000073975.3 | Olfr550 | -4.526027236 | 7.30E-20 | 5.94E-18 |
| ENSMUSG00000026525.9 | OPN3 | 1.935341831 | 1.81E-15 | 8.78E-14 |
| ENSMUSG00000050511.1 | OPRD1 | 1.942998121 | 2.15E-14 | 8.73E-13 |
| ENSMUSG00000025905.14 | OPRK1 | 1.781330296 | 5.60E-05 | 4.39E-04 |
| ENSMUSG00000059729.2 | OR2Y1 | 1.223812912 | 1.92E-06 | 2.16E-05 |
| ENSMUSG00000070417.6 | OR6A2 | 3.57463952 | 1.17E-09 | 2.34E-08 |
| ENSMUSG00000039747.11 | ORAI2 | -1.278340777 | 8.21E-33 | 2.42E-30 |
| ENSMUSG00000040875.12 | OSBPL10 | 1.034229981 | 7.59E-17 | 4.32E-15 |
| ENSMUSG00000029822.15 | OSBPL3 | 1.105380869 | 5.05E-10 | 1.07E-08 |
| ENSMUSG00000034990.15 | OTOA | 1.358018101 | 6.72E-04 | 0.003793216 |
| ENSMUSG00000050201.9 | OTOP2 | 3.672861069 | 2.52E-16 | 1.34E-14 |
| ENSMUSG00000018862.11 | OTOP3 | 2.53690259 | 1.10E-04 | 7.94E-04 |
| ENSMUSG00000044055.5 | OTOS | -2.343750504 | 4.42E-05 | 3.54E-04 |
| ENSMUSG00000021848.16 | OTX2 | -2.268273668 | 6.05E-04 | 0.003474426 |
| ENSMUSG00000037279.13 | OVOL2 | 3.145331786 | 2.61E-11 | 6.81E-10 |

| | | | | |
|---|---|---|---|---|
| ENSMUSG00000027257.13 | PACSIN3 | 1.096972143 | 1.28E-11 | 3.53E-10 |
| ENSMUSG00000028927.6 | PADI2 | -1.071985544 | 3.17E-08 | 5.04E-07 |
| ENSMUSG00000027188.8 | PAMR1 | 1.02182702 | 1.89E-09 | 3.63E-08 |
| ENSMUSG00000021223.13 | PAPLN | 1.683875768 | 8.80E-08 | 1.29E-06 |
| ENSMUSG00000028370.7 | PAPPA | 2.620722771 | 1.71E-04 | 0.001171791 |
| ENSMUSG00000073530.11 | PAPPA2 | 2.086476254 | 7.53E-05 | 5.67E-04 |
| ENSMUSG00000022439.9 | PARVG | 1.056112173 | 3.14E-10 | 6.84E-09 |
| ENSMUSG00000035873.8 | PAWR | 1.077745569 | 0.002481723 | 0.011479882 |
| ENSMUSG00000086029.7 | Pax6os1 | -1.621269974 | 5.71E-05 | 4.47E-04 |
| ENSMUSG00000038718.15 | PBX3 | 1.059142042 | 1.75E-04 | 0.001191473 |
| ENSMUSG00000034755.18 | PCDH11X | 1.035296502 | 7.28E-07 | 8.93E-06 |
| ENSMUSG00000052613.16 | PCDH15 | 1.749357934 | 1.93E-17 | 1.17E-15 |
| ENSMUSG00000051323.16 | PCDH19 | -1.229637366 | 3.19E-10 | 6.94E-09 |
| ENSMUSG00000050505.7 | PCDH20 | -3.04772991 | 5.80E-51 | 4.79E-48 |
| ENSMUSG00000029108.14 | PCDH7 | 1.253266162 | 8.06E-15 | 3.51E-13 |
| ENSMUSG00000004630.18 | PCP2 | -1.05161109 | 0.007564012 | 0.028968483 |
| ENSMUSG00000044254.6 | PCSK9 | 5.252640623 | 3.30E-17 | 1.94E-15 |
| ENSMUSG00000023868.17 | PDE10A | 1.158640059 | 3.92E-06 | 4.12E-05 |
| ENSMUSG00000075270.11 | PDE11A | -2.009496519 | 1.47E-21 | 1.53E-19 |
| ENSMUSG00000110195.1 | PDE2A | 1.249719006 | 2.52E-09 | 4.77E-08 |
| ENSMUSG00000019990.15 | PDE7B | 1.090579833 | 4.19E-06 | 4.37E-05 |
| ENSMUSG00000055044.12 | PDLIM1 | 1.597866628 | 3.79E-07 | 4.96E-06 |
| ENSMUSG00000031636.7 | Pdlim3 | 1.798047079 | 4.12E-05 | 3.33E-04 |
| ENSMUSG00000027400.11 | PDYN | 1.691941313 | 1.86E-06 | 2.10E-05 |
| ENSMUSG00000074818.13 | PDZD7 | 1.096907181 | 2.14E-04 | 0.001426172 |
| ENSMUSG00000035357.17 | PDZRN3 | 1.80865013 | 6.32E-25 | 1.00E-22 |
| ENSMUSG00000092035.8 | PEG10 | 1.855450358 | 3.59E-12 | 1.07E-10 |
| ENSMUSG00000045573.9 | PENK | 2.288269191 | 3.59E-07 | 4.71E-06 |
| ENSMUSG00000031212.3 | Pgr15l | 1.6676856 | 0.006849976 | 0.026660502 |
| ENSMUSG00000086889.1 | Phf2os1 | 1.247624344 | 0.001851743 | 0.008959768 |
| ENSMUSG00000067780.3 | PI15 | -2.148141194 | 7.76E-10 | 1.59E-08 |
| ENSMUSG00000046207.14 | PIK3R6 | 1.972480797 | 5.84E-19 | 4.15E-17 |
| ENSMUSG00000024014.8 | PIM1 | 2.292462408 | 1.04E-04 | 7.59E-04 |
| ENSMUSG00000024867.14 | PIP5K1B | -1.635271563 | 8.53E-33 | 2.47E-30 |
| ENSMUSG00000046854.16 | PIP5KL1 | 1.939314842 | 9.05E-17 | 5.12E-15 |
| ENSMUSG00000048070.4 | PIRT | -1.74887986 | 8.45E-04 | 0.004588378 |
| ENSMUSG00000082286.11 | Pisd-ps1 | 1.148135076 | 3.91E-09 | 7.22E-08 |
| ENSMUSG00000040543.16 | PITPNM3 | 1.269486615 | 2.56E-19 | 1.90E-17 |
| ENSMUSG00000029423.10 | PIWIL1 | 1.818305837 | 3.23E-07 | 4.28E-06 |
| ENSMUSG00000048827.12 | PKD1L3 | -1.1960719 | 2.88E-05 | 2.44E-04 |
| ENSMUSG00000041957.15 | PKP2 | -2.973866824 | 2.37E-69 | 7.82E-66 |
| ENSMUSG00000021822.3 | PLAU | 1.606267895 | 1.87E-15 | 9.04E-14 |
| ENSMUSG00000046223.10 | PLAUR | 2.378011974 | 2.13E-04 | 0.001419531 |
| ENSMUSG00000029134.14 | PLB1 | 1.584031179 | 3.15E-05 | 2.64E-04 |
| ENSMUSG00000029055.17 | PLCH2 | -1.001240952 | 6.53E-10 | 1.36E-08 |

| | | | | |
|---|---|---|---|---|
| ENSMUSG00000064247.14 | Plcxd1 | 1.777242818 | 9.27E-18 | 5.82E-16 |
| ENSMUSG00000087141.1 | PLCXD2 | 1.208726801 | 9.70E-20 | 7.77E-18 |
| ENSMUSG00000055214.15 | PLD5 | 1.053537035 | 3.92E-06 | 4.12E-05 |
| ENSMUSG00000040428.19 | PLEKHA4 | 2.435216736 | 1.02E-18 | 7.08E-17 |
| ENSMUSG00000014782.16 | PLEKHG4 | 1.699684881 | 1.59E-05 | 1.44E-04 |
| ENSMUSG00000032068.14 | PLET1 | 2.957879658 | 1.45E-05 | 1.33E-04 |
| ENSMUSG00000035486.14 | PLK5 | -2.513171153 | 7.18E-12 | 2.05E-10 |
| ENSMUSG00000017417.14 | PLXDC1 | 1.523930536 | 5.33E-26 | 9.57E-24 |
| ENSMUSG00000030084.11 | PLXNA1 | -1.195164196 | 2.02E-09 | 3.87E-08 |
| ENSMUSG00000035383.7 | PMCH | 3.100808736 | 0.001209914 | 0.006238062 |
| ENSMUSG00000038400.15 | PMEPA1 | 1.4553758 | 5.51E-06 | 5.59E-05 |
| ENSMUSG00000046008.8 | PNLIP | 1.683535206 | 3.87E-08 | 6.00E-07 |
| ENSMUSG00000038216.7 | PNMT | 2.166283134 | 2.37E-04 | 0.001559419 |
| ENSMUSG00000019848.14 | POPDC3 | -2.707137974 | 4.73E-21 | 4.41E-19 |
| ENSMUSG00000090125.3 | Pou3f1 | -1.506427865 | 3.20E-12 | 9.58E-11 |
| ENSMUSG00000009734.18 | Pou6f2 | 1.148862472 | 0.001351753 | 0.006870684 |
| ENSMUSG00000000440.12 | PPARG | 1.930528433 | 3.28E-14 | 1.29E-12 |
| ENSMUSG00000039457.4 | PPL | -1.175574358 | 4.13E-05 | 3.34E-04 |
| ENSMUSG00000002228.7 | PPM1J | 4.964708137 | 1.92E-20 | 1.67E-18 |
| ENSMUSG00000040734.14 | PPP1R13L | 1.007034242 | 7.87E-04 | 0.004329742 |
| ENSMUSG00000061718.12 | PPP1R1B | 2.339456541 | 6.91E-24 | 9.50E-22 |
| ENSMUSG00000038976.12 | PPP1R9B | -1.066693288 | 2.29E-15 | 1.09E-13 |
| ENSMUSG00000038151.13 | PRDM1 | 1.183989127 | 1.98E-04 | 0.001332565 |
| ENSMUSG00000085069.2 | Prdm16os | -1.244433526 | 6.34E-05 | 4.89E-04 |
| ENSMUSG00000029913.14 | PRDM5 | -1.464115127 | 2.21E-14 | 8.96E-13 |
| ENSMUSG00000035456.9 | PRDM8 | -1.270522509 | 5.07E-12 | 1.47E-10 |
| ENSMUSG00000041577.5 | PRELP | -1.38811305 | 1.17E-10 | 2.75E-09 |
| ENSMUSG00000006014.16 | Prg4 | 1.486508547 | 0.003381007 | 0.014853406 |
| ENSMUSG00000041669.15 | PRIMA1 | 1.389979548 | 1.02E-07 | 1.48E-06 |
| ENSMUSG00000050965.14 | PRKCA | -1.304186556 | 1.43E-14 | 5.99E-13 |
| ENSMUSG00000021948.17 | PRKCD | -1.402746536 | 1.54E-06 | 1.77E-05 |
| ENSMUSG00000078816.9 | PRKCG | -1.311632493 | 6.81E-15 | 2.99E-13 |
| ENSMUSG00000029334.14 | PRKG2 | 1.68748598 | 1.45E-08 | 2.45E-07 |
| ENSMUSG00000010175.13 | PROX1 | -2.49274172 | 4.31E-30 | 9.88E-28 |
| ENSMUSG00000079045.3 | Prox1os | -2.187738889 | 1.53E-16 | 8.47E-15 |
| ENSMUSG00000093629.1 | Prox2os | 1.545450937 | 2.44E-06 | 2.68E-05 |
| ENSMUSG00000045725.5 | PRR15 | 1.234354296 | 1.31E-06 | 1.52E-05 |
| ENSMUSG00000037086.3 | PRR32 | -2.695800538 | 0.006477814 | 0.025482283 |
| ENSMUSG00000045027.7 | PRSS22 | 4.711384551 | 1.04E-34 | 3.73E-32 |
| ENSMUSG00000039405.7 | PRSS23 | 1.423870155 | 5.75E-11 | 1.42E-09 |
| ENSMUSG00000030623.4 | Prss23os | 1.166078765 | 0.00130592 | 0.006662331 |
| ENSMUSG00000024124.10 | Prss30 | 3.629217801 | 1.88E-11 | 5.04E-10 |
| ENSMUSG00000033491.13 | PRSS35 | -1.004928425 | 7.25E-05 | 5.49E-04 |
| ENSMUSG00000036480.9 | PRSS56 | 3.980718585 | 9.29E-14 | 3.37E-12 |
| ENSMUSG00000036030.9 | PRTG | -1.052258802 | 6.58E-04 | 0.003723743 |

| | | | | |
|---|---|---|---|---|
| ENSMUSG00000057729.12 | PRTN3 | 1.909152774 | 1.02E-13 | 3.71E-12 |
| ENSMUSG00000068744.12 | PSRC1 | -1.426971276 | 1.38E-06 | 1.60E-05 |
| ENSMUSG00000032322.14 | PSTPIP1 | 1.569916679 | 5.57E-13 | 1.84E-11 |
| ENSMUSG00000040016.16 | PTGER3 | 1.719817798 | 6.11E-10 | 1.27E-08 |
| ENSMUSG00000097487.7 | PTGES3L | 1.265798588 | 8.09E-06 | 7.88E-05 |
| ENSMUSG00000027864.9 | PTGFRN | 1.281562899 | 1.08E-13 | 3.88E-12 |
| ENSMUSG00000028378.6 | PTGR1 | 1.01859093 | 3.02E-04 | 0.001924272 |
| ENSMUSG00000047250.13 | PTGS1 | 1.477621988 | 2.14E-07 | 2.94E-06 |
| ENSMUSG00000032487.8 | PTGS2 | 1.893146949 | 0.004743767 | 0.019643572 |
| ENSMUSG00000097028.2 | Ptgs2os | 1.451649791 | 0.005490198 | 0.02220552 |
| ENSMUSG00000059456.13 | PTK2B | -1.365897818 | 4.37E-12 | 1.29E-10 |
| ENSMUSG00000027843.13 | PTPN22 | 1.081218414 | 2.22E-04 | 0.001476229 |
| ENSMUSG00000031506.11 | PTPN7 | 3.42512848 | 2.93E-08 | 4.69E-07 |
| ENSMUSG00000035429.13 | PTPRH | 1.757194068 | 1.09E-07 | 1.58E-06 |
| ENSMUSG00000028909.17 | PTPRU | -1.154746182 | 3.82E-10 | 8.23E-09 |
| ENSMUSG00000097993.7 | Ptprv | 3.695097958 | 5.44E-11 | 1.35E-09 |
| ENSMUSG00000060260.13 | PWWP2B | -1.440353698 | 4.85E-20 | 4.00E-18 |
| ENSMUSG00000034910.5 | PYGO1 | -1.226920168 | 4.32E-19 | 3.10E-17 |
| ENSMUSG00000058400.13 | QRFPR | 2.684421049 | 3.14E-08 | 5.00E-07 |
| ENSMUSG00000020175.15 | RAB36 | 1.022758656 | 4.41E-10 | 9.44E-09 |
| ENSMUSG00000030559.8 | RAB38 | -2.351841454 | 7.89E-06 | 7.70E-05 |
| ENSMUSG00000019066.13 | RAB3D | -1.004243389 | 7.14E-07 | 8.79E-06 |
| ENSMUSG00000023015.14 | RACGAP1 | 1.802330384 | 1.57E-24 | 2.34E-22 |
| ENSMUSG00000086022.1 | RAD51AP2 | 1.491962965 | 8.48E-06 | 8.21E-05 |
| ENSMUSG00000061311.6 | RAG1 | -1.501228552 | 6.63E-04 | 0.003750309 |
| ENSMUSG00000041046.7 | RAMP3 | 1.054135794 | 2.48E-04 | 0.001620494 |
| ENSMUSG00000017491.9 | RARB | 1.359277293 | 2.71E-07 | 3.65E-06 |
| ENSMUSG00000049404.7 | RARRES1 | -1.037742903 | 1.99E-05 | 1.75E-04 |
| ENSMUSG00000004952.13 | RASA4 | 1.797913722 | 1.52E-20 | 1.35E-18 |
| ENSMUSG00000049892.7 | RASD1 | -2.775844711 | 1.22E-14 | 5.17E-13 |
| ENSMUSG00000089809.9 | RASGEF1B | 2.012234521 | 3.19E-25 | 5.32E-23 |
| ENSMUSG00000020374.16 | RASGEF1C | 3.270146237 | 1.67E-54 | 1.73E-51 |
| ENSMUSG00000032356.12 | RASGRF1 | -1.29521969 | 8.19E-16 | 4.09E-14 |
| ENSMUSG00000032946.16 | RASGRP2 | 1.701038653 | 6.77E-18 | 4.31E-16 |
| ENSMUSG00000035275.14 | RAVER2 | -1.4869875 | 2.84E-27 | 5.52E-25 |
| ENSMUSG00000041534.9 | RBP3 | 2.091031246 | 6.06E-04 | 0.00348107 |
| ENSMUSG00000024990.13 | RBP4 | 1.163304123 | 6.99E-05 | 5.32E-04 |
| ENSMUSG00000091402.7 | RD3L | 2.70488824 | 6.06E-07 | 7.59E-06 |
| ENSMUSG00000074269.10 | REC114 | 2.260506548 | 2.96E-10 | 6.46E-09 |
| ENSMUSG00000020275.9 | REL | 1.190274924 | 1.33E-04 | 9.41E-04 |
| ENSMUSG00000040121.5 | REP15 | 1.242572273 | 2.10E-06 | 2.34E-05 |
| ENSMUSG00000030222.13 | RERG | -1.587522061 | 3.57E-14 | 1.39E-12 |
| ENSMUSG00000030110.13 | RET | 1.241960842 | 7.29E-08 | 1.09E-06 |
| ENSMUSG00000012705.16 | RETN | 3.55637067 | 1.11E-10 | 2.63E-09 |
| ENSMUSG00000051079.8 | RGS13 | 3.43965117 | 1.23E-09 | 2.44E-08 |

| | | | | |
|---|---|---|---|---|
| ENSMUSG00000052087.14 | RGS14 | -1.580218728 | 3.65E-07 | 4.79E-06 |
| ENSMUSG00000019775.17 | RGS17 | 1.048908805 | 1.17E-21 | 1.24E-19 |
| ENSMUSG00000026360.9 | RGS2 | 2.371411465 | 9.99E-11 | 2.38E-09 |
| ENSMUSG00000038530.11 | RGS4 | 1.467039661 | 1.09E-11 | 3.04E-10 |
| ENSMUSG00000021219.16 | RGS6 | 1.542849465 | 8.89E-25 | 1.36E-22 |
| ENSMUSG00000020599.13 | RGS9 | 2.967777545 | 1.27E-14 | 5.37E-13 |
| ENSMUSG00000032890.17 | RIMS3 | 1.393774518 | 1.52E-18 | 1.02E-16 |
| ENSMUSG00000044456.16 | RIN3 | 1.249490899 | 1.63E-08 | 2.74E-07 |
| ENSMUSG00000022221.14 | RIPK3 | 2.318227545 | 2.03E-10 | 4.58E-09 |
| ENSMUSG00000035896.5 | RNASE1 | -1.32410447 | 5.86E-05 | 4.58E-04 |
| ENSMUSG00000066800.11 | RNASEL | 1.074838197 | 2.24E-05 | 1.94E-04 |
| ENSMUSG00000031438.11 | RNF128 | -1.003602943 | 4.24E-13 | 1.42E-11 |
| ENSMUSG00000083695.6 | Rnf138rt1 | -1.007399046 | 0.008986848 | 0.033402837 |
| ENSMUSG00000047496.6 | RNF152 | 1.340647485 | 2.78E-10 | 6.14E-09 |
| ENSMUSG00000044164.2 | RNF182 | -1.880406048 | 1.11E-43 | 6.82E-41 |
| ENSMUSG00000058498.12 | RNF207 | 1.247924957 | 2.18E-04 | 0.001446524 |
| ENSMUSG00000036492.12 | RNF39 | 2.219898476 | 2.42E-10 | 5.38E-09 |
| ENSMUSG00000035305.5 | ROR1 | -2.139542618 | 5.42E-22 | 5.85E-20 |
| ENSMUSG00000110866.1 | RP23-212C9.3 | 1.058581957 | 3.96E-04 | 0.002415837 |
| ENSMUSG00000111325.1 | RP23-438P19.7 | 1.015054498 | 0.001016297 | 0.005362023 |
| ENSMUSG00000112255.1 | RP24-323H7.5 | 1.208423832 | 2.17E-05 | 1.89E-04 |
| ENSMUSG00000111000.1 | RP24-93F20.6 | 1.315413712 | 7.94E-04 | 0.004361588 |
| ENSMUSG00000111844.1 | RP24-94A19.4 | 2.702572725 | 3.58E-06 | 3.81E-05 |
| ENSMUSG00000028174.12 | RPE65 | 1.38462052 | 0.008464748 | 0.031748093 |
| ENSMUSG00000037593.12 | RSKR | 1.399997577 | 8.74E-05 | 6.50E-04 |
| ENSMUSG00000028871.7 | RSPO1 | 1.429707515 | 5.35E-07 | 6.80E-06 |
| ENSMUSG00000085925.3 | RTL1 | 3.211281707 | 7.44E-05 | 5.61E-04 |
| ENSMUSG00000047686.9 | RTL3 | -5.922158366 | 3.51E-20 | 2.97E-18 |
| ENSMUSG00000071679.4 | RTL4 | -1.422836664 | 2.91E-08 | 4.67E-07 |
| ENSMUSG00000033383.6 | RTP1 | -1.312326394 | 0.001933599 | 0.009290504 |
| ENSMUSG00000034009.14 | RXFP1 | 1.394663185 | 1.25E-06 | 1.47E-05 |
| ENSMUSG00000053368.13 | RXFP2 | 2.155819931 | 2.71E-07 | 3.65E-06 |
| ENSMUSG00000015843.10 | RXRG | 1.143968826 | 2.07E-04 | 0.001388283 |
| ENSMUSG00000030592.18 | RYR1 | -1.047993731 | 5.34E-04 | 0.003120398 |
| ENSMUSG00000041959.14 | S100A10 | 1.119144134 | 4.60E-04 | 0.002742132 |
| ENSMUSG00000047181.12 | SAMD14 | -1.173627346 | 4.47E-18 | 2.92E-16 |
| ENSMUSG00000051354.14 | SAMD3 | 2.067777284 | 1.02E-04 | 7.45E-04 |
| ENSMUSG00000087064.1 | Sap30bpos | -1.396026039 | 0.010606802 | 0.038320001 |
| ENSMUSG00000038331.15 | SATB2 | 2.085161853 | 3.04E-16 | 1.61E-14 |
| ENSMUSG00000046056.15 | SBSN | 1.912305979 | 9.47E-08 | 1.39E-06 |
| ENSMUSG00000012017.4 | SCARF2 | 1.191473397 | 5.96E-08 | 9.01E-07 |
| ENSMUSG00000038936.13 | SCCPDH | 1.52861658 | 2.60E-24 | 3.77E-22 |
| ENSMUSG00000050195.8 | Scd4 | 2.297777131 | 2.96E-04 | 0.001892145 |
| ENSMUSG00000022123.9 | SCEL | 1.008434634 | 1.37E-04 | 9.61E-04 |
| ENSMUSG00000050711.7 | SCG2 | 1.235713602 | 8.69E-07 | 1.05E-05 |

| | | | | |
|---|---|---|---|---|
| ENSMUSG00000034115.10 | SCN11A | 3.930423119 | 3.04E-07 | 4.04E-06 |
| ENSMUSG00000049281.16 | SCN3B | -1.598039153 | 1.25E-14 | 5.30E-13 |
| ENSMUSG00000046480.6 | SCN4B | 2.652455189 | 1.32E-12 | 4.14E-11 |
| ENSMUSG00000032511.17 | SCN5A | 1.546726499 | 6.89E-05 | 5.26E-04 |
| ENSMUSG00000034810.7 | SCN7A | 2.274248076 | 4.93E-09 | 8.98E-08 |
| ENSMUSG00000038580.13 | Sct | 3.986428245 | 8.11E-14 | 2.97E-12 |
| ENSMUSG00000034161.8 | SCX | 1.710242421 | 3.58E-08 | 5.58E-07 |
| ENSMUSG00000039683.16 | SDK1 | 1.235658294 | 6.52E-09 | 1.17E-07 |
| ENSMUSG00000001103.7 | SEBOX | -1.833070384 | 4.84E-10 | 1.03E-08 |
| ENSMUSG00000026589.14 | SEC16B | 1.708082483 | 3.15E-05 | 2.64E-04 |
| ENSMUSG00000039234.11 | SEC24D | -1.238645755 | 2.17E-19 | 1.61E-17 |
| ENSMUSG00000028883.17 | SEMA3A | 3.150389689 | 8.93E-34 | 2.83E-31 |
| ENSMUSG00000028064.17 | SEMA4A | 1.687564759 | 1.56E-19 | 1.19E-17 |
| ENSMUSG00000022231.10 | SEMA5A | -1.558435282 | 1.56E-16 | 8.56E-15 |
| ENSMUSG00000063232.16 | SERPINA11 | 3.28380901 | 9.44E-06 | 9.04E-05 |
| ENSMUSG00000058260.2 | SERPINA9 | 3.726789207 | 1.12E-09 | 2.24E-08 |
| ENSMUSG00000037411.10 | SERPINE1 | 2.6741953 | 9.35E-04 | 0.0050067 |
| ENSMUSG00000008384.8 | SERTAD1 | 1.854790048 | 2.49E-06 | 2.74E-05 |
| ENSMUSG00000058153.15 | SEZ6L | -1.032360641 | 2.90E-12 | 8.72E-11 |
| ENSMUSG00000027996.13 | SFRP2 | 2.160833756 | 1.26E-08 | 2.16E-07 |
| ENSMUSG00000018822.7 | SFRP5 | 1.282636586 | 0.002272277 | 0.010666395 |
| ENSMUSG00000112343.1 | Sfta3-ps | 2.090223121 | 8.32E-06 | 8.08E-05 |
| ENSMUSG00000039539.13 | SGCZ | 1.572846147 | 3.67E-10 | 7.93E-09 |
| ENSMUSG00000022436.16 | SH3BP1 | 1.005600554 | 3.65E-04 | 0.002253161 |
| ENSMUSG00000057719.11 | SH3RF2 | 3.737637194 | 1.55E-08 | 2.61E-07 |
| ENSMUSG00000022623.17 | SHANK3 | -1.143919462 | 6.40E-18 | 4.12E-16 |
| ENSMUSG00000044461.6 | SHISA2 | -1.333810868 | 7.90E-10 | 1.62E-08 |
| ENSMUSG00000053930.13 | SHISA6 | -2.788109867 | 1.88E-44 | 1.19E-41 |
| ENSMUSG00000096883.2 | SHISA8 | 1.23918442 | 1.47E-04 | 0.001023192 |
| ENSMUSG00000020534.14 | SHMT1 | -1.242290986 | 1.29E-10 | 3.01E-09 |
| ENSMUSG00000027833.16 | SHOX2 | -2.339269627 | 0.00818898 | 0.030889177 |
| ENSMUSG00000030583.16 | SIPA1L3 | -1.757621282 | 1.03E-24 | 1.55E-22 |
| ENSMUSG00000038805.10 | SIX3 | 2.379682356 | 0.001262854 | 0.00647061 |
| ENSMUSG00000054074.9 | SKIDA1 | -1.141973425 | 2.71E-13 | 9.27E-12 |
| ENSMUSG00000025161.16 | SLC16A3 | 1.862147285 | 3.10E-11 | 8.01E-10 |
| ENSMUSG00000025094.8 | SLC18A2 | -1.484907732 | 0.003399646 | 0.01490753 |
| ENSMUSG00000100241.1 | SLC18A3 | 3.17340749 | 3.58E-05 | 2.96E-04 |
| ENSMUSG00000063652.10 | Slc22a21 | 1.109024716 | 0.001442983 | 0.0072695 |
| ENSMUSG00000023828.3 | SLC22A3 | 1.216883209 | 1.25E-04 | 8.90E-04 |
| ENSMUSG00000026205.8 | SLC23A3 | 1.586359844 | 0.001962833 | 0.009409062 |
| ENSMUSG00000063873.10 | SLC24A3 | -1.339891817 | 2.05E-30 | 4.83E-28 |
| ENSMUSG00000020651.8 | SLC26A4 | -2.298567671 | 1.55E-23 | 2.04E-21 |
| ENSMUSG00000027932.14 | SLC27A3 | -1.173413877 | 1.12E-07 | 1.63E-06 |
| ENSMUSG00000005107.13 | SLC2A9 | -1.892130011 | 1.03E-20 | 9.31E-19 |
| ENSMUSG00000050473.5 | SLC35D3 | 2.701741274 | 5.95E-09 | 1.07E-07 |

| | | | | |
|---|---|---|---|---|
| ENSMUSG00000022464.14 | SLC38A4 | 1.554359899 | 1.95E-07 | 2.70E-06 |
| ENSMUSG00000034224.14 | SLC38A8 | 2.427775929 | 3.14E-10 | 6.84E-09 |
| ENSMUSG00000028360.10 | SLC44A5 | -2.223288753 | 5.42E-15 | 2.42E-13 |
| ENSMUSG00000074796.10 | SLC4A11 | 1.459065477 | 2.97E-04 | 0.001895145 |
| ENSMUSG00000011034.6 | SLC5A1 | -3.754290925 | 5.39E-16 | 2.74E-14 |
| ENSMUSG00000000792.2 | SLC5A5 | 1.353911696 | 5.96E-14 | 2.24E-12 |
| ENSMUSG00000023945.7 | SLC5A7 | 2.935017445 | 1.48E-05 | 1.35E-04 |
| ENSMUSG00000039728.15 | SLC6A5 | 6.157752009 | 8.70E-09 | 1.53E-07 |
| ENSMUSG00000026062.12 | SLC9A2 | -2.736747885 | 1.46E-19 | 1.12E-17 |
| ENSMUSG00000036123.8 | SLC9A3 | -1.37941846 | 1.62E-06 | 1.86E-05 |
| ENSMUSG00000026065.8 | SLC9A4 | -5.102851895 | 1.64E-63 | 2.47E-60 |
| ENSMUSG00000014786.8 | SLC9A5 | 1.445470377 | 8.94E-15 | 3.86E-13 |
| ENSMUSG00000032548.14 | SLCO2A1 | -2.741917892 | 1.83E-21 | 1.85E-19 |
| ENSMUSG00000025020.11 | SLIT1 | -1.641830036 | 4.53E-14 | 1.74E-12 |
| ENSMUSG00000078439.9 | SMIM24 | -1.041633195 | 9.15E-04 | 0.004910043 |
| ENSMUSG00000085007.3 | Smim43 | 3.39271861 | 1.80E-47 | 1.35E-44 |
| ENSMUSG00000001761.7 | SMO | -1.318173072 | 1.25E-19 | 9.75E-18 |
| ENSMUSG00000023886.10 | SMOC2 | -1.126908046 | 5.76E-08 | 8.72E-07 |
| ENSMUSG00000041476.12 | SMPX | -1.88879587 | 0.004760563 | 0.019708183 |
| ENSMUSG00000045667.14 | SMTNL2 | -1.407359422 | 2.88E-05 | 2.44E-04 |
| ENSMUSG00000006587.6 | SNAI3 | 7.471761014 | 1.11E-18 | 7.59E-17 |
| ENSMUSG00000044349.16 | Snhg11 | 1.234499814 | 2.44E-04 | 0.001598255 |
| ENSMUSG00000043531.16 | SORCS1 | 2.052671611 | 2.41E-21 | 2.41E-19 |
| ENSMUSG00000036169.6 | SOSTDC1 | -2.682807861 | 6.43E-04 | 0.00365216 |
| ENSMUSG00000044352.6 | SOWAHA | -1.654510044 | 1.22E-26 | 2.24E-24 |
| ENSMUSG00000045314.5 | SOWAHB | 2.416479251 | 1.48E-21 | 1.53E-19 |
| ENSMUSG00000053153.14 | SPAG16 | -1.460391647 | 7.82E-04 | 0.00430431 |
| ENSMUSG00000085244.8 | Spag17os | -4.517248338 | 6.11E-11 | 1.50E-09 |
| ENSMUSG00000029155.16 | SPATA18 | -1.131423664 | 0.006048388 | 0.024033467 |
| ENSMUSG00000072663.12 | SPEF2 | -1.918658179 | 4.54E-12 | 1.33E-10 |
| ENSMUSG00000061878.16 | SPHK1 | 2.06734702 | 9.61E-06 | 9.18E-05 |
| ENSMUSG00000050074.12 | SPINK8 | -3.010710488 | 2.09E-19 | 1.56E-17 |
| ENSMUSG00000051457.7 | SPN | -1.720359754 | 4.68E-06 | 4.81E-05 |
| ENSMUSG00000074899.6 | SPTBN5 | 1.47341823 | 7.19E-13 | 2.35E-11 |
| ENSMUSG00000071112.12 | SPX | 1.257943465 | 2.94E-06 | 3.19E-05 |
| ENSMUSG00000002007.5 | SRPK3 | 1.032540995 | 9.93E-04 | 0.005258614 |
| ENSMUSG00000031253.15 | SRPX2 | 1.564817065 | 3.80E-10 | 8.20E-09 |
| ENSMUSG00000004366.4 | SST | 1.044334515 | 5.91E-07 | 7.43E-06 |
| ENSMUSG00000047904.6 | SSTR2 | 1.707015542 | 8.92E-05 | 6.61E-04 |
| ENSMUSG00000037014.4 | SSTR4 | -1.464220409 | 5.61E-11 | 1.38E-09 |
| ENSMUSG00000050824.12 | SSTR5 | 2.043594713 | 3.22E-09 | 6.03E-08 |
| ENSMUSG00000033740.17 | ST18 | -2.245151093 | 3.36E-31 | 8.68E-29 |
| ENSMUSG00000024172.10 | ST6GAL2 | 1.380314554 | 2.26E-12 | 6.85E-11 |
| ENSMUSG00000039037.5 | ST6GALNAC5 | -1.535884629 | 5.17E-31 | 1.29E-28 |
| ENSMUSG00000030688.15 | STARD10 | 1.230851815 | 1.84E-31 | 4.82E-29 |

| | | | | |
|---|---|---|---|---|
| ENSMUSG00000016128.14 | STARD13 | -1.119022699 | 6.63E-16 | 3.35E-14 |
| ENSMUSG00000046027.17 | STARD5 | -1.282450508 | 1.82E-13 | 6.43E-12 |
| ENSMUSG00000014813.9 | STC1 | -1.176884932 | 1.04E-05 | 9.82E-05 |
| ENSMUSG00000020303.2 | STC2 | 2.042775641 | 1.12E-04 | 8.08E-04 |
| ENSMUSG00000015652.9 | STEAP1 | -2.135237905 | 0.014342837 | 0.048797989 |
| ENSMUSG00000039156.19 | STIM2 | -1.2173123 | 8.56E-36 | 3.29E-33 |
| ENSMUSG00000026094.14 | STK17B | 1.165635417 | 7.79E-13 | 2.53E-11 |
| ENSMUSG00000027744.4 | STOML3 | -1.683386701 | 0.00622806 | 0.024639873 |
| ENSMUSG00000032327.14 | STRA6 | -1.08336282 | 1.30E-04 | 9.24E-04 |
| ENSMUSG00000000739.14 | Sult5a1 | -2.116391651 | 1.85E-11 | 4.99E-10 |
| ENSMUSG00000006342.15 | SUSD2 | 1.742494358 | 1.98E-10 | 4.49E-09 |
| ENSMUSG00000021133.9 | SUSD6 | -1.174765654 | 5.11E-27 | 9.59E-25 |
| ENSMUSG00000051111.16 | SV2C | 1.409124491 | 1.36E-08 | 2.33E-07 |
| ENSMUSG00000003824.12 | SYCE2 | -1.137275248 | 7.77E-05 | 5.83E-04 |
| ENSMUSG00000085957.4 | Syna | -3.664855513 | 4.83E-16 | 2.49E-14 |
| ENSMUSG00000071234.3 | SYNDIG1L | 2.045204045 | 6.64E-06 | 6.60E-05 |
| ENSMUSG00000027887.11 | SYPL2 | -1.199168483 | 1.03E-07 | 1.50E-06 |
| ENSMUSG00000058420.8 | SYT17 | -1.135872719 | 1.56E-07 | 2.19E-06 |
| ENSMUSG00000026452.15 | SYT2 | 1.899007983 | 7.57E-14 | 2.79E-12 |
| ENSMUSG00000027849.19 | SYT6 | 2.368531098 | 8.72E-15 | 3.79E-13 |
| ENSMUSG00000028860.13 | SYTL1 | 3.269976257 | 3.90E-24 | 5.46E-22 |
| ENSMUSG00000031255.14 | SYTL4 | -1.49642701 | 6.89E-07 | 8.52E-06 |
| ENSMUSG00000061762.12 | TAC1 | 2.268186445 | 2.30E-07 | 3.15E-06 |
| ENSMUSG00000030043.11 | TACR1 | 1.77282154 | 1.42E-08 | 2.42E-07 |
| ENSMUSG00000055865.8 | TAFA3 | 3.311917934 | 4.01E-11 | 1.02E-09 |
| ENSMUSG00000035168.16 | TANC1 | -1.585595037 | 2.01E-19 | 1.51E-17 |
| ENSMUSG00000020096.21 | TBATA | -1.484623988 | 7.39E-13 | 2.41E-11 |
| ENSMUSG00000039813.14 | TBC1D2 | 1.100639211 | 1.50E-04 | 0.001041539 |
| ENSMUSG00000003134.10 | TBC1D8 | 1.210614371 | 1.14E-13 | 4.09E-12 |
| ENSMUSG00000042473.8 | TBC1D8B | -1.693043258 | 5.32E-25 | 8.62E-23 |
| ENSMUSG00000035033.15 | TBR1 | 1.459552172 | 9.42E-10 | 1.91E-08 |
| ENSMUSG00000027868.11 | TBX15 | 1.059999917 | 8.22E-04 | 0.004485673 |
| ENSMUSG00000021187.14 | TC2N | -1.775575457 | 0.01446064 | 0.049065751 |
| ENSMUSG00000007877.2 | TCAP | 1.444806216 | 5.82E-07 | 7.34E-06 |
| ENSMUSG00000038932.13 | TCFL5 | 2.307179061 | 1.85E-05 | 1.65E-04 |
| ENSMUSG00000028011.16 | TDO2 | -3.879466066 | 8.94E-19 | 6.20E-17 |
| ENSMUSG00000054003.13 | TDRD9 | 5.428965697 | 1.65E-21 | 1.69E-19 |
| ENSMUSG00000029217.16 | TEC | 1.19597337 | 6.52E-05 | 5.01E-04 |
| ENSMUSG00000028845.15 | TEKT2 | 1.411021587 | 2.63E-06 | 2.86E-05 |
| ENSMUSG00000052616.10 | TERB1 | 2.240978811 | 2.12E-15 | 1.01E-13 |
| ENSMUSG00000025927.13 | TFAP2B | 1.554859323 | 5.53E-07 | 7.00E-06 |
| ENSMUSG00000042596.7 | TFAP2D | 3.199500101 | 2.48E-05 | 2.12E-04 |
| ENSMUSG00000002603.15 | TGFB1 | 1.164034559 | 8.73E-07 | 1.05E-05 |
| ENSMUSG00000030782.17 | TGFB1I1 | 1.291048238 | 8.97E-07 | 1.08E-05 |
| ENSMUSG00000039239.14 | TGFB2 | -2.069103883 | 2.30E-51 | 2.11E-48 |

| | | | | |
|---|---|---|---|---|
| ENSMUSG00000027401.9 | TGM3 | 2.248911511 | 3.30E-22 | 3.66E-20 |
| ENSMUSG00000000214.11 | TH | 2.143034458 | 7.08E-06 | 6.98E-05 |
| ENSMUSG00000020317.14 | THEG | 1.684857775 | 6.27E-05 | 4.85E-04 |
| ENSMUSG00000029248.14 | THEGL | -2.02297792 | 4.20E-07 | 5.46E-06 |
| ENSMUSG00000037731.5 | THEMIS2 | 1.697242519 | 2.67E-09 | 5.03E-08 |
| ENSMUSG00000035686.8 | THRSP | 1.039561057 | 3.51E-07 | 4.61E-06 |
| ENSMUSG00000032289.15 | THSD4 | -2.219813378 | 5.02E-14 | 1.91E-12 |
| ENSMUSG00000042581.14 | THSD7B | -1.637691719 | 2.22E-13 | 7.73E-12 |
| ENSMUSG00000023800.15 | TIAM2 | 1.213666784 | 1.54E-09 | 3.01E-08 |
| ENSMUSG00000034917.8 | TJP3 | -1.990354979 | 2.06E-06 | 2.30E-05 |
| ENSMUSG00000034758.12 | TLE6 | 1.16504294 | 9.81E-08 | 1.44E-06 |
| ENSMUSG00000053626.5 | TLL1 | 1.819185269 | 2.49E-04 | 0.001623188 |
| ENSMUSG00000025013.15 | TLL2 | 1.431601958 | 0.004866729 | 0.020052184 |
| ENSMUSG00000027355.15 | TMCO5A | 1.003062485 | 0.00114093 | 0.005921218 |
| ENSMUSG00000022715.3 | TMEM114 | -1.728626068 | 4.23E-09 | 7.78E-08 |
| ENSMUSG00000034310.8 | TMEM132D | 1.279806795 | 2.18E-17 | 1.32E-15 |
| ENSMUSG00000056498.13 | TMEM154 | 4.313581217 | 8.39E-08 | 1.24E-06 |
| ENSMUSG00000070720.4 | TMEM200B | 1.08850398 | 2.49E-07 | 3.37E-06 |
| ENSMUSG00000049526.8 | TMEM202 | 1.131128727 | 2.68E-07 | 3.61E-06 |
| ENSMUSG00000045036.15 | TMEM232 | 2.662272851 | 2.24E-20 | 1.94E-18 |
| ENSMUSG00000059900.14 | TMEM40 | 4.620710801 | 2.90E-24 | 4.13E-22 |
| ENSMUSG00000023153.9 | TMEM52 | -1.11734533 | 3.20E-04 | 0.00201158 |
| ENSMUSG00000028786.15 | TMEM54 | -1.359409839 | 3.85E-14 | 1.49E-12 |
| ENSMUSG00000048108.13 | TMEM72 | -1.850232621 | 0.00705859 | 0.027375661 |
| ENSMUSG00000061702.10 | TMEM91 | 2.398675552 | 9.87E-21 | 9.00E-19 |
| ENSMUSG00000072845.3 | TMPRSS11A | -2.56661635 | 0.007057954 | 0.027375661 |
| ENSMUSG00000024034.13 | TMPRSS3 | 1.333212335 | 2.89E-04 | 0.001853876 |
| ENSMUSG00000033177.13 | TMPRSS7 | 1.329384958 | 1.97E-08 | 3.26E-07 |
| ENSMUSG00000074345.4 | TNFAIP8L3 | -1.278073797 | 1.18E-11 | 3.28E-10 |
| ENSMUSG00000063727.3 | TNFRSF11B | 1.475651713 | 2.38E-05 | 2.05E-04 |
| ENSMUSG00000023905.15 | TNFRSF12A | 1.551647804 | 8.41E-04 | 0.004576606 |
| ENSMUSG00000029075.6 | TNFRSF4 | 1.388988019 | 1.82E-04 | 0.001230654 |
| ENSMUSG00000028965.13 | TNFRSF9 | 1.43030065 | 6.28E-05 | 4.85E-04 |
| ENSMUSG00000026725.17 | TNN | 1.964437861 | 2.42E-04 | 0.001584725 |
| ENSMUSG00000091898.8 | TNNC1 | 4.174506717 | 1.65E-67 | 3.41E-64 |
| ENSMUSG00000035458.15 | TNNI3 | 2.652395123 | 3.19E-08 | 5.06E-07 |
| ENSMUSG00000026414.13 | TNNT2 | 3.271476004 | 8.22E-46 | 5.66E-43 |
| ENSMUSG00000033327.18 | TNXB | -1.68953326 | 1.61E-08 | 2.70E-07 |
| ENSMUSG00000022510.14 | TP63 | -1.421301899 | 8.80E-09 | 1.54E-07 |
| ENSMUSG00000029026.17 | TP73 | -1.999193237 | 9.26E-13 | 2.96E-11 |
| ENSMUSG00000035274.13 | TPBG | 1.901160502 | 1.40E-11 | 3.84E-10 |
| ENSMUSG00000028464.16 | Tpm2 | 1.297796698 | 1.74E-08 | 2.90E-07 |
| ENSMUSG00000021277.16 | TRAF3 | -1.105164443 | 1.43E-21 | 1.50E-19 |
| ENSMUSG00000062296.8 | TRANK1 | 1.146382101 | 2.23E-04 | 0.001479208 |
| ENSMUSG00000076498.2 | Trbc2 | 2.12987468 | 4.37E-07 | 5.65E-06 |

| | | | | |
|---|---|---|---|---|
| ENSMUSG00000038760.14 | TRHR | -1.009436627 | 0.007398987 | 0.028455257 |
| ENSMUSG00000039079.1 | Trhr2 | 3.824444616 | 1.45E-25 | 2.55E-23 |
| ENSMUSG00000032715.9 | TRIB3 | 1.451205898 | 4.97E-04 | 0.002931528 |
| ENSMUSG00000020773.11 | TRIM47 | 1.087157008 | 0.001726139 | 0.008428582 |
| ENSMUSG00000031026.15 | TRIM66 | 1.345772303 | 5.49E-11 | 1.36E-09 |
| ENSMUSG00000042828.12 | TRIM72 | 1.107029035 | 0.006380142 | 0.025139982 |
| ENSMUSG00000019487.11 | TRIP10 | 1.02989982 | 3.06E-04 | 0.001938946 |
| ENSMUSG00000056596.8 | TRNP1 | 1.001116513 | 3.30E-12 | 9.86E-11 |
| ENSMUSG00000027716.13 | TRPC3 | 1.971638801 | 1.16E-35 | 4.34E-33 |
| ENSMUSG00000027748.11 | TRPC4 | -1.446060734 | 3.44E-24 | 4.86E-22 |
| ENSMUSG00000041710.4 | TRPC5 | -1.815585727 | 9.21E-18 | 5.81E-16 |
| ENSMUSG00000021217.7 | TSHZ3 | 1.355667519 | 3.25E-21 | 3.18E-19 |
| ENSMUSG00000027217.13 | TSPAN18 | -2.514719397 | 2.19E-35 | 8.03E-33 |
| ENSMUSG00000061808.4 | TTR | -3.168513088 | 0.005385172 | 0.02183984 |
| ENSMUSG00000001473.7 | TUBB6 | 2.29670215 | 5.20E-04 | 0.003053683 |
| ENSMUSG00000035799.6 | TWIST1 | -2.831769215 | 1.36E-22 | 1.58E-20 |
| ENSMUSG00000027298.17 | TYRO3 | 1.186661738 | 3.17E-15 | 1.46E-13 |
| ENSMUSG00000010492.10 | Uckl1os | -1.168974804 | 0.004293453 | 0.018138165 |
| ENSMUSG00000026668.10 | UCMA | 2.599671325 | 1.41E-08 | 2.39E-07 |
| ENSMUSG00000054134.14 | UMODL1 | -1.520743557 | 7.86E-07 | 9.57E-06 |
| ENSMUSG00000018845.14 | UNC45B | 1.127077245 | 2.24E-06 | 2.48E-05 |
| ENSMUSG00000049436.5 | UPK1B | -1.289572157 | 0.002364764 | 0.011031557 |
| ENSMUSG00000022435.6 | UPK3A | 1.429763369 | 1.34E-06 | 1.56E-05 |
| ENSMUSG00000045288.10 | USH1G | 2.162414888 | 3.65E-06 | 3.87E-05 |
| ENSMUSG00000056900.13 | USP13 | 1.097728206 | 6.58E-14 | 2.44E-12 |
| ENSMUSG00000020905.11 | USP43 | 1.748347066 | 3.39E-08 | 5.33E-07 |
| ENSMUSG00000047712.8 | UST | -1.310913149 | 2.50E-21 | 2.48E-19 |
| ENSMUSG00000019820.11 | UTRN | 1.05686477 | 4.65E-21 | 4.36E-19 |
| ENSMUSG00000022479.15 | VDR | 2.885469226 | 1.20E-09 | 2.39E-08 |
| ENSMUSG00000037428.14 | VGF | 1.791213604 | 2.47E-10 | 5.51E-09 |
| ENSMUSG00000026175.12 | VIL1 | 1.30478698 | 4.15E-04 | 0.002518215 |
| ENSMUSG00000019772.5 | VIP | 1.161591891 | 1.55E-04 | 0.001075819 |
| ENSMUSG00000032528.5 | VIPR1 | 1.526502907 | 5.91E-11 | 1.45E-09 |
| ENSMUSG00000011171.11 | VIPR2 | -1.276740139 | 2.64E-04 | 0.00170644 |
| ENSMUSG00000024076.10 | VIT | -2.31761261 | 4.40E-15 | 2.00E-13 |
| ENSMUSG00000070601.5 | Vmn2r88 (includes | 1.821375272 | 7.75E-05 | 5.82E-04 |
| ENSMUSG00000050122.19 | VWA3B | -3.087900484 | 3.40E-29 | 7.30E-27 |
| ENSMUSG00000046613.19 | VWA5B2 | 1.0630866 | 1.17E-04 | 8.40E-04 |
| ENSMUSG00000007030.8 | VWA7 | 2.018656398 | 2.22E-08 | 3.63E-07 |
| ENSMUSG00000045648.15 | VWC2L | 1.42481952 | 2.41E-08 | 3.94E-07 |
| ENSMUSG00000044976.17 | WDR72 | -2.530557959 | 0.009883461 | 0.036173643 |
| ENSMUSG00000055235.11 | WDR86 | -1.737072295 | 0.003950556 | 0.016923343 |
| ENSMUSG00000000983.13 | Wfdc18 | 3.880855283 | 1.45E-21 | 1.52E-19 |
| ENSMUSG00000039474.13 | WFS1 | 1.291509753 | 3.29E-08 | 5.20E-07 |
| ENSMUSG00000086040.8 | WIPF3 | -1.600621157 | 1.50E-12 | 4.68E-11 |

| | | | | |
|---|---|---|---|---|
| ENSMUSG00000026167.14 | WNT10A | 2.302400527 | 2.96E-14 | 1.18E-12 |
| ENSMUSG00000022996.15 | WNT10B | 1.221730049 | 7.43E-04 | 0.004118479 |
| ENSMUSG00000015957.12 | WNT11 | -1.44383981 | 6.16E-05 | 4.78E-04 |
| ENSMUSG00000029671.15 | WNT16 | 1.568919738 | 8.84E-05 | 6.56E-04 |
| ENSMUSG00000036856.4 | WNT4 | 1.046430336 | 8.47E-06 | 8.21E-05 |
| ENSMUSG00000000126.11 | WNT9A | 1.067003758 | 5.36E-09 | 9.73E-08 |
| ENSMUSG00000018486.2 | WNT9B | 2.721387162 | 3.15E-04 | 0.001985508 |
| ENSMUSG00000079243.4 | XIRP1 | 3.148952794 | 4.34E-05 | 3.49E-04 |
| ENSMUSG00000042631.7 | XKR7 | 1.482525218 | 2.77E-08 | 4.46E-07 |
| ENSMUSG00000026117.10 | ZAP70 | -1.328392667 | 6.48E-09 | 1.16E-07 |
| ENSMUSG00000056586.8 | ZAR1L | 4.432838749 | 6.44E-18 | 4.12E-16 |
| ENSMUSG00000063659.11 | ZBTB18 | -1.428090151 | 1.06E-27 | 2.11E-25 |
| ENSMUSG00000022708.16 | ZBTB20 | -1.56276265 | 4.50E-12 | 1.32E-10 |
| ENSMUSG00000045064.5 | ZC2HC1C | 1.213030591 | 2.11E-09 | 4.04E-08 |
| ENSMUSG00000039981.6 | ZC3H12D | 1.610171366 | 6.00E-06 | 6.02E-05 |
| ENSMUSG00000048483.6 | ZDHHC22 | 2.329061889 | 1.24E-39 | 5.85E-37 |
| ENSMUSG00000036304.14 | ZDHHC23 | -1.030791591 | 2.73E-11 | 7.09E-10 |
| ENSMUSG00000026872.18 | ZEB2 | -1.09791432 | 4.39E-17 | 2.56E-15 |
| ENSMUSG00000074731.3 | Zfp345 (includes o | 3.162367792 | 0.014707046 | 0.049799498 |
| ENSMUSG00000053080.10 | ZFTA | -1.152927151 | 1.10E-17 | 6.83E-16 |
| ENSMUSG00000039634.12 | ZNF189 | -1.699740191 | 1.60E-14 | 6.67E-13 |
| ENSMUSG00000014198.15 | ZNF385C | 1.173340132 | 8.84E-05 | 6.56E-04 |
| ENSMUSG00000045333.15 | ZNF423 | -1.066470701 | 7.70E-08 | 1.14E-06 |
| ENSMUSG00000043903.4 | ZNF469 | 1.315910292 | 3.70E-06 | 3.91E-05 |
| ENSMUSG00000020193.3 | ZPBP | 1.115874068 | 4.86E-07 | 6.24E-06 |

**CB**

| ID | Symbol | Expr Log Ratio | Expr p-value | Expr FDR |
|---|---|---|---|---|
| ENSMUSG00000054091 | 1810037I17Rik/Gr | 0.481509384 | 9.94E-05 | 0.002636214 |
| ENSMUSG00000086841 | 2410006H16Rik | 0.507895519 | 2.14E-04 | 0.004417341 |
| ENSMUSG00000099881 | 2810013P06Rik | 0.415200995 | 2.07E-04 | 0.00431494 |
| ENSMUSG00000102386 | 2900022M07Rik | -1.495276207 | 3.14E-06 | 2.28E-04 |
| ENSMUSG00000099696 | 2900052N01Rik | 1.304923639 | 2.98E-05 | 0.001171341 |
| ENSMUSG00000109089 | 4833411C07Rik | 1.071580257 | 6.17E-04 | 0.009097627 |
| ENSMUSG00000114282 | 5330431K02Rik | -1.152084634 | 4.63E-05 | 0.00159614 |
| ENSMUSG00000106855 | 5330437M03Rik | -1.677577834 | 1.20E-07 | 1.90E-05 |
| ENSMUSG00000106951 | 5930430L01Rik | 0.81492266 | 1.91E-05 | 8.50E-04 |
| ENSMUSG00000109536 | 9330162G02Rik | -0.795495507 | 5.94E-05 | 0.001896804 |
| ENSMUSG00000108402 | 9430064I24Rik | -0.811426212 | 2.88E-04 | 0.005451446 |
| ENSMUSG00000109394 | A230057D06Rik | -0.604874324 | 1.93E-05 | 8.56E-04 |
| ENSMUSG00000109122 | A230103L15Rik | -1.100847795 | 1.40E-05 | 6.86E-04 |
| ENSMUSG00000096929 | A330023F24Rik | -0.922041317 | 1.58E-13 | 3.40E-10 |
| ENSMUSG00000028794 | A3GALT2 | -1.588826733 | 2.52E-07 | 3.47E-05 |
| ENSMUSG00000057230 | AAK1 | -0.418204628 | 3.44E-04 | 0.006220931 |
| ENSMUSG00000015243 | ABCA1 | -0.530538397 | 5.48E-06 | 3.36E-04 |
| ENSMUSG00000035722 | ABCA7 | -0.563393713 | 5.19E-06 | 3.27E-04 |
| ENSMUSG00000032842 | ABCC10 | -0.541615667 | 7.62E-05 | 0.00226248 |
| ENSMUSG00000022822 | ABCC5 | -0.550338388 | 6.20E-07 | 6.86E-05 |
| ENSMUSG00000040136 | ABCC8 | -0.763558272 | 2.68E-09 | 8.23E-07 |
| ENSMUSG00000032131 | ABCG4 | -0.435128927 | 3.18E-05 | 0.001230469 |
| ENSMUSG00000042073 | ABHD14B | -0.522469639 | 1.10E-04 | 0.002812978 |
| ENSMUSG00000113184 | AC125351.1 | -0.527760613 | 5.85E-04 | 0.008802883 |
| ENSMUSG00000118029 | AC132288.2 | -1.196160858 | 1.50E-05 | 7.15E-04 |
| ENSMUSG00000118550 | AC135859.1 | -0.998810685 | 6.00E-04 | 0.008946205 |
| ENSMUSG00000113328 | AC154347.1 | -1.015221006 | 1.96E-05 | 8.60E-04 |
| ENSMUSG00000026003 | ACADL | 0.416998191 | 3.73E-05 | 0.001357036 |
| ENSMUSG00000022185 | ACIN1 | -0.479411869 | 8.69E-06 | 4.81E-04 |
| ENSMUSG00000037872 | ACKR1 | -0.537428197 | 2.68E-06 | 2.03E-04 |
| ENSMUSG00000019948 | ACTR6 | 0.482040448 | 6.34E-04 | 0.009273289 |
| ENSMUSG00000008822 | ACYP1 | 0.468746293 | 3.90E-04 | 0.006820396 |
| ENSMUSG00000060923 | ACYP2 | 0.425675876 | 5.50E-04 | 0.00855549 |
| ENSMUSG00000020926 | ADAM11 | -0.706191725 | 1.32E-08 | 3.19E-06 |
| ENSMUSG00000072647 | Adam1a | -0.800454764 | 7.85E-05 | 0.002299423 |
| ENSMUSG00000024299 | ADAMTS10 | -0.903462935 | 5.46E-08 | 1.01E-05 |
| ENSMUSG00000049538 | ADAMTS16 | -1.021497606 | 2.15E-05 | 9.05E-04 |
| ENSMUSG00000043635 | ADAMTS3 | -0.69139643 | 7.10E-05 | 0.002153094 |
| ENSMUSG00000027951 | ADAR | -0.430347843 | 2.47E-04 | 0.004887592 |
| ENSMUSG00000022994 | ADCY6 | -0.492664703 | 5.08E-06 | 3.26E-04 |
| ENSMUSG00000037605 | ADGRL3 | -0.361512346 | 6.15E-04 | 0.009095975 |
| ENSMUSG00000039167 | ADGRL4 | 0.649529624 | 6.50E-07 | 7.00E-05 |
| ENSMUSG00000029313 | AFF1 | -0.413570578 | 2.35E-04 | 0.004726005 |

| | | | | |
|---|---|---|---|---|
| ENSMUSG00000041936 | AGRN | -0.579544317 | 1.87E-06 | 1.57E-04 |
| ENSMUSG00000090086 | AI480526 | -1.245412083 | 1.06E-16 | 5.30E-13 |
| ENSMUSG00000022763 | AIFM3 | -0.587835114 | 5.07E-07 | 6.00E-05 |
| ENSMUSG00000028029 | AIMP1 | 0.329628099 | 5.28E-04 | 0.008341841 |
| ENSMUSG00000029419 | AJM1 | -0.506670664 | 8.61E-05 | 0.002370781 |
| ENSMUSG00000024045 | AKAP8 | -0.712478482 | 2.03E-11 | 1.53E-08 |
| ENSMUSG00000002625 | AKAP8L | -0.64371716 | 6.61E-07 | 7.04E-05 |
| ENSMUSG00000040407 | Akap9 | -0.424228476 | 6.38E-04 | 0.009307819 |
| ENSMUSG00000053279 | ALDH1A1 | 0.38213736 | 2.19E-04 | 0.00448912 |
| ENSMUSG00000002661 | ALKBH7 | 0.442321543 | 3.14E-04 | 0.005811328 |
| ENSMUSG00000027889 | AMPD2 | -0.375064636 | 6.26E-04 | 0.009188555 |
| ENSMUSG00000025135 | ANAPC11 | 0.408685905 | 9.43E-05 | 0.002522112 |
| ENSMUSG00000035048 | ANAPC13 | 0.393406025 | 5.54E-04 | 0.008593437 |
| ENSMUSG00000031465 | ANGPT2 | 0.881230182 | 3.96E-04 | 0.006870605 |
| ENSMUSG00000002289 | ANGPTL4 | 0.894595181 | 2.35E-05 | 9.74E-04 |
| ENSMUSG00000032826 | Ank2 | -0.453397822 | 1.68E-04 | 0.003757229 |
| ENSMUSG00000024483 | ANKHD1/ANKHD1- | -0.493108217 | 1.18E-05 | 6.08E-04 |
| ENSMUSG00000035569 | ANKRD11 | -0.436049142 | 3.07E-04 | 0.005696385 |
| ENSMUSG00000037907 | ANKRD13B | -0.47925267 | 1.67E-05 | 7.71E-04 |
| ENSMUSG00000047909 | ANKRD16 | -0.64176657 | 2.60E-06 | 1.98E-04 |
| ENSMUSG00000054708 | ANKRD24 | -0.460609787 | 3.53E-04 | 0.006312887 |
| ENSMUSG00000049097 | ANKRD34A | -0.352157575 | 1.71E-04 | 0.003811675 |
| ENSMUSG00000014498 | ANKRD52 | -0.693165119 | 7.63E-08 | 1.29E-05 |
| ENSMUSG00000034863 | ANO8 | -0.397458175 | 2.83E-04 | 0.005386077 |
| ENSMUSG00000032231 | ANXA2 | 0.714904696 | 7.53E-05 | 0.002240932 |
| ENSMUSG00000021458 | AOPEP | -0.35777464 | 4.12E-04 | 0.007051437 |
| ENSMUSG00000040701 | AP1G2 | -0.899084968 | 1.97E-06 | 1.63E-04 |
| ENSMUSG00000029207 | APBB2 | -0.460862882 | 1.79E-05 | 8.21E-04 |
| ENSMUSG00000022548 | APOD | 0.379610898 | 3.77E-04 | 0.006663826 |
| ENSMUSG00000020263 | APPL2 | -0.416042908 | 1.52E-04 | 0.00355335 |
| ENSMUSG00000043144 | AQP6 | -0.650739977 | 3.05E-04 | 0.005681988 |
| ENSMUSG00000032812 | ARAP1 | -0.383267129 | 3.49E-04 | 0.006265556 |
| ENSMUSG00000040459 | ARGLU1 | -0.45656908 | 3.86E-06 | 2.71E-04 |
| ENSMUSG00000036452 | ARHGAP26 | -0.494357962 | 2.45E-06 | 1.93E-04 |
| ENSMUSG00000073433 | ARHGDIG | 0.949991505 | 8.19E-05 | 0.002331043 |
| ENSMUSG00000040940 | ARHGEF1 | -0.744012894 | 3.32E-10 | 1.39E-07 |
| ENSMUSG00000041977 | ARHGEF11 | -0.377896272 | 3.73E-04 | 0.006609229 |
| ENSMUSG00000019467 | ARHGEF25 | -0.365581038 | 5.61E-04 | 0.0086356 |
| ENSMUSG00000045094 | ARHGEF37 | -0.455905819 | 5.81E-04 | 0.008786783 |
| ENSMUSG00000007880 | ARID1A | -0.533589543 | 8.84E-05 | 0.002410934 |
| ENSMUSG00000069729 | ARID1B | -0.384494905 | 2.79E-04 | 0.005314345 |
| ENSMUSG00000060904 | ARL1 | 0.459488766 | 3.21E-05 | 0.001230469 |
| ENSMUSG00000049804 | ARMCX4 | -0.631337768 | 1.02E-04 | 0.002702126 |
| ENSMUSG00000015522 | ARNT | -0.318200231 | 6.22E-04 | 0.009160436 |
| ENSMUSG00000023017 | ASIC1 | -0.447306388 | 2.99E-04 | 0.005606734 |

| | | | | |
|---|---|---|---|---|
| ENSMUSG00000042548 | ASXL1 | -0.459227917 | 3.98E-05 | 0.001426173 |
| ENSMUSG00000099083 | ATF7 | -0.528372779 | 2.50E-04 | 0.004921781 |
| ENSMUSG00000047767 | ATG16L2 | -0.608319745 | 4.03E-06 | 2.79E-04 |
| ENSMUSG00000041341 | ATG2B | -0.398893829 | 2.59E-04 | 0.005052147 |
| ENSMUSG00000037400 | ATP11B | -0.339593228 | 2.63E-04 | 0.005114705 |
| ENSMUSG00000031862 | ATP13A1 | -0.355893694 | 2.22E-04 | 0.004525956 |
| ENSMUSG00000040907 | ATP1A3 | -0.447931937 | 1.20E-04 | 0.003028523 |
| ENSMUSG00000038690 | ATP5MF | 0.4787417 | 2.32E-04 | 0.004675686 |
| ENSMUSG00000004285 | ATP6V1F | 0.403072678 | 1.86E-04 | 0.004028856 |
| ENSMUSG00000031229 | ATRX | -0.443159819 | 5.28E-04 | 0.008341841 |
| ENSMUSG00000042605 | ATXN2 | -0.400050488 | 4.10E-05 | 0.001461022 |
| ENSMUSG00000032637 | ATXN2L | -0.772511732 | 1.38E-09 | 4.63E-07 |
| ENSMUSG00000048997 | ATXN7L2 | -0.778959299 | 2.80E-07 | 3.76E-05 |
| ENSMUSG00000065990 | AURKAIP1 | 0.372815138 | 5.77E-04 | 0.008767869 |
| ENSMUSG00000029673 | AUTS2 | -0.547737642 | 1.16E-05 | 6.01E-04 |
| ENSMUSG00000097428 | AW047730 | 0.370299386 | 6.86E-05 | 0.002112559 |
| ENSMUSG00000022296 | BAALC | 0.370020851 | 2.96E-04 | 0.005567282 |
| ENSMUSG00000040054 | BAZ2A | -0.462422961 | 4.80E-05 | 0.001635798 |
| ENSMUSG00000115783 | Bc1 | 0.827187245 | 5.89E-05 | 0.001895607 |
| ENSMUSG00000013523 | BCAS1 | 0.787387691 | 1.77E-06 | 1.50E-04 |
| ENSMUSG00000038256 | BCL9 | -0.496644062 | 2.95E-05 | 0.001168731 |
| ENSMUSG00000063382 | BCL9L | -0.406524545 | 5.60E-04 | 0.0086356 |
| ENSMUSG00000037608 | BCLAF1 | -0.411314499 | 8.06E-05 | 0.002312173 |
| ENSMUSG00000028167 | BDH2 | 0.864433167 | 5.80E-04 | 0.008786366 |
| ENSMUSG00000070808 | BICRA | -0.516497763 | 3.26E-04 | 0.005966137 |
| ENSMUSG00000036568 | BICRAL | -0.431239361 | 1.87E-04 | 0.004028856 |
| ENSMUSG00000022098 | BMP1 | -0.730155275 | 9.35E-07 | 8.85E-05 |
| ENSMUSG00000061755 | BOD1L1 | -0.444583927 | 9.59E-05 | 0.002561777 |
| ENSMUSG00000024002 | Brd4 | -0.549719829 | 1.24E-05 | 6.28E-04 |
| ENSMUSG00000033940 | BRK1 | 0.379908479 | 1.34E-04 | 0.003272115 |
| ENSMUSG00000001632 | BRPF1 | -0.326360855 | 5.34E-04 | 0.008385423 |
| ENSMUSG00000032589 | BSN | -0.54031161 | 8.02E-05 | 0.002312173 |
| ENSMUSG00000104339 | C130089K02Rik | -1.47532972 | 1.81E-05 | 8.21E-04 |
| ENSMUSG00000020133 | C19orf25 | 0.425399916 | 9.81E-05 | 0.002609492 |
| ENSMUSG00000000581 | C1D | 0.460660683 | 7.27E-05 | 0.002190239 |
| ENSMUSG00000036887 | C1QA | 0.719303962 | 1.72E-06 | 1.47E-04 |
| ENSMUSG00000036905 | C1QB | 0.745491757 | 2.14E-06 | 1.76E-04 |
| ENSMUSG00000110523 | C230057M02Rik | -0.463576565 | 4.46E-04 | 0.007450672 |
| ENSMUSG00000058706 | C2orf68 | -0.477315303 | 4.82E-04 | 0.007851613 |
| ENSMUSG00000073418 | C4A/C4B | 0.985954489 | 1.48E-07 | 2.25E-05 |
| ENSMUSG00000110710 | C78859 | -1.04834295 | 7.42E-09 | 2.03E-06 |
| ENSMUSG00000009075 | CABP7 | 0.468552983 | 2.81E-06 | 2.09E-04 |
| ENSMUSG00000010066 | CACNA2D2 | -0.422708055 | 2.03E-04 | 0.004261143 |
| ENSMUSG00000013629 | CAD | -0.454009047 | 1.95E-04 | 0.004168127 |
| ENSMUSG00000053819 | CAMK2D | -0.428751937 | 3.39E-05 | 0.001261965 |

| | | | | |
|---|---|---|---|---|
| ENSMUSG00000020785 | CAMKK1 | -0.340733665 | 6.23E-05 | 0.001968283 |
| ENSMUSG00000026933 | CAMSAP1 | -0.421746638 | 2.76E-04 | 0.005276112 |
| ENSMUSG00000056737 | CAPG | 0.938277189 | 1.39E-05 | 6.83E-04 |
| ENSMUSG00000022211 | CARMIL3 | -0.628409169 | 2.42E-06 | 1.93E-04 |
| ENSMUSG00000078676 | CASC3 | -0.509872698 | 9.33E-07 | 8.85E-05 |
| ENSMUSG00000025888 | CASP1 | 0.874775865 | 3.48E-04 | 0.006252666 |
| ENSMUSG00000007655 | CAV1 | 0.779735754 | 9.14E-07 | 8.81E-05 |
| ENSMUSG00000034342 | CBL | -0.489357879 | 6.76E-05 | 0.002098897 |
| ENSMUSG00000024647 | CBLN2 | 1.668715925 | 1.78E-04 | 0.003904244 |
| ENSMUSG00000031150 | CCDC120 | -0.444200862 | 2.91E-04 | 0.0054973 |
| ENSMUSG00000026676 | CCDC3 | 0.829924644 | 4.56E-06 | 3.05E-04 |
| ENSMUSG00000048701 | CCDC6 | -0.392728509 | 4.70E-04 | 0.007719686 |
| ENSMUSG00000041375 | CCDC9 | -0.359109331 | 6.59E-04 | 0.009520404 |
| ENSMUSG00000030898 | CCKBR | 0.982470361 | 1.48E-04 | 0.003481205 |
| ENSMUSG00000027829 | CCNL1 | -0.357424527 | 8.11E-05 | 0.002317862 |
| ENSMUSG00000029068 | CCNL2 | -0.524106464 | 2.18E-07 | 3.12E-05 |
| ENSMUSG00000026349 | CCNT2 | -0.313036682 | 6.64E-04 | 0.009562184 |
| ENSMUSG00000025510 | CD151 | 0.422682627 | 2.74E-04 | 0.005268967 |
| ENSMUSG00000047139 | Cd24a | 0.854042624 | 3.05E-05 | 0.001192614 |
| ENSMUSG00000060703 | CD302 | 0.565485974 | 1.79E-04 | 0.003904244 |
| ENSMUSG00000025351 | CD63 | 0.459572344 | 3.34E-04 | 0.006071117 |
| ENSMUSG00000015396 | CD83 | 0.625192907 | 7.62E-09 | 2.04E-06 |
| ENSMUSG00000030342 | CD9 | 0.511525604 | 9.95E-06 | 5.38E-04 |
| ENSMUSG00000027284 | CDAN1 | -0.39704816 | 2.77E-04 | 0.005295482 |
| ENSMUSG00000024769 | CDC42BPG | -0.343130574 | 8.05E-05 | 0.002312173 |
| ENSMUSG00000050910 | CDR2L | -0.350550964 | 7.53E-05 | 0.002240932 |
| ENSMUSG00000005506 | Celf1 | -0.436865905 | 9.02E-05 | 0.00244406 |
| ENSMUSG00000028137 | CELF3 | -0.535023672 | 4.50E-05 | 0.001566513 |
| ENSMUSG00000034818 | CELF5 | -0.65822735 | 4.19E-06 | 2.87E-04 |
| ENSMUSG00000023473 | CELSR3 | -0.838992197 | 1.07E-08 | 2.74E-06 |
| ENSMUSG00000039781 | CEP131 | -0.532547277 | 1.05E-04 | 0.002743595 |
| ENSMUSG00000043987 | CEP164 | -0.681679145 | 2.05E-08 | 4.73E-06 |
| ENSMUSG00000038241 | CEP250 | -0.634064404 | 2.41E-05 | 9.91E-04 |
| ENSMUSG00000018372 | CEP95 | -0.501183693 | 1.56E-04 | 0.003599362 |
| ENSMUSG00000078490 | CFAP74 | -0.906018799 | 3.86E-07 | 4.87E-05 |
| ENSMUSG00000001128 | CFP | -0.57922072 | 2.26E-04 | 0.00458579 |
| ENSMUSG00000068876 | CGN | -0.779514256 | 4.71E-04 | 0.007720886 |
| ENSMUSG00000032232 | CGNL1 | -0.502598463 | 9.77E-07 | 9.12E-05 |
| ENSMUSG00000030086 | CHCHD6 | 0.434378344 | 4.84E-04 | 0.007865248 |
| ENSMUSG00000063870 | CHD4 | -0.427484823 | 1.37E-04 | 0.003322569 |
| ENSMUSG00000057133 | CHD6 | -0.448147611 | 1.04E-04 | 0.002724288 |
| ENSMUSG00000014668 | CHFR | -0.341282847 | 3.24E-04 | 0.005927339 |
| ENSMUSG00000028419 | CHMP5 | 0.357425928 | 2.50E-04 | 0.004921781 |
| ENSMUSG00000038181 | CHPF2 | -0.39260671 | 4.06E-04 | 0.006963461 |
| ENSMUSG00000006958 | CHRD | -0.774701906 | 1.36E-10 | 6.83E-08 |

| | | | | |
|---|---|---|---|---|
| ENSMUSG00000037493 | CIB2 | 0.429124919 | 1.26E-04 | 0.003141708 |
| ENSMUSG00000039205 | CIZ1 | -0.530669634 | 2.30E-07 | 3.24E-05 |
| ENSMUSG00000064302 | CLASP1 | -0.370848662 | 6.66E-04 | 0.009582672 |
| ENSMUSG00000061028 | Clasrp | -0.645481299 | 2.11E-08 | 4.74E-06 |
| ENSMUSG00000029862 | CLCN1 | -0.710554007 | 3.54E-05 | 0.001309114 |
| ENSMUSG00000022132 | CLDN10 | 0.459284034 | 2.05E-05 | 8.80E-04 |
| ENSMUSG00000049550 | CLIP1 | -0.416275875 | 8.28E-05 | 0.002331043 |
| ENSMUSG00000013921 | CLIP3 | -0.48184054 | 3.22E-05 | 0.001231135 |
| ENSMUSG00000020385 | CLK4 | -0.376254657 | 2.39E-04 | 0.004790653 |
| ENSMUSG00000025545 | CLYBL | 0.488975165 | 8.37E-05 | 0.002339439 |
| ENSMUSG00000040759 | CMTM5 | 0.464297103 | 7.68E-05 | 0.002271947 |
| ENSMUSG00000035632 | CNOT3 | -0.566906104 | 5.96E-06 | 3.61E-04 |
| ENSMUSG00000044681 | CNPY1 | -0.824345451 | 2.33E-10 | 1.06E-07 |
| ENSMUSG00000025381 | CNPY2 | 0.482198108 | 8.76E-05 | 0.002399469 |
| ENSMUSG00000017188 | COA3 | 0.441916817 | 3.28E-04 | 0.005982584 |
| ENSMUSG00000024330 | COL11A2 | -0.620729232 | 3.28E-04 | 0.005982584 |
| ENSMUSG00000001435 | COL18A1 | -0.4894822 | 1.67E-05 | 7.72E-04 |
| ENSMUSG00000045672 | COL27A1 | -1.532493338 | 2.34E-12 | 3.91E-09 |
| ENSMUSG00000026837 | COL5A1 | -1.191046138 | 5.07E-12 | 5.86E-09 |
| ENSMUSG00000001119 | COL6A1 | -0.76161459 | 5.25E-04 | 0.008323206 |
| ENSMUSG00000025650 | COL7A1 | -1.687817702 | 6.96E-12 | 6.97E-09 |
| ENSMUSG00000051154 | COMMD3 | 0.334270677 | 5.50E-04 | 0.00855549 |
| ENSMUSG00000075486 | Commd6 | 0.398903978 | 4.25E-04 | 0.007193338 |
| ENSMUSG00000073616 | COPS9 | 0.496744249 | 1.65E-04 | 0.003729993 |
| ENSMUSG00000020836 | CORO6 | -0.953766496 | 7.49E-13 | 1.41E-09 |
| ENSMUSG00000039801 | CPLANE1 | -0.462637154 | 8.51E-05 | 0.002352671 |
| ENSMUSG00000025867 | CPLX2 | -0.400143545 | 2.08E-04 | 0.004322372 |
| ENSMUSG00000024008 | CPNE5 | 1.739497558 | 1.39E-04 | 0.003346496 |
| ENSMUSG00000039007 | CPQ | 0.505108302 | 3.31E-05 | 0.001247629 |
| ENSMUSG00000034022 | CPSF1 | -0.385038214 | 5.41E-04 | 0.008463273 |
| ENSMUSG00000029625 | CPSF4 | -0.507648456 | 1.89E-05 | 8.50E-04 |
| ENSMUSG00000055531 | CPSF6 | -0.423643026 | 6.18E-04 | 0.009110373 |
| ENSMUSG00000034820 | CPSF7 | -0.357773221 | 2.46E-04 | 0.00488453 |
| ENSMUSG00000038002 | CRAMP1 | -0.430162765 | 2.86E-04 | 0.005433428 |
| ENSMUSG00000022521 | CREBBP | -0.459236593 | 1.48E-04 | 0.003498019 |
| ENSMUSG00000051451 | CREBZF | -0.303312916 | 3.46E-04 | 0.006236602 |
| ENSMUSG00000021680 | CRHBP | 3.952527159 | 4.48E-20 | 3.37E-16 |
| ENSMUSG00000006356 | Crip2 | 0.460125794 | 4.38E-04 | 0.007346812 |
| ENSMUSG00000027936 | CRTC2 | -0.444701842 | 8.66E-06 | 4.81E-04 |
| ENSMUSG00000026421 | CSRP1 | 0.453817022 | 1.22E-04 | 0.003065766 |
| ENSMUSG00000027447 | CST3 | 0.55876723 | 3.44E-06 | 2.46E-04 |
| ENSMUSG00000031256 | CSTF2 | -0.394702251 | 1.03E-05 | 5.54E-04 |
| ENSMUSG00000113226 | CT009765.1 | -1.184779402 | 5.91E-04 | 0.008871447 |
| ENSMUSG00000044258 | Ctla2a/Ctla2b | 0.954813189 | 4.32E-05 | 0.001525562 |
| ENSMUSG00000038642 | CTSS | 0.574566439 | 7.87E-05 | 0.002299423 |

| | | | | |
|---|---|---|---|---|
| ENSMUSG00000000416 | CTTNBP2 | -0.393550333 | 1.90E-04 | 0.004075474 |
| ENSMUSG00000021508 | CXCL14 | 0.369818514 | 2.78E-05 | 0.001117851 |
| ENSMUSG00000024646 | CYB5A | 0.568577986 | 1.12E-07 | 1.80E-05 |
| ENSMUSG00000046727 | Cystm1 | 0.343490215 | 3.86E-04 | 0.006782538 |
| ENSMUSG00000094910 | D430019H16Rik | 1.113733306 | 2.46E-04 | 0.004887592 |
| ENSMUSG00000028519 | DAB1 | -0.426220481 | 1.24E-05 | 6.28E-04 |
| ENSMUSG00000000889 | DBH | 1.786679449 | 1.21E-05 | 6.17E-04 |
| ENSMUSG00000027797 | DCLK1 | -0.369145236 | 2.68E-04 | 0.005195082 |
| ENSMUSG00000028078 | DCLK2 | -0.386914952 | 5.17E-04 | 0.008206561 |
| ENSMUSG00000055065 | DDX17 | -0.491483095 | 5.16E-06 | 3.27E-04 |
| ENSMUSG00000021500 | DDX46 | -0.436676612 | 1.90E-05 | 8.50E-04 |
| ENSMUSG00000054763 | Defb42 | 0.986880969 | 4.72E-06 | 3.07E-04 |
| ENSMUSG00000015377 | DENND6B | -0.610583925 | 5.32E-07 | 6.20E-05 |
| ENSMUSG00000003531 | Dgcr6 | 0.331936209 | 5.22E-04 | 0.008285543 |
| ENSMUSG00000022861 | DGKG | -0.378640119 | 3.99E-04 | 0.006914468 |
| ENSMUSG00000004815 | DGKQ | -0.425493198 | 1.21E-04 | 0.003049262 |
| ENSMUSG00000034926 | DHCR24 | 0.314478075 | 5.29E-04 | 0.008348481 |
| ENSMUSG00000042569 | DHRS7B | 0.493345746 | 2.54E-04 | 0.004978129 |
| ENSMUSG00000041415 | DICER1 | -0.437935021 | 6.63E-04 | 0.009556248 |
| ENSMUSG00000038914 | DIDO1 | -0.334568123 | 2.01E-04 | 0.004246312 |
| ENSMUSG00000030409 | DMPK | -0.498949229 | 6.82E-05 | 0.002111064 |
| ENSMUSG00000036052 | DNAJB5 | -0.379875817 | 5.44E-05 | 0.001807828 |
| ENSMUSG00000030882 | DNHD1 | -0.868943077 | 5.79E-05 | 0.001874552 |
| ENSMUSG00000004099 | DNMT1 | -0.399454435 | 1.37E-04 | 0.003316671 |
| ENSMUSG00000020661 | DNMT3A | -0.65321095 | 3.02E-06 | 2.22E-04 |
| ENSMUSG00000025558 | DOCK9 | -0.419094374 | 1.85E-04 | 0.004006798 |
| ENSMUSG00000035711 | DOK3 | -0.755894187 | 4.47E-04 | 0.007450672 |
| ENSMUSG00000034973 | DOP1A | -0.333402084 | 1.58E-04 | 0.003632905 |
| ENSMUSG00000061589 | DOT1L | -0.930658988 | 4.79E-12 | 5.86E-09 |
| ENSMUSG00000025478 | DPYSL4 | -0.623696661 | 5.05E-10 | 1.95E-07 |
| ENSMUSG00000030002 | DUSP11 | -0.673326501 | 3.25E-10 | 1.39E-07 |
| ENSMUSG00000003233 | DVL3 | -0.572122259 | 6.46E-08 | 1.17E-05 |
| ENSMUSG00000024137 | E4F1 | -0.406429273 | 1.43E-04 | 0.003403032 |
| ENSMUSG00000053898 | ECH1 | 0.45313643 | 6.44E-05 | 0.00201294 |
| ENSMUSG00000025465 | ECHS1 | 0.3968058 | 1.28E-04 | 0.003164637 |
| ENSMUSG00000024132 | ECI1 | 0.498800479 | 3.92E-05 | 0.001405956 |
| ENSMUSG00000036270 | EDC4 | -0.374555388 | 6.10E-05 | 0.001933985 |
| ENSMUSG00000037742 | EEF1A1 | 0.441060834 | 5.42E-04 | 0.008463273 |
| ENSMUSG00000025967 | EEF1B2 | 0.411193065 | 1.61E-04 | 0.003678007 |
| ENSMUSG00000035064 | EEF2K | -0.386466455 | 1.58E-04 | 0.003633193 |
| ENSMUSG00000003934 | EFNB3 | 0.352935333 | 6.24E-04 | 0.009173637 |
| ENSMUSG00000022312 | EIF3H | 0.423113064 | 3.22E-04 | 0.005905704 |
| ENSMUSG00000028798 | EIF3I | 0.39407655 | 5.64E-04 | 0.008651476 |
| ENSMUSG00000045983 | EIF4G1 | -0.460128507 | 1.94E-05 | 8.56E-04 |
| ENSMUSG00000028760 | EIF4G3 | -0.556631348 | 1.02E-06 | 9.45E-05 |

| | | | | |
|---|---|---|---|---|
| ENSMUSG00000026083 | EIF5B | -0.371712181 | 5.55E-04 | 0.008597182 |
| ENSMUSG00000048988 | ELFN1 | 0.971926973 | 5.62E-04 | 0.0086356 |
| ENSMUSG00000051166 | EML5 | -0.797214044 | 1.09E-10 | 6.07E-08 |
| ENSMUSG00000028445 | ENHO | 0.454179662 | 7.85E-06 | 4.41E-04 |
| ENSMUSG00000026927 | ENTR1 | -0.476122679 | 1.46E-06 | 1.28E-04 |
| ENSMUSG00000032446 | EOMES | -1.07443111 | 1.41E-05 | 6.86E-04 |
| ENSMUSG00000006276 | EPS15L1 | -0.354399265 | 9.39E-05 | 0.002520287 |
| ENSMUSG00000021996 | ESD | 0.43764361 | 9.50E-06 | 5.15E-04 |
| ENSMUSG00000061286 | EXOSC5 | 0.494797378 | 5.98E-04 | 0.008932034 |
| ENSMUSG00000027533 | FABP5 | 0.555767787 | 2.15E-05 | 9.05E-04 |
| ENSMUSG00000021750 | FAM107A | 0.548544304 | 2.24E-06 | 1.80E-04 |
| ENSMUSG00000037210 | FAM193A | -0.402918114 | 5.54E-04 | 0.008593437 |
| ENSMUSG00000021495 | Fam193b | -0.702894027 | 2.01E-08 | 4.71E-06 |
| ENSMUSG00000070047 | FAT1 | -0.533045254 | 4.05E-04 | 0.006963461 |
| ENSMUSG00000038274 | FAU | 0.418249116 | 4.01E-04 | 0.006940415 |
| ENSMUSG00000042423 | FBRS | -0.502877014 | 4.48E-06 | 3.03E-04 |
| ENSMUSG00000043323 | FBRSL1 | -0.453465715 | 6.86E-06 | 3.98E-04 |
| ENSMUSG00000030811 | FBXL19 | -0.428272998 | 2.36E-04 | 0.004736697 |
| ENSMUSG00000050503 | Fbxl22 | -1.135800608 | 5.14E-05 | 0.001723063 |
| ENSMUSG00000028920 | FBXO42 | -0.433373378 | 7.28E-05 | 0.002190239 |
| ENSMUSG00000033703 | FCSK | -0.386735304 | 4.02E-04 | 0.006940415 |
| ENSMUSG00000021273 | FDFT1 | 0.386193448 | 5.98E-04 | 0.008932034 |
| ENSMUSG00000044465 | FHIP1B | -0.418438028 | 1.96E-04 | 0.004171854 |
| ENSMUSG00000085396 | Firre | -0.731345569 | 1.49E-10 | 7.21E-08 |
| ENSMUSG00000031328 | FLNA | -0.427661354 | 4.85E-05 | 0.001644745 |
| ENSMUSG00000025278 | FLNB | -0.622613761 | 1.12E-08 | 2.81E-06 |
| ENSMUSG00000019689 | FMC1 | 0.515305074 | 1.45E-04 | 0.003449153 |
| ENSMUSG00000008200 | FNBP4 | -0.531242773 | 2.08E-08 | 4.73E-06 |
| ENSMUSG00000003154 | FOXJ2 | -0.547583325 | 4.54E-06 | 3.05E-04 |
| ENSMUSG00000026657 | FRMD4A | -0.416252372 | 1.08E-04 | 0.002796518 |
| ENSMUSG00000024661 | FTH1 | 0.624093577 | 1.89E-06 | 1.58E-04 |
| ENSMUSG00000030795 | Fus | -0.485823366 | 3.02E-06 | 2.22E-04 |
| ENSMUSG00000036570 | FXYD1 | 0.54467738 | 4.00E-07 | 4.98E-05 |
| ENSMUSG00000009687 | FXYD5 | 0.76602547 | 6.33E-04 | 0.009267885 |
| ENSMUSG00000049551 | FZD9 | 0.846928167 | 5.67E-07 | 6.49E-05 |
| ENSMUSG00000038766 | GABPB2 | -0.471539175 | 7.38E-05 | 0.002214003 |
| ENSMUSG00000001260 | GABRG1 | 0.466799293 | 4.99E-05 | 0.001686872 |
| ENSMUSG00000028270 | GBP2 | 1.075460311 | 8.05E-05 | 0.002312173 |
| ENSMUSG00000041638 | GCN1 | -0.432344667 | 1.93E-04 | 0.004139393 |
| ENSMUSG00000034424 | GCSH | 0.387915999 | 4.28E-04 | 0.007206888 |
| ENSMUSG00000058624 | GDA | 1.507559126 | 1.04E-04 | 0.002724288 |
| ENSMUSG00000035314 | GDPD5 | -0.367911109 | 5.72E-05 | 0.001859711 |
| ENSMUSG00000020932 | GFAP | 0.879489837 | 1.56E-06 | 1.35E-04 |
| ENSMUSG00000020740 | GGA3 | -0.533620388 | 1.56E-07 | 2.34E-05 |
| ENSMUSG00000041625 | GGACT | 0.319830176 | 6.85E-04 | 0.009777308 |

| | | | | |
|---|---|---|---|---|
| ENSMUSG00000029714 | GIGYF1 | -0.753301429 | 3.16E-08 | 6.68E-06 |
| ENSMUSG00000040055 | GJB6 | 0.34607409 | 4.84E-04 | 0.007865248 |
| ENSMUSG00000087042 | Gm11611 | -0.803418665 | 2.63E-05 | 0.001068276 |
| ENSMUSG00000085334 | Gm12940 | -0.767143092 | 4.64E-06 | 3.06E-04 |
| ENSMUSG00000084904 | Gm14827 | -0.801322205 | 2.03E-04 | 0.004261143 |
| ENSMUSG00000091177 | Gm15494 | -1.278813444 | 8.25E-05 | 0.002331043 |
| ENSMUSG00000090785 | Gm17116 | -1.077488023 | 8.39E-05 | 0.002339439 |
| ENSMUSG00000097042 | Gm17491 | -0.706248752 | 2.10E-04 | 0.004352697 |
| ENSMUSG00000111092 | Gm17875 | -1.524081697 | 5.74E-07 | 6.49E-05 |
| ENSMUSG00000109724 | Gm18194 | -1.133507692 | 2.56E-04 | 0.005011761 |
| ENSMUSG00000110332 | Gm19935 | 0.997516342 | 1.52E-05 | 7.15E-04 |
| ENSMUSG00000103983 | Gm20045 | -0.928852552 | 3.53E-06 | 2.52E-04 |
| ENSMUSG00000095123 | Gm21781 | -0.728806067 | 1.50E-04 | 0.003523842 |
| ENSMUSG00000101356 | Gm28876 | -1.162089622 | 7.21E-05 | 0.002180813 |
| ENSMUSG00000100201 | Gm29093 | -0.967859344 | 4.82E-04 | 0.007851613 |
| ENSMUSG00000100417 | Gm29595 | 1.081244967 | 1.51E-04 | 0.003541627 |
| ENSMUSG00000108804 | Gm30437 | -0.914449971 | 4.94E-04 | 0.00794112 |
| ENSMUSG00000092090 | Gm3294 | -0.949630812 | 1.45E-07 | 2.23E-05 |
| ENSMUSG00000110390 | Gm33320 | -1.363300469 | 3.76E-11 | 2.26E-08 |
| ENSMUSG00000109179 | Gm35339 | -0.939396994 | 2.79E-06 | 2.08E-04 |
| ENSMUSG00000102953 | Gm37019 | -1.525380521 | 3.50E-04 | 0.00627791 |
| ENSMUSG00000103651 | Gm37206 | -0.896746796 | 2.87E-05 | 0.001138765 |
| ENSMUSG00000103046 | Gm37309 | -1.677607217 | 5.44E-06 | 3.36E-04 |
| ENSMUSG00000102747 | Gm37602 | -1.930470803 | 1.27E-09 | 4.44E-07 |
| ENSMUSG00000097156 | Gm3764 | -0.83273258 | 4.11E-08 | 8.24E-06 |
| ENSMUSG00000102813 | Gm37795 | -1.250845269 | 6.71E-07 | 7.05E-05 |
| ENSMUSG00000103928 | Gm37893 | -1.065594521 | 3.20E-05 | 0.001230469 |
| ENSMUSG00000104399 | Gm37963 | -1.291854105 | 2.48E-06 | 1.93E-04 |
| ENSMUSG00000104154 | Gm38104 | -1.330986034 | 8.53E-07 | 8.33E-05 |
| ENSMUSG00000104277 | Gm38299 | -1.202226806 | 2.56E-04 | 0.005007609 |
| ENSMUSG00000111389 | Gm39465 | -0.947126209 | 1.99E-04 | 0.004214777 |
| ENSMUSG00000106099 | Gm42664 | -1.079838148 | 1.32E-05 | 6.57E-04 |
| ENSMUSG00000105742 | Gm42748 | -1.263425086 | 7.82E-06 | 4.40E-04 |
| ENSMUSG00000105983 | Gm42770 | -1.64688751 | 3.25E-08 | 6.68E-06 |
| ENSMUSG00000107021 | Gm42853 | -1.214507127 | 4.86E-06 | 3.15E-04 |
| ENSMUSG00000104737 | Gm42937 | -1.299607345 | 5.19E-05 | 0.001732936 |
| ENSMUSG00000104814 | Gm42979 | -1.045528139 | 1.03E-06 | 9.54E-05 |
| ENSMUSG00000104563 | Gm43041 | -1.2156969 | 1.74E-07 | 2.57E-05 |
| ENSMUSG00000107374 | Gm43172 | -1.550973214 | 3.05E-07 | 4.06E-05 |
| ENSMUSG00000107000 | Gm43481 | -1.22521109 | 1.40E-04 | 0.003356404 |
| ENSMUSG00000105368 | Gm43759 | -1.312066663 | 4.15E-07 | 5.07E-05 |
| ENSMUSG00000104546 | Gm43858 | -1.153702336 | 1.81E-05 | 8.21E-04 |
| ENSMUSG00000108297 | Gm44167 | -1.689743213 | 7.20E-07 | 7.41E-05 |
| ENSMUSG00000107620 | Gm44256 | -1.383925811 | 8.99E-05 | 0.002438907 |
| ENSMUSG00000108943 | Gm44559 | -1.86001384 | 1.94E-07 | 2.83E-05 |

| | | | | |
|---|---|---|---|---|
| ENSMUSG00000109362 | Gm44562 | -1.536618178 | 5.61E-07 | 6.49E-05 |
| ENSMUSG00000109555 | Gm44891 | -1.419443766 | 2.26E-05 | 9.43E-04 |
| ENSMUSG00000092178 | Gm45351 | 1.03263364 | 6.90E-04 | 0.009818627 |
| ENSMUSG00000110077 | Gm45632 | -1.231695233 | 2.14E-05 | 9.05E-04 |
| ENSMUSG00000113427 | Gm46378 | -0.912796663 | 4.55E-04 | 0.007549612 |
| ENSMUSG00000096768 | Gm47283 | -0.753096 | 1.62E-04 | 0.003684718 |
| ENSMUSG00000109378 | Gm49396 | -1.060087319 | 1.50E-05 | 7.15E-04 |
| ENSMUSG00000024869 | Gm49405 | -1.020285021 | 1.10E-04 | 0.002812978 |
| ENSMUSG00000074280 | Gm6166 | 0.730597102 | 3.35E-05 | 0.001254219 |
| ENSMUSG00000066553 | Gm6969 | 0.501451467 | 5.17E-04 | 0.008206561 |
| ENSMUSG00000106237 | Gm8066 | -0.83753733 | 2.02E-04 | 0.004258817 |
| ENSMUSG00000063696 | Gm8730 | 0.47790746 | 2.78E-04 | 0.005295482 |
| ENSMUSG00000053214 | Gm9899 | -0.386372871 | 4.33E-04 | 0.007286968 |
| ENSMUSG00000033021 | GMPPA | -0.439270137 | 4.55E-06 | 3.05E-04 |
| ENSMUSG00000032766 | GNG11 | 0.659408219 | 4.25E-05 | 0.001504985 |
| ENSMUSG00000029502 | GOLGA3 | -0.357096102 | 2.96E-04 | 0.005575499 |
| ENSMUSG00000038708 | GOLGA4 | -0.446514289 | 2.69E-04 | 0.00520685 |
| ENSMUSG00000034243 | GOLGB1 | -0.508491368 | 1.69E-06 | 1.45E-04 |
| ENSMUSG00000054199 | GON4L | -0.469120555 | 1.99E-05 | 8.69E-04 |
| ENSMUSG00000034621 | GPATCH8 | -0.515281059 | 2.85E-06 | 2.11E-04 |
| ENSMUSG00000063856 | GPX1 | 0.468151018 | 8.32E-05 | 0.002337637 |
| ENSMUSG00000018339 | GPX3 | 0.895037568 | 1.25E-04 | 0.003120886 |
| ENSMUSG00000075706 | GPX4 | 0.475730602 | 4.98E-04 | 0.007979318 |
| ENSMUSG00000040111 | GRAMD1B | -0.465609567 | 2.63E-05 | 0.001068276 |
| ENSMUSG00000059003 | GRIN2A | -0.458274881 | 5.34E-06 | 3.33E-04 |
| ENSMUSG00000020734 | GRIN2C | -0.412366027 | 3.93E-04 | 0.006854783 |
| ENSMUSG00000029198 | GRPEL1 | 0.383562489 | 2.25E-04 | 0.004585292 |
| ENSMUSG00000032348 | Gsta4 | 0.477695951 | 1.38E-04 | 0.003327256 |
| ENSMUSG00000004035 | GSTM2 | 0.6399311 | 6.37E-05 | 0.002001947 |
| ENSMUSG00000004032 | GSTM3 | 0.525803401 | 4.65E-05 | 0.00159614 |
| ENSMUSG00000034345 | GTF2H5 | 0.350573793 | 6.60E-05 | 0.002057798 |
| ENSMUSG00000023952 | GTPBP2 | -0.495557604 | 1.09E-06 | 9.97E-05 |
| ENSMUSG00000020444 | GUK1 | 0.473382232 | 5.80E-05 | 0.001874552 |
| ENSMUSG00000036181 | H1-2 | 0.870385215 | 5.16E-06 | 3.27E-04 |
| ENSMUSG00000051627 | H1f4 | 0.96990444 | 4.45E-05 | 0.001556764 |
| ENSMUSG00000041126 | H2AZ2 | 0.43845893 | 5.07E-05 | 0.001709213 |
| ENSMUSG00000018102 | H2BC10 | 0.669361796 | 1.09E-05 | 5.76E-04 |
| ENSMUSG00000056895 | H2BC5 | 0.567251416 | 7.59E-07 | 7.66E-05 |
| ENSMUSG00000060639 | H4-16 | 0.839915147 | 6.60E-04 | 0.009521487 |
| ENSMUSG00000059447 | HADHB | 0.382871698 | 6.52E-04 | 0.00944569 |
| ENSMUSG00000031386 | HCFC1 | -0.388060522 | 6.58E-04 | 0.009506346 |
| ENSMUSG00000031161 | HDAC6 | -0.407080197 | 2.36E-04 | 0.004736697 |
| ENSMUSG00000022475 | HDAC7 | -0.656299234 | 3.23E-07 | 4.20E-05 |
| ENSMUSG00000030532 | HDDC3 | 0.398828056 | 5.81E-04 | 0.008786783 |
| ENSMUSG00000002833 | HDGFL2 | -0.403500554 | 3.18E-05 | 0.001230469 |

| | | | | |
|---|---|---|---|---|
| ENSMUSG00000042770 | HEBP1 | 0.640208397 | 1.75E-04 | 0.003863565 |
| ENSMUSG00000035247 | HECTD1 | -0.434027285 | 3.20E-04 | 0.005889107 |
| ENSMUSG00000042744 | HECTD4 | -0.599455302 | 3.28E-05 | 0.001243861 |
| ENSMUSG00000031209 | HEPH | 0.701064826 | 4.23E-04 | 0.00716837 |
| ENSMUSG00000030451 | HERC2 | -0.500220188 | 5.93E-05 | 0.001896804 |
| ENSMUSG00000021665 | HEXB | 0.382962307 | 3.86E-04 | 0.006786995 |
| ENSMUSG00000073405 | HLA-E | 1.10726209 | 3.12E-04 | 0.005792085 |
| ENSMUSG00000027875 | HMGCS2 | 0.922435319 | 4.97E-08 | 9.64E-06 |
| ENSMUSG00000003038 | Hmgn2 (includes c | 0.385101239 | 1.90E-04 | 0.0040806 |
| ENSMUSG00000059208 | HNRNPM | -0.412236613 | 3.43E-05 | 0.001268571 |
| ENSMUSG00000039630 | HNRNPU | -0.488647889 | 5.10E-06 | 3.26E-04 |
| ENSMUSG00000078591 | HS3ST4 | 1.250588851 | 1.22E-04 | 0.003065744 |
| ENSMUSG00000031839 | HSBP1 | 0.397763506 | 1.81E-04 | 0.003930889 |
| ENSMUSG00000025260 | HSD17B10 | 0.502704032 | 5.67E-05 | 0.001853646 |
| ENSMUSG00000029311 | HSD17B11 | 0.454691128 | 2.26E-05 | 9.43E-04 |
| ENSMUSG00000039745 | HTATIP2 | 0.70116013 | 5.76E-04 | 0.008756534 |
| ENSMUSG00000034525 | ICE1 | -0.386690818 | 4.38E-05 | 0.001538007 |
| ENSMUSG00000025950 | IDH1 | 0.385845441 | 3.70E-04 | 0.006580748 |
| ENSMUSG00000058258 | IDI1 | 0.482259768 | 8.70E-06 | 4.81E-04 |
| ENSMUSG00000023830 | IGF2R | -0.378961924 | 9.21E-05 | 0.002481169 |
| ENSMUSG00000026185 | IGFBP5 | -0.593714384 | 6.00E-07 | 6.73E-05 |
| ENSMUSG00000034275 | IGSF9B | -0.705790246 | 6.71E-06 | 3.94E-04 |
| ENSMUSG00000031537 | IKBKB | -0.323139003 | 5.71E-04 | 0.008716871 |
| ENSMUSG00000044244 | IL20RB | -0.462844802 | 7.45E-05 | 0.002226046 |
| ENSMUSG00000032968 | INHA | -0.658029115 | 8.89E-06 | 4.89E-04 |
| ENSMUSG00000028894 | INPP5B | -0.417879355 | 1.43E-05 | 6.93E-04 |
| ENSMUSG00000026925 | INPP5E | -0.318559241 | 3.94E-04 | 0.006854783 |
| ENSMUSG00000034570 | INPP5J | -0.57420563 | 4.83E-07 | 5.81E-05 |
| ENSMUSG00000029547 | INTS1 | -0.50472823 | 6.39E-05 | 0.002005173 |
| ENSMUSG00000035967 | INTS6L | -0.487088745 | 6.21E-06 | 3.72E-04 |
| ENSMUSG00000051495 | IRF2BP2 | -0.362298911 | 1.54E-04 | 0.003577454 |
| ENSMUSG00000055980 | IRS1 | -0.379007422 | 6.34E-04 | 0.009273289 |
| ENSMUSG00000051243 | ISLR2 | 1.157271345 | 7.35E-07 | 7.51E-05 |
| ENSMUSG00000025348 | ITGA7 | -0.449155667 | 9.41E-06 | 5.14E-04 |
| ENSMUSG00000026223 | ITM2C | 0.354152203 | 6.91E-04 | 0.009818627 |
| ENSMUSG00000062785 | KCNC3 | -0.425787588 | 2.19E-04 | 0.00448912 |
| ENSMUSG00000036760 | KCNK9 | -0.681023875 | 1.45E-06 | 1.28E-04 |
| ENSMUSG00000101609 | KCNQ1OT1 | -0.653196081 | 6.52E-04 | 0.00944569 |
| ENSMUSG00000016346 | KCNQ2 | -0.461549573 | 8.51E-05 | 0.002352671 |
| ENSMUSG00000058740 | KCNT1 | -0.660382623 | 3.24E-07 | 4.20E-05 |
| ENSMUSG00000018476 | KDM6B | -0.438192614 | 2.21E-04 | 0.004517336 |
| ENSMUSG00000028060 | KHDC4 | -0.474447784 | 9.58E-08 | 1.60E-05 |
| ENSMUSG00000047153 | KHNYN | -0.390412319 | 3.15E-04 | 0.00582063 |
| ENSMUSG00000007670 | KHSRP | -0.597270188 | 6.99E-08 | 1.22E-05 |
| ENSMUSG00000031824 | KIAA0513 | -0.395655775 | 8.07E-05 | 0.002312173 |

| | | | | |
|---|---|---|---|---|
| ENSMUSG00000107877 | KIAA0753 | -0.808749847 | 6.09E-04 | 0.009039363 |
| ENSMUSG00000060012 | KIF13B | -0.324677014 | 4.40E-04 | 0.007366527 |
| ENSMUSG00000041642 | KIF21B | -0.404918267 | 7.02E-04 | 0.009949925 |
| ENSMUSG00000036915 | KIRREL2 | 0.737385878 | 6.70E-05 | 0.002086415 |
| ENSMUSG00000002028 | KMT2A | -0.534514917 | 2.47E-04 | 0.004887592 |
| ENSMUSG00000006307 | KMT2B | -0.534599756 | 2.00E-05 | 8.70E-04 |
| ENSMUSG00000048154 | KMT2D | -0.82577278 | 3.16E-07 | 4.17E-05 |
| ENSMUSG00000029004 | KMT2E | -0.511893957 | 1.04E-04 | 0.002724288 |
| ENSMUSG00000059851 | KMT5C | -0.677103833 | 6.51E-07 | 7.00E-05 |
| ENSMUSG00000066129 | KNDC1 | -0.483850765 | 5.91E-05 | 0.001896804 |
| ENSMUSG00000042810 | KRBA1 | -0.497690875 | 2.59E-06 | 1.98E-04 |
| ENSMUSG00000015647 | LAMA5 | -0.755811003 | 1.54E-06 | 1.34E-04 |
| ENSMUSG00000002900 | LAMB1 | -0.509303025 | 6.91E-06 | 3.99E-04 |
| ENSMUSG00000030842 | LAMTOR1 | 0.356324897 | 4.92E-04 | 0.007936394 |
| ENSMUSG00000037331 | LARP1 | -0.420948212 | 4.20E-04 | 0.007142516 |
| ENSMUSG00000037295 | LDLRAP1 | -0.443253179 | 5.09E-04 | 0.008106899 |
| ENSMUSG00000035545 | LENG8 | -1.163784799 | 1.65E-14 | 4.14E-11 |
| ENSMUSG00000018698 | LHX1 | -0.619385577 | 2.02E-06 | 1.67E-04 |
| ENSMUSG00000024781 | LIPA | 0.425701638 | 5.06E-04 | 0.008068595 |
| ENSMUSG00000062044 | LMTK3 | -0.596042775 | 4.61E-06 | 3.06E-04 |
| ENSMUSG00000018451 | LOC728392 | 0.69750986 | 1.10E-07 | 1.77E-05 |
| ENSMUSG00000038668 | LPAR1 | 0.50259961 | 8.43E-05 | 0.002341404 |
| ENSMUSG00000040249 | LRP1 | -0.535695638 | 6.84E-04 | 0.009777308 |
| ENSMUSG00000025145 | LRRC45 | -0.873927081 | 7.28E-11 | 4.21E-08 |
| ENSMUSG00000002020 | LTBP2 | -0.926668419 | 5.70E-09 | 1.65E-06 |
| ENSMUSG00000040488 | LTBP4 | -0.530918696 | 1.10E-06 | 9.97E-05 |
| ENSMUSG00000024188 | LUC7L | -0.330724234 | 1.41E-04 | 0.003373398 |
| ENSMUSG00000020863 | LUC7L3 | -0.502251308 | 3.95E-06 | 2.75E-04 |
| ENSMUSG00000001089 | LUZP1 | -0.464202741 | 1.16E-05 | 6.01E-04 |
| ENSMUSG00000069516 | LYZ | 0.776027264 | 4.79E-05 | 0.001635798 |
| ENSMUSG00000051510 | MAFG | -0.517937815 | 4.54E-07 | 5.50E-05 |
| ENSMUSG00000045095 | MAGI1 | -0.402201412 | 4.50E-05 | 0.001566513 |
| ENSMUSG00000027375 | MAL | 0.526908697 | 8.35E-05 | 0.002339439 |
| ENSMUSG00000026941 | MAMDC4 | -0.834202308 | 3.60E-05 | 0.001319803 |
| ENSMUSG00000059401 | Mamld1 | -0.489466669 | 4.16E-06 | 2.87E-04 |
| ENSMUSG00000032295 | MAN2C1 | -0.390868675 | 9.66E-05 | 0.002576131 |
| ENSMUSG00000032575 | MANF | 0.327732609 | 5.73E-04 | 0.008734236 |
| ENSMUSG00000052727 | MAP1B | -0.563802805 | 3.87E-05 | 0.001392878 |
| ENSMUSG00000033618 | MAP3K13 | -0.617736605 | 6.00E-06 | 3.62E-04 |
| ENSMUSG00000020700 | MAP3K3 | -0.40071906 | 5.76E-04 | 0.008756534 |
| ENSMUSG00000042724 | MAP3K9 | -0.513535539 | 5.22E-06 | 3.27E-04 |
| ENSMUSG00000024948 | MAP4K2 | -0.468940391 | 3.12E-06 | 2.28E-04 |
| ENSMUSG00000019996 | MAP7 | -0.380554386 | 3.03E-04 | 0.005643502 |
| ENSMUSG00000053137 | MAPK11 | -0.538771916 | 2.54E-05 | 0.001038466 |
| ENSMUSG00000001034 | MAPK7 | -0.546347089 | 8.97E-05 | 0.002438907 |

| | | | | |
|---|---|---|---|---|
| ENSMUSG00000033902 | MAPKBP1 | -0.516134143 | 4.72E-06 | 3.07E-04 |
| ENSMUSG00000024969 | MARK2 | -0.386722232 | 4.91E-04 | 0.007936394 |
| ENSMUSG00000053693 | MAST1 | -0.443572066 | 2.42E-04 | 0.004839485 |
| ENSMUSG00000004933 | MATK | -0.563645669 | 2.08E-04 | 0.004322789 |
| ENSMUSG00000024561 | MBD1 | -0.371467533 | 2.11E-04 | 0.004375319 |
| ENSMUSG00000025409 | MBD6 | -0.593714406 | 1.20E-05 | 6.14E-04 |
| ENSMUSG00000041607 | MBP | 0.7901779 | 5.20E-07 | 6.11E-05 |
| ENSMUSG00000001150 | MCM3AP | -0.324273429 | 4.18E-04 | 0.007124692 |
| ENSMUSG00000004567 | MCOLN1 | -0.344680622 | 4.06E-04 | 0.006963461 |
| ENSMUSG00000061607 | MDC1 | -0.423580002 | 4.05E-04 | 0.006963461 |
| ENSMUSG00000054387 | MDM4 | -0.458180236 | 1.15E-05 | 6.01E-04 |
| ENSMUSG00000002768 | MEA1 | 0.385263168 | 3.00E-04 | 0.00560938 |
| ENSMUSG00000079487 | MED12 | -0.682031937 | 1.75E-09 | 5.60E-07 |
| ENSMUSG00000012114 | MED15 | -0.342145993 | 2.13E-04 | 0.004395305 |
| ENSMUSG00000017210 | MED24 | -0.421961135 | 2.69E-04 | 0.005195082 |
| ENSMUSG00000015804 | MED28 | 0.35079631 | 3.14E-04 | 0.005811328 |
| ENSMUSG00000001419 | MEF2D | -0.421980028 | 1.10E-04 | 0.002812978 |
| ENSMUSG00000021268 | Meg3 | -1.130852132 | 5.50E-15 | 1.65E-11 |
| ENSMUSG00000024593 | MEGF10 | -0.403971542 | 1.72E-04 | 0.003824447 |
| ENSMUSG00000004896 | METTL25B | -0.470869694 | 9.98E-06 | 5.38E-04 |
| ENSMUSG00000054619 | METTL7A | 0.377590895 | 3.74E-04 | 0.006609229 |
| ENSMUSG00000001082 | MFSD10 | -0.576104281 | 2.67E-04 | 0.005183705 |
| ENSMUSG00000008540 | MGST1 | 0.494142249 | 4.97E-04 | 0.007971569 |
| ENSMUSG00000029060 | MIB2 | -0.397412164 | 1.42E-04 | 0.003395138 |
| ENSMUSG00000038244 | MICAL2 | -0.46567515 | 5.15E-05 | 0.001723063 |
| ENSMUSG00000051586 | MICAL3 | -0.427117845 | 5.80E-04 | 0.008786366 |
| ENSMUSG00000049760 | MICOS13 | 0.468955958 | 6.82E-05 | 0.002111064 |
| ENSMUSG00000035299 | MID1 | -0.64506517 | 8.37E-05 | 0.002339439 |
| ENSMUSG00000002580 | MIEN1 | 0.500230693 | 2.78E-05 | 0.001117851 |
| ENSMUSG00000074415 | Mir100hg | -0.554345474 | 2.45E-04 | 0.004881538 |
| ENSMUSG00000097545 | Mir124a-1hg | -0.903989028 | 5.00E-12 | 5.86E-09 |
| ENSMUSG00000092981 | Mir5125 | -1.159062879 | 5.21E-08 | 9.81E-06 |
| ENSMUSG00000097391 | Mirg | -0.986529017 | 2.17E-11 | 1.53E-08 |
| ENSMUSG00000074217 | Misp3 | 1.265462388 | 1.67E-05 | 7.71E-04 |
| ENSMUSG00000038342 | MLXIP | -0.505155731 | 3.25E-05 | 0.001233497 |
| ENSMUSG00000039533 | MMD2 | 0.369161761 | 1.39E-04 | 0.003346496 |
| ENSMUSG00000032517 | MOBP | 0.88970706 | 9.43E-09 | 2.49E-06 |
| ENSMUSG00000023861 | MPC1 | 0.48588792 | 8.20E-05 | 0.002331043 |
| ENSMUSG00000026568 | MPC2 | 0.396336148 | 4.56E-04 | 0.007560543 |
| ENSMUSG00000071711 | MPST | 0.447665918 | 1.74E-04 | 0.003843466 |
| ENSMUSG00000022558 | MROH1 | -0.498632533 | 3.21E-05 | 0.001230469 |
| ENSMUSG00000022370 | MRPL13 | 0.502799313 | 1.16E-04 | 0.002949472 |
| ENSMUSG00000030879 | MRPL17 | 0.427296882 | 1.69E-04 | 0.003773434 |
| ENSMUSG00000030612 | MRPL46 | 0.490012659 | 2.03E-04 | 0.004261143 |
| ENSMUSG00000034932 | MRPL54 | 0.505874672 | 4.15E-04 | 0.007091696 |

| | | | | |
|---|---|---|---|---|
| ENSMUSG00000049960 | MRPS16 | 0.465695075 | 2.51E-04 | 0.004939019 |
| ENSMUSG00000029918 | MRPS33 | 0.438791087 | 3.00E-04 | 0.00560938 |
| ENSMUSG00000009569 | MRTFB | -0.376984413 | 1.54E-04 | 0.003585467 |
| ENSMUSG00000007035 | MSH5 | -0.896442401 | 3.57E-05 | 0.001313702 |
| ENSMUSG00000031604 | MSMO1 | 0.493031926 | 2.55E-06 | 1.97E-04 |
| ENSMUSG00000031765 | Mt1 | 0.504980971 | 1.44E-04 | 0.003423779 |
| ENSMUSG00000031760 | Mt3 | 0.519541789 | 8.66E-07 | 8.40E-05 |
| ENSMUSG00000073481 | MTARC2 | 0.360246493 | 6.11E-04 | 0.009064327 |
| ENSMUSG00000052105 | MTCL1 | -0.548720217 | 4.15E-05 | 0.001475659 |
| ENSMUSG00000020900 | MYH10 | -0.551034432 | 1.51E-05 | 7.15E-04 |
| ENSMUSG00000074652 | MYH7B | -0.542293517 | 1.06E-05 | 5.66E-04 |
| ENSMUSG00000022443 | MYH9 | -0.495287806 | 5.81E-06 | 3.54E-04 |
| ENSMUSG00000042678 | MYO15A | -1.489582056 | 7.20E-07 | 7.41E-05 |
| ENSMUSG00000020527 | MYO19 | -0.733746553 | 2.01E-04 | 0.004246312 |
| ENSMUSG00000004677 | MYO9B | -0.486501672 | 3.03E-05 | 0.001185893 |
| ENSMUSG00000031652 | N4BP1 | -0.363902154 | 5.36E-04 | 0.008406521 |
| ENSMUSG00000024764 | NAA40 | -0.514020659 | 1.05E-05 | 5.59E-04 |
| ENSMUSG00000025402 | NAB2 | -0.59230126 | 1.19E-09 | 4.25E-07 |
| ENSMUSG00000061315 | NACA | 0.384621402 | 4.57E-04 | 0.007565952 |
| ENSMUSG00000025588 | NAT2 | 0.845826658 | 6.74E-04 | 0.009665507 |
| ENSMUSG00000052512 | NAV2 | -0.563492101 | 7.48E-07 | 7.60E-05 |
| ENSMUSG00000031505 | NAXD | 0.371778064 | 5.15E-04 | 0.008199597 |
| ENSMUSG00000056724 | NBEAL2 | -0.645422895 | 4.40E-04 | 0.007366527 |
| ENSMUSG00000038252 | NCAPD2 | -0.449277397 | 3.89E-04 | 0.006816387 |
| ENSMUSG00000020647 | NCOA1 | -0.373691313 | 6.47E-04 | 0.009411581 |
| ENSMUSG00000027678 | NCOA3 | -0.435808548 | 4.72E-04 | 0.007728931 |
| ENSMUSG00000038369 | Ncoa6 | -0.491991293 | 2.80E-05 | 0.001120716 |
| ENSMUSG00000004558 | NDRG2 | 0.544159531 | 2.69E-07 | 3.64E-05 |
| ENSMUSG00000027971 | NDST4 | 1.506831364 | 1.32E-05 | 6.57E-04 |
| ENSMUSG00000014294 | NDUFA2 | 0.447674476 | 4.22E-05 | 0.001495434 |
| ENSMUSG00000023089 | NDUFA5 | 0.340830997 | 6.28E-04 | 0.00921093 |
| ENSMUSG00000041881 | NDUFA7 | 0.467168562 | 4.86E-04 | 0.00787775 |
| ENSMUSG00000026032 | NDUFB3 | 0.356168409 | 6.94E-04 | 0.009853873 |
| ENSMUSG00000033938 | NDUFB7 | 0.381352889 | 5.60E-04 | 0.0086356 |
| ENSMUSG00000037152 | NDUFC1 | 0.389439851 | 3.90E-04 | 0.006820396 |
| ENSMUSG00000081824 | Ndufs5-ps | 0.437988657 | 1.75E-04 | 0.003861763 |
| ENSMUSG00000021606 | NDUFS6 | 0.411526075 | 5.21E-05 | 0.001737376 |
| ENSMUSG00000003847 | NFAT5 | -0.57696926 | 1.97E-05 | 8.62E-04 |
| ENSMUSG00000027574 | NKAIN4 | 0.542849062 | 3.81E-07 | 4.85E-05 |
| ENSMUSG00000032525 | NKTR | -0.579470648 | 1.43E-07 | 2.22E-05 |
| ENSMUSG00000051790 | NLGN2 | -0.423374871 | 4.96E-04 | 0.007958452 |
| ENSMUSG00000067786 | NNAT | 0.621776259 | 5.22E-08 | 9.81E-06 |
| ENSMUSG00000095567 | NOC2L | -0.351929424 | 1.62E-04 | 0.003681282 |
| ENSMUSG00000015468 | NOTCH4 | -0.626448884 | 4.34E-04 | 0.007300218 |
| ENSMUSG00000020447 | NPC1L1 | -1.219041292 | 4.83E-04 | 0.007851613 |

| | | | | |
|---|---|---|---|---|
| ENSMUSG00000021242 | NPC2 | 0.394010572 | 2.63E-04 | 0.005114705 |
| ENSMUSG00000028469 | NPR2 | -0.487212229 | 1.90E-05 | 8.50E-04 |
| ENSMUSG00000059991 | NPTX2 | 1.424186686 | 3.24E-08 | 6.68E-06 |
| ENSMUSG00000029819 | NPY | 0.534745134 | 5.61E-04 | 0.0086356 |
| ENSMUSG00000026826 | NR4A2 | -1.507736907 | 4.05E-04 | 0.006963461 |
| ENSMUSG00000075590 | Nrbp2 | -0.45779543 | 2.11E-04 | 0.004375319 |
| ENSMUSG00000053310 | Nrgn | 1.679410496 | 7.04E-08 | 1.22E-05 |
| ENSMUSG00000048978 | NRSN1 | 0.521896453 | 7.00E-07 | 7.31E-05 |
| ENSMUSG00000021488 | NSD1 | -0.466099454 | 4.61E-05 | 0.00159614 |
| ENSMUSG00000030750 | NSMCE1 | 0.54513627 | 5.67E-04 | 0.008689413 |
| ENSMUSG00000020032 | NUAK1 | -0.43322405 | 2.99E-04 | 0.005608807 |
| ENSMUSG00000037857 | NUFIP2 | -0.379074955 | 6.06E-04 | 0.009023781 |
| ENSMUSG00000066306 | NUMA1 | -0.557363068 | 1.94E-05 | 8.56E-04 |
| ENSMUSG00000030091 | NUP210 | -0.464547279 | 3.56E-04 | 0.006356061 |
| ENSMUSG00000090061 | NWD2 | -0.461504392 | 2.18E-04 | 0.004472854 |
| ENSMUSG00000010097 | NXF1 | -0.504645669 | 7.88E-07 | 7.89E-05 |
| ENSMUSG00000011179 | ODC1 | 0.414281415 | 6.88E-04 | 0.009816849 |
| ENSMUSG00000026790 | ODF2 | -0.710995498 | 6.24E-09 | 1.77E-06 |
| ENSMUSG00000034160 | OGT | -0.512133149 | 7.50E-06 | 4.27E-04 |
| ENSMUSG00000029822 | OSBPL3 | -0.612377704 | 3.34E-05 | 0.001254219 |
| ENSMUSG00000036990 | OTUD4 | -0.457184946 | 1.04E-04 | 0.002724288 |
| ENSMUSG00000033510 | OTUD7A | -0.350901073 | 1.82E-04 | 0.00396324 |
| ENSMUSG00000039670 | OXLD1 | 0.641776155 | 6.17E-04 | 0.009097627 |
| ENSMUSG00000023191 | P3H3 | -0.441273408 | 1.03E-04 | 0.002724288 |
| ENSMUSG00000022194 | PABPN1 | -0.83656304 | 1.33E-09 | 4.53E-07 |
| ENSMUSG00000040276 | PACSIN1 | -0.432512497 | 5.02E-04 | 0.008026451 |
| ENSMUSG00000074923 | PAK6 | 0.97849939 | 1.67E-04 | 0.003746092 |
| ENSMUSG00000005682 | PAN2 | -0.573750714 | 2.46E-06 | 1.93E-04 |
| ENSMUSG00000029647 | PAN3 | -0.416097069 | 1.87E-04 | 0.004041701 |
| ENSMUSG00000054509 | PARP4 | -0.49018704 | 6.91E-04 | 0.009818627 |
| ENSMUSG00000022974 | PAXBP1 | -0.482686381 | 7.37E-06 | 4.21E-04 |
| ENSMUSG00000021496 | PCBD2 | 0.671850703 | 8.69E-05 | 0.002386118 |
| ENSMUSG00000032527 | PCCB | 0.318961751 | 5.45E-04 | 0.008495291 |
| ENSMUSG00000018537 | PCGF2 | -0.600716931 | 3.58E-06 | 2.54E-04 |
| ENSMUSG00000025050 | PCGF6 | -0.526315105 | 8.27E-05 | 0.002331043 |
| ENSMUSG00000001151 | Pcnt | -0.491950878 | 1.68E-04 | 0.003753686 |
| ENSMUSG00000054874 | PCNX3 | -0.505180906 | 2.24E-05 | 9.40E-04 |
| ENSMUSG00000021576 | PDCD6 | 0.378400531 | 4.64E-04 | 0.007638215 |
| ENSMUSG00000004347 | PDE1C | -0.481178583 | 1.18E-04 | 0.002984889 |
| ENSMUSG00000032006 | PDGFD | 0.482623558 | 3.81E-04 | 0.006717965 |
| ENSMUSG00000022090 | PDLIM2 | 0.748279045 | 2.38E-05 | 9.81E-04 |
| ENSMUSG00000074305 | PEAK1 | -0.476516876 | 5.51E-05 | 0.001820873 |
| ENSMUSG00000002265 | PEG3 | -0.69571567 | 3.64E-09 | 1.07E-06 |
| ENSMUSG00000020893 | PER1 | -0.47657269 | 6.67E-04 | 0.009588458 |
| ENSMUSG00000028957 | PER3 | -0.407397625 | 2.76E-04 | 0.005276112 |

| | | | | |
|---|---|---|---|---|
| ENSMUSG00000024346 | PFDN1 | 0.378283044 | 8.79E-05 | 0.002403913 |
| ENSMUSG00000001289 | PFDN5 | 0.479404762 | 6.13E-05 | 0.001941247 |
| ENSMUSG00000006373 | PGRMC1 | 0.347449294 | 6.45E-04 | 0.00938975 |
| ENSMUSG00000040669 | PHC1 | -0.374333387 | 4.61E-04 | 0.007603814 |
| ENSMUSG00000037791 | PHF12 | -0.362696179 | 3.02E-04 | 0.005627503 |
| ENSMUSG00000016624 | PHF21B | -0.510895597 | 7.79E-05 | 0.002292711 |
| ENSMUSG00000086889 | Phf2os1 | -1.114726784 | 4.82E-05 | 0.001638196 |
| ENSMUSG00000026664 | PHYH | 0.430403046 | 2.87E-05 | 0.001138765 |
| ENSMUSG00000022940 | PIGP | 0.40967471 | 3.83E-04 | 0.006738695 |
| ENSMUSG00000032855 | PKD1 | -0.448009981 | 2.73E-04 | 0.005257639 |
| ENSMUSG00000023913 | PLA2G7 | 0.357948426 | 2.88E-04 | 0.005451446 |
| ENSMUSG00000060675 | PLAAT3 | 0.421044282 | 5.68E-04 | 0.008691174 |
| ENSMUSG00000016933 | PLCG1 | -0.429184258 | 8.70E-05 | 0.002386118 |
| ENSMUSG00000022565 | PLEC | -0.766618488 | 6.79E-08 | 1.20E-05 |
| ENSMUSG00000040268 | PLEKHA1 | -0.410286659 | 6.53E-04 | 0.009454557 |
| ENSMUSG00000030231 | PLEKHA5 | -0.41106993 | 2.27E-05 | 9.43E-04 |
| ENSMUSG00000030701 | PLEKHB1 | 0.596266379 | 6.74E-06 | 3.94E-04 |
| ENSMUSG00000066438 | PLEKHD1 | -0.365904268 | 3.58E-04 | 0.006385514 |
| ENSMUSG00000014782 | PLEKHG4 | -0.980737823 | 4.01E-07 | 4.98E-05 |
| ENSMUSG00000078485 | PLEKHN1 | -0.411783209 | 2.65E-05 | 0.001073006 |
| ENSMUSG00000031775 | PLLP | 0.620545621 | 3.78E-05 | 0.001367114 |
| ENSMUSG00000031398 | PLXNA3 | -0.799389394 | 9.90E-08 | 1.64E-05 |
| ENSMUSG00000029765 | PLXNA4 | -0.427717405 | 6.00E-04 | 0.008946205 |
| ENSMUSG00000028248 | PNISR | -0.759653311 | 3.76E-10 | 1.53E-07 |
| ENSMUSG00000020994 | PNN | -0.623834504 | 5.89E-10 | 2.21E-07 |
| ENSMUSG00000028675 | PNRC2 | 0.340722331 | 1.17E-04 | 0.002984889 |
| ENSMUSG00000038902 | POGZ | -0.330182636 | 5.64E-04 | 0.008651476 |
| ENSMUSG00000039176 | POLG | -0.355641682 | 5.57E-05 | 0.00183749 |
| ENSMUSG00000005198 | POLR2A | -0.453983772 | 4.54E-04 | 0.007549244 |
| ENSMUSG00000033020 | POLR2F | 0.469671412 | 1.89E-04 | 0.004070457 |
| ENSMUSG00000071662 | POLR2G | 0.38387073 | 5.72E-04 | 0.008734236 |
| ENSMUSG00000025280 | POLR3A | -0.432146959 | 3.87E-04 | 0.006790884 |
| ENSMUSG00000026458 | Ppfia4 | -0.477509584 | 9.18E-05 | 0.002477059 |
| ENSMUSG00000032383 | PPIB | 0.4273546 | 5.95E-04 | 0.008911249 |
| ENSMUSG00000062729 | PPOX | -0.468396412 | 8.40E-05 | 0.002339439 |
| ENSMUSG00000039220 | PPP1R10 | -0.377369228 | 1.32E-04 | 0.003241919 |
| ENSMUSG00000061950 | PPP4R1 | -0.370354013 | 6.26E-04 | 0.009188555 |
| ENSMUSG00000036561 | PPP6R2 | -0.372265037 | 4.70E-04 | 0.007719686 |
| ENSMUSG00000055491 | PPRC1 | -0.587913779 | 2.87E-05 | 0.001138765 |
| ENSMUSG00000050271 | PRAG1 | -0.513010335 | 8.07E-07 | 8.04E-05 |
| ENSMUSG00000057637 | PRDM2 | -0.378720251 | 5.91E-04 | 0.008871447 |
| ENSMUSG00000028691 | PRDX1 | 0.389261495 | 6.51E-04 | 0.00944569 |
| ENSMUSG00000005161 | PRDX2 | 0.418215399 | 4.61E-04 | 0.007603814 |
| ENSMUSG00000078816 | PRKCG | -0.547280888 | 2.39E-06 | 1.91E-04 |
| ENSMUSG00000049504 | PROSER1 | -0.612308957 | 5.74E-07 | 6.49E-05 |

| | | | | |
|---|---|---|---|---|
| ENSMUSG00000093629 | Prox2os | -1.175131066 | 1.27E-04 | 0.003157373 |
| ENSMUSG00000027881 | PRPF38B | -0.654217079 | 2.25E-11 | 1.53E-08 |
| ENSMUSG00000035597 | PRPF39 | -0.566648541 | 2.58E-07 | 3.52E-05 |
| ENSMUSG00000023007 | PRPF40B | -0.559347005 | 5.00E-07 | 5.97E-05 |
| ENSMUSG00000021413 | PRPF4B | -0.460201908 | 1.28E-05 | 6.46E-04 |
| ENSMUSG00000020528 | PRPSAP2 | 0.410010212 | 2.51E-04 | 0.004939019 |
| ENSMUSG00000046574 | PRR12 | -0.53984345 | 3.65E-05 | 0.001333099 |
| ENSMUSG00000054280 | PRR14L | -0.35048531 | 5.33E-04 | 0.008385423 |
| ENSMUSG00000055945 | PRR18 | 0.752212674 | 2.76E-08 | 6.01E-06 |
| ENSMUSG00000040225 | PRRC2C | -0.62150003 | 9.57E-07 | 8.99E-05 |
| ENSMUSG00000015476 | Prrt1 | -0.480607382 | 2.26E-04 | 0.00458579 |
| ENSMUSG00000021792 | PRXL2A | 0.421676113 | 3.40E-05 | 0.001261965 |
| ENSMUSG00000029059 | PRXL2B | 0.372125421 | 3.39E-04 | 0.006136383 |
| ENSMUSG00000024640 | PSAT1 | 0.396337403 | 3.00E-04 | 0.00560938 |
| ENSMUSG00000096727 | PSMB9 | 0.946486386 | 1.50E-04 | 0.003529657 |
| ENSMUSG00000026914 | PSMD14 | 0.341216061 | 3.94E-04 | 0.006854783 |
| ENSMUSG00000072946 | PTGR2 | 0.406299543 | 1.32E-05 | 6.57E-04 |
| ENSMUSG00000059456 | PTK2B | 1.142511146 | 6.10E-04 | 0.009052864 |
| ENSMUSG00000027843 | PTPN22 | -0.796696448 | 1.10E-06 | 9.97E-05 |
| ENSMUSG00000036057 | PTPN23 | -0.596120608 | 6.15E-06 | 3.70E-04 |
| ENSMUSG00000038764 | PTPN3 | -0.402582451 | 1.47E-04 | 0.003480769 |
| ENSMUSG00000013236 | PTPRS | -0.442796453 | 8.97E-05 | 0.002438907 |
| ENSMUSG00000029528 | PXN | -0.898436043 | 2.75E-15 | 1.03E-11 |
| ENSMUSG00000015806 | QDPR | 0.430782176 | 3.92E-04 | 0.006837134 |
| ENSMUSG00000056211 | R3HDM1 | -0.417836582 | 7.86E-05 | 0.002299423 |
| ENSMUSG00000079316 | RAB9A | 0.442168917 | 1.34E-04 | 0.003269649 |
| ENSMUSG00000034353 | RAMP1 | 0.649117547 | 2.69E-06 | 2.04E-04 |
| ENSMUSG00000037415 | RANBP10 | -0.430264559 | 1.57E-05 | 7.36E-04 |
| ENSMUSG00000009281 | RARRES2 | 0.590462169 | 1.79E-04 | 0.003904244 |
| ENSMUSG00000010608 | Rbm25 | -0.583278166 | 2.39E-07 | 3.33E-05 |
| ENSMUSG00000029701 | RBM28 | -0.359272311 | 4.26E-04 | 0.00719468 |
| ENSMUSG00000048271 | Rbm33 | -0.702593355 | 2.70E-10 | 1.19E-07 |
| ENSMUSG00000032580 | RBM5 | -0.434046168 | 3.35E-05 | 0.001254219 |
| ENSMUSG00000032582 | RBM6 | -0.348013202 | 4.85E-04 | 0.007869268 |
| ENSMUSG00000052751 | REPIN1 | -0.410317133 | 8.50E-05 | 0.002352671 |
| ENSMUSG00000039852 | RERE | -0.582938654 | 1.07E-05 | 5.71E-04 |
| ENSMUSG00000030222 | RERG | 0.898037146 | 3.94E-04 | 0.006855866 |
| ENSMUSG00000047417 | REXO1 | -0.384098654 | 1.31E-04 | 0.003231462 |
| ENSMUSG00000031706 | RFX1 | -0.679223535 | 2.20E-06 | 1.79E-04 |
| ENSMUSG00000022018 | RGCC | 0.608794508 | 7.05E-05 | 0.002140958 |
| ENSMUSG00000041354 | RGL2 | -0.583781702 | 6.09E-07 | 6.78E-05 |
| ENSMUSG00000022323 | RIDA | 0.414269189 | 3.20E-05 | 0.001230469 |
| ENSMUSG00000041670 | RIMS1 | -0.440053721 | 1.62E-05 | 7.55E-04 |
| ENSMUSG00000038604 | RIPOR1 | -0.347023892 | 3.78E-04 | 0.0066699 |
| ENSMUSG00000001313 | RND2 | 0.487899979 | 6.33E-06 | 3.76E-04 |

| | | | | |
|---|---|---|---|---|
| ENSMUSG00000010086 | RNF112 | -0.518190915 | 2.10E-05 | 8.91E-04 |
| ENSMUSG00000051234 | RNF7 | 0.42606487 | 1.99E-04 | 0.004214777 |
| ENSMUSG00000027981 | RNPC3 | -0.605198488 | 2.07E-04 | 0.004313494 |
| ENSMUSG00000067847 | ROMO1 | 0.429386052 | 3.38E-04 | 0.006129603 |
| ENSMUSG00000110928 | RP23-114G13.1 | -1.39499934 | 4.96E-06 | 3.20E-04 |
| ENSMUSG00000112505 | RP23-118L13.6 | 0.833289988 | 4.63E-04 | 0.00762794 |
| ENSMUSG00000111619 | RP23-319C8.5 | -1.041003919 | 4.11E-07 | 5.06E-05 |
| ENSMUSG00000111394 | RP23-320D23.6 | -1.037015209 | 1.57E-07 | 2.34E-05 |
| ENSMUSG00000111496 | RP23-331E5.10 | -1.384473609 | 1.81E-05 | 8.21E-04 |
| ENSMUSG00000111837 | RP23-361B11.3 | -1.122561034 | 2.22E-06 | 1.80E-04 |
| ENSMUSG00000034032 | RPAP1 | -0.379837189 | 3.07E-04 | 0.005696385 |
| ENSMUSG00000037805 | RPL10A | 0.399237723 | 2.45E-04 | 0.004881538 |
| ENSMUSG00000059291 | RPL11 | 0.462813191 | 1.18E-04 | 0.002991246 |
| ENSMUSG00000038900 | RPL12 | 0.474571245 | 1.28E-04 | 0.003171033 |
| ENSMUSG00000000740 | RPL13 | 0.426597534 | 1.77E-04 | 0.00389201 |
| ENSMUSG00000062328 | RPL17 | 0.490886284 | 1.79E-04 | 0.003904244 |
| ENSMUSG00000045128 | RPL18A | 0.401790271 | 1.37E-04 | 0.003318949 |
| ENSMUSG00000017404 | RPL19 | 0.471249462 | 1.04E-04 | 0.002724288 |
| ENSMUSG00000041453 | RPL21 | 0.360573223 | 3.47E-04 | 0.006240871 |
| ENSMUSG00000058546 | Rpl23a | 0.426685744 | 1.79E-04 | 0.003904244 |
| ENSMUSG00000060938 | RPL26 | 0.439239536 | 1.07E-04 | 0.002779316 |
| ENSMUSG00000046364 | RPL27A | 0.461161506 | 8.01E-05 | 0.002312173 |
| ENSMUSG00000060036 | RPL3 | 0.369067695 | 5.42E-04 | 0.008463273 |
| ENSMUSG00000084349 | Rpl3-ps1 | 0.396476532 | 1.41E-04 | 0.003367 |
| ENSMUSG00000058600 | RPL30 | 0.383542693 | 5.61E-05 | 0.001845577 |
| ENSMUSG00000073702 | RPL31 | 0.497955874 | 1.23E-05 | 6.26E-04 |
| ENSMUSG00000057841 | Rpl32 | 0.422689378 | 2.96E-04 | 0.005567282 |
| ENSMUSG00000062997 | RPL35 | 0.45193863 | 1.96E-04 | 0.004168127 |
| ENSMUSG00000060636 | RPL35A | 0.413583567 | 4.75E-04 | 0.007755082 |
| ENSMUSG00000057863 | Rpl36 | 0.452171538 | 1.65E-04 | 0.003719893 |
| ENSMUSG00000079435 | Rpl36a | 0.440921674 | 1.52E-05 | 7.16E-04 |
| ENSMUSG00000041841 | RPL37 | 0.403887271 | 1.37E-04 | 0.003318949 |
| ENSMUSG00000046330 | RPL37A | 0.482639424 | 2.77E-05 | 0.001117851 |
| ENSMUSG00000057322 | RPL38 | 0.408329563 | 1.52E-04 | 0.003549111 |
| ENSMUSG00000079641 | RPL39 | 0.481887748 | 1.63E-04 | 0.003691526 |
| ENSMUSG00000058558 | RPL5 | 0.462877964 | 7.28E-05 | 0.002190239 |
| ENSMUSG00000029614 | RPL6 | 0.359799751 | 5.41E-04 | 0.008463273 |
| ENSMUSG00000043716 | RPL7 | 0.414535699 | 2.44E-04 | 0.004877454 |
| ENSMUSG00000003970 | RPL8 | 0.390351361 | 5.85E-04 | 0.008802883 |
| ENSMUSG00000047215 | RPL9 | 0.49911792 | 8.24E-05 | 0.002331043 |
| ENSMUSG00000067274 | RPLP0 | 0.393112913 | 4.23E-04 | 0.00716837 |
| ENSMUSG00000007892 | Rplp1 (includes otl | 0.43490354 | 2.13E-04 | 0.004402486 |
| ENSMUSG00000025508 | RPLP2 | 0.390633175 | 3.19E-04 | 0.005875976 |
| ENSMUSG00000003429 | RPS11 | 0.523042792 | 1.80E-05 | 8.21E-04 |
| ENSMUSG00000061983 | RPS12 | 0.463551224 | 2.04E-04 | 0.004278493 |

| | | | | |
|---|---|---|---|---|
| ENSMUSG00000090862 | RPS13 | 0.452951135 | 1.36E-04 | 0.003316671 |
| ENSMUSG00000024608 | RPS14 | 0.495052773 | 1.40E-04 | 0.003356404 |
| ENSMUSG00000063457 | RPS15 | 0.462343499 | 1.32E-04 | 0.003241757 |
| ENSMUSG00000037563 | RPS16 | 0.516848147 | 5.87E-05 | 0.001894458 |
| ENSMUSG00000061787 | RPS17 | 0.407485114 | 2.29E-04 | 0.004640632 |
| ENSMUSG00000008668 | RPS18 | 0.506296789 | 5.62E-05 | 0.001845577 |
| ENSMUSG00000069117 | Rps18-ps6 | 0.526711165 | 2.05E-05 | 8.80E-04 |
| ENSMUSG00000040952 | RPS19 | 0.432904148 | 1.68E-04 | 0.003757229 |
| ENSMUSG00000028234 | RPS20 | 0.444233254 | 6.94E-05 | 0.00212573 |
| ENSMUSG00000039001 | RPS21 | 0.490517462 | 5.69E-05 | 0.001855367 |
| ENSMUSG00000049517 | RPS23 | 0.460406648 | 3.80E-05 | 0.001370509 |
| ENSMUSG00000025290 | RPS24 | 0.481295315 | 7.01E-05 | 0.002140958 |
| ENSMUSG00000009927 | RPS25 | 0.441897922 | 4.22E-04 | 0.007168328 |
| ENSMUSG00000025362 | RPS26 | 0.361889217 | 5.01E-04 | 0.008024202 |
| ENSMUSG00000090733 | Rps27/Rps27rt | 0.589445989 | 9.28E-06 | 5.09E-04 |
| ENSMUSG00000020460 | RPS27A | 0.486684768 | 2.96E-05 | 0.001169251 |
| ENSMUSG00000067288 | RPS28 | 0.367662476 | 1.60E-04 | 0.003672348 |
| ENSMUSG00000034892 | RPS29 | 0.383796307 | 4.21E-04 | 0.007154441 |
| ENSMUSG00000028081 | Rps3a1 | 0.423915168 | 6.41E-05 | 0.002008845 |
| ENSMUSG00000031320 | RPS4Y1 | 0.372103012 | 4.00E-04 | 0.006921866 |
| ENSMUSG00000012848 | RPS5 | 0.502249645 | 1.11E-05 | 5.84E-04 |
| ENSMUSG00000028495 | RPS6 | 0.421220248 | 1.72E-04 | 0.003824447 |
| ENSMUSG00000024830 | RPS6KB2 | -0.534033923 | 8.43E-07 | 8.28E-05 |
| ENSMUSG00000061477 | RPS7 | 0.496228664 | 7.73E-05 | 0.002284293 |
| ENSMUSG00000095597 | Rps7-ps3 | 0.486302982 | 3.62E-04 | 0.006448168 |
| ENSMUSG00000047675 | RPS8 | 0.432504929 | 8.01E-05 | 0.002312173 |
| ENSMUSG00000006333 | RPS9 | 0.4495313 | 5.09E-05 | 0.001712444 |
| ENSMUSG00000032518 | RPSA | 0.474934942 | 9.38E-05 | 0.002520287 |
| ENSMUSG00000039087 | RREB1 | -0.961736192 | 6.58E-12 | 6.97E-09 |
| ENSMUSG00000037266 | RSRP1 | -0.454797937 | 7.95E-05 | 0.002311564 |
| ENSMUSG00000048617 | RTBDN | -1.200720569 | 5.14E-04 | 0.008184836 |
| ENSMUSG00000021807 | RTRAF | 0.310542344 | 5.68E-04 | 0.008689413 |
| ENSMUSG00000041263 | RUSC1 | -0.354283779 | 1.25E-04 | 0.003116384 |
| ENSMUSG00000030592 | RYR1 | -0.953906246 | 2.05E-10 | 9.65E-08 |
| ENSMUSG00000057378 | RYR3 | -1.241684604 | 1.36E-10 | 6.83E-08 |
| ENSMUSG00000044080 | S100A1 | 0.602261651 | 8.60E-06 | 4.81E-04 |
| ENSMUSG00000074457 | S100A16 | 0.449586981 | 2.63E-04 | 0.005114705 |
| ENSMUSG00000071054 | SAFB | -0.763542626 | 2.19E-09 | 6.85E-07 |
| ENSMUSG00000042625 | SAFB2 | -0.738183539 | 3.95E-10 | 1.56E-07 |
| ENSMUSG00000078427 | SARNP | 0.32879813 | 6.50E-04 | 0.009440091 |
| ENSMUSG00000039148 | SART1 | -0.451791664 | 1.19E-05 | 6.11E-04 |
| ENSMUSG00000085272 | SBK3 | -1.030175322 | 2.74E-04 | 0.005265212 |
| ENSMUSG00000035673 | SBNO2 | -0.467643096 | 4.35E-04 | 0.007301742 |
| ENSMUSG00000046056 | SBSN | -1.188004333 | 3.31E-04 | 0.00602708 |
| ENSMUSG00000022983 | SCAF4 | -0.657357364 | 1.53E-09 | 5.01E-07 |

| | | | | |
|---|---|---|---|---|
| ENSMUSG00000028603 | SCP2 | 0.351358027 | 6.17E-04 | 0.009097627 |
| ENSMUSG00000022568 | SCRIB | -0.362680159 | 1.96E-04 | 0.004168127 |
| ENSMUSG00000002064 | SDF2 | 0.405887009 | 4.51E-04 | 0.007503927 |
| ENSMUSG00000074211 | SDHAF1 | 0.465153606 | 3.45E-04 | 0.006224234 |
| ENSMUSG00000026924 | SEC16A | -0.417650212 | 2.03E-04 | 0.004261143 |
| ENSMUSG00000034473 | SEC22A | 0.456279166 | 4.55E-04 | 0.007549612 |
| ENSMUSG00000037072 | SELENOF | 0.442854324 | 5.52E-04 | 0.008577474 |
| ENSMUSG00000042682 | SELENOK | 0.460549418 | 4.63E-05 | 0.00159614 |
| ENSMUSG00000064373 | SELENOP | 0.609395269 | 2.55E-06 | 1.97E-04 |
| ENSMUSG00000041571 | SELENOW | 0.40884129 | 5.91E-04 | 0.008871447 |
| ENSMUSG00000019647 | SEMA6A | -0.454280758 | 5.45E-04 | 0.008495291 |
| ENSMUSG00000058013 | SEPTIN11 | -0.393117445 | 1.79E-04 | 0.003904244 |
| ENSMUSG00000060147 | SERPINB6 | 0.563179142 | 3.89E-06 | 2.72E-04 |
| ENSMUSG00000026249 | SERPINE2 | 0.490235543 | 1.11E-06 | 9.97E-05 |
| ENSMUSG00000042272 | SESTD1 | -0.347094102 | 6.71E-04 | 0.009620257 |
| ENSMUSG00000042308 | SETD1A | -0.560726595 | 1.76E-04 | 0.003868706 |
| ENSMUSG00000038384 | Setd1b | -0.624275109 | 1.79E-06 | 1.52E-04 |
| ENSMUSG00000015697 | SETDB1 | -0.364642101 | 7.88E-05 | 0.002299423 |
| ENSMUSG00000024949 | Sf1 | -0.483405257 | 4.40E-05 | 0.001540562 |
| ENSMUSG00000078348 | SF3B5 | 0.43626703 | 2.62E-04 | 0.005113055 |
| ENSMUSG00000037361 | SF3B6 | 0.429767855 | 9.42E-05 | 0.002522112 |
| ENSMUSG00000028820 | SFPQ | -0.449109945 | 6.87E-05 | 0.002113182 |
| ENSMUSG00000018822 | SFRP5 | 1.350340099 | 5.94E-05 | 0.001896804 |
| ENSMUSG00000029439 | SFSWAP | -0.644245047 | 2.34E-08 | 5.17E-06 |
| ENSMUSG00000025036 | SFXN2 | -0.712409975 | 2.21E-04 | 0.004511136 |
| ENSMUSG00000042594 | SH2B3 | -0.38385214 | 1.86E-04 | 0.004026077 |
| ENSMUSG00000038884 | SHFL | -0.547681337 | 8.29E-07 | 8.20E-05 |
| ENSMUSG00000041889 | SHISA4 | 0.402584824 | 2.75E-04 | 0.005276112 |
| ENSMUSG00000053930 | SHISA6 | -0.5590801 | 1.91E-04 | 0.00408788 |
| ENSMUSG00000089832 | SHKBP1 | -0.650157131 | 6.08E-04 | 0.009033709 |
| ENSMUSG00000034908 | SIDT2 | -0.349006155 | 6.90E-04 | 0.009818627 |
| ENSMUSG00000029524 | SIRT4 | -0.569926963 | 3.24E-05 | 0.001233497 |
| ENSMUSG00000025138 | SIRT7 | -0.353553244 | 5.73E-04 | 0.008734236 |
| ENSMUSG00000040938 | SLC16A11 | -0.810451148 | 1.16E-11 | 1.03E-08 |
| ENSMUSG00000040414 | SLC25A28 | -0.408356664 | 4.70E-05 | 0.001610025 |
| ENSMUSG00000028982 | SLC25A33 | 0.385914622 | 1.18E-04 | 0.002991246 |
| ENSMUSG00000002346 | SLC25A42 | -0.46357461 | 7.94E-05 | 0.002311564 |
| ENSMUSG00000036298 | SLC2A13 | -0.448467727 | 7.79E-06 | 4.40E-04 |
| ENSMUSG00000085028 | Slc2a4rg-ps | -1.079837274 | 1.03E-07 | 1.68E-05 |
| ENSMUSG00000037089 | SLC35B2 | 0.363516639 | 4.94E-04 | 0.00794112 |
| ENSMUSG00000036949 | SLC39A12 | 0.455301097 | 1.89E-05 | 8.50E-04 |
| ENSMUSG00000072572 | SLC39A2 | -1.101900118 | 5.45E-06 | 3.36E-04 |
| ENSMUSG00000057193 | SLC44A2 | -0.364042595 | 1.56E-04 | 0.003599362 |
| ENSMUSG00000023032 | SLC4A8 | -0.594942984 | 1.32E-04 | 0.003241919 |
| ENSMUSG00000030307 | SLC6A11 | 0.529722858 | 6.51E-07 | 7.00E-05 |

| | | | | |
|---|---|---|---|---|
| ENSMUSG00000030096 | SLC6A6 | -0.431638473 | 3.23E-04 | 0.005919811 |
| ENSMUSG00000014786 | SLC9A5 | -0.460437428 | 4.01E-05 | 0.001430419 |
| ENSMUSG00000036097 | SLF2 | -0.443405593 | 3.75E-05 | 0.001361927 |
| ENSMUSG00000032212 | SLTM | -0.565269077 | 6.30E-07 | 6.91E-05 |
| ENSMUSG00000039738 | SLX4 | -0.404282153 | 4.46E-04 | 0.007450672 |
| ENSMUSG00000032481 | SMARCC1 | -0.377355332 | 6.70E-04 | 0.009620257 |
| ENSMUSG00000022452 | SMDT1 | 0.395797547 | 1.65E-04 | 0.003727904 |
| ENSMUSG00000030655 | SMG1 | -0.479199996 | 5.56E-04 | 0.008602691 |
| ENSMUSG00000044600 | SMIM7 | 0.34404378 | 5.98E-04 | 0.008932034 |
| ENSMUSG00000005899 | SMPD4 | -0.514013512 | 3.20E-06 | 2.31E-04 |
| ENSMUSG00000019872 | SMPDL3A | 0.477764872 | 6.94E-05 | 0.00212573 |
| ENSMUSG00000033419 | SNAP91 | -0.400815737 | 6.77E-04 | 0.009684726 |
| ENSMUSG00000036281 | SNAPC4 | -0.576206574 | 1.18E-06 | 1.05E-04 |
| ENSMUSG00000044349 | Snhg11 | -1.260464708 | 1.60E-21 | 2.41E-17 |
| ENSMUSG00000001158 | SNRNP27 | 0.41172809 | 5.66E-05 | 0.001853646 |
| ENSMUSG00000021431 | SNRNP48 | -0.462355763 | 5.62E-04 | 0.0086356 |
| ENSMUSG00000063511 | SNRNP70 | -0.813177593 | 6.87E-10 | 2.52E-07 |
| ENSMUSG00000028385 | SNX30 | -0.454283676 | 4.05E-04 | 0.006963461 |
| ENSMUSG00000056185 | SNX32 | -0.327548262 | 6.35E-04 | 0.009277164 |
| ENSMUSG00000045314 | SOWAHB | -0.556525272 | 2.07E-05 | 8.83E-04 |
| ENSMUSG00000080316 | Spaca6 | -0.967172788 | 8.28E-12 | 7.78E-09 |
| ENSMUSG00000018593 | SPARC | 0.528242169 | 2.71E-06 | 2.04E-04 |
| ENSMUSG00000021917 | SPCS1 | 0.389986766 | 3.37E-04 | 0.006121287 |
| ENSMUSG00000026207 | SPEG | -0.439625935 | 3.77E-05 | 0.001366668 |
| ENSMUSG00000040761 | SPEN | -0.62886822 | 8.23E-05 | 0.002331043 |
| ENSMUSG00000045671 | SPRED2 | -0.504802675 | 5.45E-05 | 0.001809482 |
| ENSMUSG00000037239 | SPRED3 | -0.541257495 | 1.36E-05 | 6.74E-04 |
| ENSMUSG00000011751 | SPTBN4 | -0.712896964 | 7.39E-08 | 1.26E-05 |
| ENSMUSG00000032621 | SREK1 | -0.458021437 | 7.32E-06 | 4.20E-04 |
| ENSMUSG00000020121 | SRGAP1 | -0.435231171 | 1.51E-04 | 0.003542024 |
| ENSMUSG00000026425 | SRGAP2 | -0.471515877 | 3.15E-05 | 0.00122804 |
| ENSMUSG00000009549 | SRP14 | 0.409647017 | 1.25E-04 | 0.003120886 |
| ENSMUSG00000028809 | Srrm1 | -0.566565581 | 3.27E-07 | 4.20E-05 |
| ENSMUSG00000039218 | Srrm2 | -1.000499997 | 1.46E-11 | 1.22E-08 |
| ENSMUSG00000039860 | Srrm3 | -0.590919769 | 1.42E-05 | 6.88E-04 |
| ENSMUSG00000063919 | SRRM4 | -0.748936227 | 2.83E-08 | 6.07E-06 |
| ENSMUSG00000037364 | SRRT | -0.49035693 | 6.27E-06 | 3.74E-04 |
| ENSMUSG00000018379 | SRSF1 | -0.43099859 | 1.19E-04 | 0.002999505 |
| ENSMUSG00000055436 | SRSF11 | -0.356406818 | 2.30E-04 | 0.004640632 |
| ENSMUSG00000034120 | SRSF2 | -0.396315054 | 3.30E-05 | 0.001247629 |
| ENSMUSG00000021134 | Srsf5 | -0.535395674 | 4.65E-06 | 3.06E-04 |
| ENSMUSG00000016921 | SRSF6 | -0.414113181 | 1.09E-04 | 0.002804334 |
| ENSMUSG00000034616 | SSH3 | -0.832935428 | 3.72E-11 | 2.26E-08 |
| ENSMUSG00000004366 | SST | 1.456242376 | 1.50E-05 | 7.15E-04 |
| ENSMUSG00000040287 | STAC3 | -0.842039275 | 2.53E-04 | 0.004956057 |

| | | | | |
|---|---|---|---|---|
| ENSMUSG00000040033 | STAT2 | -0.449438374 | 1.48E-05 | 7.09E-04 |
| ENSMUSG00000037885 | STK35 | -0.405973549 | 1.44E-04 | 0.003420653 |
| ENSMUSG00000024006 | STK38 | -0.38671402 | 1.95E-04 | 0.004160959 |
| ENSMUSG00000022044 | STMN4 | 0.467525175 | 2.45E-05 | 0.001004776 |
| ENSMUSG00000026915 | STRBP | -0.442376637 | 4.19E-04 | 0.007140925 |
| ENSMUSG00000027522 | STX16 | -0.350263924 | 2.88E-04 | 0.005451446 |
| ENSMUSG00000026797 | STXBP1 | -0.4192487 | 1.55E-04 | 0.003585772 |
| ENSMUSG00000004626 | STXBP2 | -0.67889124 | 2.16E-07 | 3.12E-05 |
| ENSMUSG00000028369 | SVEP1 | -0.449882066 | 1.60E-04 | 0.003672348 |
| ENSMUSG00000037217 | SYN1 | -0.427829036 | 1.52E-04 | 0.003549111 |
| ENSMUSG00000096054 | SYNE1 | -0.714124683 | 4.39E-06 | 2.99E-04 |
| ENSMUSG00000019737 | SYNE4 | -0.81765952 | 3.69E-06 | 2.60E-04 |
| ENSMUSG00000067629 | SYNGAP1 | -0.692906399 | 1.00E-08 | 2.60E-06 |
| ENSMUSG00000056296 | SYNPR | 0.820457953 | 3.16E-11 | 2.07E-08 |
| ENSMUSG00000033253 | SZT2 | -0.641414582 | 6.71E-07 | 7.05E-05 |
| ENSMUSG00000061762 | TAC1 | 2.474405042 | 2.28E-07 | 3.24E-05 |
| ENSMUSG00000059187 | TAFA1 | 1.199006447 | 4.48E-04 | 0.007463532 |
| ENSMUSG00000026547 | TAGLN2 | 0.798208954 | 6.62E-08 | 1.18E-05 |
| ENSMUSG00000023051 | TARBP2 | -0.392327367 | 4.12E-04 | 0.007051437 |
| ENSMUSG00000037410 | TBC1D2B | -0.413361316 | 2.30E-04 | 0.004642246 |
| ENSMUSG00000042043 | TBCA | 0.354571483 | 5.93E-04 | 0.008886666 |
| ENSMUSG00000018604 | TBX3 | -0.801757983 | 2.33E-04 | 0.004693988 |
| ENSMUSG00000024498 | TCERG1 | -0.517727896 | 9.36E-07 | 8.85E-05 |
| ENSMUSG00000024985 | TCF7L2 | 0.837787864 | 5.82E-04 | 0.008787521 |
| ENSMUSG00000024613 | TCOF1 | -0.398141638 | 4.27E-04 | 0.007206888 |
| ENSMUSG00000039461 | TCTA | 0.400519064 | 4.94E-04 | 0.00794112 |
| ENSMUSG00000066621 | TECPR1 | -0.441481817 | 2.98E-04 | 0.005598157 |
| ENSMUSG00000039179 | TEKT5 | 0.946793613 | 1.06E-04 | 0.002762564 |
| ENSMUSG00000024170 | TELO2 | -0.549511226 | 7.63E-06 | 4.33E-04 |
| ENSMUSG00000040943 | TET2 | -0.454673851 | 1.96E-05 | 8.60E-04 |
| ENSMUSG00000034832 | TET3 | -0.50429501 | 4.64E-04 | 0.007638215 |
| ENSMUSG00000021359 | TFAP2A | -0.693844145 | 2.70E-04 | 0.00520685 |
| ENSMUSG00000025927 | TFAP2B | -0.540292677 | 2.01E-05 | 8.70E-04 |
| ENSMUSG00000000134 | TFE3 | -0.378930855 | 8.61E-05 | 0.002370781 |
| ENSMUSG00000089736 | TGFBR3L | -1.104560131 | 1.31E-05 | 6.57E-04 |
| ENSMUSG00000000214 | TH | 1.66435322 | 2.20E-06 | 1.79E-04 |
| ENSMUSG00000022847 | THPO | -0.961380956 | 6.93E-06 | 3.99E-04 |
| ENSMUSG00000002489 | TIAM1 | -0.489103017 | 4.35E-05 | 0.001529777 |
| ENSMUSG00000039016 | TIMM8B | 0.450505069 | 3.71E-04 | 0.006580748 |
| ENSMUSG00000110218 | Tincr | -1.084946888 | 1.17E-08 | 2.89E-06 |
| ENSMUSG00000034771 | TLE2 | -0.527316266 | 7.68E-05 | 0.002271947 |
| ENSMUSG00000052698 | TLN2 | -0.427499838 | 6.40E-04 | 0.009331042 |
| ENSMUSG00000031556 | TM2D2 | 0.398552887 | 5.99E-05 | 0.001907207 |
| ENSMUSG00000078681 | TM2D3 | 0.40910815 | 2.74E-04 | 0.005268004 |
| ENSMUSG00000027800 | TM4SF1 | 0.757891862 | 7.05E-05 | 0.002140958 |

| | | | | |
|---|---|---|---|---|
| ENSMUSG00000036151 | TM6SF2 | -0.977714504 | 5.41E-04 | 0.008463273 |
| ENSMUSG00000091537 | TMA7 | 0.395823621 | 4.60E-04 | 0.007603814 |
| ENSMUSG00000019734 | TMC4 | -1.219965975 | 2.04E-05 | 8.80E-04 |
| ENSMUSG00000052428 | TMCO1 | 0.446172056 | 2.05E-05 | 8.80E-04 |
| ENSMUSG00000032353 | TMED3 | 0.503806351 | 1.61E-04 | 0.00367563 |
| ENSMUSG00000043843 | TMEM145 | -0.474365269 | 6.25E-05 | 0.00196947 |
| ENSMUSG00000021361 | TMEM14C | 0.364480298 | 4.93E-04 | 0.00794112 |
| ENSMUSG00000023367 | TMEM176A | 0.750689424 | 1.61E-04 | 0.003678007 |
| ENSMUSG00000029810 | TMEM176B | 0.465973457 | 2.17E-04 | 0.004458552 |
| ENSMUSG00000040883 | TMEM205 | 0.472696012 | 5.30E-04 | 0.008362936 |
| ENSMUSG00000004945 | TMEM242 | 0.538766156 | 3.43E-06 | 2.46E-04 |
| ENSMUSG00000070394 | TMEM256 | 0.537887377 | 1.27E-04 | 0.003157373 |
| ENSMUSG00000028822 | TMEM50A | 0.413547782 | 1.70E-05 | 7.83E-04 |
| ENSMUSG00000045282 | TMEM86B | -0.956268937 | 1.42E-07 | 2.22E-05 |
| ENSMUSG00000020747 | TMEM94 | -0.469084325 | 1.36E-04 | 0.003316671 |
| ENSMUSG00000037278 | TMEM97 | 0.578152856 | 1.15E-04 | 0.002939236 |
| ENSMUSG00000049555 | TMIE | 0.676844312 | 2.28E-04 | 0.00462683 |
| ENSMUSG00000005628 | TMOD4 | -0.921853009 | 3.70E-04 | 0.006580444 |
| ENSMUSG00000049775 | Tmsb4x (includes | 0.510070129 | 1.91E-05 | 8.50E-04 |
| ENSMUSG00000027692 | TNIK | -0.531015897 | 4.34E-06 | 2.96E-04 |
| ENSMUSG00000015829 | TNR | -0.639382664 | 5.00E-08 | 9.64E-06 |
| ENSMUSG00000052707 | TNRC6A | -0.446527555 | 4.65E-05 | 0.00159614 |
| ENSMUSG00000025571 | TNRC6C | -0.489205754 | 1.61E-04 | 0.003678007 |
| ENSMUSG00000022427 | TOMM22 | 0.379480311 | 2.03E-04 | 0.004261143 |
| ENSMUSG00000028998 | TOMM7 | 0.397501955 | 5.05E-04 | 0.008068595 |
| ENSMUSG00000000296 | TPD52L1 | 1.082580822 | 1.47E-05 | 7.08E-04 |
| ENSMUSG00000014846 | TPPP3 | 0.458657276 | 2.98E-05 | 0.001171341 |
| ENSMUSG00000006005 | TPR | -0.414211248 | 1.18E-04 | 0.002991246 |
| ENSMUSG00000002871 | TPRA1 | -0.438521964 | 4.99E-05 | 0.001686872 |
| ENSMUSG00000060126 | TPT1 | 0.408789012 | 4.16E-04 | 0.007106384 |
| ENSMUSG00000062296 | TRANK1 | -1.137035387 | 6.48E-08 | 1.17E-05 |
| ENSMUSG00000015013 | TRAPPC2L | 0.422474533 | 2.26E-04 | 0.00458579 |
| ENSMUSG00000020455 | TRIM11 | -0.432980977 | 1.10E-04 | 0.002812978 |
| ENSMUSG00000036989 | TRIM3 | -0.424987395 | 3.39E-05 | 0.001261965 |
| ENSMUSG00000045409 | TRIM39 | -0.708198142 | 3.25E-12 | 4.88E-09 |
| ENSMUSG00000042766 | TRIM46 | -0.378691042 | 3.74E-04 | 0.006609229 |
| ENSMUSG00000021071 | TRIM9 | -0.473186239 | 5.75E-05 | 0.001865435 |
| ENSMUSG00000022263 | TRIO | -0.396111114 | 2.94E-04 | 0.005561023 |
| ENSMUSG00000025272 | TRO | -0.436352251 | 3.22E-04 | 0.005905704 |
| ENSMUSG00000045482 | TRRAP | -0.488159437 | 2.15E-04 | 0.004430003 |
| ENSMUSG00000002496 | TSC2 | -0.385950055 | 5.82E-04 | 0.008786783 |
| ENSMUSG00000006736 | TSPAN31 | 0.347165508 | 4.07E-04 | 0.006974245 |
| ENSMUSG00000034156 | TSPOAP1 | -0.430798901 | 9.06E-05 | 0.00244949 |
| ENSMUSG00000027677 | TTC14 | -0.53827979 | 1.46E-06 | 1.28E-04 |
| ENSMUSG00000051786 | TUBGCP6 | -0.498163686 | 1.39E-05 | 6.83E-04 |

| | | | | |
|---|---|---|---|---|
| ENSMUSG00000014177 | TVP23B | 0.414552904 | 7.45E-05 | 0.002226046 |
| ENSMUSG00000053841 | TXLNA | -0.321234955 | 6.14E-04 | 0.00909294 |
| ENSMUSG00000030579 | TYROBP | 0.776898472 | 4.88E-04 | 0.007894704 |
| ENSMUSG00000086228 | UBAP1L | -1.013386534 | 1.41E-05 | 6.87E-04 |
| ENSMUSG00000042520 | Ubap2l | -0.379837262 | 2.49E-04 | 0.004917745 |
| ENSMUSG00000028960 | UBE4B | -0.421972799 | 1.56E-04 | 0.003599362 |
| ENSMUSG00000009741 | UBP1 | -0.543463918 | 4.82E-08 | 9.53E-06 |
| ENSMUSG00000066036 | UBR4 | -0.533787785 | 1.56E-04 | 0.003599362 |
| ENSMUSG00000037487 | UBR5 | -0.456369779 | 3.50E-04 | 0.00627791 |
| ENSMUSG00000029223 | UCHL1 | 0.508732703 | 3.57E-05 | 0.001313702 |
| ENSMUSG00000039512 | UHRF1BP1 | -0.533957721 | 1.34E-06 | 1.19E-04 |
| ENSMUSG00000034799 | UNC13A | -0.506382192 | 4.66E-06 | 3.06E-04 |
| ENSMUSG00000020770 | UNK | -0.360634536 | 4.74E-04 | 0.007755082 |
| ENSMUSG00000058301 | UPF1 | -0.423655771 | 1.80E-04 | 0.003912714 |
| ENSMUSG00000036572 | UPF3B | -0.456282246 | 1.67E-04 | 0.003746092 |
| ENSMUSG00000071654 | UQCC3 | 0.409508947 | 5.34E-04 | 0.008385423 |
| ENSMUSG00000059534 | UQCR10 | 0.472875124 | 2.47E-04 | 0.004887592 |
| ENSMUSG00000021520 | UQCRB | 0.360989348 | 9.90E-05 | 0.002630203 |
| ENSMUSG00000002395 | USE1 | 0.388800512 | 1.07E-04 | 0.002777095 |
| ENSMUSG00000068284 | USF3 | -0.435024104 | 3.44E-04 | 0.006214938 |
| ENSMUSG00000006676 | USP19 | -0.471010656 | 2.40E-05 | 9.88E-04 |
| ENSMUSG00000028514 | USP24 | -0.413103933 | 1.47E-04 | 0.003474702 |
| ENSMUSG00000032267 | USP28 | -0.388283686 | 7.79E-05 | 0.002292711 |
| ENSMUSG00000032376 | USP3 | -0.38557969 | 1.27E-04 | 0.003146969 |
| ENSMUSG00000033909 | USP36 | -0.395234867 | 1.09E-04 | 0.002805144 |
| ENSMUSG00000043411 | USP48 | -0.482324996 | 5.71E-06 | 3.49E-04 |
| ENSMUSG00000022710 | USP7 | -0.421970967 | 4.65E-05 | 0.00159614 |
| ENSMUSG00000037355 | UVSSA | -0.553317975 | 4.82E-04 | 0.007851613 |
| ENSMUSG00000007029 | VARS1 | -0.474544383 | 6.68E-06 | 3.94E-04 |
| ENSMUSG00000023951 | VEGFA | -0.404627639 | 3.24E-05 | 0.001233497 |
| ENSMUSG00000019772 | VIP | 2.491231247 | 5.17E-06 | 3.27E-04 |
| ENSMUSG00000035284 | VPS13C | -0.426759985 | 2.06E-04 | 0.0043106 |
| ENSMUSG00000046613 | VWA5B2 | -0.920347398 | 1.95E-08 | 4.65E-06 |
| ENSMUSG00000058997 | VWA8 | -0.320884057 | 3.16E-04 | 0.00582063 |
| ENSMUSG00000073434 | WDR90 | -0.579137213 | 2.82E-05 | 0.001125755 |
| ENSMUSG00000037989 | WNK2 | -0.483328535 | 1.63E-04 | 0.003691526 |
| ENSMUSG00000000131 | XPO6 | -0.411553798 | 1.03E-04 | 0.002721095 |
| ENSMUSG00000018554 | YBX2 | -1.496284026 | 2.96E-09 | 8.89E-07 |
| ENSMUSG00000024875 | YIF1A | 0.500468938 | 7.04E-05 | 0.002140958 |
| ENSMUSG00000021244 | YLPM1 | -0.508352372 | 1.65E-05 | 7.69E-04 |
| ENSMUSG00000034059 | YPEL4 | -0.459090437 | 6.67E-06 | 3.94E-04 |
| ENSMUSG00000094410 | ZBED6 | -0.685823002 | 9.49E-06 | 5.15E-04 |
| ENSMUSG00000022000 | ZC3H13 | -0.554231484 | 5.45E-06 | 3.36E-04 |
| ENSMUSG00000017478 | ZC3H18 | -0.307508901 | 5.84E-04 | 0.008802883 |
| ENSMUSG00000034163 | ZFC3H1 | -0.548666957 | 2.25E-05 | 9.41E-04 |

| | | | | |
|---|---|---|---|---|
| ENSMUSG00000040721 | ZFHX2 | -0.606450875 | 6.84E-05 | 0.002111064 |
| ENSMUSG00000093452 | Zfhx2os | -0.735863575 | 8.07E-05 | 0.002312173 |
| ENSMUSG00000034949 | ZFR2 | -0.485714126 | 6.00E-05 | 0.001907207 |
| ENSMUSG00000021286 | ZFYVE21 | 0.555265088 | 5.48E-05 | 0.00181456 |
| ENSMUSG00000066440 | ZFYVE26 | -0.577684047 | 1.15E-05 | 6.00E-04 |
| ENSMUSG00000027582 | ZGPAT | -0.653202775 | 3.59E-08 | 7.28E-06 |
| ENSMUSG00000032368 | ZIC1 | -0.396893372 | 3.36E-04 | 0.006100281 |
| ENSMUSG00000030757 | ZKSCAN2 | -0.640928554 | 1.11E-05 | 5.84E-04 |
| ENSMUSG00000027663 | ZMAT3 | -0.363815367 | 6.48E-04 | 0.009418998 |
| ENSMUSG00000031310 | ZMYM3 | -0.466192994 | 2.08E-05 | 8.87E-04 |
| ENSMUSG00000049764 | ZNF280B | -0.43743466 | 1.73E-04 | 0.003837447 |
| ENSMUSG00000015597 | ZNF318 | -0.509290301 | 6.76E-06 | 3.94E-04 |
| ENSMUSG00000039834 | ZNF335 | -0.424249 | 8.27E-05 | 0.002331043 |
| ENSMUSG00000059842 | ZNF341 | -0.535108268 | 1.36E-04 | 0.003307764 |
| ENSMUSG00000037855 | ZNF365 | 0.375775394 | 1.04E-04 | 0.002724288 |
| ENSMUSG00000038346 | ZNF384 | -0.439394583 | 1.52E-05 | 7.15E-04 |
| ENSMUSG00000027016 | ZNF385B | -0.359192814 | 6.75E-04 | 0.009668113 |
| ENSMUSG00000014198 | ZNF385C | -0.45156214 | 1.36E-05 | 6.74E-04 |
| ENSMUSG00000047036 | ZNF445 | -0.396178791 | 1.67E-04 | 0.003746092 |
| ENSMUSG00000000823 | ZNF512B | -0.720494212 | 7.04E-09 | 1.96E-06 |
| ENSMUSG00000005621 | ZNF592 | -0.373037753 | 5.34E-04 | 0.008385423 |
| ENSMUSG00000041130 | ZNF598 | -0.678875882 | 2.21E-11 | 1.53E-08 |
| ENSMUSG00000040524 | ZNF609 | -0.426350271 | 1.05E-04 | 0.002739848 |
| ENSMUSG00000037243 | ZNF692 | -0.717130113 | 1.36E-10 | 6.83E-08 |
| ENSMUSG00000020526 | ZNHIT3 | 0.484073218 | 3.63E-04 | 0.006461317 |
| ENSMUSG00000021819 | ZSWIM8 | -0.533348765 | 2.48E-06 | 1.93E-04 |
