## Supplementary material for "WWOX P47T loss-of-function mutation induces epilepsy, progressive neuroinflammation, and cerebellar degeneration in mice phenocopying human SCAR12": Wwox-P47T_Supplementary File 2.pdf

| | | PFC | | | CTX | | | HPC | | |
|---|---|---|---|---|---|---|---|---|---|---|
| Symbol | ID | Log Ratio | p-value | FDR | Log Ratio | p-value | FDR | Log Ratio | p-value | FDR |
| 2310002F09Rik | ENSMUSG0000 | 2.096 | 2.73E-09 | 4.79E-07 | 1.051 | 0.00196 | 0.0315 | -2.117 | 1.64E-10 | 3.75E-09 |
| 2410004I01Rik | ENSMUSG0000 | 1.561 | 2.02E-05 | 0.00089 | 1.068 | 0.00218 | 0.0339 | 1.273 | 0.000287 | 0.00184 |
| 5031425F14Rik | ENSMUSG0000 | 3.587 | 4.54E-09 | 7.01E-07 | 3.064 | 5.19E-07 | 5.14E-05 | 2.291 | 8.85E-05 | 0.000657 |
| 9630028I04Rik | ENSMUSG0000 | 2.403 | 5.37E-15 | 4.43E-12 | 1.842 | 1.42E-09 | 3.73E-07 | 2.536 | 2.92E-15 | 1.35E-13 |
| Acaa1b | ENSMUSG0000 | 2.319 | 8.62E-24 | 1.78E-20 | 1.939 | 1.51E-17 | 2.54E-14 | 2.291 | 5.17E-23 | 6.42E-21 |
| ADAM19 | ENSMUSG0000 | 1.208 | 6.32E-07 | 5.22E-05 | 1.118 | 3.93E-06 | 0.000272 | 1.636 | 2.73E-11 | 7.1E-10 |
| ADAMTS18 | ENSMUSG0000 | 1.254 | 0.00131 | 0.0231 | 1.401 | 0.000379 | 0.00956 | 1.772 | 9.16E-06 | 0.000088 |
| ADCYAP1 | ENSMUSG0000 | 1.851 | 4.49E-13 | 2.25E-10 | 1.882 | 2.01E-13 | 1.66E-10 | 3.146 | 1.47E-31 | 3.99E-29 |
| ADGRF4 | ENSMUSG0000 | 1.873 | 1.23E-06 | 9.08E-05 | 2.318 | 3.63E-09 | 8.11E-07 | 1.323 | 0.000453 | 0.00271 |
| AGMAT | ENSMUSG0000 | 1.783 | 3.71E-07 | 3.33E-05 | 2.253 | 2.28E-10 | 7.86E-08 | 3.587 | 4.07E-21 | 3.89E-19 |
| AI847159 | ENSMUSG0000 | 1.399 | 0.000209 | 0.00573 | 1.443 | 0.000222 | 0.00636 | 2.257 | 9.12E-08 | 1.34E-06 |
| AKR1C3 | ENSMUSG0000 | 1.561 | 1.9E-06 | 0.000132 | 1.482 | 4.88E-06 | 0.00032 | 1.899 | 8.32E-09 | 1.46E-07 |
| ALDH3A1 | ENSMUSG0000 | 3.339 | 1.02E-29 | 5.6E-26 | 2.583 | 5.35E-19 | 1.1E-15 | 4.143 | 1.77E-39 | 8.13E-37 |
| ALOX12B | ENSMUSG0000 | -1.809 | 2.38E-07 | 2.39E-05 | -1.535 | 1.03E-05 | 0.000582 | -2.238 | 2.13E-10 | 4.82E-09 |
| ALOX5 | ENSMUSG0000 | 2.692 | 5.2E-14 | 3.43E-11 | 2.009 | 7.13E-09 | 1.37E-06 | 2.002 | 8.05E-09 | 1.42E-07 |
| Apol9a/Apol9b | ENSMUSG0000 | 1.852 | 0.00058 | 0.0126 | 2.428 | 1.75E-05 | 0.000893 | 4.077 | 1.38E-10 | 3.22E-09 |
| ASCL2 | ENSMUSG0000 | 4.122 | 9.33E-06 | 0.000485 | 4.234 | 5.5E-06 | 0.000348 | 3.874 | 2.03E-05 | 0.000179 |
| ASPG | ENSMUSG0000 | 1.606 | 1.17E-06 | 8.74E-05 | 1.418 | 1.57E-05 | 0.000825 | 1.06 | 0.00113 | 0.00586 |
| ATP2A1 | ENSMUSG0000 | 3.723 | 6.93E-17 | 7.93E-14 | 3.567 | 3.19E-15 | 4.05E-12 | 2.475 | 5.09E-09 | 9.26E-08 |
| ATP2C2 | ENSMUSG0000 | 2.12 | 3.47E-06 | 0.000217 | 2.034 | 7.74E-06 | 0.000465 | 1.95 | 1.74E-05 | 0.000156 |
| B3GNT7 | ENSMUSG0000 | 2.472 | 6.3E-11 | 1.68E-08 | 2.11 | 1.38E-08 | 2.44E-06 | 2.433 | 1.18E-10 | 2.78E-09 |
| B3GNT8 | ENSMUSG0000 | 1.793 | 1.26E-14 | 9.49E-12 | 1.774 | 6E-14 | 5.51E-11 | 1.171 | 3.07E-07 | 4.09E-06 |
| B430212C06Rik | ENSMUSG0000 | 1.294 | 0.00063 | 0.0133 | 1.178 | 0.00146 | 0.0259 | -2.223 | 7.82E-12 | 2.22E-10 |
| BAZ1A | ENSMUSG0000 | 1.882 | 0.000219 | 0.00589 | 1.456 | 0.00368 | 0.0479 | 2.562 | 9.33E-07 | 1.12E-05 |
| BCOR | ENSMUSG0000 | 1.068 | 0.00178 | 0.0287 | 1.003 | 0.0033 | 0.0447 | 1.369 | 6.91E-05 | 0.000527 |
| BDNF | ENSMUSG0000 | 2.861 | 4.46E-08 | 5.5E-06 | 3.159 | 2.43E-09 | 5.82E-07 | 2.268 | 8.97E-06 | 8.63E-05 |
| BLNK | ENSMUSG0000 | 1.221 | 3.59E-05 | 0.00138 | 1.875 | 4.59E-10 | 1.4E-07 | 1.835 | 1.02E-09 | 2.06E-08 |
| BPIFB1 | ENSMUSG0000 | 2.671 | 1.84E-09 | 3.45E-07 | 2.631 | 1.65E-09 | 4.22E-07 | 2.34 | 3.52E-08 | 5.49E-07 |
| CAVIN4 | ENSMUSG0000 | 1.492 | 0.00192 | 0.0303 | 1.672 | 0.000511 | 0.0118 | 1.781 | 0.000223 | 0.00148 |
| CCNF | ENSMUSG0000 | 1.565 | 2.06E-05 | 0.000903 | 1.703 | 3.86E-06 | 0.00027 | 1.589 | 1.52E-05 | 0.000138 |
| CD164L2 | ENSMUSG0000 | 1.321 | 7.69E-09 | 1.11E-06 | 1.357 | 2.24E-09 | 5.52E-07 | 1.641 | 1.12E-12 | 3.55E-11 |

| | | | | | | | | | | |
|---|---|---|---|---|---|---|---|---|---|---|
| CD1D | ENSMUSG0000 | 1.901 | 4.82E-06 | 0.000284 | 1.933 | 3.01E-06 | 0.000221 | 1.937 | 2.86E-06 | 0.000031 |
| CD200R1 | ENSMUSG0000 | 1.537 | 6.53E-05 | 0.00227 | 1.334 | 0.000436 | 0.0106 | 3.256 | 5.26E-14 | 1.99E-12 |
| CD28 | ENSMUSG0000 | 2.971 | 4.53E-12 | 1.74E-09 | 3.082 | 1.15E-12 | 8.22E-10 | 4.382 | 4.64E-20 | 3.85E-18 |
| CDKN1A | ENSMUSG0000 | 2.214 | 5.41E-05 | 0.00196 | 2.218 | 5.26E-05 | 0.00209 | 3.366 | 4.8E-09 | 8.77E-08 |
| CDKN3 | ENSMUSG0000 | 1.906 | 0.00119 | 0.0215 | 2.085 | 0.000409 | 0.0101 | 3.51 | 3.37E-08 | 5.31E-07 |
| CEP295NL | ENSMUSG0000 | 1.907 | 0.000286 | 0.00722 | 2.072 | 7.48E-05 | 0.00273 | 4.553 | 3.72E-13 | 1.26E-11 |
| CHGA | ENSMUSG0000 | 1.266 | 3.71E-21 | 6.81E-18 | 1.382 | 9.09E-25 | 3E-21 | 1.513 | 3.74E-29 | 7.81E-27 |
| CIBAR2 | ENSMUSG0000 | 1.602 | 1.41E-06 | 0.000104 | 1.228 | 0.0002 | 0.00587 | 1.709 | 5.58E-07 | 7.05E-06 |
| CLEC18B | ENSMUSG0000 | 1.671 | 6.22E-07 | 5.21E-05 | 1.805 | 8.13E-08 | 0.000011 | 2.555 | 1.77E-13 | 6.28E-12 |
| COL10A1 | ENSMUSG0000 | 2.23 | 0.00232 | 0.0349 | 2.831 | 9.72E-05 | 0.00337 | 5.584 | 6.21E-11 | 1.52E-09 |
| COL5A3 | ENSMUSG0000 | 1.917 | 6.62E-06 | 0.000369 | 2.098 | 8.3E-07 | 7.66E-05 | 2.273 | 1.16E-07 | 1.67E-06 |
| CPA4 | ENSMUSG0000 | 1.601 | 0.000248 | 0.0065 | 1.604 | 0.000118 | 0.00391 | 3.438 | 8.75E-13 | 2.82E-11 |
| CRHBP | ENSMUSG0000 | 1.476 | 1.31E-08 | 1.8E-06 | 1.477 | 1.25E-08 | 2.25E-06 | 1.693 | 9.11E-11 | 2.18E-09 |
| CRYBA2 | ENSMUSG0000 | 2.424 | 5.49E-06 | 0.000316 | 3.227 | 4.78E-09 | 9.87E-07 | 4.482 | 3.49E-14 | 1.36E-12 |
| CRYBG2 | ENSMUSG0000 | 1.264 | 3.03E-05 | 0.00121 | 1.648 | 4.36E-08 | 6.61E-06 | 2.658 | 4.14E-17 | 2.42E-15 |
| D7Bwg0826e | ENSMUSG0000 | 2.034 | 1.03E-06 | 7.93E-05 | 2.167 | 1.86E-07 | 2.12E-05 | 2.654 | 6.92E-10 | 1.43E-08 |
| DHRS9 | ENSMUSG0000 | 2.268 | 0.000878 | 0.0171 | 2.94 | 2.15E-05 | 0.00106 | 3.729 | 2.98E-07 | 3.98E-06 |
| DIO3 | ENSMUSG0000 | -2.036 | 2.74E-05 | 0.00113 | -2.113 | 6.88E-06 | 0.000419 | -2.83 | 2.87E-09 | 5.4E-08 |
| DMKN | ENSMUSG0000 | 2.064 | 1.86E-08 | 2.44E-06 | 1.187 | 0.000819 | 0.0169 | 3.395 | 3.11E-18 | 2.07E-16 |
| DNMT3L | ENSMUSG0000 | 2.389 | 6.76E-09 | 1.01E-06 | 2.615 | 3.45E-11 | 1.58E-08 | 2.161 | 2.01E-08 | 3.31E-07 |
| DPM3 | ENSMUSG0000 | 1.259 | 5.65E-06 | 0.000322 | 1.448 | 1.93E-07 | 2.17E-05 | 1.617 | 7.13E-09 | 1.26E-07 |
| DRD4 | ENSMUSG0000 | 3.493 | 3.57E-07 | 3.22E-05 | 3.837 | 3.48E-08 | 5.42E-06 | 5.065 | 9.81E-12 | 2.76E-10 |
| DSP | ENSMUSG0000 | 2.139 | 4.46E-11 | 1.27E-08 | 1.335 | 0.000021 | 0.00104 | -2.547 | 2.58E-15 | 1.21E-13 |
| EFCAB6 | ENSMUSG0000 | 3.018 | 6.1E-10 | 1.27E-07 | 3.415 | 5.92E-12 | 3.15E-09 | 2.164 | 4.45E-06 | 0.000046 |
| ESRRB | ENSMUSG0000 | 1.351 | 2.73E-06 | 0.000178 | 1.163 | 5.59E-05 | 0.00219 | 1.251 | 1.55E-05 | 0.000141 |
| ETV3 | ENSMUSG0000 | 1.223 | 0.000162 | 0.00467 | 1.264 | 9.82E-05 | 0.00339 | 1.235 | 0.00014 | 0.000984 |
| ETV3L | ENSMUSG0000 | 5.965 | 8.24E-06 | 0.000439 | 5.889 | 1.21E-05 | 0.000653 | 4.196 | 0.000438 | 0.00264 |
| F2RL2 | ENSMUSG0000 | 1.482 | 6.56E-05 | 0.00228 | 1.942 | 4.09E-07 | 0.000042 | 4.131 | 1.18E-20 | 1.07E-18 |
| FCRL6 | ENSMUSG0000 | 3.279 | 0.00202 | 0.0315 | 4.569 | 1.12E-07 | 1.43E-05 | 3.834 | 2.91E-06 | 3.15E-05 |
| FGF10 | ENSMUSG0000 | -1.997 | 6.93E-12 | 2.44E-09 | -1.307 | 1.58E-06 | 0.000127 | -3.678 | 9.49E-38 | 4.02E-35 |
| FGR | ENSMUSG0000 | 4.917 | 8.78E-26 | 2.07E-22 | 4.057 | 1.48E-19 | 3.49E-16 | 4.711 | 8.1E-24 | 1.1E-21 |
| FHAD1 | ENSMUSG0000 | 1.163 | 4.41E-08 | 5.47E-06 | 1.168 | 5.21E-08 | 7.62E-06 | 1.109 | 2.22E-07 | 3.05E-06 |
| FNDC7 | ENSMUSG0000 | 2.168 | 3.26E-06 | 0.000207 | 2.057 | 6.44E-06 | 0.000398 | 3.574 | 1.94E-12 | 5.93E-11 |

| | | | | | | | | | | |
|---|---|---|---|---|---|---|---|---|---|---|
| FNDC9 | ENSMUSG0000 | 3.431 | 9.54E-06 | 0.000491 | 3.056 | 5.99E-05 | 0.0023 | 4.087 | 3.08E-07 | 4.1E-06 |
| FRZB | ENSMUSG0000 | -1.425 | 1.38E-09 | 2.64E-07 | -1.385 | 4.98E-09 | 1.02E-06 | -2.013 | 5.21E-17 | 3.03E-15 |
| GADL1 | ENSMUSG0000 | 1.111 | 0.0015 | 0.0255 | 1.422 | 5.11E-05 | 0.00205 | 1.047 | 0.00219 | 0.0103 |
| GAL | ENSMUSG0000 | 3.805 | 9.47E-07 | 7.37E-05 | 3.169 | 2.39E-05 | 0.00114 | 2.98 | 0.000062 | 0.00048 |
| GJD3 | ENSMUSG0000 | 1.144 | 3.06E-05 | 0.00122 | 1.068 | 0.000079 | 0.00285 | 1.261 | 4.55E-06 | 4.69E-05 |
| GLRA1 | ENSMUSG0000 | 3.833 | 5.1E-12 | 1.91E-09 | 2.637 | 5E-07 | 4.98E-05 | 3.744 | 1.4E-11 | 3.85E-10 |
| Gm11681 | ENSMUSG0000 | -1.34 | 0.00152 | 0.0256 | -1.309 | 0.000589 | 0.0131 | -3.633 | 4.22E-21 | 4E-19 |
| Gm13571 | ENSMUSG0000 | 2.599 | 0.00279 | 0.04 | 3.18 | 0.000268 | 0.00736 | 3.378 | 0.000173 | 0.00118 |
| Gm14637 | ENSMUSG0000 | 3.494 | 3.04E-05 | 0.00122 | 3.613 | 8.9E-06 | 0.000521 | 5.712 | 4.45E-10 | 9.51E-09 |
| Gm16048 | ENSMUSG0000 | 2.458 | 8.81E-11 | 2.24E-08 | 2.41 | 2.65E-09 | 6.17E-07 | 1.961 | 5.63E-07 | 7.11E-06 |
| Gm16701 | ENSMUSG0000 | 1.256 | 3.43E-05 | 0.00133 | 1.08 | 0.000278 | 0.00755 | 1.774 | 9.91E-09 | 1.73E-07 |
| Gm18786 | ENSMUSG0000 | 1.259 | 6.86E-05 | 0.00236 | 1.534 | 5.48E-06 | 0.000348 | 1.52 | 0.000007 | 6.91E-05 |
| Gm26916 | ENSMUSG0000 | 1.354 | 0.000288 | 0.00726 | 1.084 | 0.00155 | 0.0269 | 1.037 | 0.0024 | 0.0112 |
| Gm28626 | ENSMUSG0000 | 3.176 | 2.34E-09 | 4.19E-07 | 3.114 | 3.91E-09 | 8.5E-07 | 3.115 | 3.91E-09 | 7.22E-08 |
| Gm29100 | ENSMUSG0000 | 1.788 | 0.00353 | 0.0474 | 2.02 | 0.0011 | 0.0211 | 2.306 | 0.000229 | 0.00151 |
| Gm32224 | ENSMUSG0000 | 1.168 | 0.00344 | 0.0466 | 1.146 | 0.00375 | 0.0485 | 2.756 | 1.81E-10 | 4.13E-09 |
| Gm35853 | ENSMUSG0000 | 2.277 | 4.02E-06 | 0.000244 | 3.12 | 1.04E-09 | 2.76E-07 | 5.784 | 1.18E-22 | 1.4E-20 |
| Gm39168 | ENSMUSG0000 | 2.499 | 5.64E-09 | 8.46E-07 | 1.934 | 5.26E-06 | 0.000339 | 3.59 | 2.39E-13 | 8.24E-12 |
| Gm42864 | ENSMUSG0000 | -2.332 | 1.35E-12 | 6.02E-10 | -2.041 | 2.35E-10 | 7.87E-08 | -2.525 | 5.29E-15 | 2.37E-13 |
| Gm42865 | ENSMUSG0000 | -2.334 | 2.37E-07 | 2.39E-05 | -1.799 | 3.82E-05 | 0.00166 | -2.027 | 2.61E-06 | 2.84E-05 |
| Gm42901 | ENSMUSG0000 | 1.525 | 5.58E-06 | 0.000319 | 1.257 | 0.000224 | 0.0064 | 1.383 | 5.37E-05 | 0.000424 |
| Gm45187 | ENSMUSG0000 | -1.721 | 1.73E-06 | 0.000121 | -1.443 | 7.23E-05 | 0.00267 | -3.17 | 8.33E-19 | 5.81E-17 |
| Gm45346 | ENSMUSG0000 | 2.109 | 7.22E-06 | 0.000395 | 1.877 | 4.73E-05 | 0.00192 | 2.493 | 2.38E-07 | 3.23E-06 |
| Gm5912 | ENSMUSG0000 | 1.479 | 0.00048 | 0.0109 | 1.73 | 0.000023 | 0.00111 | 4.105 | 2.4E-16 | 1.28E-14 |
| Gm7276 | ENSMUSG0000 | 1.03 | 0.00153 | 0.0256 | 1.24 | 0.000148 | 0.00468 | 1.735 | 1.91E-07 | 2.66E-06 |
| GPR139 | ENSMUSG0000 | 2.073 | 3.41E-09 | 5.75E-07 | 1.182 | 0.000394 | 0.00981 | 3.201 | 1.48E-18 | 9.99E-17 |
| GPR150 | ENSMUSG0000 | -1.703 | 3.16E-08 | 4.01E-06 | -1.219 | 0.000045 | 0.00186 | -1.071 | 0.000366 | 0.00226 |
| GPR3 | ENSMUSG0000 | 1.393 | 0.0034 | 0.0462 | 1.667 | 0.000504 | 0.0117 | 2.234 | 4.96E-06 | 5.08E-05 |
| GPRC5C | ENSMUSG0000 | 2.211 | 3.57E-09 | 5.95E-07 | 1.756 | 1.8E-06 | 0.00014 | 2.061 | 2.89E-08 | 4.63E-07 |
| GUCY2C | ENSMUSG0000 | 1.521 | 0.000114 | 0.00356 | 1.251 | 0.00172 | 0.0288 | 1.314 | 0.00106 | 0.00559 |
| GXYLT2 | ENSMUSG0000 | 1.321 | 3.32E-05 | 0.0013 | 1.352 | 1.66E-05 | 0.000855 | 1.428 | 5.86E-06 | 5.89E-05 |
| HCN4 | ENSMUSG0000 | 1.666 | 2.16E-09 | 3.91E-07 | 1.096 | 6.49E-05 | 0.00245 | 2.239 | 4.55E-15 | 2.05E-13 |
| HELT | ENSMUSG0000 | 2.092 | 0.000169 | 0.00484 | 2.156 | 0.000151 | 0.00473 | 5.209 | 9.2E-14 | 3.35E-12 |

| | | | | | | | | | | |
|---|---|---|---|---|---|---|---|---|---|---|
| HHIPL1 | ENSMUSG0000 | 1.253 | 2.03E-11 | 6.22E-09 | 1.186 | 2.08E-10 | 7.62E-08 | 1.557 | 1.53E-16 | 8.47E-15 |
| HMCN2 | ENSMUSG0000 | 3.276 | 1.92E-09 | 3.56E-07 | 3.117 | 7.12E-09 | 1.37E-06 | 3.458 | 2.72E-10 | 6E-09 |
| HP | ENSMUSG0000 | 2.127 | 0.00074 | 0.0149 | 2.488 | 3.73E-06 | 0.000263 | 1.85 | 0.000368 | 0.00227 |
| HSPB3 | ENSMUSG0000 | 1.814 | 6.13E-07 | 5.19E-05 | 2.041 | 5.91E-08 | 8.34E-06 | 2.989 | 9E-14 | 3.28E-12 |
| HTRA4 | ENSMUSG0000 | 2.954 | 4.1E-10 | 9.03E-08 | 3.178 | 2.88E-11 | 1.36E-08 | 1.952 | 1.63E-05 | 0.000146 |
| IGHG1 | ENSMUSG0000 | 6.239 | 1.81E-13 | 9.98E-11 | 3.752 | 2.65E-07 | 2.88E-05 | 4.602 | 2.42E-09 | 4.58E-08 |
| IL17RB | ENSMUSG0000 | 3.574 | 5.32E-10 | 1.16E-07 | 3.39 | 2.71E-09 | 6.22E-07 | 3.818 | 5.46E-11 | 1.35E-09 |
| INHBA | ENSMUSG0000 | 2.547 | 0.000138 | 0.00417 | 3.113 | 5.35E-06 | 0.000342 | 3.016 | 9.57E-06 | 9.15E-05 |
| JSRP1 | ENSMUSG0000 | 1.584 | 0.000144 | 0.00429 | 2.017 | 1.75E-06 | 0.000138 | 2.971 | 1.35E-11 | 3.72E-10 |
| KCNQ1 | ENSMUSG0000 | 1.907 | 4.68E-06 | 0.000278 | 1.712 | 2.96E-05 | 0.00136 | 1.774 | 1.56E-05 | 0.000142 |
| KLHL40 | ENSMUSG0000 | 1.737 | 0.000181 | 0.00512 | 1.879 | 5.54E-05 | 0.00217 | 2.101 | 7.69E-06 | 7.53E-05 |
| Klk2-ps | ENSMUSG0000 | 3.468 | 1.09E-10 | 2.68E-08 | 3.644 | 1.7E-10 | 6.54E-08 | 2.141 | 3.04E-05 | 0.000256 |
| KLK8 | ENSMUSG0000 | 1.455 | 1.16E-07 | 1.29E-05 | 1.119 | 3.55E-05 | 0.00158 | -1.788 | 2.88E-11 | 7.45E-10 |
| KRT7 | ENSMUSG0000 | 1.829 | 0.000207 | 0.00569 | 2.911 | 2.18E-08 | 3.71E-06 | 3.779 | 3.89E-11 | 9.95E-10 |
| KRT75 | ENSMUSG0000 | 5.376 | 1.08E-07 | 1.21E-05 | 6.094 | 9.91E-09 | 1.82E-06 | 4.404 | 3.19E-06 | 3.43E-05 |
| LHX3 | ENSMUSG0000 | 4.069 | 3.38E-12 | 1.39E-09 | 3.147 | 1.66E-09 | 4.22E-07 | 4.271 | 3.74E-14 | 1.45E-12 |
| LIPG | ENSMUSG0000 | 1.586 | 0.000423 | 0.00986 | 1.485 | 0.000938 | 0.0188 | 2.59 | 2.89E-08 | 4.64E-07 |
| LLGL2 | ENSMUSG0000 | 1.495 | 6.54E-08 | 7.66E-06 | 1.319 | 1.22E-06 | 0.000104 | 1.277 | 2.32E-06 | 2.56E-05 |
| Lrrc74b | ENSMUSG0000 | 2.086 | 5.06E-11 | 1.39E-08 | 1.299 | 2.18E-05 | 0.00107 | 1.364 | 8.85E-06 | 8.53E-05 |
| LSP1 | ENSMUSG0000 | 1.012 | 6.31E-06 | 0.000356 | 1.175 | 1.94E-07 | 2.17E-05 | 1.926 | 1.06E-16 | 6E-15 |
| Ly6g6e | ENSMUSG0000 | 1.917 | 0.000147 | 0.00436 | 2.836 | 5.49E-08 | 7.96E-06 | 2.356 | 4.08E-06 | 4.27E-05 |
| LYZL4 | ENSMUSG0000 | 2.434 | 9.82E-05 | 0.00317 | 2.617 | 3.02E-05 | 0.00138 | 3.673 | 2.84E-08 | 4.56E-07 |
| MAB21L3 | ENSMUSG0000 | 3.318 | 3.88E-06 | 0.000238 | 1.988 | 0.00279 | 0.04 | 2.951 | 2.27E-05 | 0.000196 |
| MARCHF11 | ENSMUSG0000 | 1.274 | 1.2E-08 | 1.68E-06 | 1.083 | 1.08E-06 | 9.55E-05 | 1.636 | 5.63E-13 | 1.86E-11 |
| MCEMP1 | ENSMUSG0000 | 2.527 | 6.82E-10 | 1.41E-07 | 2.16 | 2.55E-08 | 4.25E-06 | 3.166 | 7.57E-14 | 2.79E-12 |
| MEDAG | ENSMUSG0000 | 1.418 | 1.05E-06 | 0.00008 | 1.557 | 1.03E-07 | 1.33E-05 | 2.746 | 4.31E-19 | 3.1E-17 |
| MEI1 | ENSMUSG0000 | 1.045 | 0.000626 | 0.0133 | 1.023 | 0.000831 | 0.0171 | 2.919 | 1.26E-18 | 8.58E-17 |
| Morrbid | ENSMUSG0000 | 2.094 | 1.59E-08 | 2.14E-06 | 1.557 | 9.7E-06 | 0.000558 | 1.76 | 7.22E-07 | 8.88E-06 |
| MYBPC1 | ENSMUSG0000 | 2.669 | 3.23E-32 | 2.66E-28 | 3.113 | 5.15E-42 | 8.51E-38 | 4.836 | 1.95E-81 | 1.07E-77 |
| MYBPC3 | ENSMUSG0000 | 3.037 | 1.3E-28 | 3.57E-25 | 3.052 | 7.17E-28 | 3.95E-24 | 3.511 | 6E-34 | 2.02E-31 |
| MYL4 | ENSMUSG0000 | 1.551 | 7.2E-17 | 7.93E-14 | 1.395 | 6.48E-14 | 5.64E-11 | 4.252 | 2.03E-88 | 3.35E-84 |
| MYO15A | ENSMUSG0000 | 2.73 | 2.9E-14 | 1.99E-11 | 2.164 | 7.1E-10 | 2.02E-07 | 2.459 | 8.15E-12 | 2.31E-10 |
| MYOC | ENSMUSG0000 | -1.894 | 0.000187 | 0.00528 | -2.113 | 2.72E-05 | 0.00127 | -2.643 | 2.51E-07 | 3.4E-06 |

| | | | | | | | | | | |
|---|---|---|---|---|---|---|---|---|---|---|
| NKX1-2 | ENSMUSG0000 | 6.887 | 7.34E-15 | 5.77E-12 | 7.154 | 1.79E-15 | 2.47E-12 | 8.075 | 2.47E-17 | 1.48E-15 |
| NLRP6 | ENSMUSG0000 | -1.785 | 4.17E-06 | 0.000251 | -1.302 | 0.000395 | 0.00981 | -1.184 | 0.00135 | 0.00687 |
| NPPA | ENSMUSG0000 | 1.79 | 0.000224 | 0.00598 | 2.818 | 1.46E-07 | 1.71E-05 | 2.691 | 4.99E-07 | 6.4E-06 |
| NPTX2 | ENSMUSG0000 | 3.4 | 1.05E-06 | 0.00008 | 3.21 | 3.35E-06 | 0.000244 | 4.046 | 1.7E-08 | 2.84E-07 |
| NPW | ENSMUSG0000 | 5.124 | 3.29E-11 | 9.69E-09 | 4.965 | 8.57E-11 | 3.72E-08 | 4.93 | 1.1E-10 | 2.6E-09 |
| NPY | ENSMUSG0000 | 1.137 | 4.33E-05 | 0.00162 | 1.58 | 1.9E-08 | 3.3E-06 | 2.315 | 9.18E-16 | 4.57E-14 |
| NRAP | ENSMUSG0000 | 1.338 | 2.41E-05 | 0.00102 | 1.848 | 2.75E-09 | 6.22E-07 | 3.255 | 2.73E-22 | 3.08E-20 |
| NTF3 | ENSMUSG0000 | -2.147 | 1.21E-07 | 1.32E-05 | -1.186 | 0.0015 | 0.0263 | -2.789 | 4.26E-13 | 1.42E-11 |
| NUGGC | ENSMUSG0000 | 3.633 | 1.21E-13 | 6.89E-11 | 3.349 | 4.28E-12 | 2.62E-09 | 4.359 | 3.6E-16 | 1.88E-14 |
| Oas1f | ENSMUSG0000 | 7.272 | 9.7E-09 | 1.38E-06 | 6.016 | 7.78E-07 | 0.000073 | 7.13 | 7.83E-08 | 1.16E-06 |
| OTOP2 | ENSMUSG0000 | 1.614 | 5.29E-05 | 0.00192 | 2.222 | 7.29E-08 | 0.00001 | 3.673 | 2.52E-16 | 1.34E-14 |
| OTOP3 | ENSMUSG0000 | 2.679 | 5.33E-05 | 0.00193 | 2.747 | 3.23E-05 | 0.00146 | 2.537 | 0.00011 | 0.000794 |
| PAPPA | ENSMUSG0000 | 3.201 | 9.3E-06 | 0.000484 | 3.503 | 1.39E-06 | 0.000116 | 2.621 | 0.000171 | 0.00117 |
| PARVG | ENSMUSG0000 | 1.94 | 1.48E-29 | 6.12E-26 | 1.566 | 3.32E-20 | 9.13E-17 | 1.056 | 3.14E-10 | 6.84E-09 |
| PDLIM1 | ENSMUSG0000 | 1.242 | 7.13E-05 | 0.00242 | 1.325 | 2.19E-05 | 0.00108 | 1.598 | 3.79E-07 | 4.96E-06 |
| PIK3R6 | ENSMUSG0000 | 1.456 | 1.45E-11 | 4.71E-09 | 1.369 | 2.38E-10 | 7.87E-08 | 1.972 | 5.84E-19 | 4.15E-17 |
| PIWIL1 | ENSMUSG0000 | 1.529 | 1.19E-06 | 8.82E-05 | 2.36 | 3.85E-10 | 1.2E-07 | 1.818 | 3.23E-07 | 4.28E-06 |
| PLAU | ENSMUSG0000 | 1.074 | 6.85E-08 | 7.86E-06 | 1.154 | 6.24E-09 | 1.23E-06 | 1.606 | 1.87E-15 | 9.04E-14 |
| PLB1 | ENSMUSG0000 | 1.577 | 3.51E-05 | 0.00136 | 1.778 | 3.52E-06 | 0.000254 | 1.584 | 3.15E-05 | 0.000264 |
| PLET1 | ENSMUSG0000 | 3.149 | 7.97E-06 | 0.000427 | 3.251 | 2.84E-06 | 0.000211 | 2.958 | 1.45E-05 | 0.000133 |
| PLK5 | ENSMUSG0000 | -2.204 | 1.48E-09 | 2.81E-07 | -2.525 | 5.9E-12 | 3.15E-09 | -2.513 | 7.18E-12 | 2.05E-10 |
| PPM1J | ENSMUSG0000 | 3.316 | 1.13E-11 | 3.81E-09 | 3.455 | 2.58E-12 | 1.65E-09 | 4.965 | 1.92E-20 | 1.67E-18 |
| PPP1R13L | ENSMUSG0000 | 1.043 | 0.000508 | 0.0114 | 1.039 | 0.000538 | 0.0123 | 1.007 | 0.000787 | 0.00433 |
| PRSS22 | ENSMUSG0000 | 1.108 | 6.13E-05 | 0.00215 | 1.291 | 6.72E-06 | 0.000411 | 4.711 | 1.04E-34 | 3.73E-32 |
| PRSS23 | ENSMUSG0000 | 1.752 | 1.24E-15 | 1.2E-12 | 1.879 | 1.54E-17 | 2.54E-14 | 1.424 | 5.75E-11 | 1.42E-09 |
| Prss23os | ENSMUSG0000 | 1.752 | 2.26E-07 | 2.32E-05 | 1.698 | 7.07E-06 | 0.000429 | 1.166 | 0.00131 | 0.00666 |
| PRTN3 | ENSMUSG0000 | 1.899 | 6.5E-14 | 4.13E-11 | 1.132 | 4.89E-06 | 0.00032 | 1.909 | 1.02E-13 | 3.71E-12 |
| PSRC1 | ENSMUSG0000 | -2.049 | 1.64E-11 | 5.2E-09 | -1.062 | 0.000303 | 0.00801 | -1.427 | 1.38E-06 | 0.000016 |
| PSTPIP1 | ENSMUSG0000 | 1.8 | 1.31E-16 | 1.35E-13 | 1.375 | 1.69E-10 | 6.54E-08 | 1.57 | 5.57E-13 | 1.84E-11 |
| PTGR1 | ENSMUSG0000 | 2.158 | 6.68E-13 | 3.15E-10 | 2.223 | 5.67E-14 | 5.51E-11 | 1.019 | 0.000302 | 0.00192 |
| PTGS1 | ENSMUSG0000 | 1.632 | 1.16E-08 | 1.64E-06 | 1.509 | 1.21E-07 | 1.49E-05 | 1.478 | 2.14E-07 | 2.94E-06 |
| PTGS2 | ENSMUSG0000 | 2.388 | 0.000465 | 0.0107 | 2.999 | 0.000018 | 0.00091 | 1.893 | 0.00474 | 0.0196 |
| RASA4 | ENSMUSG0000 | 1.423 | 9.61E-14 | 5.88E-11 | 1.312 | 5.73E-12 | 3.15E-09 | 1.798 | 1.52E-20 | 1.35E-18 |

| | | | | | | | | | | |
|---|---|---|---|---|---|---|---|---|---|---|
| RBP3 | ENSMUSG0000 | 4.3 | 5.22E-09 | 7.98E-07 | 2.573 | 3.35E-05 | 0.0015 | 2.091 | 0.000606 | 0.00348 |
| REC114 | ENSMUSG0000 | 1.751 | 1.32E-06 | 0.000097 | 2.106 | 4.29E-09 | 9.07E-07 | 2.261 | 2.96E-10 | 6.46E-09 |
| RET | ENSMUSG0000 | 1.791 | 2.89E-14 | 1.99E-11 | 1.295 | 2E-08 | 3.44E-06 | 1.242 | 7.29E-08 | 1.09E-06 |
| RETN | ENSMUSG0000 | 2.346 | 1.29E-05 | 0.000619 | 2.344 | 5.71E-06 | 0.00036 | 3.556 | 1.11E-10 | 2.63E-09 |
| RGS13 | ENSMUSG0000 | 3.09 | 2.69E-08 | 3.47E-06 | 2.769 | 4.49E-07 | 4.55E-05 | 3.44 | 1.23E-09 | 2.44E-08 |
| RGS2 | ENSMUSG0000 | 1.416 | 6.81E-05 | 0.00234 | 1.435 | 5.46E-05 | 0.00214 | 2.371 | 9.99E-11 | 2.38E-09 |
| RIN3 | ENSMUSG0000 | 1.342 | 1.98E-09 | 3.63E-07 | 1.047 | 0.000002 | 0.000154 | 1.249 | 1.63E-08 | 2.74E-07 |
| RIPK3 | ENSMUSG0000 | 3.081 | 3.59E-15 | 3.12E-12 | 2.755 | 3.01E-13 | 2.36E-10 | 2.318 | 2.03E-10 | 4.58E-09 |
| RNF39 | ENSMUSG0000 | 1.163 | 0.000569 | 0.0124 | 1.33 | 8.86E-05 | 0.00312 | 2.22 | 2.42E-10 | 5.38E-09 |
| SCCPDH | ENSMUSG0000 | 1.685 | 5.54E-29 | 1.83E-25 | 1.585 | 6.13E-26 | 2.53E-22 | 1.529 | 2.6E-24 | 3.77E-22 |
| SCG2 | ENSMUSG0000 | 1.435 | 1.33E-08 | 1.81E-06 | 1.36 | 6.74E-08 | 9.43E-06 | 1.236 | 8.69E-07 | 1.05E-05 |
| SCN7A | ENSMUSG0000 | 2.406 | 5.95E-10 | 1.27E-07 | 1.786 | 2.81E-06 | 0.00021 | 2.274 | 4.93E-09 | 8.98E-08 |
| Sct | ENSMUSG0000 | 3.361 | 9.35E-10 | 1.88E-07 | 3.571 | 5.26E-12 | 3.1E-09 | 3.986 | 8.11E-14 | 2.97E-12 |
| SEMA4A | ENSMUSG0000 | 1.081 | 4.04E-09 | 6.38E-07 | 1.185 | 1.21E-10 | 4.87E-08 | 1.688 | 1.56E-19 | 1.19E-17 |
| SERPINA11 | ENSMUSG0000 | 3.172 | 2.36E-05 | 0.001 | 3.731 | 1.19E-06 | 0.000102 | 3.284 | 9.44E-06 | 9.04E-05 |
| SFRP2 | ENSMUSG0000 | 1.624 | 1.25E-05 | 0.000606 | 1.604 | 0.000016 | 0.000829 | 2.161 | 1.26E-08 | 2.16E-07 |
| SLC38A8 | ENSMUSG0000 | 3.187 | 2.63E-15 | 2.41E-12 | 1.84 | 4.6E-07 | 4.63E-05 | 2.428 | 3.14E-10 | 6.84E-09 |
| SLC6A5 | ENSMUSG0000 | 5.032 | 7.43E-07 | 5.93E-05 | 3.943 | 4.32E-05 | 0.0018 | 6.158 | 8.7E-09 | 1.53E-07 |
| STC2 | ENSMUSG0000 | 2.836 | 2.46E-07 | 2.43E-05 | 2.988 | 5.73E-08 | 8.16E-06 | 2.043 | 0.000112 | 0.000808 |
| SYTL1 | ENSMUSG0000 | 1.358 | 4.72E-06 | 0.00028 | 1.248 | 2.28E-05 | 0.0011 | 3.27 | 3.9E-24 | 5.46E-22 |
| TAFA3 | ENSMUSG0000 | 2.798 | 3.76E-10 | 8.45E-08 | 3.134 | 2.21E-10 | 7.75E-08 | 3.312 | 4.01E-11 | 1.02E-09 |
| TBC1D2 | ENSMUSG0000 | 1.03 | 0.000373 | 0.00883 | 1.028 | 0.000384 | 0.00963 | 1.101 | 0.00015 | 0.00104 |
| TDRD9 | ENSMUSG0000 | 3.557 | 1.27E-12 | 5.84E-10 | 3.335 | 1.96E-11 | 9.52E-09 | 5.429 | 1.65E-21 | 1.69E-19 |
| TGFB1 | ENSMUSG0000 | 1.479 | 5.98E-10 | 1.27E-07 | 1.263 | 1.01E-07 | 1.32E-05 | 1.164 | 8.73E-07 | 1.05E-05 |
| TGFB1I1 | ENSMUSG0000 | 1.339 | 3.56E-07 | 3.22E-05 | 1.155 | 1.03E-05 | 0.000583 | 1.291 | 8.97E-07 | 1.08E-05 |
| THEG | ENSMUSG0000 | 1.288 | 0.00143 | 0.0246 | 1.79 | 2.31E-05 | 0.00111 | 1.685 | 6.27E-05 | 0.000485 |
| THEMIS2 | ENSMUSG0000 | 1.526 | 5.41E-08 | 6.5E-06 | 1.148 | 4.05E-05 | 0.00172 | 1.697 | 2.67E-09 | 5.03E-08 |
| THRSP | ENSMUSG0000 | 1.042 | 3.5E-07 | 3.22E-05 | 1.385 | 1.67E-11 | 8.53E-09 | 1.04 | 3.51E-07 | 4.61E-06 |
| TLL1 | ENSMUSG0000 | 1.741 | 0.000479 | 0.0109 | 2.25 | 8.26E-06 | 0.000487 | 1.819 | 0.000249 | 0.00162 |
| TMEM154 | ENSMUSG0000 | 4.759 | 8.34E-09 | 1.2E-06 | 3.787 | 1.37E-06 | 0.000115 | 4.314 | 8.39E-08 | 1.24E-06 |
| TMEM40 | ENSMUSG0000 | 2.194 | 5.56E-08 | 6.6E-06 | 3.237 | 1.89E-14 | 2.23E-11 | 4.621 | 2.9E-24 | 4.13E-22 |
| TMEM91 | ENSMUSG0000 | 1.604 | 1.43E-10 | 3.41E-08 | 1.211 | 9.65E-07 | 8.66E-05 | 2.399 | 9.87E-21 | 9E-19 |
| TMPRSS3 | ENSMUSG0000 | 2.488 | 3.92E-09 | 6.28E-07 | 1.58 | 2.97E-05 | 0.00136 | 1.333 | 0.000289 | 0.00185 |

| | | | | | | | | | | |
|---|---|---|---|---|---|---|---|---|---|---|
| TNNC1 | ENSMUSG0000 | 1.793 | 1.06E-17 | 1.34E-14 | 2.508 | 1.4E-30 | 1.15E-26 | 4.175 | 1.65E-67 | 3.41E-64 |
| TNNI3 | ENSMUSG0000 | 1.761 | 0.000136 | 0.0041 | 1.566 | 0.000391 | 0.00976 | 2.652 | 3.19E-08 | 5.06E-07 |
| TRNP1 | ENSMUSG0000 | 1.315 | 1.01E-19 | 1.51E-16 | 1.095 | 2.86E-14 | 3.15E-11 | 1.001 | 3.3E-12 | 9.86E-11 |
| UCMA | ENSMUSG0000 | 3.242 | 8.34E-12 | 2.87E-09 | 3.347 | 2.24E-12 | 1.54E-09 | 2.6 | 1.41E-08 | 2.39E-07 |
| UPK3A | ENSMUSG0000 | 1.223 | 3.53E-05 | 0.00136 | 1.275 | 1.29E-05 | 0.000691 | 1.43 | 1.34E-06 | 1.56E-05 |
| USH1G | ENSMUSG0000 | 2.18 | 2.68E-06 | 0.000176 | 2.5 | 1.3E-07 | 1.56E-05 | 2.162 | 3.65E-06 | 3.87E-05 |
| VDR | ENSMUSG0000 | 1.312 | 0.00331 | 0.0452 | 1.364 | 0.00205 | 0.0324 | 2.885 | 1.2E-09 | 2.39E-08 |
| VGF | ENSMUSG0000 | 1.389 | 6.86E-07 | 5.56E-05 | 1.334 | 1.79E-06 | 0.00014 | 1.791 | 2.47E-10 | 5.51E-09 |
| Wfdc18 | ENSMUSG0000 | 2.174 | 4.17E-09 | 6.5E-07 | 2.513 | 1.7E-11 | 8.53E-09 | 3.881 | 1.45E-21 | 1.52E-19 |
| WNT16 | ENSMUSG0000 | 1.561 | 0.000103 | 0.00329 | 1.348 | 0.000658 | 0.0141 | 1.569 | 8.84E-05 | 0.000656 |
| WNT9B | ENSMUSG0000 | 4.143 | 3.29E-07 | 3.07E-05 | 4.301 | 1.24E-07 | 1.49E-05 | 2.721 | 0.000315 | 0.00199 |
| ZC3H12D | ENSMUSG0000 | 1.568 | 4.97E-05 | 0.00182 | 2.099 | 8.97E-09 | 1.68E-06 | 1.61 | 0.000006 | 6.02E-05 |
