## Supplementary material for "WWOX P47T loss-of-function mutation induces epilepsy, progressive neuroinflammation, and cerebellar degeneration in mice phenocopying human SCAR12": Wwox-P47T_Supplementary Tables.pdf

### Supplementary Table 1: Genotypes of pups from *Wwox*$^{P47T/WT}$ x *Wwox*$^{P47T/WT}$ cross

| Genotype | *Wwox*$^{WT/WT}$ | *Wwox*$^{P47T/WT}$ | *Wwox*$^{P47T/P47T}$ |
|---|---|---|---|
| **Actual number of pups** | 18 | 45 | 19 |
| **Expected number of pups** | 20.5 | 41 | 20.5 |

Data represents genotyping result from 82 mice from 12 litters

### Supplementary Table 2: Blood chemistry analysis

| | ***Wwox***$^{WT/WT}$ **(n=9)** | ***Wwox***$^{P47T/WT}$ **(n=16)** | ***Wwox***$^{P47T/P47T}$ **(n=16)** | |
|---|---|---|---|---|
| | **Average ± SEM** | **Average ± SEM** | **Average ± SEM** | **p value** |
| **Albumin (g/dL)** | 3.93 ± 0.13 | 3.58 ± 0.13 | 3.53 ± 0.13 | ns |
| **Alkaline phosphatase (U/L)** | 70.11 ± 13.14 | 88.13 ± 33.62 | 55.2 ± 4.39 | ns |
| **Alanine aminotransferase (U/L)** | 79 ± 33.80 | 98.25 ± 30 | 48.73 ± 3.79 | ns |
| **Amylase (U/L)** | 1081.55 ± 48.40 | 1206.81 ± 71.01 | 1145.28 ± 40.91 | ns |
| **Total bilirubin (mg/dL)** | 0.3 ± 0 | 0.31 ± 0.04 | 0.29 ± 0.01 | ns |
| **Urea nitrogen (mg/dL)** | 19.11 ± 1.14 | 22.26 ± 2.98 | 20.86 ± 1.24 | ns |
| **Calcium (mg/dL)** | 11.57 ± 0.15 | 11.5 ± 0.31 | 11.26 ± 0.26 | ns |
| **Phosphorus (mg/dL)** | 11.66 ± 0.78 | 13.4 ± 0.65 | 12.62 ± 0.79 | ns |
| **Creatinine (U/L)** | < 0.24 | < 0.36 | < 0.31 | ns |
| **Glucose (mg/dL)** | 296 ± 25.47 | 267.25 ± 30.25 | 254.33 ± 22.52 | ns |
| **Sodium (mmol/L)** | 153.66 ± 1.25 | 154.56 ± 1.18 | 152.86 ± 1.49 | ns |
| **Potassium (mmol/L)** | >8.48 | > 9.38 | >9.22 | ns |
| **Total protein (g/dL)** | 5.51 ± 0.22 | 5.63 ± 0.14 | 5.38 ± 0.08 | ns |
| **Globulin (g/dL)** | 1.78 ± 0.08 | 2.05 ± 0.15 | 1.83 ± 0.14 | ns |

### Supplementary Table 3: Blood hematology analysis

| | *Wwox*$^{WT/WT}$ (n=9) | *Wwox*$^{P47T/WT}$ (n=16) | *Wwox*$^{P47T/P47T}$ (n=16) | |
|---|---|---|---|---|
| | **Average ± SEM** | **Average ± SEM** | **Average ± SEM** | **p value** |
| **White blood cells ($10^3$ cells/µL)** | 5.71 ± 0.74 | 4.99 ± 0.60 | 4.08 ± 0.45 | ns |
| **Neutrophils ($10^3$ cells/µL)** | 1.99 ± 0.33 | 2.15 ± 0.56 | 1.53 ± 0.26 | ns |
| **Lymphocytes ($10^3$ cells/µL)** | 3.28 ± 0.35 | 2.5 ± 0.27 | 2.28 ± 0.23 | ns |
| **Monocytes ($10^3$ cells/µL)** | 0.23 ± 0.04 | 0.2 ± 0.02 | 0.19 ± 0.02 | ns |
| **Eosinophils ($10^3$ cells/µL)** | 0.16 ± 0.06 | 0.09 ± 0.02 | 0.07 ± 0.01 | ns |
| **Basophils ($10^3$ cells/µL)** | 0.05 ± 0.02 | 0.02 ± 0.01 | 0.02 ± 0 | ns |
| **Neutrophils (%)** | 33.79 ± 2.17 | 38.27 ± 4.77 | 35.52 ± 2.89 | ns |
| **Lymphocytes (%)** | 59.18 ± 1.97 | 54.41 ± 4.39 | 56.8 ± 2.82 | ns |
| **Monocytes (%)** | 4.15 ± 0.73 | 5.02 ± 0.89 | 5.37 ± 0.64 | ns |
| **Eosinophils (%)** | 2.21 ± 0.57 | 1.81 ± 0.31 | 1.84 ± 0.23 | ns |
| **Basophils (%)** | 0.65 ± 0.18 | 0.46 ± 0.09 | 0.45 ± 0.07 | ns |
| **Red blood cells ($10^6$ cells/µl)** | 9.22 ± 0.26 | 8.99 ± 0.49 | 9.26 ± 0.30 | ns |
| **Hemoglobin (g/dL)** | 11.31 ± 0.39 | 10.93 ± 0.53 | 11.7 ± 0.45 | ns |
| **Hematocrit (%)** | 45.15 ± 1.39 | 44.32 ± 2.08 | 46.37 ± 1.40 | ns |
| **Mean corpuscular volume (fL)** | 48.95 ± 0.19 | 50.08 ± 1.17 | 50.15 ± 0.34 | ns |
| **Mean corpuscular Hb (pg)** | 12.26 ± 0.21 | 12.28 ± 0.25 | 12.65 ± 0.28 | ns |
| **MCHC (g/dL)** | 25.05 ± 0.44 | 24.59 ± 0.42 | 25.21 ± 0.55 | ns |
| **Red cell distribution width (%)** | 16.68 ± 0.15 | 19.14 ± 1.31 | 17.39 ± 0.25 | ns |
| **Platelets ($10^3$ cells/µL)** | 1342.7 ± 128.01 | 1378.87 ± 138.92 | 1098.5 ± 93.72 | ns |
| **Mean platelet volume (fL)** | 5.4 ± 0.06 | 5.51 ± 0.07 | 5.36 ± 0.09 | ns |

### Supplementary Table 4: Tissue histology analysis

| **Genotype** | **Neoplasia (%)** | **Cholestasis (%)** | ***Focal liver degeneration (%)*** | ***Nephropathy (%)*** |
|---|---|---|---|---|
| ***Wwox***$^{WT/WT}$ (n=7) | 2 (29%) | 0 | 0 | 1 (14%) |
| ***Wwox***$^{P47T/WT}$ (n=19) | 6 (33%) | 4 (22%) | 5 (28%) | 13 (72%) |
| ***Wwox***$^{P47T/P47T}$ (n=21) | 1 (5%) | 9 (43%) | 5 (24%) | 11 (52%) |
