## Supplementary material for "WWOX P47T loss-of-function mutation induces epilepsy, progressive neuroinflammation, and cerebellar degeneration in mice phenocopying human SCAR12": Wwox-P47T_Supplementray Figures .pdf

### SUPPLEMENTARY FIGURES AND SUPPLEMENTARY FIGURE LEGENDS

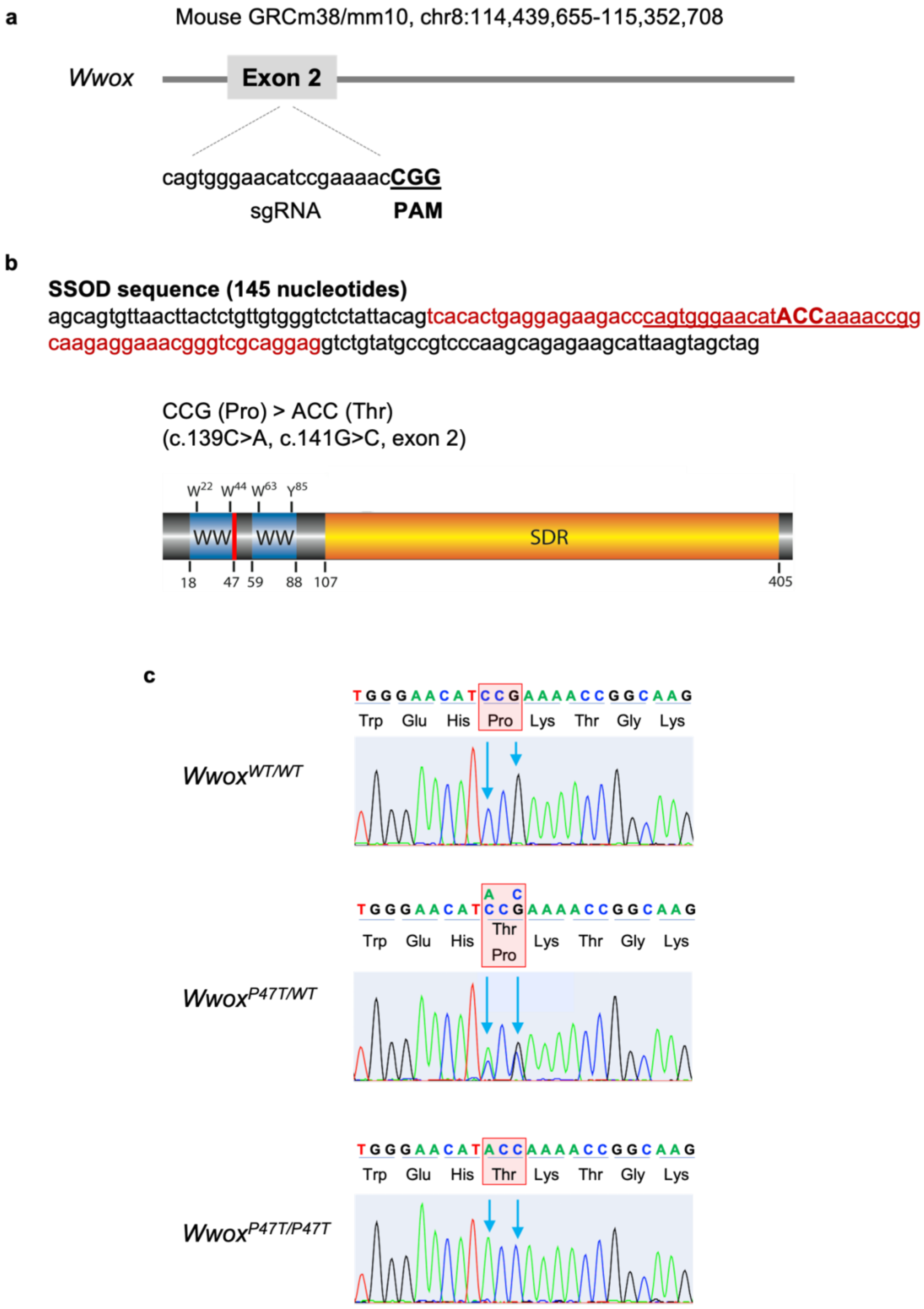

**Supplementary Figure S1 | Generation of *Wwox*$^{P47T/P47T}$ mutant mice using CRISPR/Cas9.** (a) A schematic illustration showing sgRNA and PAM sequence designed to target exon 2 of Wwox gene (Mouse GRCm38/mm10, chr8:114,439,655-115,352,708). (b) The 145 nucleotide long sequence shows SSOD, where red letters indicate exon 2 and the underline sequence shows sgRNA and PAM sequence within exon 2, red underlined capitalized ACC (Thr) codon shows the substituted sequence to replace CCG (Pro) (c.139C>A, c.141G>C). The Wwox protein illustration shows the location of p.Pro47Thr mutation at the 47 position in the first WW domain. (c) Sanger sequencing histograms showing results from *Wwox*$^{WT/WT}$, *Wwox*$^{P47T/WT}$, and *Wwox*$^{P47T/P47T}$ mice, where the wildtype shows CCG codon, the heterozygous shows overlapping peaks of c.139C/A and c.141G/C, and the homozygous shows ACC codon with a substitution of c.139C>A and c.141G>C.

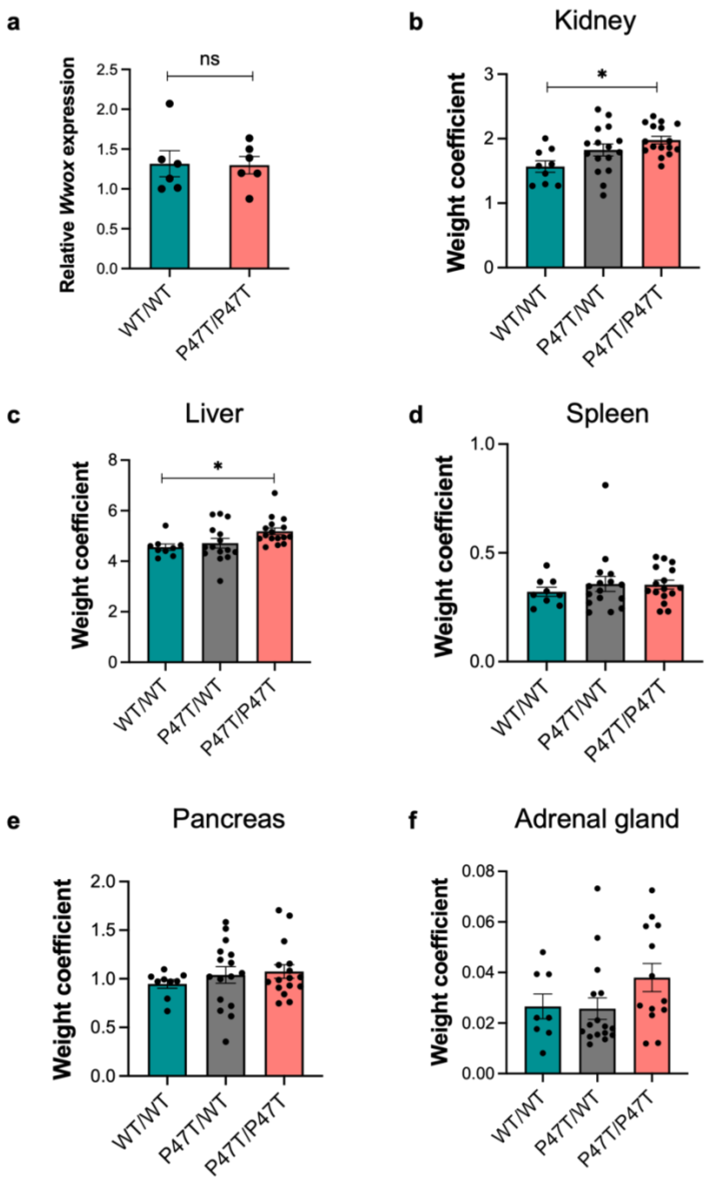


**Supplementary Figure S2 | Wwox mRNA expression in CB and comparative weight coefficient of major organs.** (a) Bar graph showing mean *Wwox* mRNA expression measured by qRT-PCR in $Wwox^{WT/WT}$ and $Wwox^{P47T/P47T}$ CB (n=6 mice/group). Error bars represent ± SEM, p-value= ns (not significant), unpaired Student's *t*-test. (b-f) Bar graphs showing relative organ weight (weight coefficient) of kidney (b), liver (c), spleen (d), pancreas (e), and adrenal gland (f) from $Wwox^{WT/WT}$ (n=9), $Wwox^{P47T/WT}$ (n=16), and $Wwox^{P47T/P47T}$ (n=16) mice. Data are represented as mean ± SEM, Kidney weight coefficient *p-value < 0.05, One-Way ANOVA (Tukey's post-hoc test), Liver weight coefficient *p-value < 0.05, Kruskal-Wallis test.

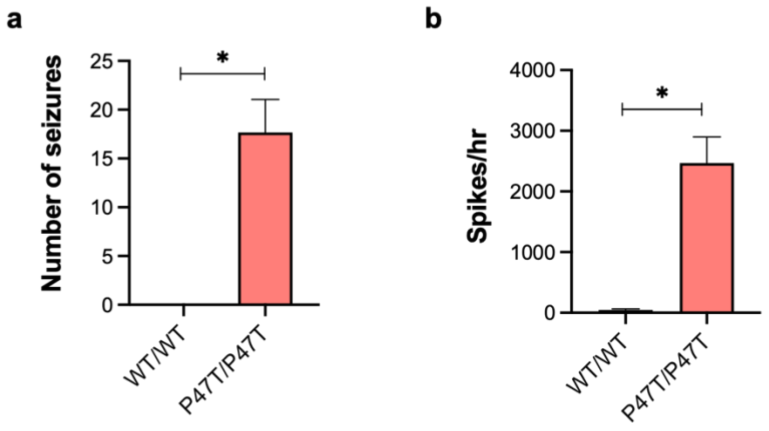


**Supplementary Figure S3 | *Wwox*$^{P47T/P47T}$ mice display spontaneous seizure activity.** Average number of seizures (a) and spikes per hour (b) in *Wwox*$^{P47T/P47T}$ mice (n=3) compared to *Wwox*$^{WT/WT}$ (n=3) control mice. Data is represented as mean ± SEM, *p-value < 0.01, unpaired Student's *t*-test.

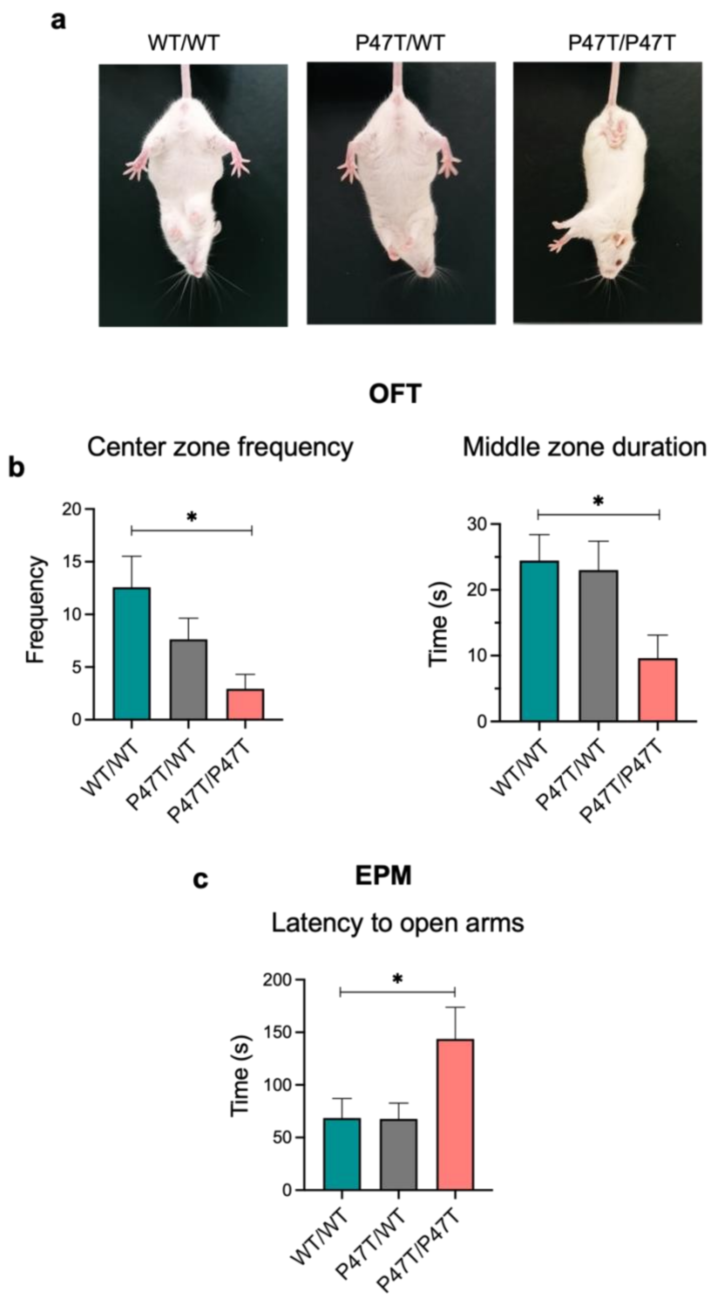


**Supplementary Figure S4 | *Wwox*$^{P47T/P47T}$ mice display abnormal social and motor behaviors.** (a) Tail suspension test shows abnormal hind limb clasping reflex in *Wwox*$^{P47T/P47T}$ mice. (b) Bar graphs showing quantitative measurement of center zone frequency and time spent by test mice in middle zone of the OFT arena. (c) Bar graph showing quantitative measurement of the amount of time taken by test mice to first enter the open arms in EPM. Data are represented as mean ± SEM, *p-value < 0.05, One-Way ANOVA (Tukey's post-hoc test).

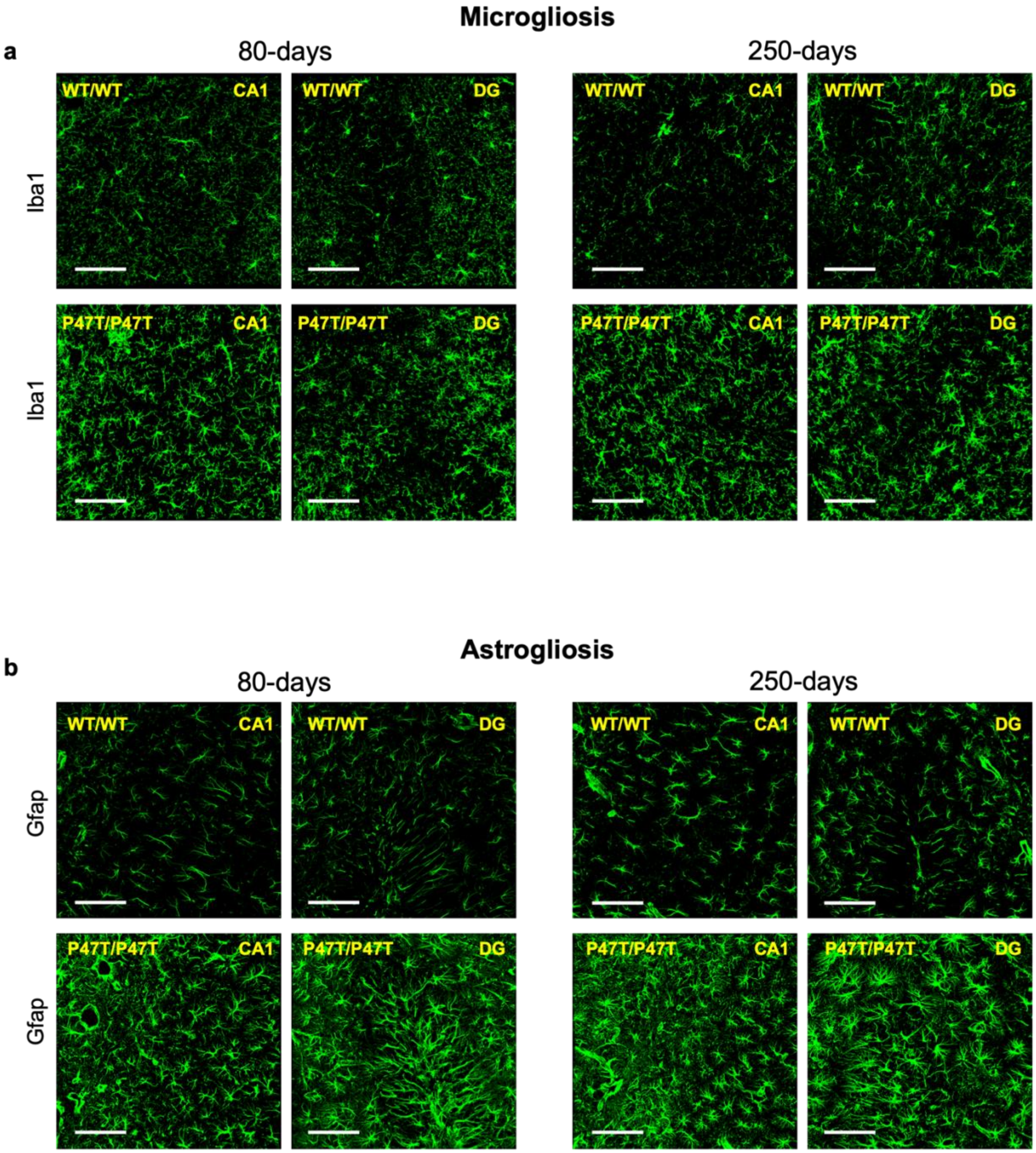


**Supplementary Figure S5 | *Wwox*$^{P47T/P47T}$ hippocampi display progressive microgliosis and astrogliosis.** Representative high magnification Iba1+ (a) microglia and Gfap+ (b) astrocytes immunostaining images in HPC CA1 and DG regions of *Wwox*$^{WT/WT}$ and *Wwox*$^{P47T/P47T}$ samples at 80 and 250 days of age as indicated, scale bar=100 μm.

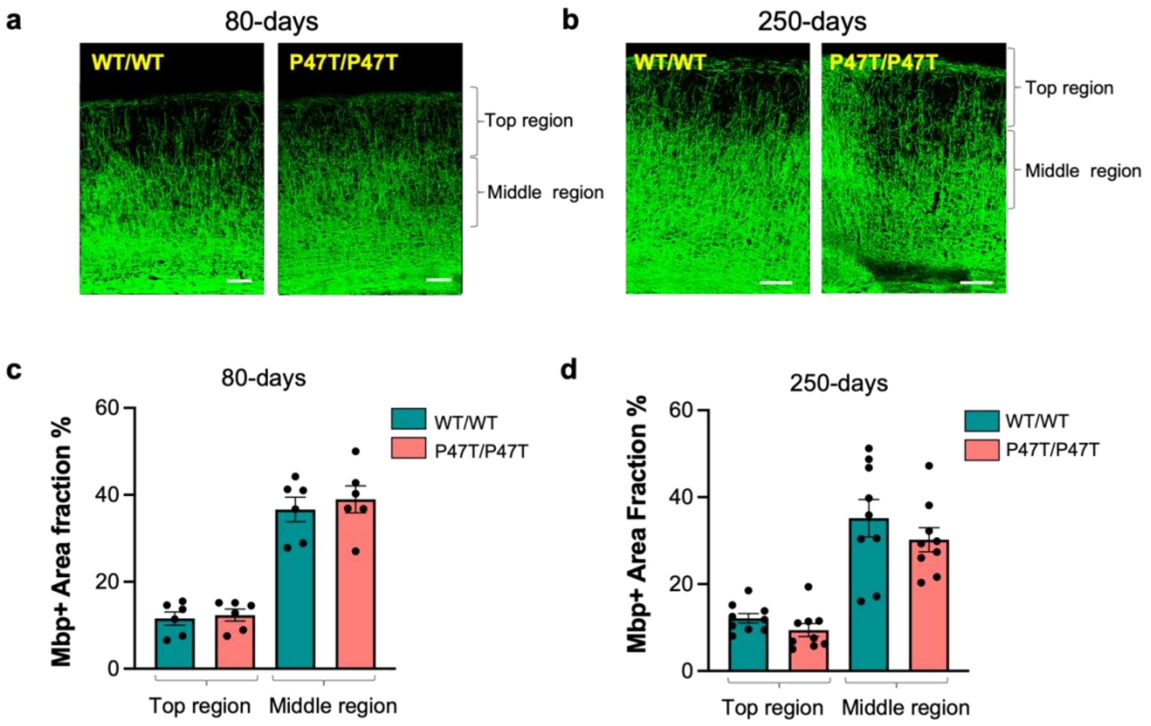


**Supplementary Figure S6 | Wwox P47T mutation does not affect myelination in parietal cortex.** Representative confocal microscopy images immunostained with Mbp illustrating myelination in the parietal cortex region above the corpus callosum of *Wwox*$^{WT/WT}$ and *Wwox*$^{P47T/P47T}$ mice from 80 and 250-days age groups, scale bar=100 µm (80-days) and scale bar=200 µm (250-days). Bar graph showing comparative quantitation of Mbp AF % in the top region (from the top of the image between 0-325 µm (80-days) and 0-450 µm (250-days) and middle region (between 326-650 µm (80-days) and 451-900 µm (250-days) of *Wwox*$^{WT/WT}$ (n=3) and *Wwox*$^{P47T/P47T}$ (n=3) mice. Data is represented as mean ± SEM, p-value is not significant, unpaired Student's *t*-test.

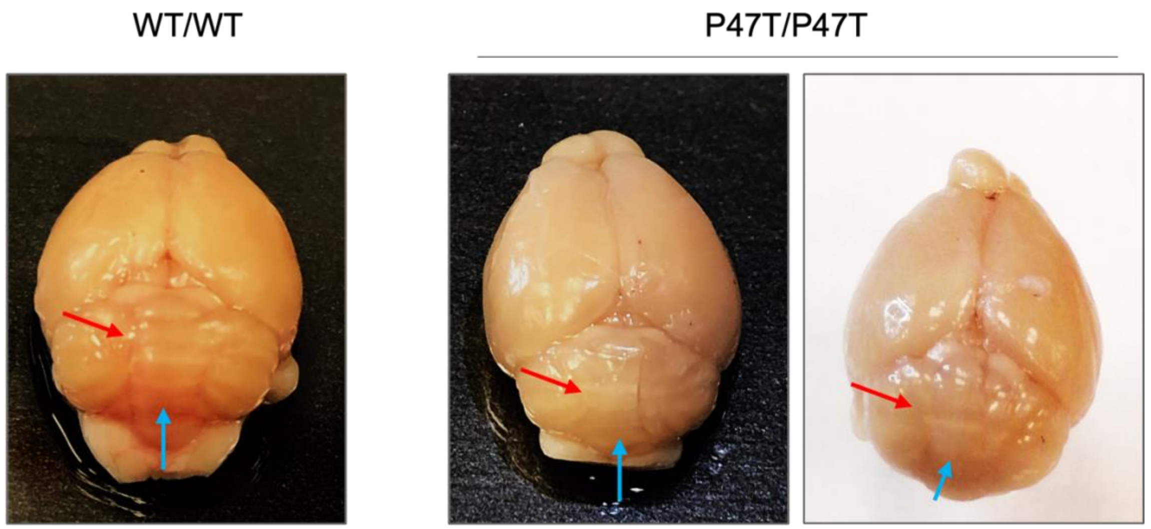


**Supplementary Figure S7 | *Wwox*$^{P47T/P47T}$ mice display aberrant cerebellar morphology.** Representative macroscopic photographs of *Wwox*$^{P47T/P47T}$ mice whole brains (right two panels) showing aberrant cerebellar morphology with fusion of interhemispheric fissure (red arrow) and cerebellar vermis lobules (blue arrow) in comparison to *Wwox*$^{WT/WT}$ (left panel).

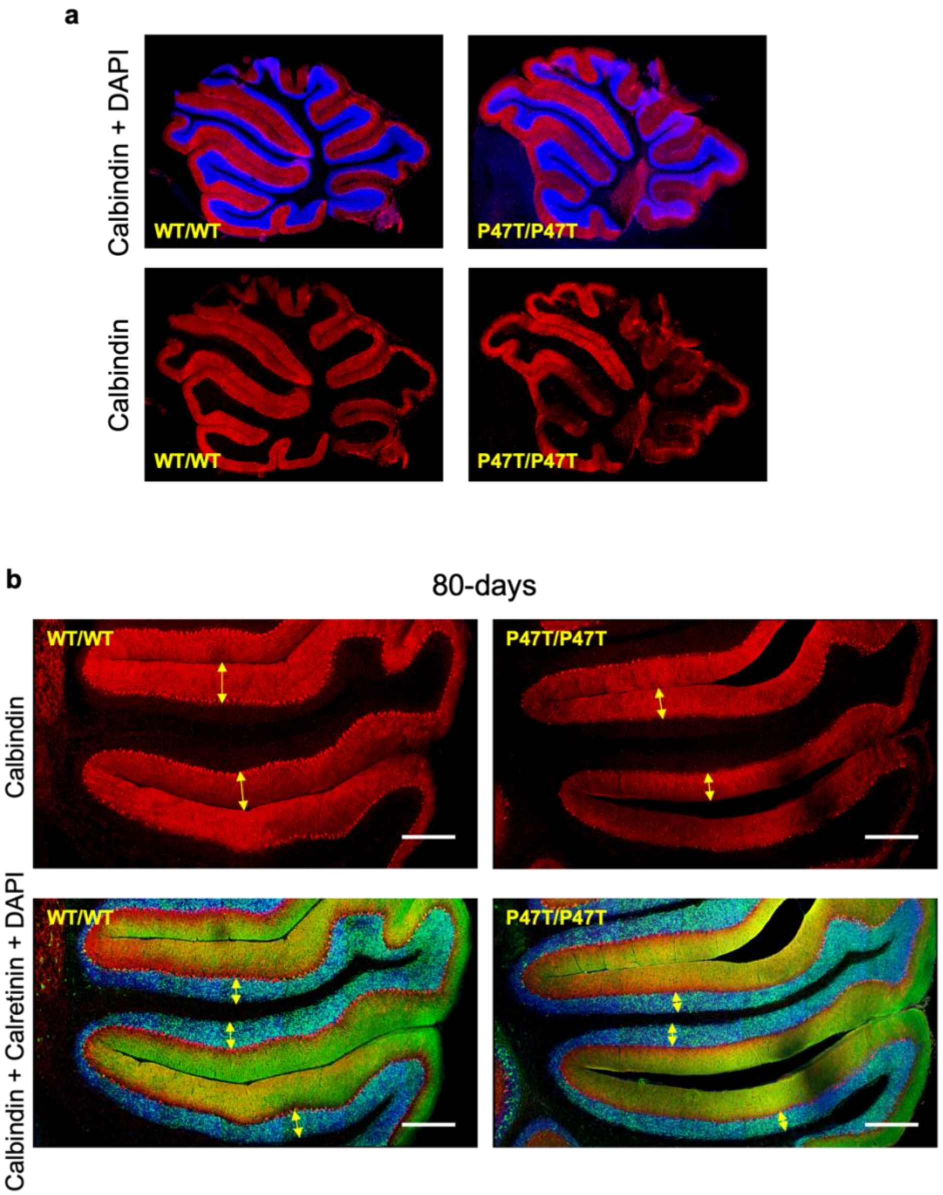


**Supplementary Figure S8 | *Wwox* LoF leads to cerebellar atrophy.** (a) Representative calbindin staining PCs (red) and DAPI stained low resolution immunofluorescence images from the mid-sagittal sections of the vermis region from *Wwox*$^{WT/WT}$ and *Wwox*$^{P47T/P47T}$ mice cerebella of 250-days age group displaying cerebellar atrophy with several regions showing loss of PC and dendrite staining. (b) Representative calbindin immunofluorescence of PCs (top panel, red) and calretinin (green) and DAPI (blue) immunofluorescence of GL (bottom panel) in preculminate and primary fissures between cerebellar lobules III-VI of 80-days old *Wwox*$^{WT/WT}$ and *Wwox*$^{P47T/P47T}$ mice. Yellow arrows in the top panel mark the ML thickness and in the lower panel mark the GL thickness, scale bar=200 μm.

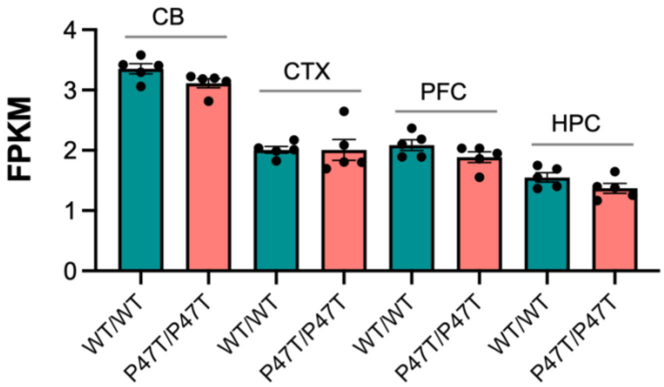


**Supplementary Figure S9 | Wwox P47T mutation does not affect *Wwox* mRNA expression.** Bar graph showing FPKM values of *Wwox* gene from the RNA-Seq transcriptome profiling data from $Wwox^{WT/WT}$ (n=5) and $Wwox^{P47T/P47T}$ (n=5) CB, CTX, PFC, and HPC tissues. Data is represented as mean ± SEM, p-value is not significant, unpaired Student's *t*-test.

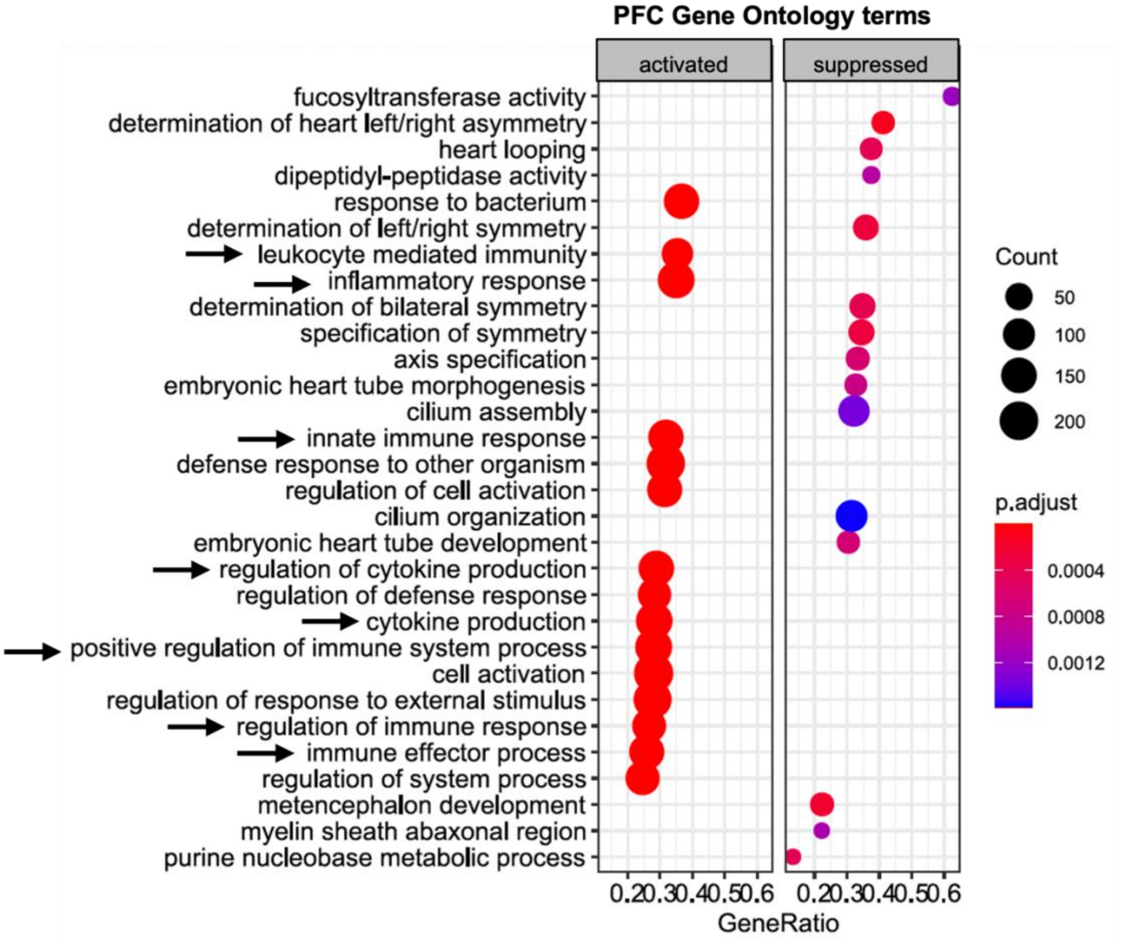


**Supplementary Figure S10 | *Wwox* LoF brains display enrichment of neuroinflammation pathways in PFC.** clusterProfiler plot showing Gene Ontology terms enriched in $Wwox^{P47T/P47T}$ PFC, black arrows indicate activation of bioprocesses related to inflammation. Color of the circle represents adjusted p-value and size represents number of genes enriched. Left panel shows activate and right panel shows suppressed GO terms.

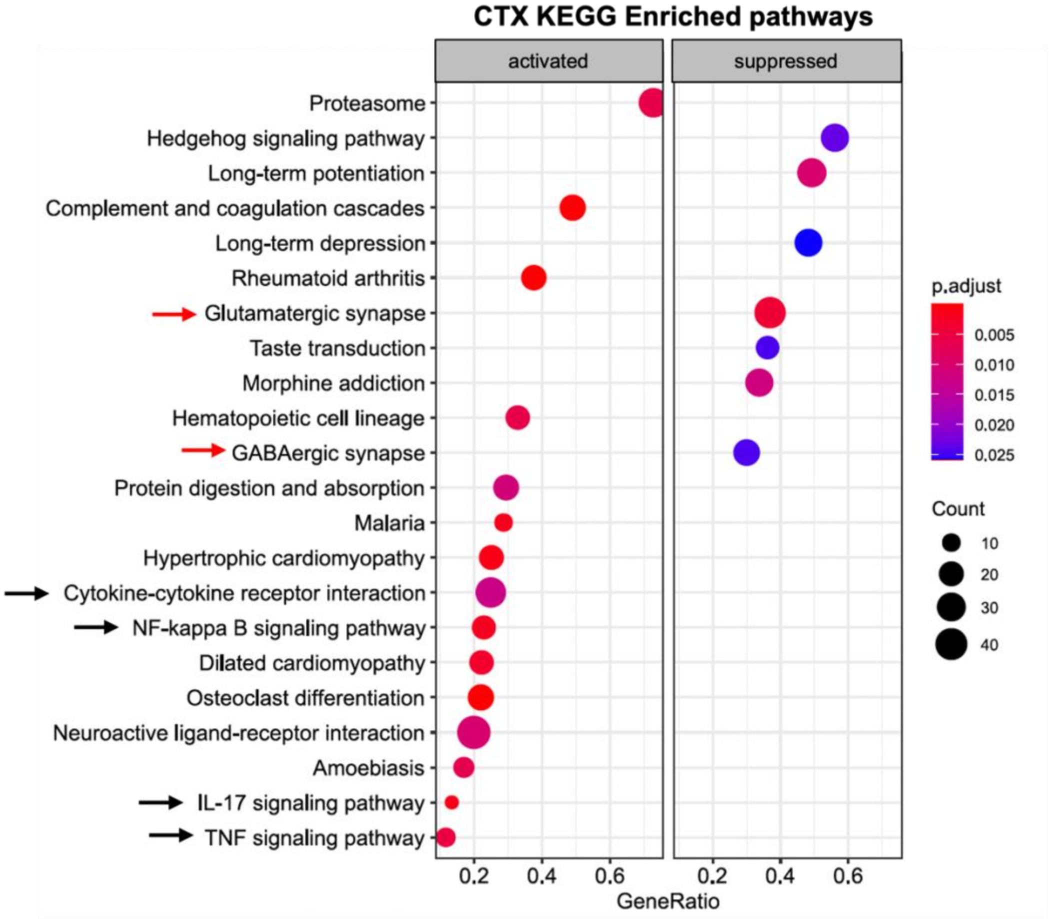


**Supplementary Figure S11 | KEGG pathway enrichment analysis in CTX.** Bar graph showing enrichment of KEGG pathways in CTX samples. Red arrows indicate suppression of GABAergic and Glutamatergic synapses and black arrows indicate activation of inflammation bioprocesses. Color of the circle represents adjusted p-value and size represents number of genes enriched. Left panel shows activate and right panel shows suppressed KEGG pathways.

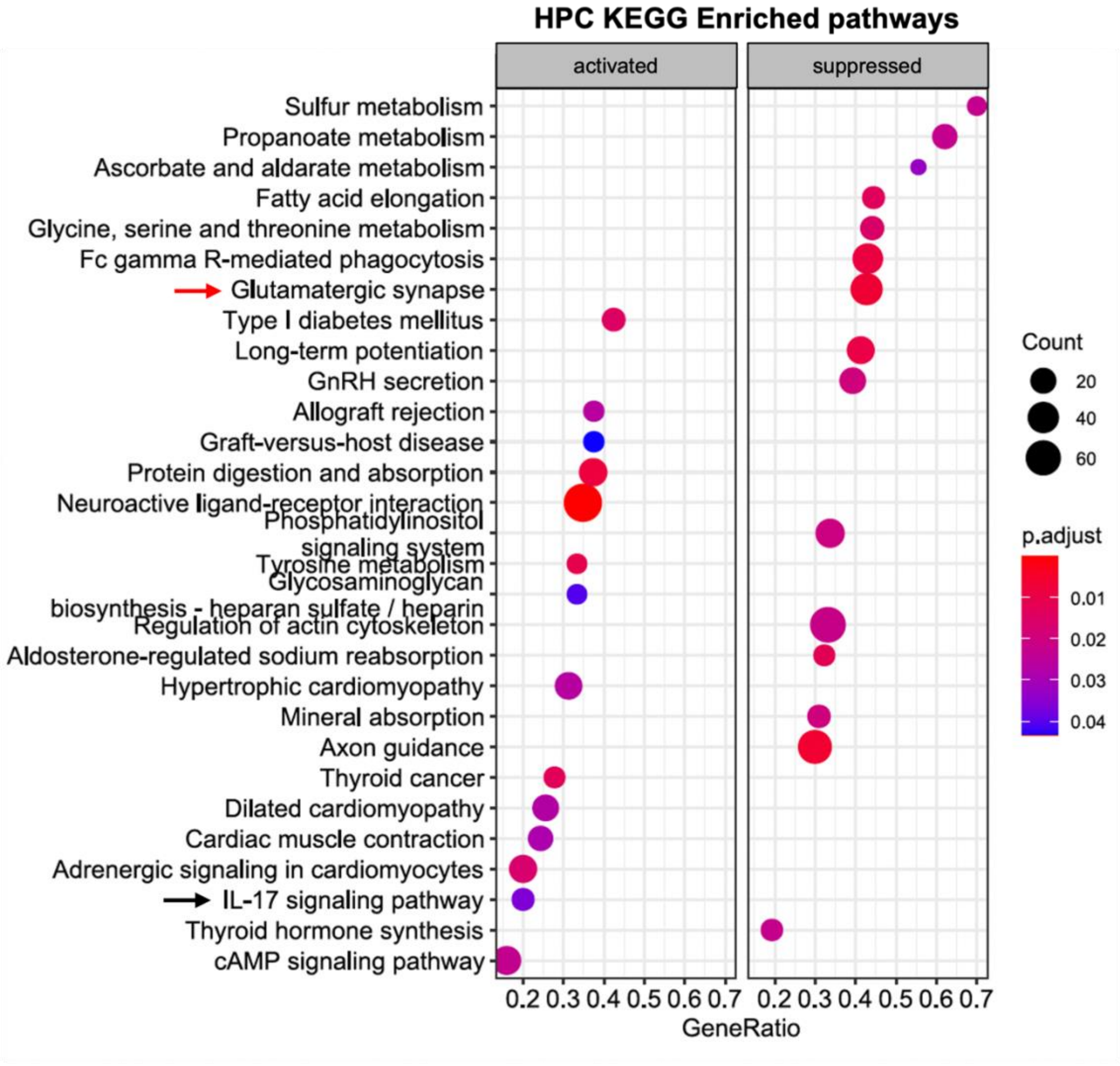


**Supplementary Figure S11 | KEGG pathway enrichment analysis in HPC.** Bar graph showing enrichment of KEGG pathways in HPC samples. Red arrow indicate suppression Glutamatergic synapses and black arrow indicate activation of IL-17 signaling. Color of the circle represents adjusted p-value and size represents number of genes enriched. Left panel shows activate and right panel shows suppressed KEGG pathways.

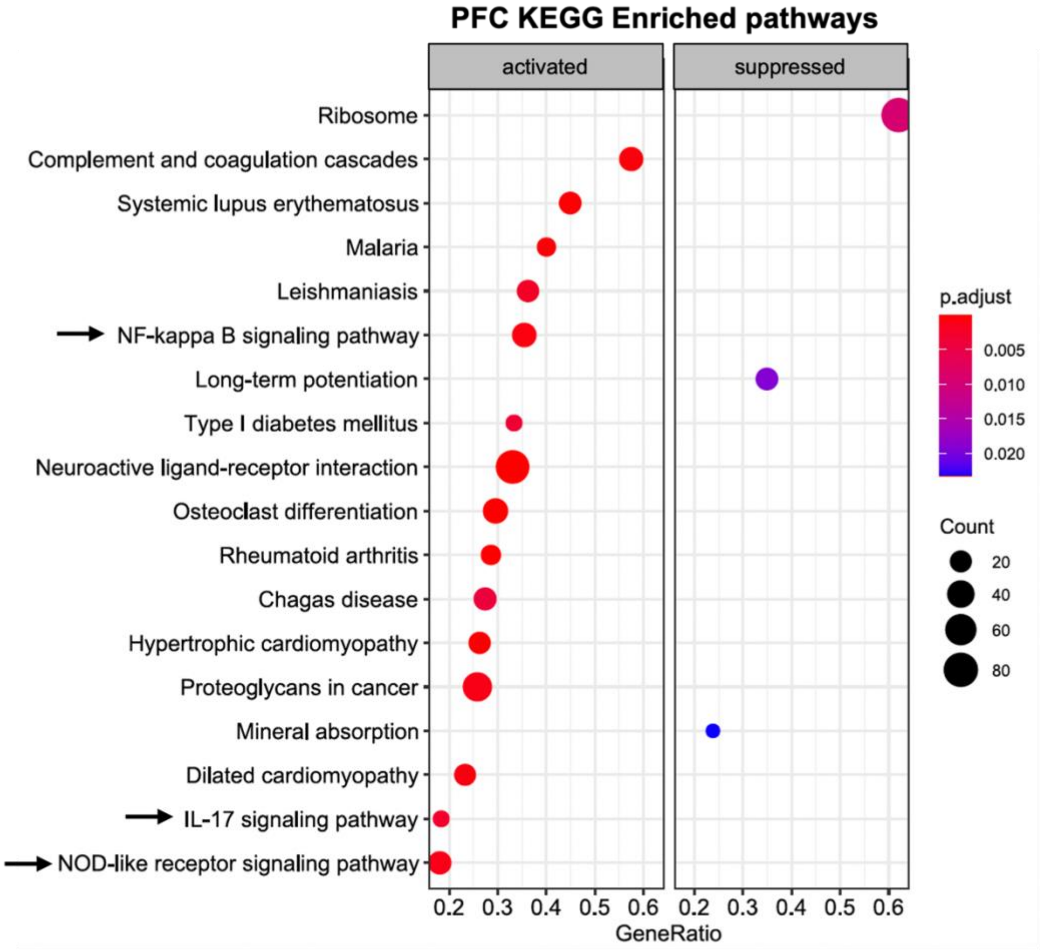


**Supplementary Figure S13 | KEGG pathway enrichment analysis in PFC.** Bar graph showing enrichment of KEGG pathways in PFC samples. Black arrows indicate enrichment of inflammation bioprocesses. Color of the circle represents adjusted p-value and size represents number of genes enriched. Left panel shows activate and right panel shows suppressed KEGG pathways.
